# A combined microphysiological-computational omics approach in dietary protein evaluation

**DOI:** 10.1101/2020.07.03.184689

**Authors:** Paulus G.M. Jochems, Willem R. Keusters, Antoine H.P. America, Pascale C.S. Rietveld, Shanna Bastiaan-Net, Renata M.C. Ariëns, Monic M.M. Tomassen, Fraser Lewis, Yang Li, Koen G.C. Westphal, Johan Garssen, Harry J. Wichers, Jeroen van Bergenhenegouwen, Rosalinde Masereeuw

## Abstract

The ever-growing world population puts pressure on food security. To tackle this, waste stream proteins and novel protein sources need to be evaluated for nutritional value, which requires information on digesta peptide composition in comparison to established protein sources and coupling to biological parameters. Here, we present a novel combined experimental and computational approach comparing seventeen protein sources with cow’s whey protein (WPC) as benchmark. *In vitro* digestion was followed by proteomics analysis and statistical model clustering based on Bayesian Information Criterion. Next, we incorporated functional protein data after evaluating the effects of eighteen protein digests on intestinal barrier integrity, viability, brush border enzyme activity and immune parameters using a bioengineered intestine. Our data show that a holistic approach allows evaluating a dietary protein’s potential for delivery of bioactive peptides, where protein source (animal, plant or novel source-derived) does not seem to be the driving force for clustering.

---

The ever-expanding world population goes hand in hand with an increase in food demand (1). One of the primary components of the human diet are proteins. In contrast to other macronutrients, proteins’ primary function is not to provide energy, but proteins are involved in almost all biological processes (2). Re-evaluating the over 90 million tons of food waste per year and exploring other protein sources may aid in identifying potential sustainable sources to assist in strengthening food security (3). A relevant example is whey; first considered a waste product during cheese production, now one of the most abundant proteins in the modern Western diet. In 2013, global whey ingredient exports were estimated 1.5 million metric tons/yr (4). Over the years, whey has been studied extensively and was found to contain a complex mixture of bioactive peptides (5), involved in *e.g.* immune modulation, maintaining intestinal barrier integrity and regulation of satiety (6, 7). Nutritional data on protein evaluation have been collected in human clinical trials, which are, however, associated with high costs, allow a limited number of samples to be tested, deliver limited mechanistic data, are not easy to control, leading to confounding outcomes and require ethical approval. *In vitro* models are cheaper, less time consuming and suited for medium to high throughput screening (8). These models could be an important supportive tool to (pre)select proteins for inclusion in a clinical trial. The first relevant model to evaluate would be a prototype of the gastrointestinal tract, where digestion takes place, and the small intestine in particular, as this is the primary site of nutrient absorption. The small intestine has many key functions in addition to selective absorption via its epithelial barrier, *e.g.* finalizing the enzymatic digestion by the brush border enzymes and a first line of defense against pathogens (9). Intestinal permeability is associated with both local (*e.g.* inflammatory bowel disease) and systemic (*e.g.* Parkinson’s) diseases (10, 11). To gain insights in these intestinal physiological processes, novel, more complex, *in vitro* models have emerged as advanced technologies with the potential to improve *in vitro-in vivo* translation (12).

Here, bioengineered intestinal tubules were used as microphysiological gut that were exposed to *in vitro* digested proteins followed by evaluation of key physiological parameters (13). Three commonly used proteins sources (acidic WPC, egg, soy) and fourteen potential dietary proteins from a variety of sustainable sources (bovine blood plasma, insect (mealworm), pea, corn, wheat, fungi, yeast, potato) were studied and compared to a benchmark whey protein concentrate (WPC). After static *in vitro* digestion, the digestive profile was evaluated via proteomics to determine the degree of overlap with WPC. Next, the digests were assessed for biological effects, *viz.* intestinal epithelial integrity, brush border enzyme activity, cell viability, cytokine and chemokine release. These data were evaluated in a statistical cluster analysis using the Bayesian Information Criterion (BIC), enabling to evaluate potential dietary protein in a holistic manner.

## Results and Discussion

### Proteomic data provides insights on dietary protein content and complexity

Details on the protein sources studied are given in Table 1. AWPC, a whey protein concentrate, is a byproduct of acidic dairy foods (14). Due to the proteolysis by rennet, a complex set of enzymes, AWPC has a lower protein content in comparison to WPC (14). The next commonly used protein is egg white (Egg), consisting mainly of water and approx. 11% of protein (77% ovalbumin, ovotransferin and ovomucoid) (15). The third frequently used protein is soy, often used to replace animal-source proteins, but considered a less ideal protein for consumption due to its low methionine content (16, 17). In addition to these three commonly used proteins, we compared animal-based proteins from bovine blood plasma, plant-derived proteins from pea, wheat, potato and corn, representing major food crops worldwide, and the novel-source proteins derived from insect, fungi and yeast (18).

**Table 1.**
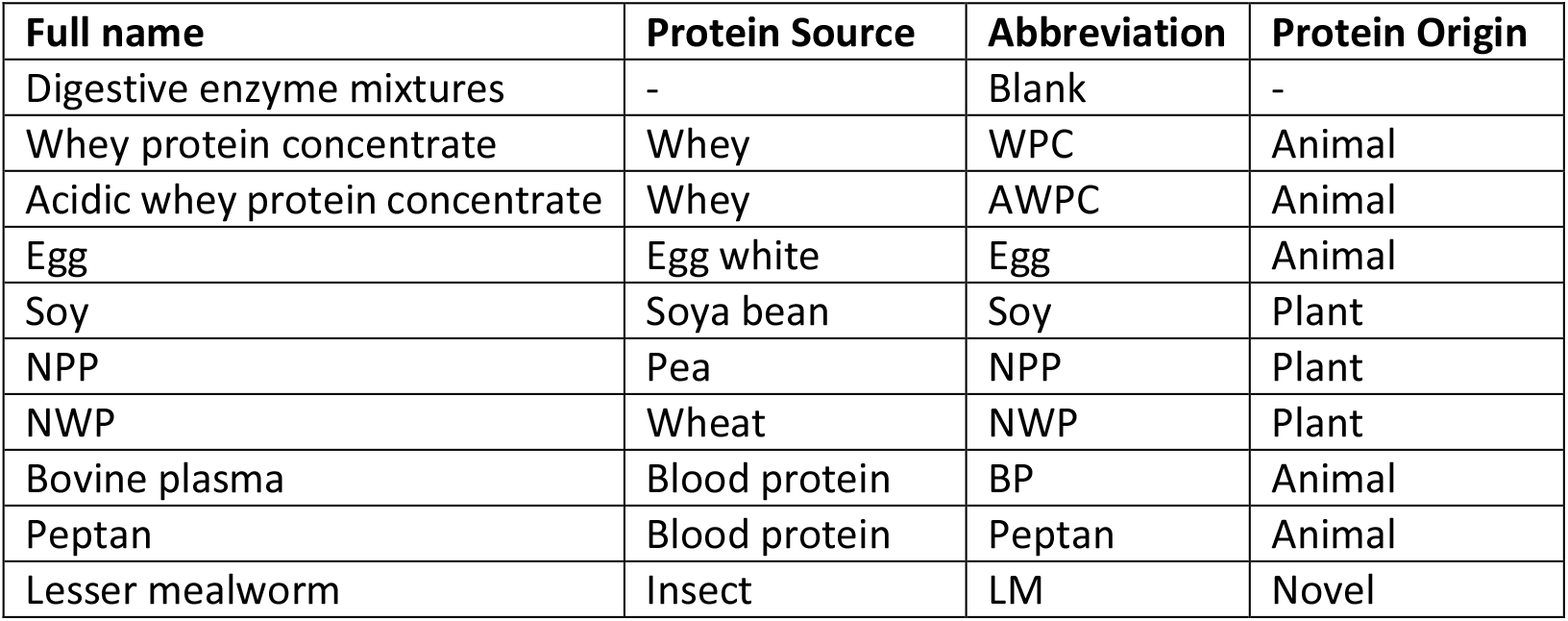
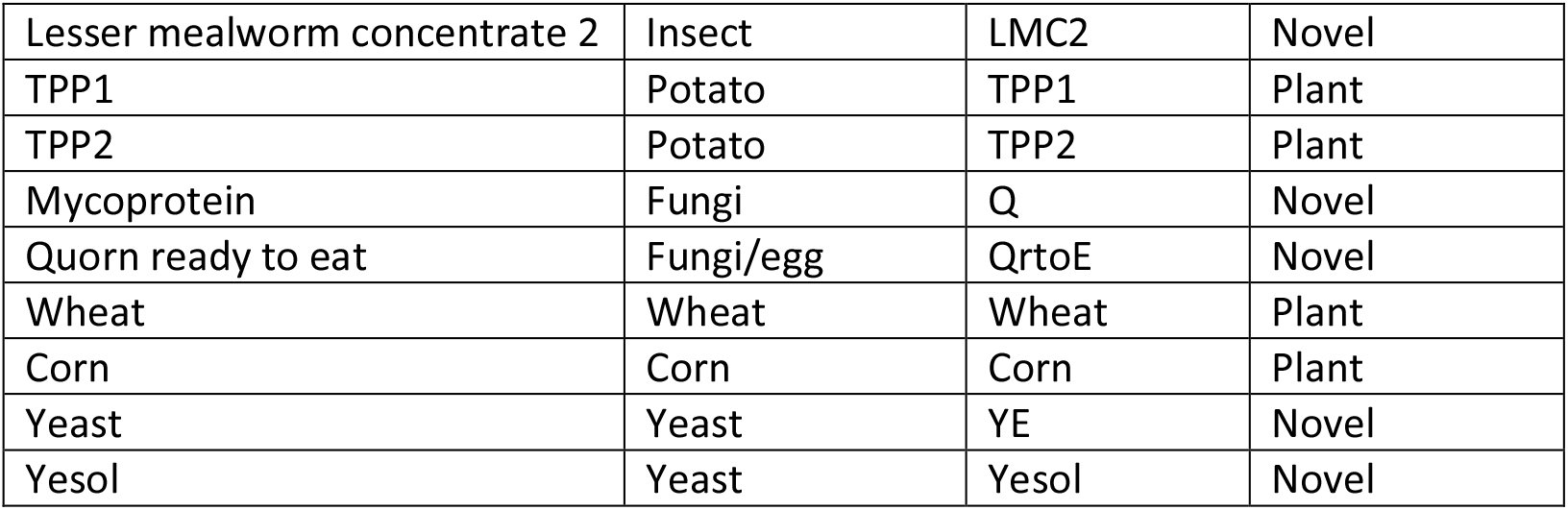
Overvi ew of the 18 dietary proteins evaluated.

After ingestion, protein digestion immediately starts and continues along the complete gastro-intestinal tract. *In vivo*, peptides from dietary proteins can be formed through hydrolysis by enzymes released from the matrix itself or contaminating microbes, gastrointestinal digestive enzyme hydrolysis and/or microbial fermentation (19). Important to note is that during static *in vitro* digestion only gastrointestinal enzymes are taken into account (without brush border enzymes), present at a total protein concentration of 2.9 – 14.8 mg/mL (avg. 9.4 mg/mL) (Suppl. Table 1).

Currently, dietary protein quality is primarily determined by presence and level of essential amino acids for humans and its digestibility, absorbability and utilization for metabolic functions (20). Here, the digestibility was examined by a proteome analysis of the digested peptide mixture. Commonly, online databases are analyzed to correlate data on digest composition to bioactivity of peptides (21, 22). Based on homology analysis, functions of unknown peptide sequences can then be predicted; however, actual biological efficacy remains to be validated. A limitation of this approach is that these databases hold data on a limited number of bioactive protein sequences from known sources. Our approach differs in that we compared complete peptide ion profiles generated by LC-MS. Comparison of novel protein sources peptides with a well-studied protein source, like WPC, will result in increased understanding of the digestive profile and the potential to induce health benefits of the novel protein hydrolysate. After enzymatic digestion and bioseperation, LC-MS ion-peak data of the filtrate, containing peptides to be taken up in the small intestine, were used to compare WPC and the other dietary proteins based on relative molar mass (m/z), charge, retention time and abundance. Each protein source was evaluated for the following characteristics: a) fraction of protein digestive products (dividing di-, tri- and oligopeptides abundancy), b) total peptide abundance (ranging from 0.12-1.99*10^8^), c) unique peptides, *i.e.* non-overlapping with WPC (ranging from 65-436 peptide ions), d) WPC peptide overlap relative to total number of peptides (ranging from 7.8-82.6%), and e) summed peptide abundance of WPC overlapping peptides relative to total peptide abundance (ranging from 5.7-95.4%) (Suppl. Fig. 1). The overlap of peptides of dietary protein digests versus WPC was visualized in tile plots (Suppl. Fig. 2), providing insights in the degree of overlap and peptide length. For each overlapping peptide, the abundance of that specific peptide was plotted in comparison to the WPC abundance (Suppl. Fig. 2). For a holistic evaluation of the dietary proteins in comparison to WPC, these data were used for a statistical cluster analysis and centered around zero proteomic data (absolute values are shown in Suppl. Fig. 1). Proteomic data were clustered using the Bayesian Information Criterion (BIC), a standard goodness of fit metric, used here to determine the optimal clustering model supported by the data. Fourteen different clustering models were compared resulting in ellipsoidal, equal volume, shape and orientation (EEE) model with 8 distinct clusters as best fit (BIC results Suppl. Fig. 3) (23).

All combinations of two components plots were plotted (Suppl. Fig. 4), of which two examples are shown in Fig. 1. As expected, AWPC protein content showed great similarity with WPC as confirmed by its direct clustering (Fig. 1, cluster 1). Both animal and plant-sourced proteins, respectively BP, Soy and TPP1 clustered together (Fig. 1, cluster 2). This cluster is primarily driven by the similarity in ratios between protein digestive products. Total peptide abundance in Corn is similar as in WPC, however, its limited overlap with WPC peptides resulted in a separate cluster (Fig. 1, cluster 3).

**Figure 1.**
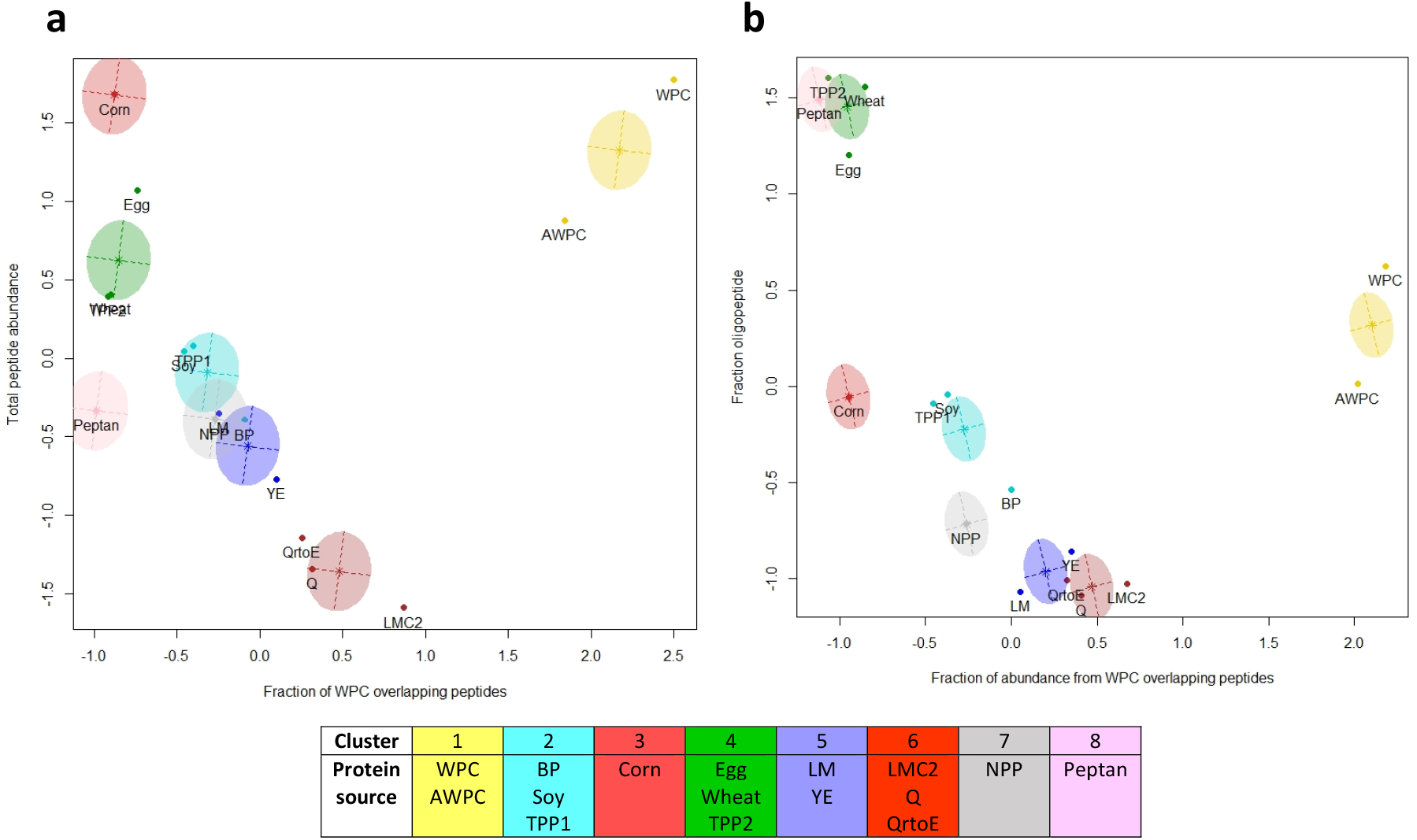
Two-component plot of proteomic data clustering potential dietary proteins. The optimal cluster method (EEE) and cluster quantity (8) were determined based on Bayesian Information Criterion. A) This two-component plot shows peptide mass overlap relative to total peptide abundance on the vertical axis and relative peptide overlap to total peptide count in WPC on the horizontal axis. B) Two-component plot showing fraction oligopeptides on the vertical axis and relative peptide abundance of WPC overlapping peptides on the horizontal axis. Supplementary figure 4 shows all combinations of two-component plots. Analysis was done based on digest peptide profiles, ion peak overlap, quantity of WPC-matched ion peaks relative to total protein ion peaks and relative abundance of WPC-matched ion peaks.

Cluster 4 contains proteins from Egg, TPP2 and Wheat, again comprising both animal and plant derived proteins (Fig.1, cluster 4), and characterized by a limited peptide mass overlap in comparison to WPC while having a high number of unique peptides and a distinct relative peptide abundance. LM (insect) and YE (yeast), both considered novel-protein sources, clustered together (Fig. 1, cluster 5) based on their average peptide abundance and fraction of peptide digestive products. Even though novel-sourced proteins, LMC2, Q and QrtoE, showed similar digestive fractioning, their low number of unique peptides, peptide abundance and high degree of WPC overlap resulted in a separate clustering (Fig. 1, cluster 6). NPP clustered by itself (Fig.1, cluster 7), due to its high tri/oligopeptide fraction in combination with an average peptide abundance. Peptan formed the final cluster (Fig.1, cluster 8) most likely due to its low level of peptide overlap with WPC in comparison with other dietary proteins while having an average peptide abundance (Fig.1, cluster 8).

Putting the proteomic data clustering in perspective, digestibility and absorbability insights are created by the evaluation of digestive peptide fractions. Hence, di- and tripeptide uptake is most efficient, as the small intestine expresses four apical peptide transporters allowing active absorption (2). Cluster 5 and 6, containing only novel-source proteins (insect, fungi and yeast) showed greatest fractions of di- and tripeptides suggestive for a favorable absorbability. On the other hand, cluster 4 and 8, a combination of plant and animal-based proteins, showed a high oligopeptide content suggestive for a lower degree of absorbability. A high protein abundance and associated peptide content suggest a more complex digestion product. In general, plant food contains less protein in comparison to animal matter (24). However, extracts used were enriched in protein and similar amounts of protein were used per digest (except for TPP1 and TPP2 that contained extra starch). Cluster 1 and 3 are composed of plant and animal-source dietary proteins showing high levels of peptide content. Cluster 6 containing novel-source proteins had a low peptide content. To further compare the concentration of overlapping peptides with WPC, their abundance was plotted for each overlapping peptide (Suppl. Fig. 2). Identification of native digests peptide sequences is not trivial, as protein database search algorithms are mostly developed for longer peptide sequences, and preferentially use a minimum sequence length of 5 or 6 amino acids (25). As a proof of concept, WPC peptides that showed overlap with another dietary protein were, if possible, linked to a specific peptide sequence using *de novo* MS/MS spectrum interpretation (26) (Table 2).

**Table 2.** Identified peptide sequences with at least a *de novo* score of >90.

| Peptide sequence | #ID | Presence in proteins |
| --- | --- | --- |
| Leu-Lys (LK) | 33 | AWPC, Egg, Soy, NPP, BP, LM, QrtoE, TPP1, TPP2, YE |
| Asp-Lys (DK) | 37 | AWPC, Egg, Q, QrtoE, Corn |
| Val-Arg (VR) | 43 | AWPC, Egg |
| Met-Lys (MK) | 50 | AWPC, Soy, NPP, BP, LM, TPP1, Corn, Wheat, YE |
| Met-Ala-Pro-Lys (MAPK) | 128 | AWPC |

Important to note is that the *de novo* identification method is not capable of distinguishing isomeric amino acids (leucine and isoleucine) (27, 28). In total 323 peptides sequences were identified with a range of 0 - 98.1 for the *de novo* sequence score. To ensure a reliable identification, only *de novo* sequence scores >90 are discussed (Table 2; abundance shown in Suppl. Fig. 5). First, sequences LK, MK and MAPK form parts of known antihypertensive peptides LKP (β-lactoglobulin), MKP (α_s2_-casein f) and MAP (β-casein f) present in whey protein (29–31). Furthermore, VR has been identified as peptide sequence with angiotensin-converting-enzyme inhibitory capacity (32). Hence, these peptide sequences might have positive cardiovascular health effects by inhibiting angiotensin I conversion into angiotensin II, preventing vasoconstrictive effects (33). Before dietary peptides can induce systemic health effects, these must be absorbed in the small intestine. Therefore, their biological efficacy was tested using bioengineered intestinal tubules (Suppl. Fig. 6).

### Cluster analysis distinguishes biological behavior of potential dietary proteins

Protein sources are inherently contaminated with microbial products such as endotoxins, of which translocation over the epithelial barrier can initiate a systemic immune response (34). Nevertheless, the endotoxin contamination in human nutrition is not subject of legislation. In addition, the small intestine is continuously exposed to endotoxins originating from its own microbiome and acquires tolerance immediately after birth by *e.g.* downregulating interleukin 1 receptor-associated kinase 1 (35). Endotoxin contamination in *in vitro* digestion used chemicals, resulted in increased endotoxin contamination in the protein digests, varying in this study between 10.4 - >65.0*10^3^ EU/mL (Suppl. Table 1). However, blank digest showed no effects on the evaluated small intestinal features (Suppl. Fig. 7), providing evidence that the enzymes and associated endotoxin content had no detrimental effects on the epithelial barrier integrity, cell viability and alkaline phosphatase activity in our bioengineered intestinal tubules.

Next, experiment-ready bioengineered intestinal tubules were exposed to dietary protein digests and impact on epithelial barrier integrity, brush border enzyme activity and cell viability were assessed (Suppl. Fig. 7). The biological data set was further supplemented by dietary protein-induced release of immune markers (*viz.* IL-6, TGFβ and NO) (Suppl. Fig. 8). The complete set of biological read-outs were clustered using the Bayesian Information Criterion (BIC), a standard goodness of fit metric, used here to determine the optimal clustering model supported by the data. This resulted in a diagonal equal shape and volume (EEI) model with 6 distinct clusters as best fit (Suppl. Fig. 9) (23). All combinations of two components plots were plotted (Suppl. Fig. 10), of which two examples are shown in Fig. 2.

**Figure 2.**
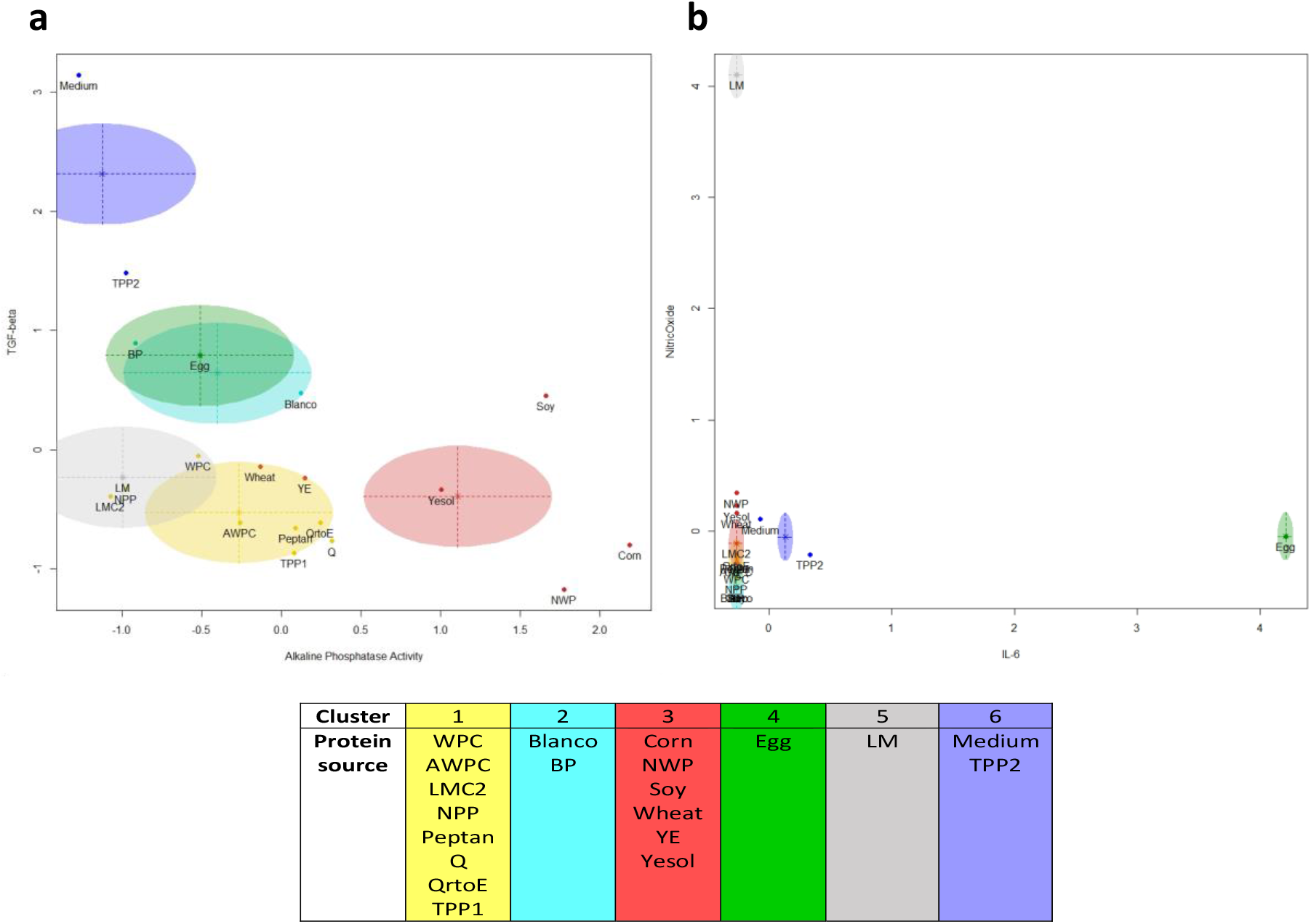
Two-component plot with 6 distinct protein clusters based on bioengineered intestinal tubule-acquired biological efficiacy data. The optimal cluster model (EEE) and cluster quantity (6) were determined based on Bayesian Information Criterion. A) Two-component plot showing TGFβ secretion (vertical axis) and alkaline phosphatase activity (horizontal axis). B) two-component plot showing nitric oxide (vertical axis) and IL-6 (horizontal axis). Suppl. Fig. 9 shows all combinations of two-component plots. Analysis was done based on inulin-FITC leakage, zonula occludens-1 (ZO-1) expression quantification, tissue viability, alkaline phosphatase activity, IL-6 secretion, TGF-β secretion and NO production.

In a subsequent analysis, dietary proteins were compared to the reference protein WPC, which showed no significant difference for most parameters tested (Suppl. Fig. 7). However, analysis of the biological data using a non-biased cluster-based approach demonstrated apparent differences. The first and largest cluster contains WPC together proteins originating from animal (WPC, AWPC, Peptan), plant (NPP and TPP1) and novel sources (LMC2, Q and QrtoE) (Fig.2, cluster 1). These proteins showed average levels of intestinal physiological parameters and low levels of cytokine and chemokine secretion/production. Cluster 2 includes the digestion mixture (blank digest) and BP, primarily based on their relative high levels of ZO-1 expression (Fig.2, cluster 2). The blank digest did significantly decrease TGFβ and NO secretion, as reported earlier for TGFβ, nitrate and nitrite (as measure for NO) (36–38). Corn, NWP, Soy, and Wheat (plant-source) next to YE and Yesol (novel-source) form the second largest cluster (Fig.2, cluster 3). Egg forms its own separate cluster because it significantly induces IL-6 secretion (Fig.2, cluster 4). Next to Egg, only medium and TPP2 also induced IL-6 secretion, however, these levels were not sufficient to allow for clustering together with Egg, and other variables accounted for the formation of a separate clustering (Fig. 2, cluster 6). Even though IL-6 concentrations were increased, these remained under an epithelial barrier disruptive concentration (1 ng/mL), confirmed by intestinal epithelial integrity assays (39). However, IL-6 itself can induce intestinal inflammation by provoking a T-cell response and IgG secretion by B-cells, and is associated with a variety of local and systemic inflammatory disease (40, 41). The capacity to induce an immune response by the increased IL-6 levels upon Egg and TPP2 digest exposure requires further research. LM clustered separately, primarily due to its induction of high levels of NO (Fig.2, cluster 5; Suppl. Fig. 8B), although these levels (avg. 155.4 μM) did not affect the epithelial barrier (Suppl. Fig. 7A and 7B). NO is able to induce beneficial regulatory effects for essential intestinal features, such as maintaining an intestinal epithelial barrier, but increased levels are associated with pathology in *e.g.* inflammatory bowel disease (42, 43). Its dual effect *in vivo* challenges translational interpretation and warrants further research focusing on the downstream effects of increased NO levels upon LM digest exposures.

### Statistical cluster analysis based on the proteome and biological efficacy reveals that protein source is not a driving factor for clustering

To create a holistic overview, both the proteomic and biological efficacy data were combined in a cluster analysis. Based on the BIC, an ellipsoidal equal volume, shape and orientation (EEE) statistical cluster model with 5 distinct clusters had a best fit (Suppl. Fig. 11) (23). All combinations of two components plots were plotted (Suppl. Fig. 12), of which two examples are shown in Fig. 3.

**Figure 3.**
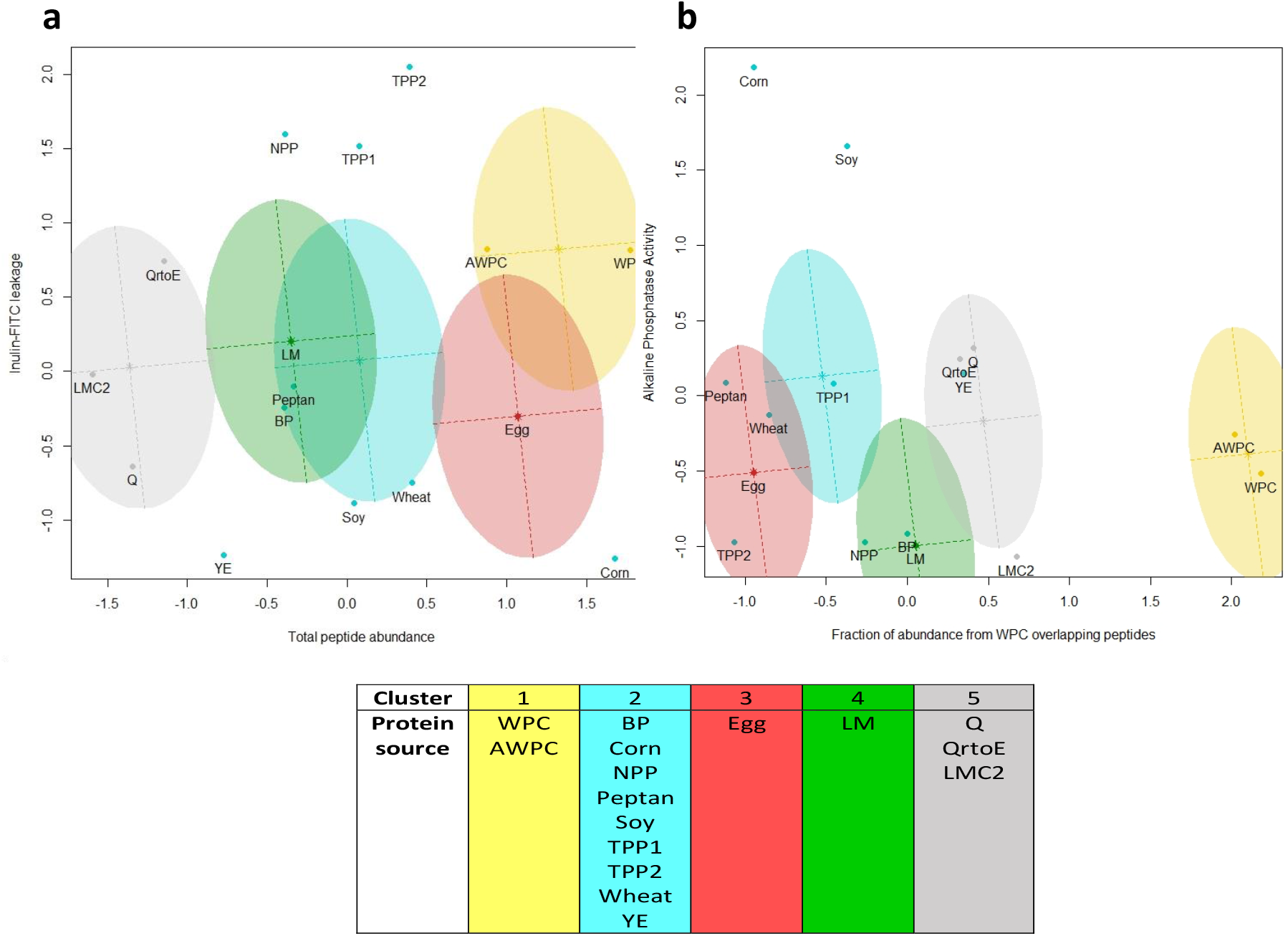
Two-component plot of proteomic data and *in vitro* biological efficacy data clustering potential dietary proteins. The optimal statistical cluster model was ellipsoidal, equal volume, shape and orientation (EEE), and cluster quantity (5) was determined based on Bayesian Information Criterion. Considered data were peptide abundance, total amount of unique peptides, relative overlap with whey protein concentrate (WPC), the relative peptide abundance of the WPC overlap, inulin-FITC leakage, ZO-1 quantification, cell viability, alkaline phosphatase activity, IL-6, TGFβ and nitric oxide. A) two-component plot showing inulin-FITC leakage assay on the vertical axis and total peptide abundance on the horizontal axis. B) two-component plot showing alkaline phosphatase activity assay on the vertical axis and relative peptide abundance of WPC overlapping peptides on the horizontal axis. Supplementary figure 11 shows all combinations of two-component plots. The division of potential dietary proteins in the different clusters is shown below the two two-component plots. Analysis was done based on proteomic data and biological efficacy data.

Important to note is that emphasis during this clustering was placed automatically on biological efficacy as these parameters outnumbered proteomic parameters in a 2:1 ratio. Placement of WPC and AWPC (Fig. 3, cluster 1) together is thought to be primarily driven by the high level of peptide overlap in proteomic data, as no significant differences in biological efficacy were detected. This underscores the potential of AWPC as alternative to WPC in providing dietary proteins. However, the production process still requires optimization (14). The largest cluster consists of animal-based protein sources, BP and Peptan, plant-based sources, Corn, NPP, Soy, TPP1, TPP2 and Wheat, and novel protein source, YE (Fig.3, cluster 2). Again, the protein source does not seem to be the driving force for clustering of proteomic data nor biological efficacy. Interestingly, a wide variety of protein-sources clustered together with the common animal-protein replacer Soy. Hence, these plant-based and novel protein sources potentially could replace soy. The separate clusters of Egg (Fig.3, cluster 3) and LM (Fig.3, cluster 4) are driven by their significant induction of IL-6 and NO, respectively. Q, QrtoE and LMC2 did not show much alterations on biological efficacy rand are grouped based on their relatively small peptide content resulting in a high degree of WPC overlap (Fig.3, cluster 5). The statistical model clustering based on proteomics and *in vitro* biological efficacy showed sensitivity for both datasets, as also clusters driven by either proteomics or *in vitro* biological efficacy appeared confirming the validity of our approach.

## Conclusion

In the present study, we combined a microphysiological intestinal evaluation of eighteen dietary protein sources with their digests using a computational approach. To create a holistic characterization of novel dietary proteins, proteomic and *in vitro* biological efficacy data were used for a statistical model clustering. The combinational clustering remained sensitive for proteomic and biological efficacy data as specific clusters could be driven by either one of them. As little is known with respect to novel protein sources, a well-characterized reference protein is instrumental for evaluation. Proteomic analysis focused on global characteristics (peptide length, abundance and distribution) rather than individual peptide sequence, therefore, *de novo* interpretation of MSMS spectra provides a valid alternative approach. However, depends on sufficient fragmentation information available in the MSMS spectra, which is not available for all peptide ions.

As limited information on the biological impact of novel source proteins is available, our approach increases the chance of identifying specific protein sources to support a targeted health benefit discovery. Overall, our holistic approach of clustering based on both proteomic and biological efficacy shows great potential as a first evaluation step in the analysis of novel dietary proteins. Moreover, this original approach showed that protein source is not a key factor in determining the holistic clustering as different sources clustered together based on digestive profile and/or biological efficacy of a (novel) protein.

## Methods

### Chemicals

All chemicals were purchased from Sigma-Aldrich (Zwijndrecht, The Netherlands) unless stated otherwise.

### Dietary proteins

A total of 18 dietary proteins were *in vitro* digested and evaluated. WPC was used as primary reference protein for 17 sustainable other dietary proteins. These sustainable proteins had a wide variety of protein source (table 1).

### Static in vitro digestion

The static *in vitro* digestions were performed according to the widely accepted INFOGEST consensus protocol (44). To allow use of the digestions in cell culture models, the pancreatin enzyme mixture was adjusted to a Trypsin activity at 10 U/ml. After the intestinal incubation phase, samples were aliquoted and snap frozen in liquid nitrogen and stored at −80 for subsequent measurements. Protein content in the digests as assessed in triplicate on a Flash EA 1112 GC (Interscience BV, Breda, The Netherlands) according the Dumas method (45). Methionine and cellulose were used as a standard and negative control respectively. Endotoxin content was determined by the EndoZyme II kit according to manufacturer’s instructions (Biomérieux company, München, Germany). Results were considered valid if the endotoxin spike recovery was in the range of 50 % to 200 %. Fluorescence readings were performed on an Infinite F200 plate reader (Tecan Group Ltd., Männedorf, Switzerland). When samples displayed fluorescence overflow values, the endotoxin content was set to > 65000 EU/ml.Next, digested samples were used for proteomics or exposure to the bioengineered intestinal tubules.

### Proteomics

Proteomic analysis has only been conducted on the small intestine absorbable peptide fraction. To that end, the digests were separated by an Amicon^®^ Bioseparations Stirred Cells device over a 1kDa ultrafiltration disc (Merck Millipore; Amsterdam, The Netherlands) into a retentate and filtrate. *In vitro* protein digests were analyzed on a UPLC-MS system (Dionex Ultimate 3000 online connected to Qexactive^PLUS^ (Thermofisher, Waltham, U.S.A). Samples were injected (10microliter) on a pentafluorophenyl F5 core-shell column (Kinetex F5, 15cm *2.1mm, 2.6 micrometer particles, Phenomenex, Torrance, USA) operated at 30°C and flow rate of 0.2 ml per minute. Total run time was developed over a 40 minutes time window: Starting with buffer A (0.1% formic acid in water) for 5 minutes and separated with a gradient of 0% to 30% buffer B (0.1% formic acid in 100% Acetonitrile) during 20 minutes, increasing to 80% B in 5 minutes, stable at 80% B for 3 minutes and back to 0% B during 2 minutes, and stable at 0% B for 5 minutes. Separated peptides were on-line injected into the Qexactive^PLUS^ using the standard ESI source in positive mode, with 3.5kV spray voltage, 290oC capillary temperature, nitrogen sheath gas flow 40 and auxillary gas flow 10 heated at 60oC. MS spectra were collected with alternating scans; first within a m/z range of 70-380, followed by MS scan within a range of 350-1200 m/z at 70000 resolution (profile) and AGC target of 3*10e6 ions maxIT for 100 milliseconds; followed with data-dependent switch to MSMS mode at 17500 resolution (centroid) at AGC target 10e5 ions, minimum AGC 8*10e3, maxIT 50 millliseconds and 4 m/z isolation window, loop count 5, a dynamic exclusion of 10 seconds without further charge exclusion.

### In vitro biological efficacy

#### Caco-2 cell culture

The human colon adenocarcinoma-derived intestinal cell line, Caco-2, (ATCC, Wesel, Germany) were maintained in high glucose Dulbecco’s Modified Eagle Medium – high glucose (Gibco) supplemented with fetal calf serum (10% v/v) and penicillin and streptomycin (1% v/v). Media was refreshed every 2-3 days, and cells were passaged and seeded on bioengineered intestinal tubules when reaching 80-90% confluency.

#### Bioengineered intestinal tubule construction, coating, seeding and cultivation

Three-dimensional polylactide chambers (schematic illustration in Suppl. Fig. 13) were printed with the Ultimaker 3 (Ultimaker, Geldermalsen, The Netherlands). Thereafter, SENUOfil type H-MF-0,2 hollow fiber capillary membranes (SENUOFIL, Tianjin, China) were cut and guided through the chamber using a steel wire. 18g blunt needles 0.5-inch length (OctoInkjet, Hoyland, United Kingdom) were put in the in- and outlet of the chambers and made leak tight using Loctite EA M-31CL glue (Henkel Adhesives, Nieuwegein, The Netherlands). Finally, the bottom of the chamber was sealed with a 24 x 60mm glass cover slip (Menzel-Gläser, Braunschweig, Germany) using Loctite EA M-31CL glue. Glue dried for 24h and GI-MASK Automix glue (Coltene, Lezennes, France) was used to make the inner parts of both syringes leak tight.

After chamber construction, extra cellular matrix (ECM) coating and cell seeding were done as described before with minor changes (13). Here, coating was directly executed in the 3D-chambers, thus 90° turning was excluded from the protocol. In short, 30min of sterilization in 70% (v/v) EtOH followed by 5h filter-sterilized L-3,4-di-hydroxy-phenylalanine (L-Dopa, 2 mg.mL^−1^ in 10 mM Tris buffer, pH 8.5) followed by 2h human collagen IV solution (25 μg.mL^−1^ in PBS) to complete the ECM-coating. Caco-2 cells were trypsinized and seeded in the chambers for 4h at a seeding density of 1*10^6^ Caco-2 cells/bioengineered intestinal tubule.

Bioengineered intestinal tubules were cultivated for 21 days in a 5% CO_2_ and 37°C incubator, and culture medium was refreshed every 2-3 days. During the final 7 days, chambers were put on a 2-dimensional rocking platform (VWR, Breda, The Netherlands) with a speed rate of 1 rotation per minute at an angle of 10°. Thereafter, bioengineered intestinal tubules were experiment ready.

#### Protein digest exposures

Protein digests were diluted 1:4 with culture medium and bioengineered intestinal tubules were exposed to 1mL for 3h. Thereafter, bioengineered intestinal tubule were assessed on barrier integrity, brush border enzyme activity, epithelial cell viability and secretory markers.

#### Inulin-FITC leakage assay

To quantify the paracellular permeability, bioengineered intestinal tubules were perfused with inulin-FITC (0.1 mg.ml^−1^ in PBS). Prior to perfusion, bioengineered intestinal tubules were washed and placed in PBS. Then, chambers were connected to a Reglo Independent Channel Control pump (Ismatec, Wertheim, Germany) and perfused with inulin-FITC for 10 min at a flow rate of 0.1 mL/min. PBS from the chamber was collected, homogenized and fluorescence was measured at excitation wavelength of 492 nm and emission wavelength of 518 nm using Tecan infinite M200PRO plate reader (Tecan Austria GmbH). Results are shown relative to medium control. After the leakage assay, bioengineered intestinal tubules were washed in wash buffer (4% FCS in HBSS (v/v)) and cut in three pieces of which one was used for immunostaining and two for cell viability and alkaline phosphate activity.

#### Immunofluorescent staining and zonula occludens-1 quantification

Staining protocol for bioengineered intestinal tubules was described previously (13). In short, cells were fixed (60% EtOH, 30% chloroform and 10% acetic acid (v/v)) for 5minutes, permeabilized (0.3% (v/v) Trition X-100 in HBSS) for 10min and blocked ((2% (w/v) bovine serium albumin (BSA) fraction V and 0.1% (v/v) Tween-20 in HBSS) for 30min. Thereafter, bioengineered intestinal tubules were incubated with zonula occludens-1 (ZO-1) antibody (1:1000 diluted in blocking solution) (Thermo Fisher Scientific, Bleiswijk, The Netherland) for 2h. After washing, secondary antibody goat-anti-rabbit 594 (1:200) (Abcam, Cambridge, United Kingdom) was added. Finally, bioengineered intestinal tubules were mounted using Prolong gold containing DAPI (Cell signaling technology, Leiden, The Netherlands) for nuclei staining. Images were acquired using the Leica TCS SP8 X (Leica Biosystems, Amsterdam, The Netherlands).

For image analyses Fuiji ImageJ was used and a full field of view image was required (Suppl. File 2 provides a step by step protocol, including representative bioengineered intestinal tubules images). Maximum intensity projection were made of the bioengineered intestinal tubules, channels were separated. ZO-1 intersections were determined using a ROI manager, lines were drawn over the area of interest. Values were corrected for background and the amount of intersections per line were counted, providing a quantitative measure for ZO-1. The number of nuclei was counted via a particle size measurement. Thereafter, each value was corrected for surface area expressed in micron^2^.

#### Cell viability

As a measure for cell viability, the mitochondrial activity was measured via PrestoBlueTM cell viability reagent assay (Thermo Fisher). PrestoBlue^TM^ reagent was mixed with culture medium at a 1:10 ratio and 100μL was put on a part of the bioengineered intestinal tubules in a 96 wells plate. The plate was placed in an incubator at 5% CO_2_ and 37°C and incubated for 1h protected from light. Bioengineered intestinal tubules were removed and fluorescence was measured at excitation wavelength of 530 nm and emission wavelength of 590 nm using Tecan infinite M200PRO plate reader. Positive control, unexposed bioengineered intestinal tubule, was set to 100% viability.

#### Brush border enzyme activity

After the cell viability assay, bioengineered tubules were washed with PBS and alkaline phosphatase activity was measured using Amplite^TM^ Colorimetric Alkaline Phosphatase Assay kit (AAT Bioquest, Sunnyvale, United States). Assay was performed according manufacturer protocol. Bioengineered intestinal tubules were exposed to reaction mixture for 30 min and absorbance was measured at 600 nm using colorimetric plate reader (iMARK™ microplate absorbance reader, Bio-Rad). Values are shown relative to medium exposed.

#### Immune response

After exposures, supernatant was collected and IL-8 (BioLegend, London, United Kingdom), IL-6 (Biolegend) and TGF-β (Biolegend) were quantified via ELISA. ELISAs were executed according manufacturer protocol. First, plates were coated and incubated overnight. Thereafter, plates were blocked for 1h and incubated with samples for 2h. This was followed by detection antibody incubation for 1h and Avidin-HRP for 30min. Wells were exposed for 15min to substrate solution followed by stop solution. Absorbance was read at 450nm. Wells were washed in between exposures. Values were corrected for background using the supernatant of the unseeded no-ECM coated bioengineered intestinal tubules, specific for each data set.

NO was determined via Griess reaction (Promega, Leiden, The Netherlands) according manufacturer protocol. In short, samples were centrifuged for 3min at 6000rpm and supernatants were transferred to a 96wells plate. Sulfanilamide solution was added to all wells and incubated for 10min. Thereafter, NED solution was added and incubated for 10min. Followed by an absorbance measurement at 490nm.

### Data processing and dietary protein clustering

#### In vitro biological data

Every experiment was at least performed in triplicate. Data was analyzed for outliers using Grubbs test, α=0.05. Statistical analysis was performed in Graphpad version 8 using t-test and one-way ANOVA. A P-value of <0.05 was considered significant.

#### Proteomic analysis

LC-MSMS data were loaded per replicate of protein source sample into Progenesis QI software (Non-linear Dynamics, Waters BV, UK). Peak detection was performed with a sensitivity setting of 4 and minimum peak length of 2 seconds. Peptide ion data were exported to a comma separated file per replicate of protein source sample. Individual peptide ion tables were further filtered and processed in Excel and R-studio.

A total of 422697 ion-peaks (over all peptide ion tables) were observed and corrected for background noise. First, data with a raw abundance <1*10^5^ was excluded, excluding 414436 ion-peaks. Second, all dietary protein digesta were corrected for digestive enzymes by excluding overlapping ion peaks with the blank. Criteria for matching ion peaks between separate samples was based on absolute differences in charge, m/z (±0.003) and retention time (±0.5min). Matching to Blank, resulted in 3542 ion-peaks exclusion, maintaining a total of 4719 ion-peaks divided over the eighteen protein digests for further analysis.

All protein digesta were matched to WPC by the matching criteria. If an ion-peak matched to multiple WPC ion-peaks, the nearest ion-peak was defined based on the two-dimensional space regarding m/z and retention time, weighing both criteria equally (resulting in 0.003 m/z = 0.5 min retention time). The distance was measured using Pythagoras equation, as shown in the formula. The match with the smallest distance was then selected.

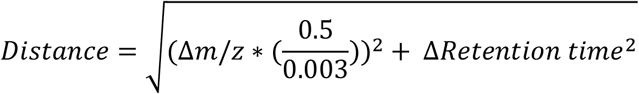

If a protein had multiple ion-peaks matched to the same ion-peak in WPC, their abundance was shown as total sum.

#### Computational statistical model clustering

Data of proteomics and *in vitro* biological efficacy were processed using R version 3.5.1 (packages data.table, ggplot2, Mclust & dplyr). Proteomic and *in vitro* biological efficacy data were used as mean per parameter and standardized and centered around zero. The Mclust package was used for an unbiased clustering method based on Bayesian Information Criterion resulting in a best fit model with preferred number of clustering based on the BIC score (R-script, Suppl. File 3). These clusters provide a 2-dimensional holistic representation of dietary protein properties.

## Abbreviations

WPC: whey protein concentrate
HFM: hollow fiber membranes
ZO-1: zonula occludens-1
ECM: extracellular matrix
L-Dopa: L-3,4-di-hydroxy-phenylalanine
BIC: Bayesian Information Criterion

## Acknowledgements

This work was supported by the Dutch Ministry of Economic Affairs program TKI-AF under grant 15269: Sustainable Future Proteins-Focus on nutritional and health promoting quality. The views expressed in this manuscript are those of the authors and do not necessarily reflect the position or policy of funding organizations: Cargill, La Saffre, PepsiCo, Inc., BASF, Mimetas, Proti-Farm, Danone Nutricia Research, Quorn Foods, Darling Ingredients, Roquette and AVEBE. Furthermore, we thank Karim El Bachrioui and Dianne B.P.M. van den Berg‐Somhorst for their assistance in the *in vitro* digestion experiments.

## Author contribution

PGMJ TA, PCSR, RMCA, MMMT and SBN conducted experimental work. PGMJ, WRK and LF conducted data analysis. PGMJ, JvB and RM wrote and revised the manuscript. All co-authors reviewed and revised the manuscript.

## References

1. Tilman D, Balzer C, Hill J, Befort BL. Global food demand and the sustainable intensification of agriculture. Proceedings of the National Academy of Sciences. 2011;108(50):20260–4.

2. Jochems P, Garssen J, van Keulen A, Masereeuw R, Jeurink P. Evaluating human intestinal cell lines for studying dietary protein absorption. Nutrients. 2018;10(3):322.

3. Ravindran R, Jaiswal AK. Exploitation of food industry waste for high-value products. Trends in Biotechnology. 2016;34(1):58–69.

4. Lagrange V, Whitsett D, Burris C. Global market for dairy proteins. Journal of food science. 2015;80(S1):A16–A22.

5. Walzem R, Dillard C, German JB. Whey components: millennia of evolution create functionalities for mammalian nutrition: what we know and what we may be overlooking. Critical reviews in food science and nutrition. 2002;42(4):353–75.

6. Keri Marshall N. Therapeutic applications of whey protein. Alternative medicine review. 2004;9(2):136–56.

7. Hering NA, Andres S, Fromm A, van Tol EA, Amasheh M, Mankertz J, et al. Transforming growth factor-β, a whey protein component, strengthens the intestinal barrier by upregulating claudin-4 in HT-29/B6 cells. The Journal of nutrition. 2011;141(5):783–9.

8. Jones D, Caballero S, Davidov-Pardo G. Bioavailability of nanotechnology-based bioactives and nutraceuticals. Advances in food and nutrition research. 2019;88:235–73.

9. Yamada T, Alpers DH, Kalloo AN, Kaplowitz N, Owyang C, Powell DW. Principles of clinical gastroenterology: Wiley-Blackwell; 2009.

10. Perez-Pardo P, Kliest T, Dodiya HB, Broersen LM, Garssen J, Keshavarzian A, et al. The gut-brain axis in Parkinson's disease: Possibilities for food-based therapies. European journal of pharmacology. 2017;817:86–95.

11. Lee SH. Intestinal permeability regulation by tight junction: implication on inflammatory bowel diseases. Intestinal research. 2015;13(1):11.

12. Esch EW, Bahinski A, Huh D. Organs-on-chips at the frontiers of drug discovery. Nature reviews Drug discovery. 2015;14(4):248–60.

13. Jochems PG, van Bergenhenegouwen J, van Genderen AM, Eis ST, Versprille LJW, Wichers HJ, et al. Development and validation of bioengineered intestinal tubules for translational research aimed at safety and efficacy testing of drugs and nutrients. Toxicology in Vitro. 2019;60:1–11.

14. Chandrapala J, Duke MC, Gray SR, Zisu B, Weeks M, Palmer M, et al. Properties of acid whey as a function of pH and temperature. Journal of Dairy Science. 2015;98(7):4352–63.

15. Mine Y. Recent advances in egg protein functionality in the food system. World's poultry science journal. 2002;58(1):31–9.

16. Friedman M, Brandon DL. Nutritional and health benefits of soy proteins. Journal of Agricultural and Food Chemistry. 2001;49(3):1069–86.

17. Montgomery KS. Soy protein. The Journal of perinatal education. 2003;12(3):42–5.

18. King JC, Slavin JL. White potatoes, human health, and dietary guidance. Advances in Nutrition. 2013;4(3):393S–401S.

19. Chakrabarti S, Guha S, Majumder K. Food-derived bioactive peptides in human health: Challenges and opportunities. Nutrients. 2018;10(11):1738.

20. Nosworthy MG, House JD. Factors influencing the quality of dietary proteins: Implications for pulses. Cereal Chemistry. 2017;94(1):49–57.

21. Wang J, Li D, Dangott LJ, Wu G. Proteomics and its role in nutrition research. The Journal of nutrition. 2006;136(7):1759–62.

22. Minkiewicz P, Dziuba J, Iwaniak A, Dziuba M, Darewicz M. BIOPEP database and other programs for processing bioactive peptide sequences. Journal of AOAC International. 2008;91(4):965–80.

23. Dolnicar S, Grün B, Leisch F. Market segmentation analysis. Market Segmentation Analysis: Springer; 2018. p. 11–22.

24. Millward DJ. The nutritional value of plant-based diets in relation to human amino acid and protein requirements. Proceedings of the Nutrition Society. 1999;58(2):249–60.

25. Cottrell JS. Protein identification using MS/MS data. Journal of proteomics. 2011;74(10):1842–51.

26. Ma B. Novor: real-time peptide de novo sequencing software. Journal of the American Society for Mass Spectrometry. 2015;26(11):1885–94.

27. Johnson RS, Searle BC, Nunn BL, Gilmore JM, Phillips M, Amemiya CT, et al. Assessing protein sequence database suitability using de novo sequencing. Molecular & Cellular Proteomics. 2020;19(1):198–208.

28. Olsen JV, Ong S-E, Mann M. Trypsin cleaves exclusively C-terminal to arginine and lysine residues. Molecular & Cellular Proteomics. 2004;3(6):608–14.

29. Sharma S, Singh R, Rana S. Bioactive peptides: a review. Int J Bioautomation. 2011;15(4):223–50.

30. Nielsen SD, Beverly RL, Qu Y, Dallas DC. Milk bioactive peptide database: A comprehensive database of milk protein-derived bioactive peptides and novel visualization. Food chemistry. 2017;232:673–82.

31. Onwulata C, Huth P. Whey processing, functionality and health benefits: John Wiley & Sons; 2009.

32. Gu R-Z, Li C-Y, Liu W-Y, Yi W-X, Cai M-Y,. Angiotensin I-converting enzyme inhibitory activity of low-molecular-weight peptides from Atlantic salmon (Salmo salar L.) skin. Food Research International. 2011;44(5):1536–40.

33. Bernstein KE, Ong FS, Blackwell W-LB, Shah KH, Giani JF, Gonzalez-Villalobos RA, et al. A modern understanding of the traditional and nontraditional biological functions of angiotensin-converting enzyme. Pharmacological reviews. 2013;65(1):1–46.

34. Ghosh SS, Wang J, Yannie PJ, Ghosh S. Intestinal Barrier Dysfunction, LPS Translocation, and Disease Development. Journal of the Endocrine Society. 2020;4(2):bvz039.

35. Lotz M, Gütle D, Walther S, Ménard S, Bogdan C, Hornef MW. Postnatal acquisition of endotoxin tolerance in intestinal epithelial cells. The Journal of experimental medicine. 2006;203(4):973–84.

36. Desser L, Holomanova D, Zavadova E, Pavelka K, Mohr T, Herbacek I. Oral therapy with proteolytic enzymes decreases excessive TGF-β levels in human blood. Cancer chemotherapy and pharmacology. 2001;47(1):S10–S5.

37. Lundberg JO, Weitzberg E, Cole JA, Benjamin N. Nitrate, bacteria and human health. Nature Reviews Microbiology. 2004;2(7):593.

38. Kim HS, Hur SJ. Changes of sodium nitrate, nitrite, and N-nitrosodiethylamine during in vitro human digestion. Food chemistry. 2017;225:197–201.

39. Jones S, Trejdosiewicz L, Banks R, Howdle P, Axon A, Dixon M, et al. Expression of interleukin-6 by intestinal enterocytes. Journal of clinical pathology. 1993;46(12):1097–100.

40. Kuhn KA, Manieri NA, Liu T-C, Stappenbeck TS. IL-6 stimulates intestinal epithelial proliferation and repair after injury. PloS one. 2014;9(12).

41. Maeda K, Mehta H, Drevets DA, Coggeshall KM. IL-6 increases B-cell IgG production in a feed-forward proinflammatory mechanism to skew hematopoiesis and elevate myeloid production. Blood, The Journal of the American Society of Hematology. 2010;115(23):4699–706.

42. Lanas A. Role of nitric oxide in the gastrointestinal tract. Arthritis research & therapy. 2008;10(2):S4.

43. Kolios G, Valatas V, Ward SG. Nitric oxide in inflammatory bowel disease: a universal messenger in an unsolved puzzle. Immunology. 2004;113(4):427–37.

44. Minekus M, Alminger M, Alvito P, Ballance S, Bohn T, Bourlieu C, et al. A standardised static in vitro digestion method suitable for food–an international consensus. Food & function. 2014;5(6):1113–24.

45. Adler-Nissen J. Enzymic hydrolysis of food proteins: Elsevier applied science publishers; 1986.

