## Supplementary data for "A combined microphysiological-computational omics approach in dietary protein evaluation"

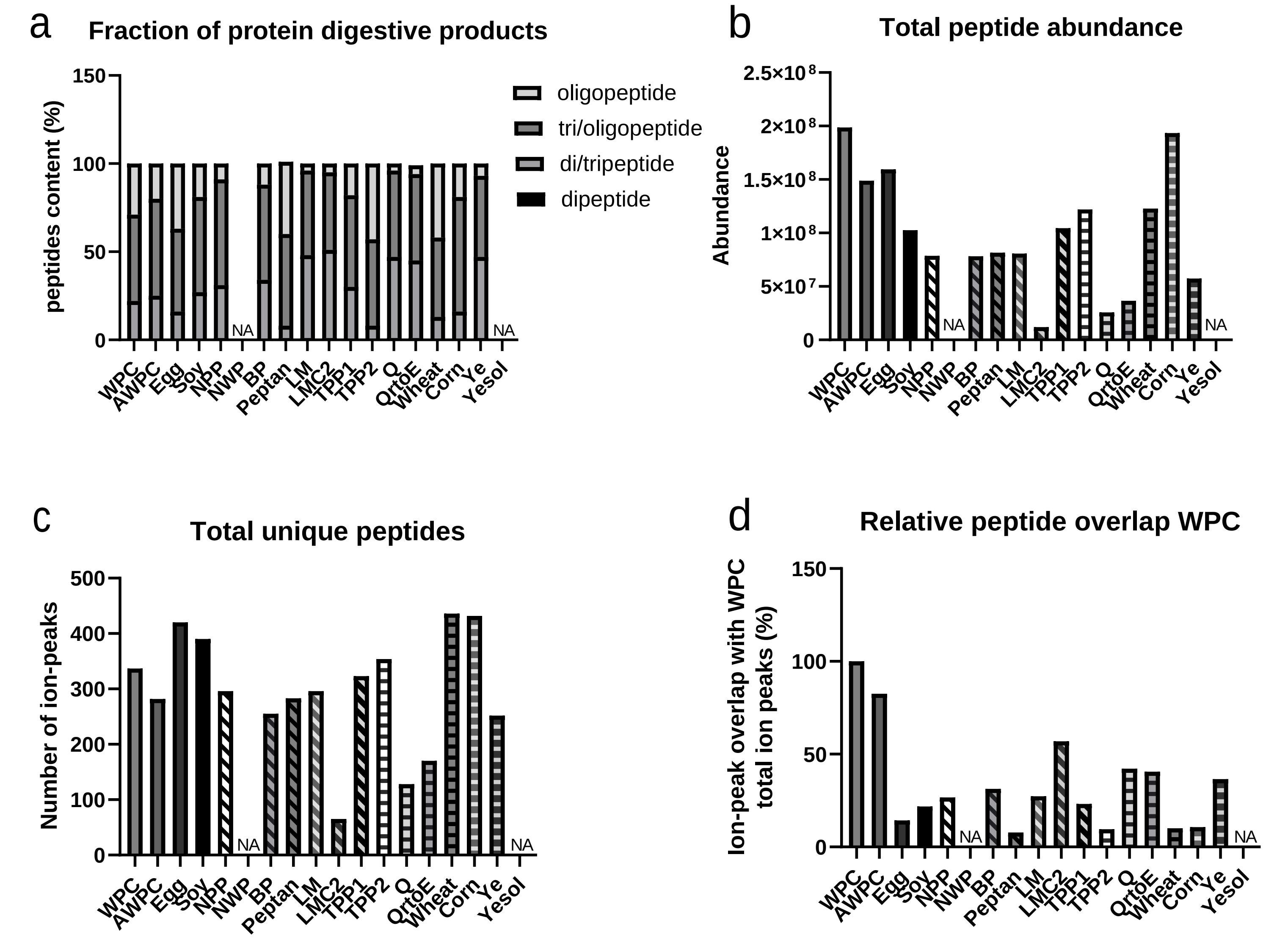

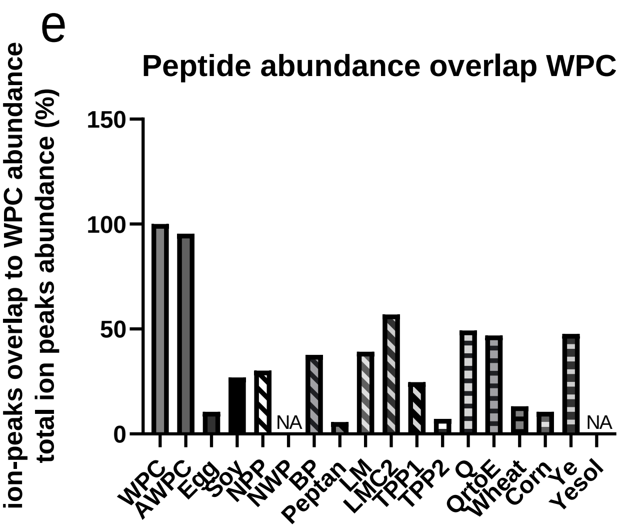

**Supplementary Figure 1.** Proteomic analysis of potential dietary protein in comparison to whey protein concentrate (WPC).

**Supplementary figure 2: overview of potential dietary protein tile plots, visualizing peptide overlap and non-overlap with WPC and WPC-matched peptides and their quantity.** Unique peptides of WPC were labelled with individual ID numbers (#1-337), if matched with another protein source, matching peptide received WPC ID#. Non overlapping peptides received a new ID# (>338). Abundance of overlapping peptides were plotted based on ID# on the right.

### AWPC

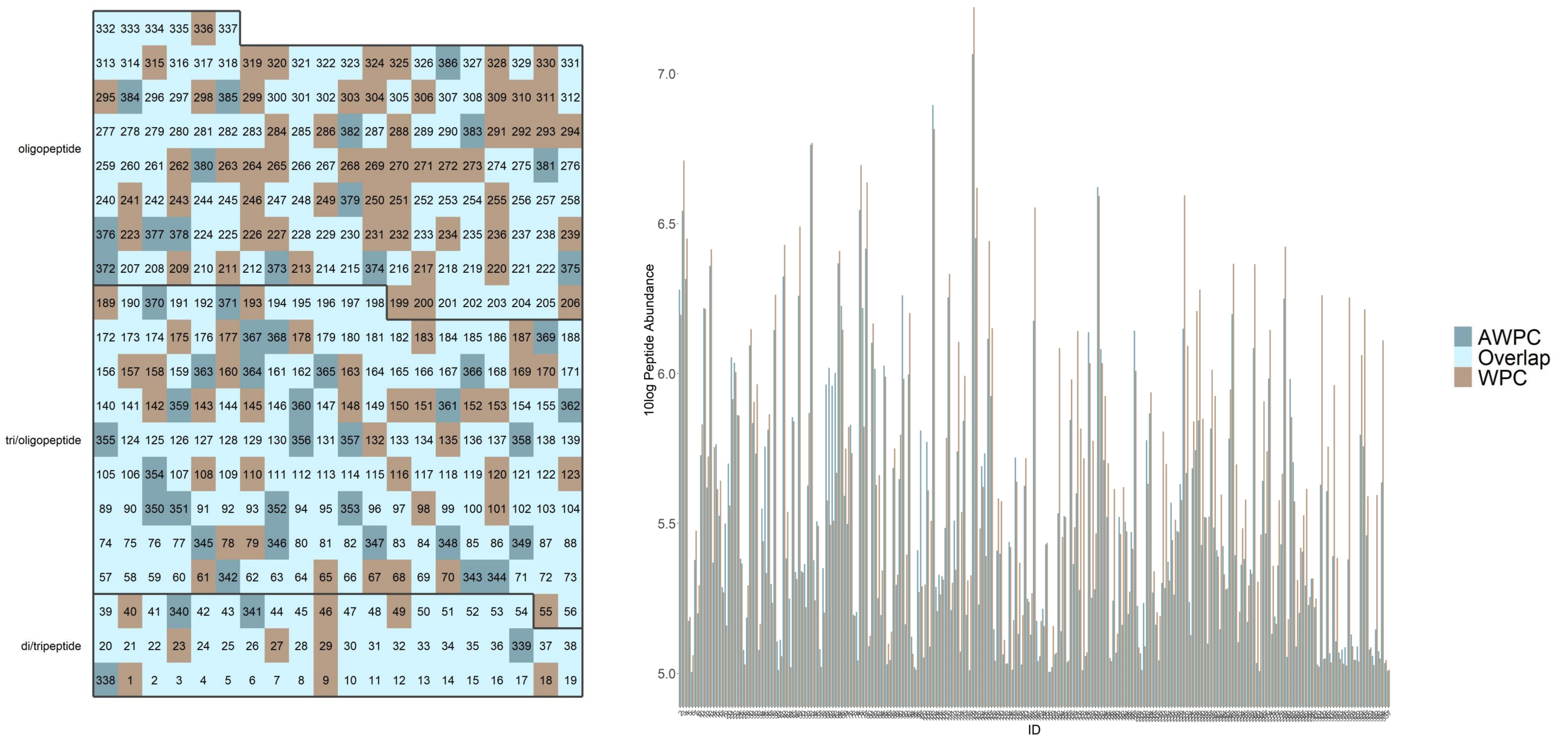

**Tile plot of peptide overlap and direct comparison in peptide concentration of AWPC in comparison to WPC.** Left shows the tile plot visualizing the amount of overlap, and non-overlapping peptides (ion peaks) sorted by ID number. The right figure visualizes quantities of WPC-matched peptides (ion peaks) sorted by ID number. ID numbers including information can be found in Suppl. File 1.

### Egg

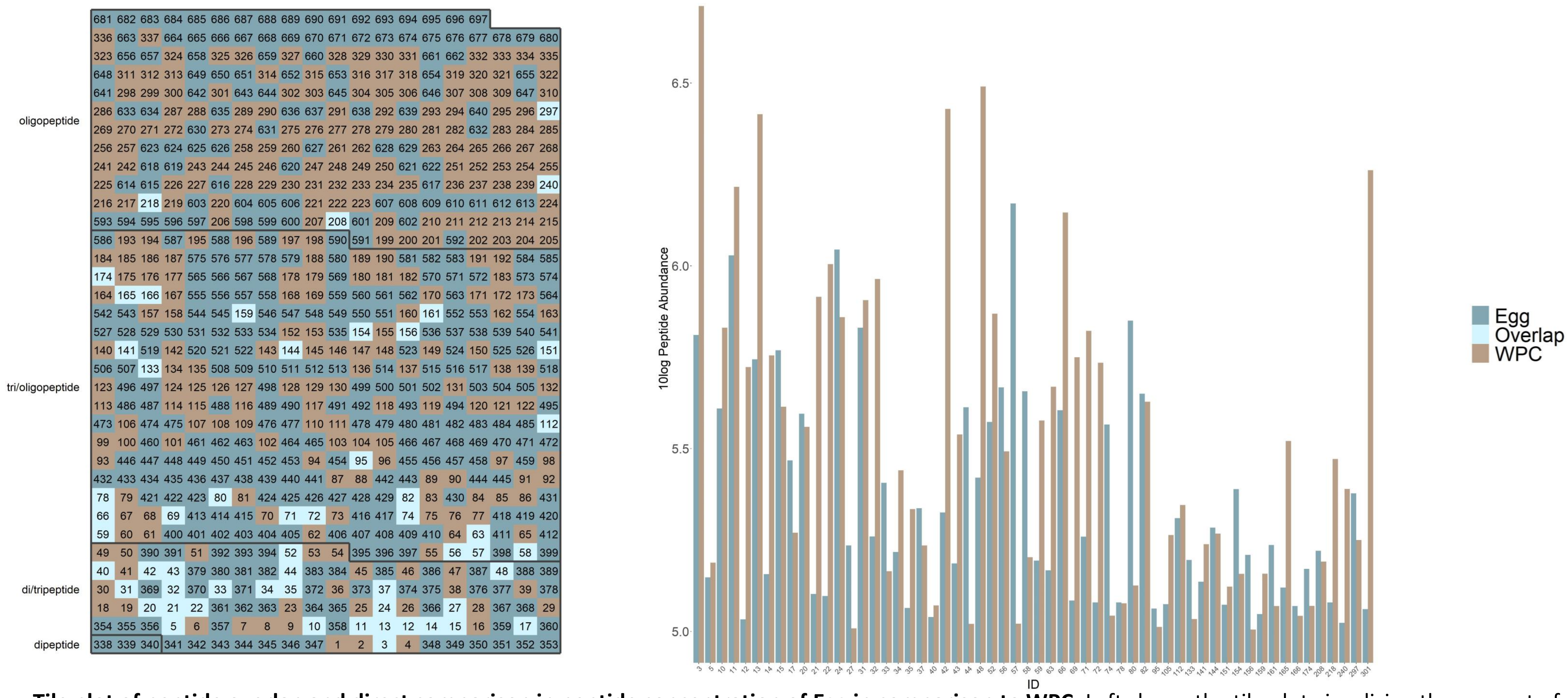

**Tile plot of peptide overlap and direct comparison in peptide concentration of Egg in comparison to WPC.** Left shows the tile plot visualizing the amount of overlap, and non-overlapping peptides (ion peaks) sorted by ID number. The right figure visualizes quantities of WPC-matched peptides (ion peaks) sorted by ID number. ID numbers including information can be found in Suppl. File 1.

### Soy

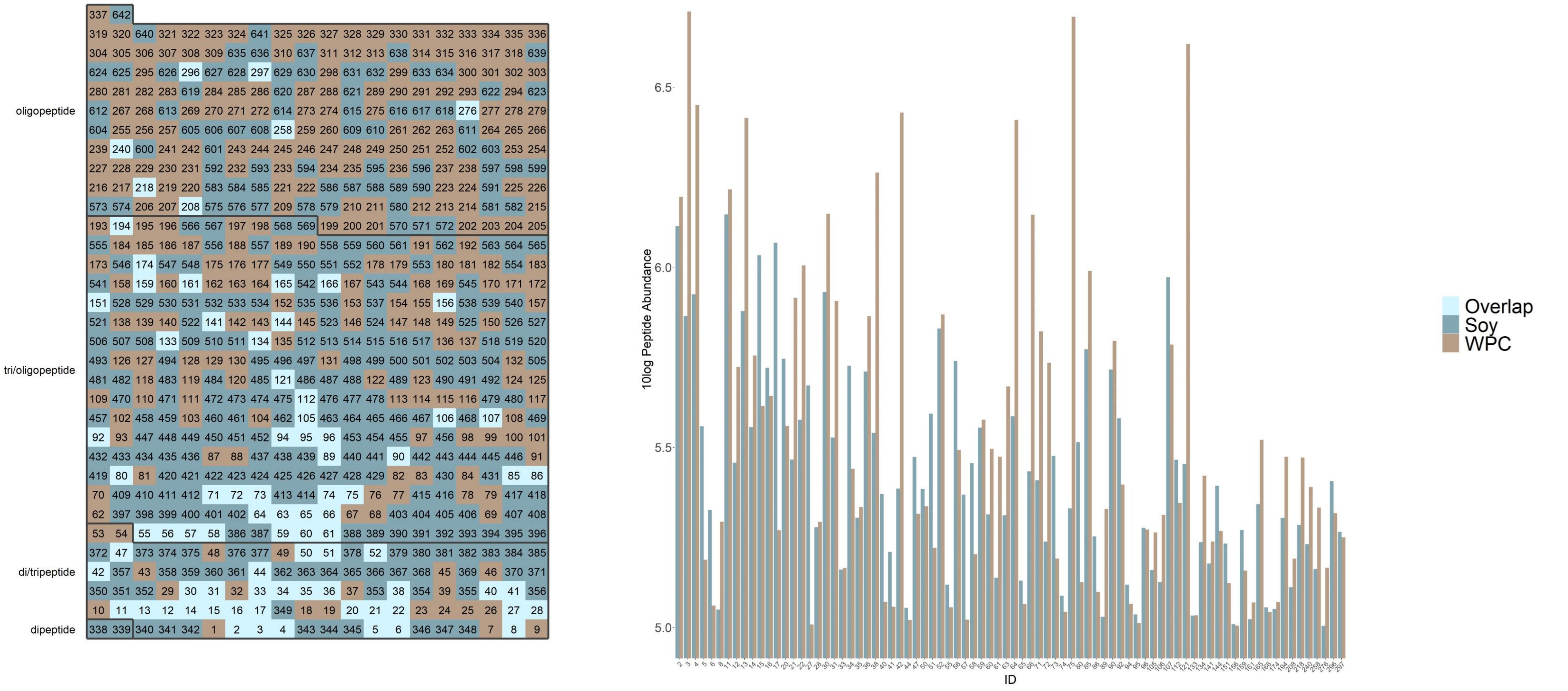

**Tile plot of peptide overlap and direct comparison in peptide concentration of Soy in comparison to WPC.** Left shows the tile plot visualizing the amount of overlap, and non-overlapping peptides (ion peaks) sorted by ID number. The right figure visualizes quantities of WPC-matched peptides (ion peaks) sorted by ID number. ID numbers including information can be found in Suppl. File 1.

### NPP

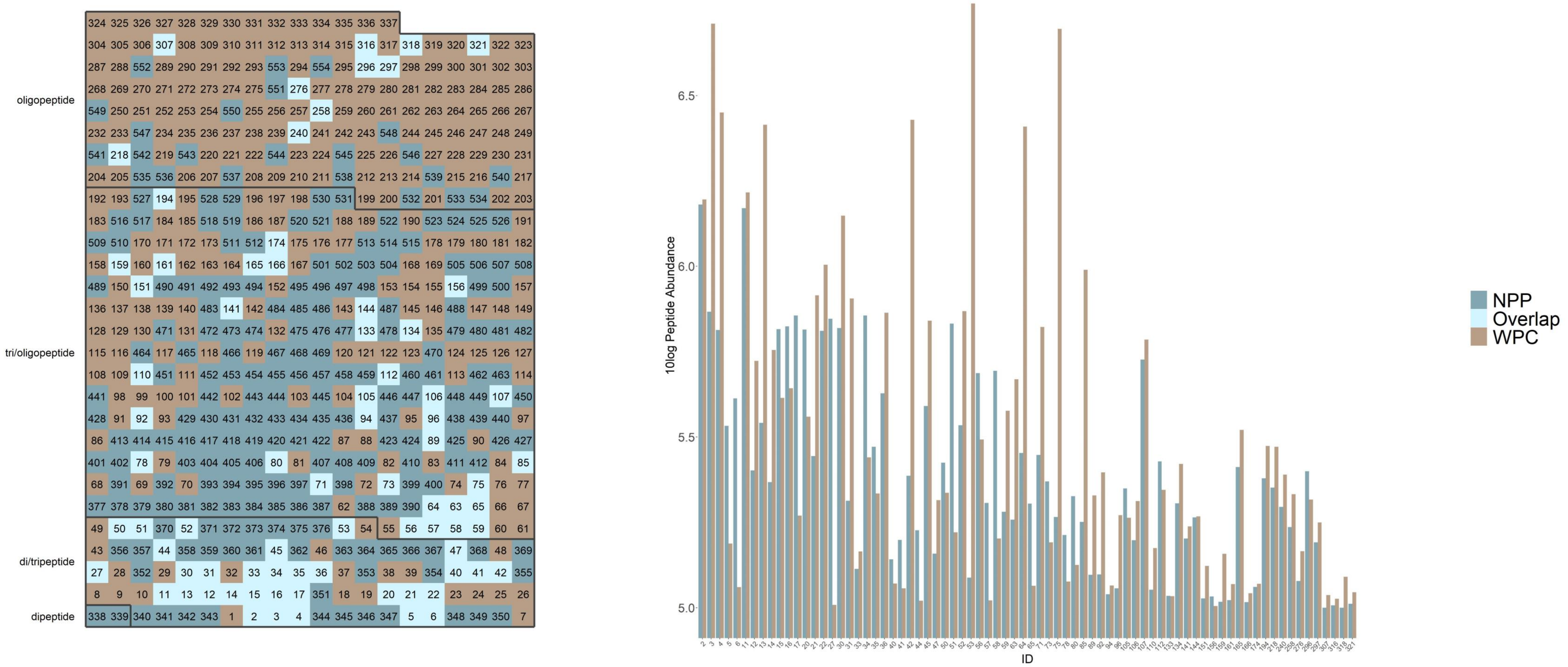

**Tile plot of peptide overlap and direct comparison in peptide concentration of NPP in comparison to WPC.** Left shows the tile plot visualizing the amount of overlap, and non-overlapping peptides (ion peaks) sorted by ID number. The right figure visualizes quantities of WPC-matched peptides (ion peaks) sorted by ID number. ID numbers including information can be found in Suppl. File 1.

# BP

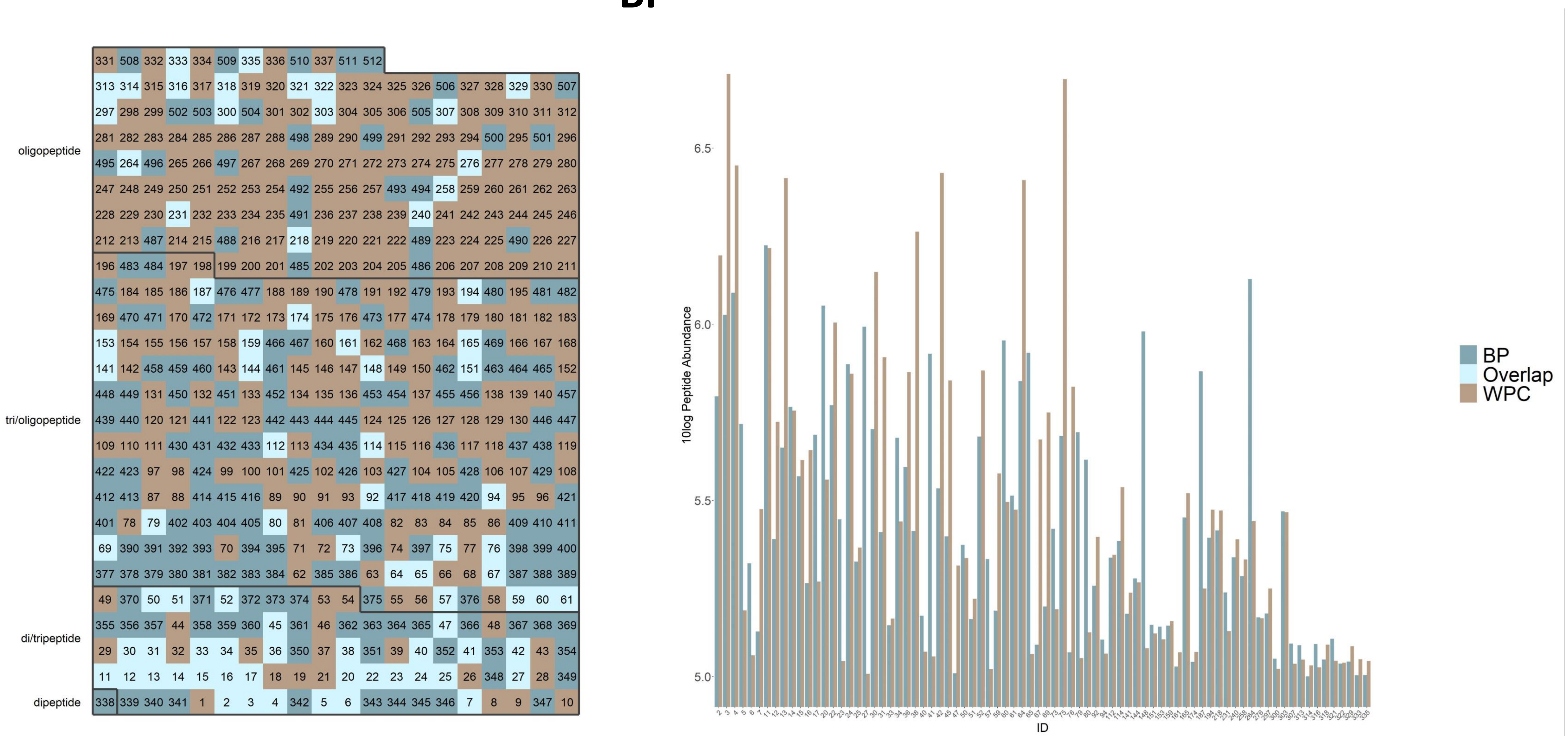

**Tile plot of peptide overlap and direct comparison in peptide concentration of BP in comparison to WPC.** Left shows the tile plot visualizing the amount of overlap, and non-overlapping peptides (ion peaks) sorted by ID number. The right figure visualizes quantities of WPC-matched peptides (ion peaks) sorted by ID number. ID numbers including information can be found in Suppl. File 1.

### Peptan

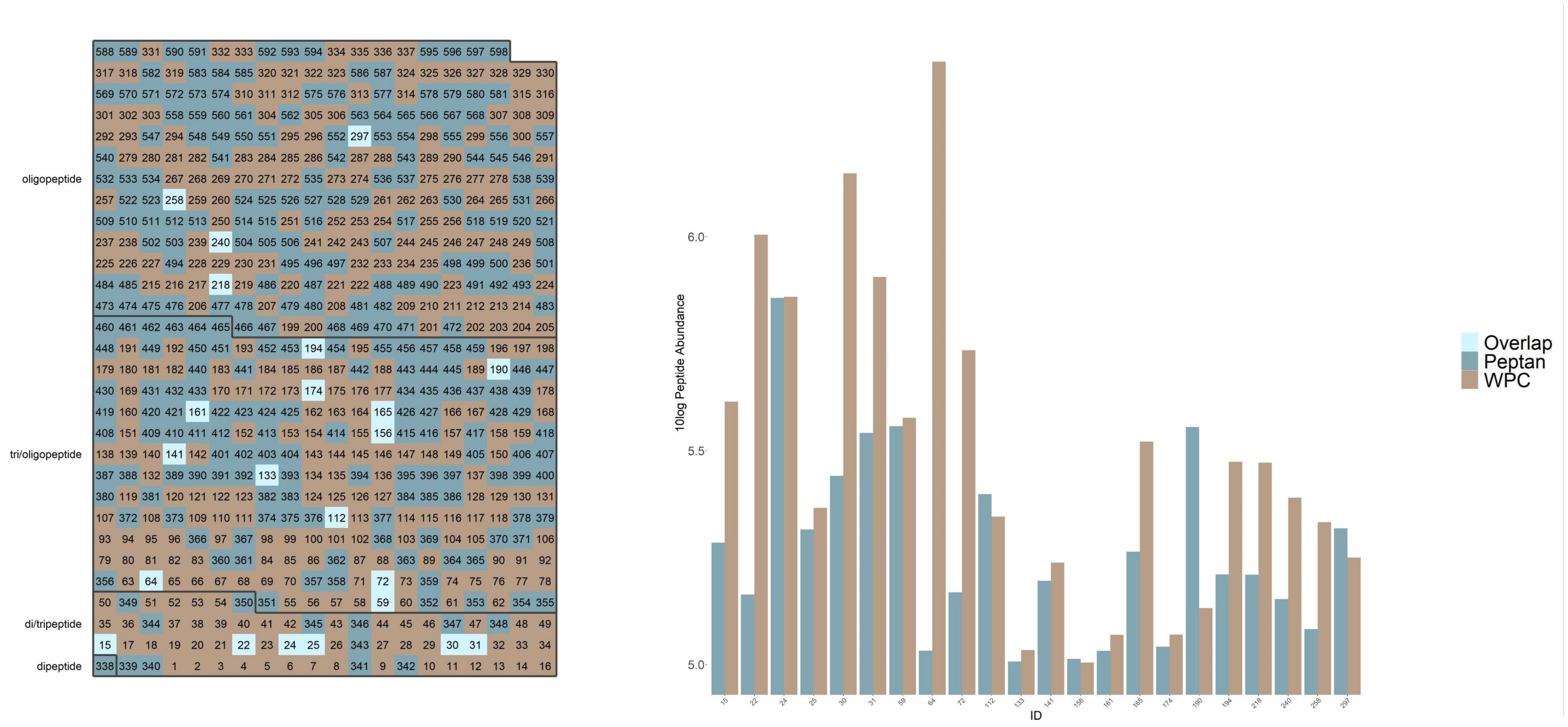

**Tile plot of peptide overlap and direct comparison in peptide concentration of Peptan in comparison to WPC.** Left shows the tile plot visualizing the amount of overlap, and non-overlapping peptides (ion peaks) sorted by ID number. The right figure visualizes quantities of WPC-matched peptides (ion peaks) sorted by ID number. ID numbers including information can be found in Suppl. File 1.

# LM

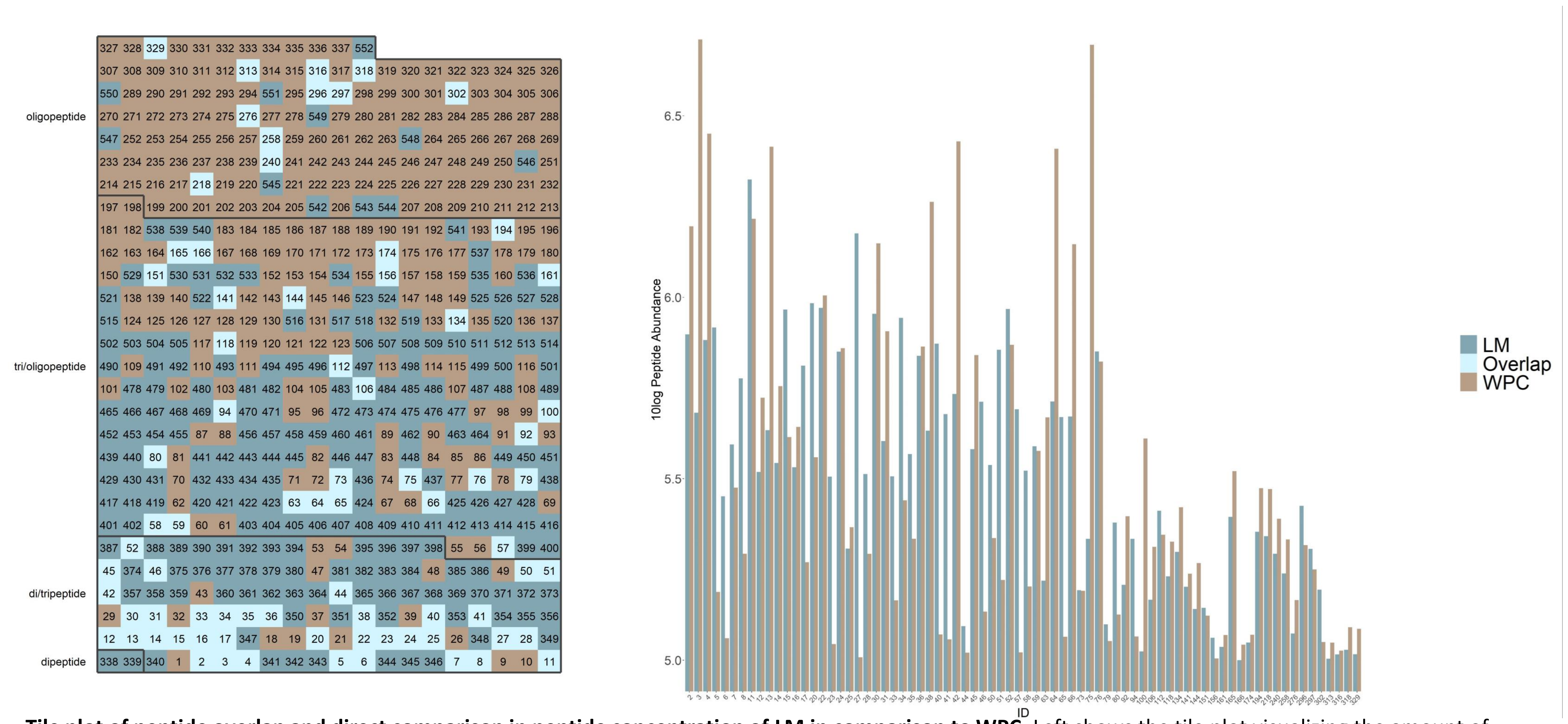

**Tile plot of peptide overlap and direct comparison in peptide concentration of LM in comparison to WPC.** Left shows the tile plot visualizing the amount of overlap, and non-overlapping peptides (ion peaks) sorted by ID number. The right figure visualizes quantities of WPC-matched peptides (ion peaks) sorted by ID number. ID numbers including information can be found in Suppl. File 1.

### LMC2

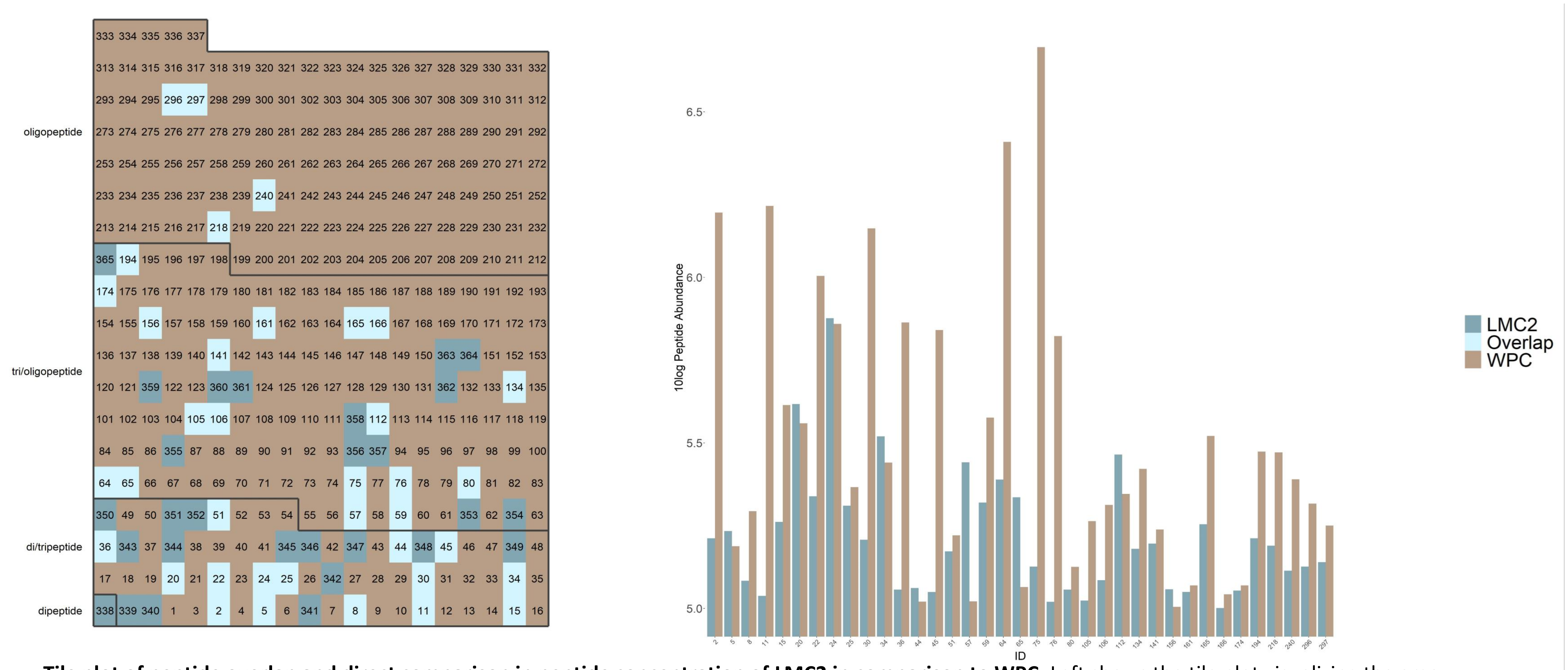

**Tile plot of peptide overlap and direct comparison in peptide concentration of LMC2 in comparison to WPC.** Left shows the tile plot visualizing the amou[illegible] overlap, and non-overlapping peptides (ion peaks) sorted by ID number. The right figure visualizes quantities of WPC-matched peptides (ion peaks) sorted by ID number. ID numbers including information can be found in Suppl. File 1.

### TPP1

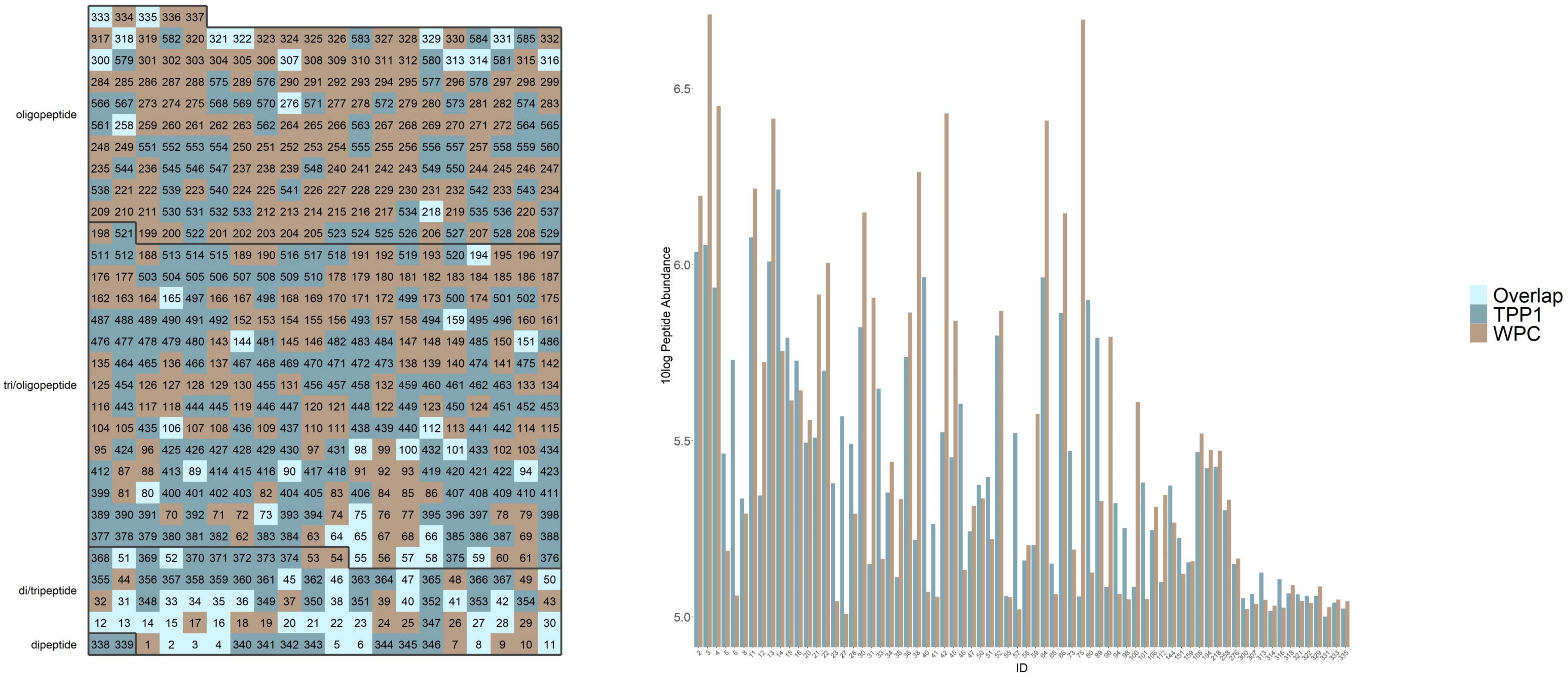

**Tile plot of peptide overlap and direct comparison in peptide concentration of TPP1 in comparison to WPC.** Left shows the tile plot visualizing the amount of overlap, and non-overlapping peptides (ion peaks) sorted by ID number. The right figure visualizes quantities of WPC-matched peptides (ion peaks) sorted by ID number. ID numbers including information can be found in Suppl. File 1.

### TPP2

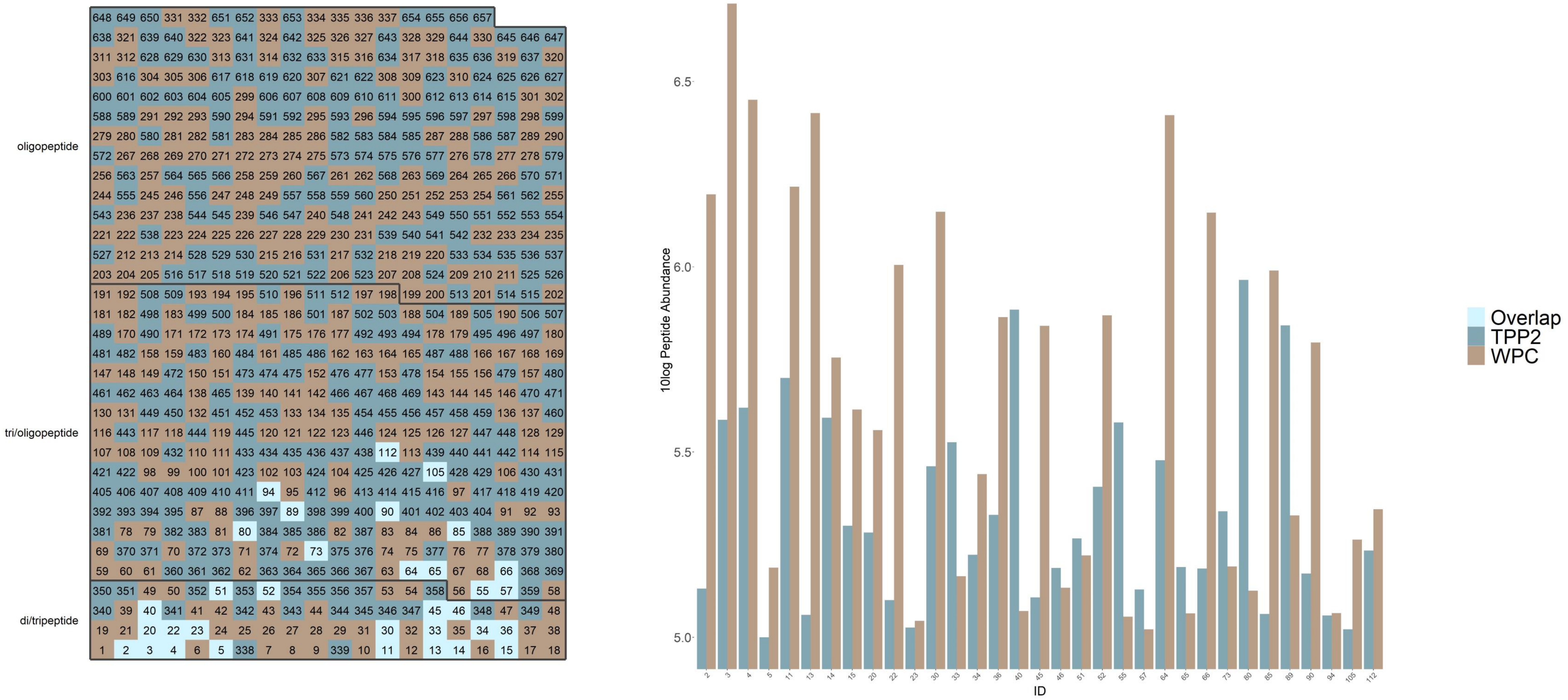

**Tile plot of peptide overlap and direct comparison in peptide concentration of TPP2 in comparison to WPC.** Left shows the tile plot visualizing the amount of overlap, and non-overlapping peptides (ion peaks) sorted by ID number. The right figure visualizes quantities of WPC-matched peptides (ion peaks) sorted by ID number. ID numbers including information can be found in Suppl. File 1.

# Q

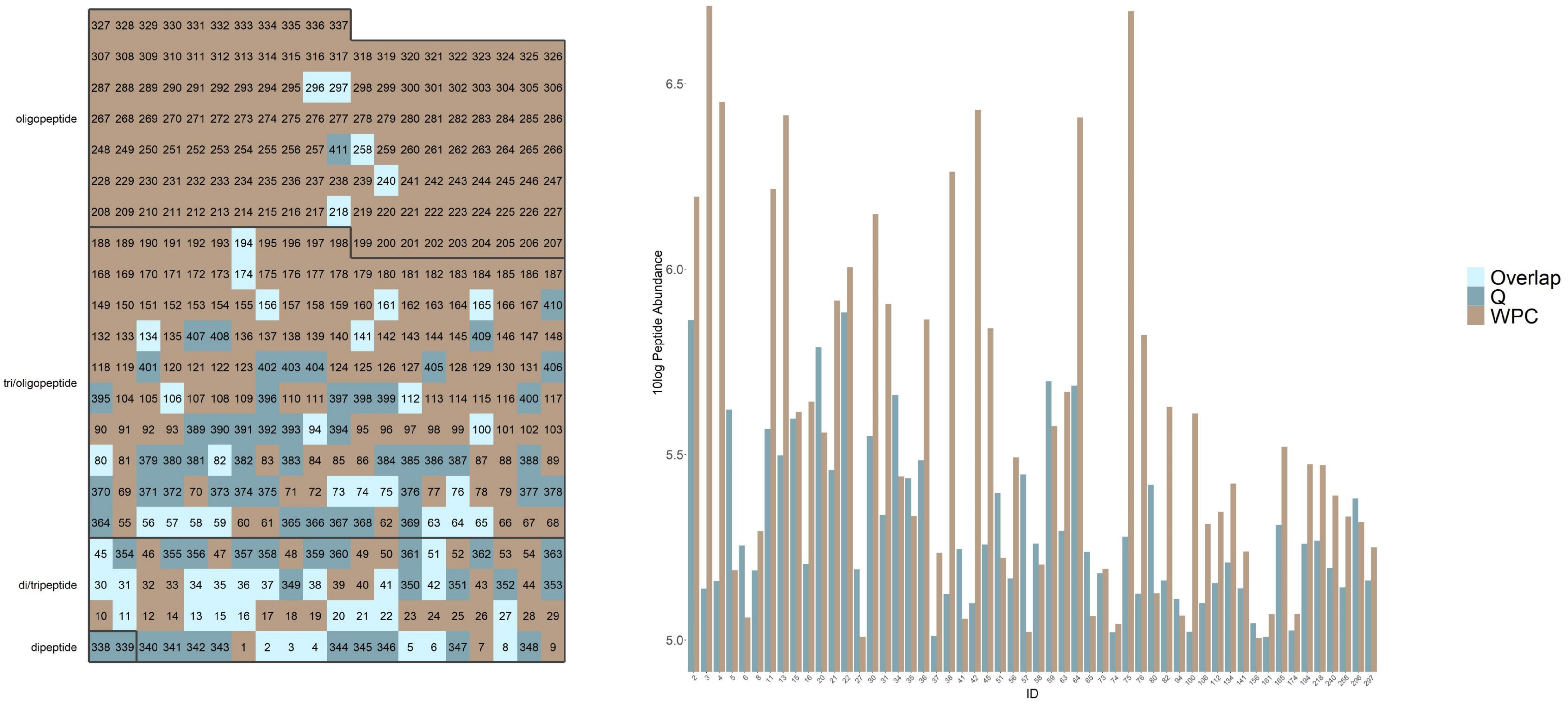

**Tile plot of peptide overlap and direct comparison in peptide concentration of Q in comparison to WPC.** Left shows the tile plot visualizing the amount of overlap, and non-overlapping peptides (ion peaks) sorted by ID number. The right figure visualizes quantities of WPC-matched peptides (ion peaks) sorted by ID number. ID numbers including information can be found in Suppl. File 1.

### QrtoE

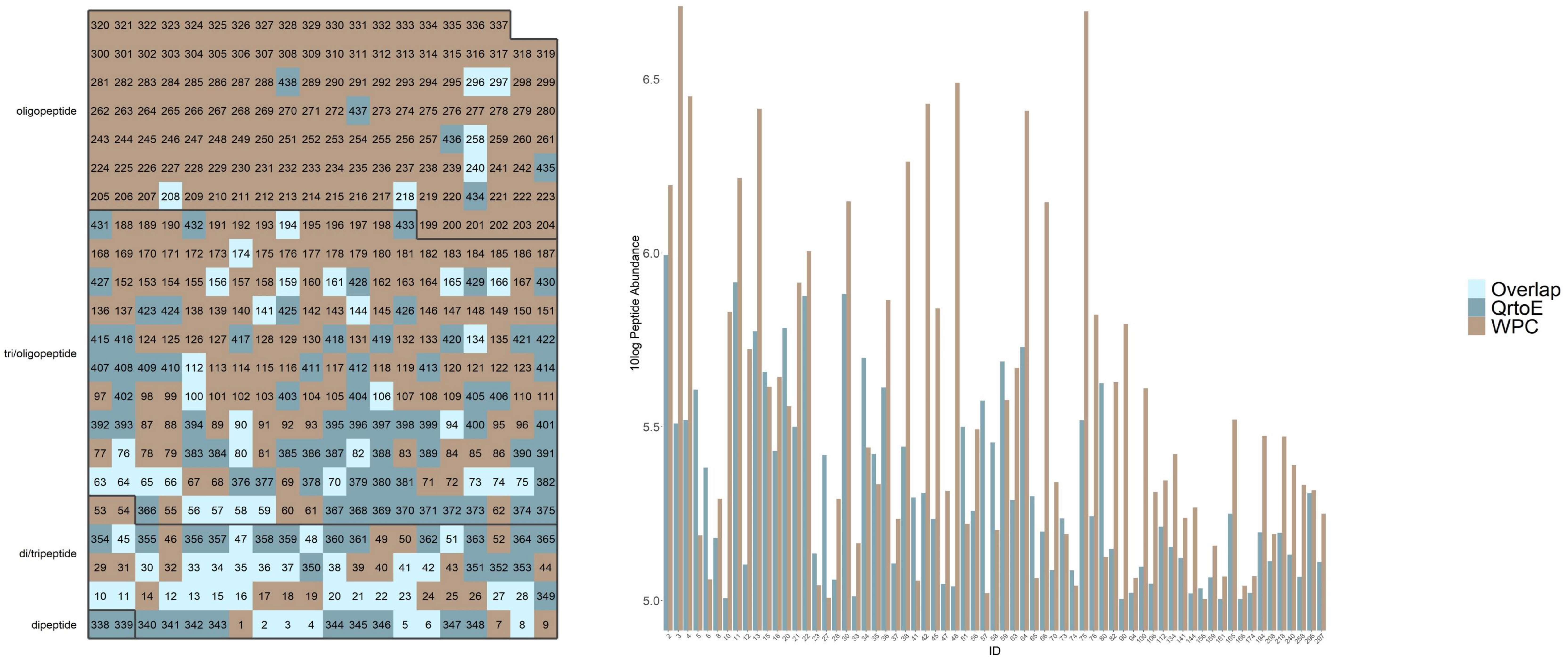

**Tile plot of peptide overlap and direct comparison in peptide concentration of QrtoE in comparison to WPC.** Left shows the tile plot visualizing the amount of overlap, and non-overlapping peptides (ion peaks) sorted by ID number. The right figure visualizes quantities of WPC-matched peptides (ion peaks) sorted by ID number. ID numbers including information can be found in Suppl. File 1.

### Wheat

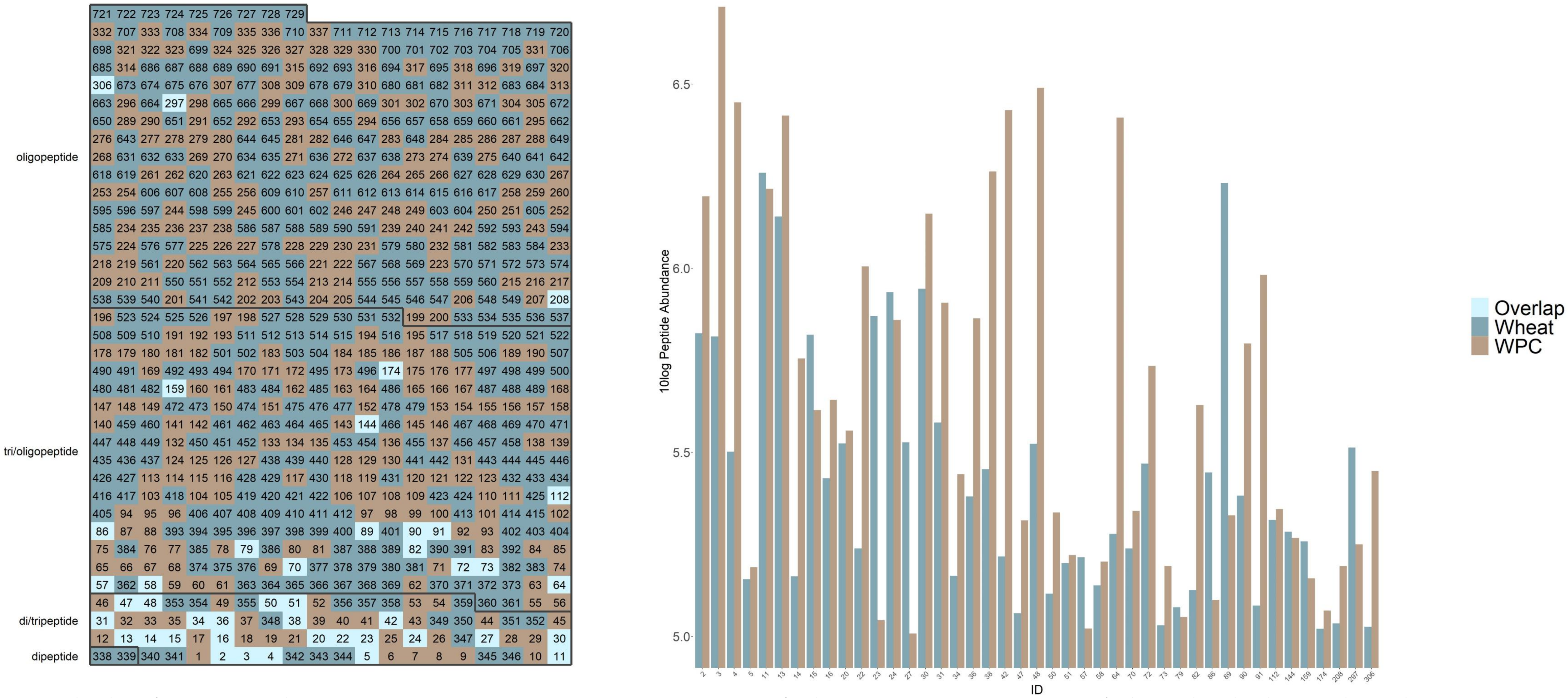

**Tile plot of peptide overlap and direct comparison in peptide concentration of Wheat in comparison to WPC.** Left shows the tile plot visualizing the amount of overlap, and non-overlapping peptides (ion peaks) sorted by ID number. The right figure visualizes quantities of WPC-matched peptides (ion peaks) sorted by ID number. ID numbers including information can be found in Suppl. File 1.

### Corn

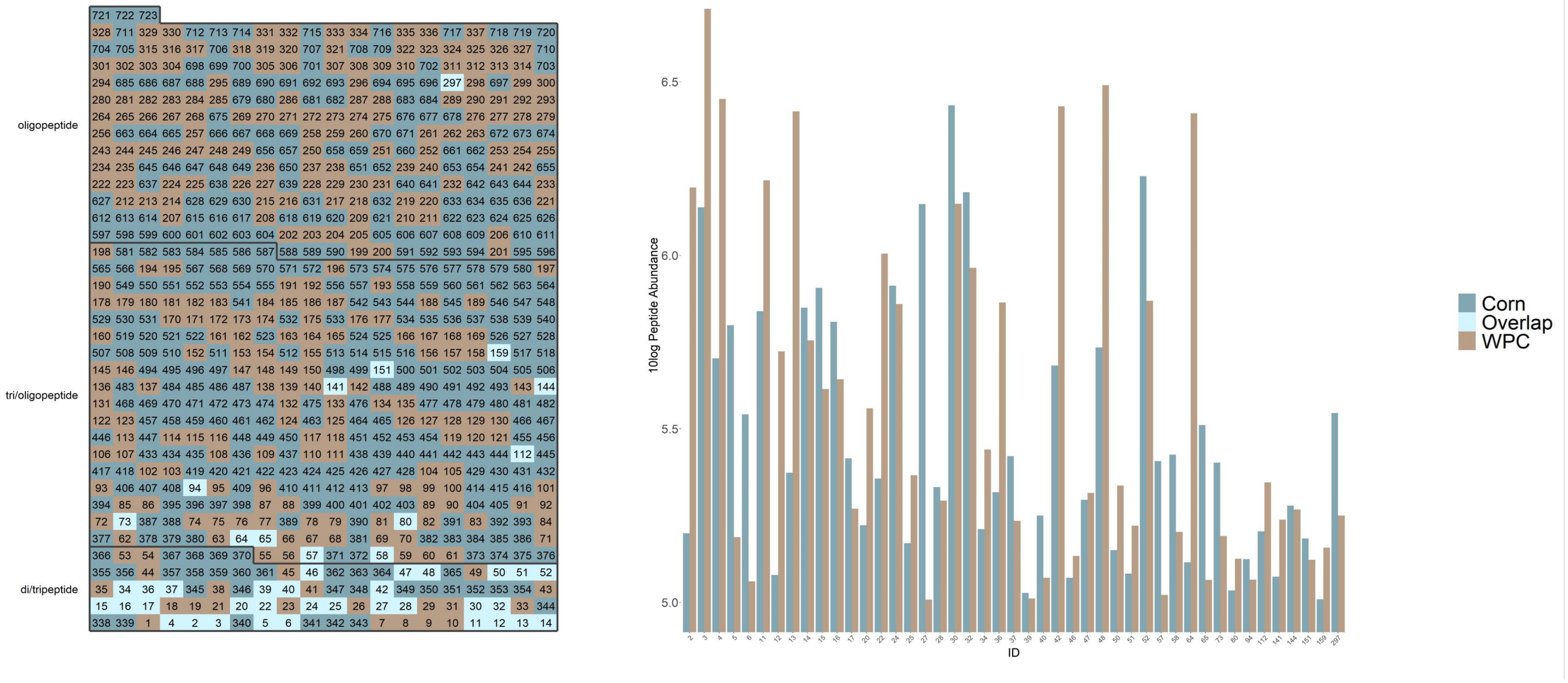

**Tile plot of peptide overlap and direct comparison in peptide concentration of Corn in comparison to WPC.** Left shows the tile plot visualizing the amount of overlap, and non-overlapping peptides (ion peaks) sorted by ID number. The right figure visualizes quantities of WPC-matched peptides (ion peaks) sorted by ID number. ID numbers including information can be found in Suppl. File 1.

# YE

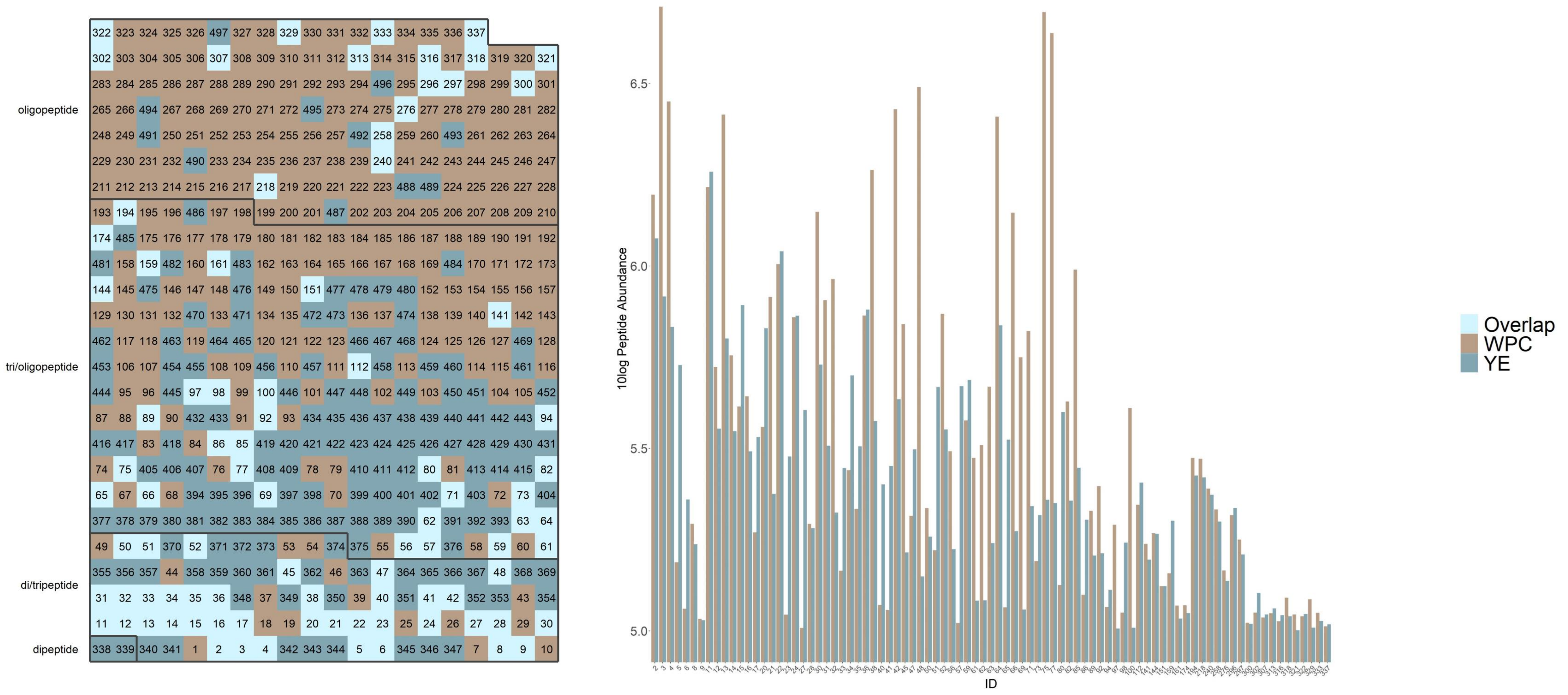

**Tile plot of peptide overlap and direct comparison in peptide concentration of YE in comparison to WPC.** Left shows the tile plot visualizing the amount of overlap, and non-overlapping peptides (ion peaks) sorted by ID number. The right figure visualizes quantities of WPC-matched peptides (ion peaks) sorted by ID number. ID numbers including information can be found in Suppl. File 1.

**Supplementary Table 1.** Details on overlapping and non-overlapping ion-peak data

| ID | Protein | mz | charge | retentiont | raw.abund | mass | type |
|---|---|---|---|---|---|---|---|
| 1 | WPC | 230.248 | 1 | 36.62458 | 218546.5 | 229.2407 | di/tripeptide |
| 2 | WPC | 231.1708 | 1 | 5.204583 | 1568927 | 230.1635 | di/tripeptide |
| 3 | WPC | 231.1708 | 1 | 9.601775 | 5129673 | 230.1635 | di/tripeptide |
| 4 | WPC | 231.1708 | 1 | 12.02929 | 2821537 | 230.1636 | di/tripeptide |
| 5 | WPC | 233.1497 | 1 | 2.2007 | 154251.9 | 232.1424 | di/tripeptide |
| 6 | WPC | 233.1498 | 1 | 2.780942 | 114979.6 | 232.1426 | di/tripeptide |
| 7 | WPC | 239.1027 | 1 | 2.18985 | 298924.1 | 238.0954 | di/tripeptide |
| 8 | WPC | 239.1029 | 1 | 2.425233 | 196607.6 | 238.0956 | di/tripeptide |
| 9 | WPC | 243.1345 | 1 | 5.418808 | 107826.2 | 242.1272 | di/tripeptide |
| 10 | WPC | 245.1133 | 1 | 1.978642 | 677213.8 | 244.106 | di/tripeptide |
| 11 | WPC | 245.1863 | 1 | 16.42797 | 1644069 | 244.179 | di/tripeptide |
| 12 | WPC | 245.1865 | 1 | 15.90611 | 528585.5 | 244.1793 | di/tripeptide |
| 13 | WPC | 245.1866 | 1 | 14.55145 | 2597514 | 244.1793 | di/tripeptide |
| 14 | WPC | 245.1866 | 1 | 17.6888 | 568858.5 | 244.1793 | di/tripeptide |
| 15 | WPC | 246.145 | 1 | 2.123375 | 411579.4 | 245.1377 | di/tripeptide |
| 16 | WPC | 246.1451 | 1 | 2.6523 | 439301.4 | 245.1378 | di/tripeptide |
| 17 | WPC | 246.1453 | 1 | 3.364567 | 186173.6 | 245.138 | di/tripeptide |
| 18 | WPC | 247.1108 | 1 | 20.3645 | 123176.6 | 246.1036 | di/tripeptide |
| 19 | WPC | 247.1113 | 1 | 2.381492 | 144495.1 | 246.1041 | di/tripeptide |
| 20 | WPC | 247.129 | 1 | 2.18985 | 362628.6 | 246.1217 | di/tripeptide |
| 21 | WPC | 247.1291 | 1 | 3.147667 | 822062.9 | 246.1218 | di/tripeptide |
| 22 | WPC | 247.1293 | 1 | 4.658808 | 1010908 | 246.122 | di/tripeptide |
| 23 | WPC | 249.1271 | 1 | 2.795658 | 110783.5 | 248.1198 | di/tripeptide |
| 24 | WPC | 250.178 | 1 | 34.86286 | 723803.2 | 249.1707 | di/tripeptide |
| 25 | WPC | 250.1781 | 1 | 37.31456 | 232458 | 249.1708 | di/tripeptide |
| 26 | WPC | 252.9628 | 1 | 4.78395 | 107139.5 | 251.9555 | di/tripeptide |
| 27 | WPC | 253.1185 | 1 | 2.301917 | 101901.8 | 252.1112 | di/tripeptide |
| 28 | WPC | 253.1188 | 1 | 9.2948 | 196395.1 | 252.1115 | di/tripeptide |
| 29 | WPC | 258.2794 | 1 | 37.62385 | 527017 | 257.2722 | di/tripeptide |
| 30 | WPC | 260.1607 | 1 | 2.18985 | 1406430 | 259.1534 | di/tripeptide |
| 31 | WPC | 260.1608 | 1 | 2.6523 | 805315.3 | 259.1535 | di/tripeptide |
| 32 | WPC | 260.161 | 1 | 3.703708 | 920707.1 | 259.1537 | di/tripeptide |
| 33 | WPC | 260.1971 | 1 | 2.18985 | 146183.2 | 259.1898 | di/tripeptide |
| 34 | WPC | 261.1447 | 1 | 2.18985 | 275836.5 | 260.1375 | di/tripeptide |
| 35 | WPC | 261.1449 | 1 | 3.2809 | 216032.6 | 260.1376 | di/tripeptide |
| 36 | WPC | 261.145 | 1 | 5.410683 | 731320.4 | 260.1378 | di/tripeptide |
| 37 | WPC | 262.1399 | 1 | 2.093217 | 171943.3 | 261.1327 | di/tripeptide |
| 38 | WPC | 263.143 | 1 | 11.50558 | 1830714 | 262.1357 | di/tripeptide |
| 39 | WPC | 263.1967 | 1 | 2.18985 | 102589.8 | 262.1895 | di/tripeptide |
| 40 | WPC | 265.1553 | 1 | 16.2901 | 117787.5 | 264.148 | di/tripeptide |
| 41 | WPC | 267.1343 | 1 | 2.404683 | 114092.9 | 266.127 | di/tripeptide |
| 42 | WPC | 269.1608 | 1 | 2.157517 | 2686633 | 268.1536 | di/tripeptide |
| 43 | WPC | 274.1875 | 1 | 1.978642 | 345626.5 | 273.1802 | di/tripeptide |
| 44 | WPC | 277.1032 | 1 | 1.86235 | 104868 | 276.096 | di/tripeptide |
| 45 | WPC | 281.1134 | 1 | 2.566517 | 693056.2 | 280.1062 | di/tripeptide |
| 46 | WPC | 281.1503 | 1 | 6.977133 | 136167.2 | 280.143 | di/tripeptide |
| 47 | WPC | 288.1925 | 1 | 16.08898 | 206592.3 | 287.1852 | di/tripeptide |
| 48 | WPC | 290.1714 | 1 | 2.712558 | 3091093 | 289.1641 | di/tripeptide |
| 49 | WPC | 292.1327 | 1 | 2.18985 | 117891.6 | 291.1254 | di/tripeptide |

| | | | | | | | |
|---|---|---|---|---|---|---|---|
| 50 | WPC | 294.1451 | 1 | 2.324467 | 217038.3 | 293.1379 | di/tripeptide |
| 51 | WPC | 295.1295 | 1 | 3.245917 | 166335.6 | 294.1222 | di/tripeptide |
| 52 | WPC | 295.1657 | 1 | 16.42797 | 739234.7 | 294.1585 | di/tripeptide |
| 53 | WPC | 300.192 | 1 | 2.5938 | 5880434 | 299.1847 | di/tripeptide |
| 54 | WPC | 300.1928 | 1 | 15.14582 | 175279.4 | 299.1855 | di/tripeptide |
| 55 | WPC | 302.208 | 1 | 16.42797 | 113690.3 | 301.2007 | tri/oligopeptide |
| 56 | WPC | 302.2082 | 1 | 15.16074 | 310777.4 | 301.2009 | tri/oligopeptide |
| 57 | WPC | 302.2084 | 1 | 19.87094 | 105064.9 | 301.2011 | tri/oligopeptide |
| 58 | WPC | 303.1666 | 1 | 2.2007 | 159566.5 | 302.1594 | tri/oligopeptide |
| 59 | WPC | 304.1506 | 1 | 2.404683 | 377347 | 303.1433 | tri/oligopeptide |
| 60 | WPC | 304.1509 | 1 | 4.823717 | 313274.7 | 303.1437 | tri/oligopeptide |
| 61 | WPC | 304.1664 | 1 | 19.00618 | 297847.9 | 303.1591 | tri/oligopeptide |
| 62 | WPC | 316.2237 | 1 | 13.87745 | 322898.2 | 315.2164 | tri/oligopeptide |
| 63 | WPC | 318.1664 | 1 | 2.531717 | 466715 | 317.1591 | tri/oligopeptide |
| 64 | WPC | 318.1665 | 1 | 2.324467 | 2564465 | 317.1592 | tri/oligopeptide |
| 65 | WPC | 318.1669 | 1 | 9.051517 | 116011.3 | 317.1596 | tri/oligopeptide |
| 66 | WPC | 318.1821 | 1 | 22.57613 | 1399162 | 317.1748 | tri/oligopeptide |
| 67 | WPC | 318.1823 | 1 | 24.72373 | 471190.9 | 317.175 | tri/oligopeptide |
| 68 | WPC | 318.1824 | 1 | 23.73726 | 113843.7 | 317.1751 | tri/oligopeptide |
| 69 | WPC | 319.1615 | 1 | 2.168392 | 561926.7 | 318.1542 | tri/oligopeptide |
| 70 | WPC | 325.1133 | 1 | 1.719867 | 219276.2 | 324.106 | tri/oligopeptide |
| 71 | WPC | 330.2396 | 1 | 18.42354 | 663790.9 | 329.2324 | tri/oligopeptide |
| 72 | WPC | 331.1613 | 1 | 2.081408 | 542874.9 | 330.1541 | tri/oligopeptide |
| 73 | WPC | 331.1657 | 1 | 2.837858 | 155454.8 | 330.1584 | tri/oligopeptide |
| 74 | WPC | 331.2851 | 1 | 38.45748 | 110434.2 | 330.2778 | tri/oligopeptide |
| 75 | WPC | 332.1818 | 1 | 2.6191 | 4958838 | 331.1745 | tri/oligopeptide |
| 76 | WPC | 332.2186 | 1 | 3.085625 | 664880.6 | 331.2113 | tri/oligopeptide |
| 77 | WPC | 332.2187 | 1 | 3.418708 | 4341485 | 331.2114 | tri/oligopeptide |
| 78 | WPC | 333.1407 | 1 | 1.7666 | 119427.6 | 332.1335 | tri/oligopeptide |
| 79 | WPC | 333.1567 | 1 | 10.97644 | 113003.9 | 332.1494 | tri/oligopeptide |
| 80 | WPC | 334.1612 | 1 | 2.21185 | 133517.7 | 333.1539 | tri/oligopeptide |
| 81 | WPC | 334.1613 | 1 | 3.245917 | 1468881 | 333.1541 | tri/oligopeptide |
| 82 | WPC | 342.2399 | 1 | 21.82051 | 424860 | 341.2326 | tri/oligopeptide |
| 83 | WPC | 343.1239 | 1 | 1.719867 | 457935 | 342.1167 | tri/oligopeptide |
| 84 | WPC | 344.1826 | 1 | 5.567342 | 220477.6 | 343.1753 | tri/oligopeptide |
| 85 | WPC | 344.2555 | 1 | 21.54997 | 976111.4 | 343.2482 | tri/oligopeptide |
| 86 | WPC | 344.2556 | 1 | 20.18498 | 125510.4 | 343.2483 | tri/oligopeptide |
| 87 | WPC | 349.1723 | 1 | 1.86235 | 137873.8 | 348.165 | tri/oligopeptide |
| 88 | WPC | 350.1197 | 1 | 1.8327 | 562475.5 | 349.1125 | tri/oligopeptide |
| 89 | WPC | 358.2712 | 1 | 24.02735 | 213384.7 | 357.2639 | tri/oligopeptide |
| 90 | WPC | 359.2293 | 1 | 2.18985 | 624779.6 | 358.2221 | tri/oligopeptide |
| 91 | WPC | 360.196 | 1 | 18.73659 | 959584.1 | 359.1888 | tri/oligopeptide |
| 92 | WPC | 360.2134 | 1 | 2.736492 | 249087.5 | 359.2061 | tri/oligopeptide |
| 93 | WPC | 360.2137 | 1 | 13.6536 | 1589841 | 359.2064 | tri/oligopeptide |
| 94 | WPC | 363.1555 | 1 | 2.18985 | 116238.7 | 362.1482 | tri/oligopeptide |
| 95 | WPC | 365.1366 | 1 | 37.82153 | 102864.7 | 364.1293 | tri/oligopeptide |
| 96 | WPC | 366.214 | 1 | 2.18985 | 186795.1 | 365.2067 | tri/oligopeptide |
| 97 | WPC | 374.2297 | 1 | 19.34277 | 195229.6 | 373.2224 | tri/oligopeptide |
| 98 | WPC | 375.2239 | 1 | 2.2007 | 112242.4 | 374.2166 | tri/oligopeptide |
| 99 | WPC | 376.1467 | 1 | 1.719867 | 197899.7 | 375.1395 | tri/oligopeptide |

| | | | | | | | |
|---|---|---|---|---|---|---|---|
| 100 | WPC | 376.1719 | 1 | 2.2007 | 407839.7 | 375.1646 | tri/oligopeptide |
| 101 | WPC | 376.2276 | 1 | 21.63163 | 112528.8 | 375.2203 | tri/oligopeptide |
| 102 | WPC | 381.0801 | 1 | 1.6744 | 322600.8 | 380.0728 | tri/oligopeptide |
| 103 | WPC | 387.2246 | 1 | 13.99581 | 6537951 | 386.2173 | tri/oligopeptide |
| 104 | WPC | 389.2306 | 1 | 13.98086 | 161396.7 | 388.2233 | tri/oligopeptide |
| 105 | WPC | 390.1876 | 1 | 2.313292 | 183419.9 | 389.1803 | tri/oligopeptide |
| 106 | WPC | 395.0394 | 1 | 1.572517 | 205423.7 | 394.0321 | tri/oligopeptide |
| 107 | WPC | 399.2612 | 1 | 18.55638 | 610178.2 | 398.254 | tri/oligopeptide |
| 108 | WPC | 401.1786 | 1 | 1.510183 | 107765.2 | 400.1713 | tri/oligopeptide |
| 109 | WPC | 402.2354 | 1 | 11.39669 | 2149924 | 401.2281 | tri/oligopeptide |
| 110 | WPC | 405.162 | 1 | 1.844392 | 149490.1 | 404.1547 | tri/oligopeptide |
| 111 | WPC | 405.199 | 1 | 4.178875 | 200487.1 | 404.1918 | tri/oligopeptide |
| 112 | WPC | 415.2118 | 1 | 36.4732 | 221664.2 | 414.2046 | tri/oligopeptide |
| 113 | WPC | 417.2352 | 1 | 22.10395 | 1276204 | 416.2279 | tri/oligopeptide |
| 114 | WPC | 420.2094 | 1 | 2.464725 | 345128.3 | 419.2021 | tri/oligopeptide |
| 115 | WPC | 421.1932 | 1 | 2.2007 | 982449.5 | 420.1859 | tri/oligopeptide |
| 116 | WPC | 423.0647 | 1 | 1.697417 | 151401.7 | 422.0574 | tri/oligopeptide |
| 117 | WPC | 428.287 | 1 | 2.2007 | 204194.7 | 427.2797 | tri/oligopeptide |
| 118 | WPC | 429.272 | 1 | 18.61096 | 212246.2 | 428.2647 | tri/oligopeptide |
| 119 | WPC | 431.2332 | 1 | 20.35361 | 16663423 | 430.2259 | tri/oligopeptide |
| 120 | WPC | 433.0939 | 1 | 1.6744 | 153852.8 | 432.0866 | tri/oligopeptide |
| 121 | WPC | 433.1931 | 1 | 2.895875 | 4165481 | 432.1858 | tri/oligopeptide |
| 122 | WPC | 434.1884 | 1 | 2.047242 | 304482.7 | 433.1812 | tri/oligopeptide |
| 123 | WPC | 435.199 | 1 | 2.889092 | 122352.1 | 434.1918 | tri/oligopeptide |
| 124 | WPC | 444.2464 | 1 | 10.52273 | 418633.2 | 443.2392 | tri/oligopeptide |
| 125 | WPC | 445.2305 | 1 | 14.19772 | 246155.8 | 444.2232 | tri/oligopeptide |
| 126 | WPC | 445.2662 | 1 | 14.23255 | 2765783 | 444.2589 | tri/oligopeptide |
| 127 | WPC | 445.3027 | 1 | 18.99421 | 1416299 | 444.2955 | tri/oligopeptide |
| 128 | WPC | 223.6254 | 2 | 2.146283 | 110258.4 | 445.2363 | tri/oligopeptide |
| 129 | WPC | 446.2611 | 1 | 2.2007 | 383107.2 | 445.2538 | tri/oligopeptide |
| 130 | WPC | 223.6342 | 2 | 2.21185 | 375341 | 445.2539 | tri/oligopeptide |
| 131 | WPC | 449.2078 | 1 | 17.60869 | 129291.1 | 448.2005 | tri/oligopeptide |
| 132 | WPC | 229.0794 | 2 | 2.895875 | 130998 | 456.1443 | tri/oligopeptide |
| 133 | WPC | 461.3004 | 1 | 37.32523 | 108090.1 | 460.2932 | tri/oligopeptide |
| 134 | WPC | 463.0268 | 1 | 1.606508 | 263864.8 | 462.0195 | tri/oligopeptide |
| 135 | WPC | 463.2038 | 1 | 2.2007 | 112764.9 | 462.1965 | tri/oligopeptide |
| 136 | WPC | 236.0739 | 2 | 2.889092 | 150618 | 470.1332 | tri/oligopeptide |
| 137 | WPC | 472.2412 | 1 | 11.73022 | 435935.1 | 471.234 | tri/oligopeptide |
| 138 | WPC | 476.2906 | 1 | 20.35361 | 233771.2 | 475.2833 | tri/oligopeptide |
| 139 | WPC | 477.183 | 1 | 2.18985 | 156430.2 | 476.1758 | tri/oligopeptide |
| 140 | WPC | 478.2307 | 1 | 14.86646 | 522714.3 | 477.2235 | tri/oligopeptide |
| 141 | WPC | 479.3103 | 1 | 37.14109 | 173144 | 478.303 | tri/oligopeptide |
| 142 | WPC | 482.2254 | 1 | 9.694933 | 106457.7 | 481.2182 | tri/oligopeptide |
| 143 | WPC | 488.2466 | 1 | 2.18985 | 210189.7 | 487.2394 | tri/oligopeptide |
| 144 | WPC | 488.2517 | 1 | 19.14056 | 185115.9 | 487.2444 | tri/oligopeptide |
| 145 | WPC | 244.6581 | 2 | 14.26878 | 188683.5 | 487.3017 | tri/oligopeptide |
| 146 | WPC | 489.2559 | 1 | 17.18248 | 3579764 | 488.2487 | tri/oligopeptide |
| 147 | WPC | 491.2873 | 1 | 22.65285 | 110004.6 | 490.28 | tri/oligopeptide |
| 148 | WPC | 491.288 | 1 | 23.14529 | 120511.5 | 490.2807 | tri/oligopeptide |
| 149 | WPC | 492.2304 | 1 | 2.2007 | 149446.9 | 491.2231 | tri/oligopeptide |

| | | | | | | | |
|---|---|---|---|---|---|---|---|
| 150 | WPC | 249.5928 | 2 | 2.880642 | 104121 | 497.171 | tri/oligopeptide |
| 151 | WPC | 500.2884 | 1 | 25.20752 | 132652.1 | 499.2811 | tri/oligopeptide |
| 152 | WPC | 255.6069 | 2 | 20.35361 | 125040 | 509.1992 | tri/oligopeptide |
| 153 | WPC | 513.2676 | 1 | 13.79717 | 127727.1 | 512.2603 | tri/oligopeptide |
| 154 | WPC | 513.3046 | 1 | 19.66069 | 143752.3 | 512.2974 | tri/oligopeptide |
| 155 | WPC | 515.2464 | 1 | 2.576092 | 272501.3 | 514.2391 | tri/oligopeptide |
| 156 | WPC | 515.2914 | 1 | 37.12918 | 101147.7 | 514.2841 | tri/oligopeptide |
| 157 | WPC | 518.2467 | 1 | 4.178875 | 119791.5 | 517.2394 | tri/oligopeptide |
| 158 | WPC | 519.2309 | 1 | 8.928583 | 134416.1 | 518.2236 | tri/oligopeptide |
| 159 | WPC | 520.2781 | 1 | 21.18686 | 143863.4 | 519.2709 | tri/oligopeptide |
| 160 | WPC | 263.5939 | 2 | 15.67608 | 158150.4 | 525.1732 | tri/oligopeptide |
| 161 | WPC | 264.1748 | 2 | 35.26535 | 117309 | 526.335 | tri/oligopeptide |
| 162 | WPC | 265.1712 | 2 | 2.511267 | 1216070 | 528.3278 | tri/oligopeptide |
| 163 | WPC | 530.3298 | 1 | 2.2007 | 166324.8 | 529.3225 | tri/oligopeptide |
| 164 | WPC | 265.6687 | 2 | 2.2007 | 285344.6 | 529.3229 | tri/oligopeptide |
| 165 | WPC | 531.0142 | 1 | 1.595175 | 331868.6 | 530.0069 | tri/oligopeptide |
| 166 | WPC | 531.3871 | 1 | 37.84241 | 110336.2 | 530.3798 | tri/oligopeptide |
| 167 | WPC | 532.2251 | 1 | 2.2007 | 956207.7 | 531.2178 | tri/oligopeptide |
| 168 | WPC | 539.2359 | 1 | 8.921633 | 307519.3 | 538.2286 | tri/oligopeptide |
| 169 | WPC | 270.6017 | 2 | 17.47979 | 177534 | 539.1889 | tri/oligopeptide |
| 170 | WPC | 544.318 | 1 | 26.99236 | 103620.7 | 543.3107 | tri/oligopeptide |
| 171 | WPC | 545.2939 | 1 | 18.70931 | 1387714 | 544.2866 | tri/oligopeptide |
| 172 | WPC | 273.1507 | 2 | 18.69747 | 655049.1 | 544.2868 | tri/oligopeptide |
| 173 | WPC | 546.2783 | 1 | 22.25869 | 521518.6 | 545.271 | tri/oligopeptide |
| 174 | WPC | 274.1677 | 2 | 35.94308 | 117483.5 | 546.3209 | tri/oligopeptide |
| 175 | WPC | 276.1349 | 2 | 17.19903 | 128044.7 | 550.2553 | tri/oligopeptide |
| 176 | WPC | 553.3361 | 1 | 23.57533 | 1081491 | 552.3288 | tri/oligopeptide |
| 177 | WPC | 277.6244 | 2 | 17.18858 | 117237.7 | 553.2342 | tri/oligopeptide |
| 178 | WPC | 561.3251 | 1 | 2.392808 | 273559.9 | 560.3178 | tri/oligopeptide |
| 179 | WPC | 562.2734 | 1 | 11.38478 | 596542.6 | 561.2661 | tri/oligopeptide |
| 180 | WPC | 565.2106 | 1 | 2.012825 | 292933.6 | 564.2033 | tri/oligopeptide |
| 181 | WPC | 565.2623 | 1 | 16.60482 | 3907241 | 564.255 | tri/oligopeptide |
| 182 | WPC | 283.1349 | 2 | 16.61216 | 1082928 | 564.2552 | tri/oligopeptide |
| 183 | WPC | 285.1298 | 2 | 18.70931 | 242415.3 | 568.245 | tri/oligopeptide |
| 184 | WPC | 286.1582 | 2 | 2.179125 | 842452 | 570.3018 | tri/oligopeptide |
| 185 | WPC | 571.3091 | 1 | 2.18985 | 502056.4 | 570.3018 | tri/oligopeptide |
| 186 | WPC | 571.3105 | 1 | 16.07581 | 109766.3 | 570.3032 | tri/oligopeptide |
| 187 | WPC | 572.2767 | 1 | 19.16763 | 177884.1 | 571.2694 | tri/oligopeptide |
| 188 | WPC | 578.232 | 1 | 9.06615 | 411848.7 | 577.2247 | tri/oligopeptide |
| 189 | WPC | 292.1242 | 2 | 18.70931 | 106683.9 | 582.2339 | tri/oligopeptide |
| 190 | WPC | 584.303 | 1 | 2.18985 | 135506.5 | 583.2957 | tri/oligopeptide |
| 191 | WPC | 295.114 | 2 | 16.61216 | 291893.9 | 588.2135 | tri/oligopeptide |
| 192 | WPC | 590.2688 | 1 | 19.6081 | 417765.5 | 589.2615 | tri/oligopeptide |
| 193 | WPC | 597.2523 | 1 | 2.381492 | 108829.2 | 596.245 | tri/oligopeptide |
| 194 | WPC | 599.002 | 1 | 1.606508 | 297704.2 | 597.9947 | tri/oligopeptide |
| 195 | WPC | 300.1609 | 2 | 1.424158 | 187409.8 | 598.3073 | tri/oligopeptide |
| 196 | WPC | 302.1085 | 2 | 16.60482 | 260133.9 | 602.2025 | tri/oligopeptide |
| 197 | WPC | 607.3834 | 1 | 20.80124 | 1020570 | 606.3761 | tri/oligopeptide |
| 198 | WPC | 304.1954 | 2 | 20.80863 | 122171 | 606.3763 | tri/oligopeptide |
| 199 | WPC | 308.0823 | 2 | 2.168392 | 104088.3 | 614.15 | oligopeptide |

| | | | | | | | |
|---|---|---|---|---|---|---|---|
| 200 | WPC | 615.1576 | 1 | 2.18985 | 106149.4 | 614.1503 | oligopeptide |
| 201 | WPC | 310.0947 | 2 | 16.60482 | 102695.6 | 618.1748 | oligopeptide |
| 202 | WPC | 314.1798 | 2 | 26.73088 | 123082.8 | 626.3451 | oligopeptide |
| 203 | WPC | 627.3524 | 1 | 26.71047 | 428634 | 626.3451 | oligopeptide |
| 204 | WPC | 628.3317 | 1 | 19.14679 | 865849.2 | 627.3245 | oligopeptide |
| 205 | WPC | 314.6695 | 2 | 19.15264 | 218800.2 | 627.3245 | oligopeptide |
| 206 | WPC | 634.2482 | 1 | 14.49353 | 113192.2 | 633.2409 | oligopeptide |
| 207 | WPC | 640.2975 | 1 | 19.9362 | 159851.2 | 639.2902 | oligopeptide |
| 208 | WPC | 643.3318 | 1 | 20.73993 | 155273.2 | 642.3246 | oligopeptide |
| 209 | WPC | 323.1825 | 2 | 2.404683 | 216816.8 | 644.3504 | oligopeptide |
| 210 | WPC | 646.3048 | 1 | 2.279217 | 640774.8 | 645.2975 | oligopeptide |
| 211 | WPC | 323.6563 | 2 | 2.279217 | 225924.3 | 645.298 | oligopeptide |
| 212 | WPC | 652.3209 | 1 | 21.66973 | 499650.8 | 651.3136 | oligopeptide |
| 213 | WPC | 654.3842 | 1 | 21.82051 | 147026 | 653.3769 | oligopeptide |
| 214 | WPC | 328.1577 | 2 | 15.64802 | 203884.4 | 654.3008 | oligopeptide |
| 215 | WPC | 658.3422 | 1 | 20.44584 | 277949 | 657.3349 | oligopeptide |
| 216 | WPC | 659.2895 | 1 | 14.00398 | 325590.3 | 658.2822 | oligopeptide |
| 217 | WPC | 660.2839 | 1 | 2.18985 | 102110.3 | 659.2767 | oligopeptide |
| 218 | WPC | 666.9895 | 1 | 1.606508 | 296182.6 | 665.9822 | oligopeptide |
| 219 | WPC | 668.3623 | 1 | 20.00583 | 378059.8 | 667.355 | oligopeptide |
| 220 | WPC | 335.6351 | 2 | 2.279217 | 186496.1 | 669.2557 | oligopeptide |
| 221 | WPC | 678.347 | 1 | 22.66756 | 3929075 | 677.3398 | oligopeptide |
| 222 | WPC | 339.6773 | 2 | 22.67454 | 1237910 | 677.34 | oligopeptide |
| 223 | WPC | 342.701 | 2 | 16.28062 | 126824.3 | 683.3875 | oligopeptide |
| 224 | WPC | 348.1777 | 2 | 2.234092 | 134224 | 694.3408 | oligopeptide |
| 225 | WPC | 349.198 | 2 | 12.31427 | 692917.7 | 696.3814 | oligopeptide |
| 226 | WPC | 350.1688 | 2 | 2.168392 | 102643.7 | 698.323 | oligopeptide |
| 227 | WPC | 350.6622 | 2 | 22.66756 | 131181.7 | 699.3098 | oligopeptide |
| 228 | WPC | 701.3478 | 1 | 16.54638 | 1618184 | 700.3406 | oligopeptide |
| 229 | WPC | 351.1776 | 2 | 16.53589 | 1903700 | 700.3407 | oligopeptide |
| 230 | WPC | 351.6563 | 2 | 22.67454 | 706602.9 | 701.298 | oligopeptide |
| 231 | WPC | 702.3688 | 1 | 18.63017 | 134673.1 | 701.3615 | oligopeptide |
| 232 | WPC | 707.2228 | 1 | 1.697417 | 464747.2 | 706.2156 | oligopeptide |
| 233 | WPC | 356.6847 | 2 | 2.024292 | 330991.7 | 711.3549 | oligopeptide |
| 234 | WPC | 239.4362 | 3 | 22.67454 | 205965.4 | 715.2867 | oligopeptide |
| 235 | WPC | 358.6509 | 2 | 22.67454 | 333158.8 | 715.2872 | oligopeptide |
| 236 | WPC | 362.1624 | 2 | 16.54638 | 153103.2 | 722.3102 | oligopeptide |
| 237 | WPC | 363.1464 | 2 | 1.955792 | 1030446 | 724.2783 | oligopeptide |
| 238 | WPC | 363.1565 | 2 | 16.53589 | 842482.8 | 724.2985 | oligopeptide |
| 239 | WPC | 366.6369 | 2 | 22.67454 | 131593.4 | 731.2593 | oligopeptide |
| 240 | WPC | 734.9771 | 1 | 1.606508 | 245470.7 | 733.9699 | oligopeptide |
| 241 | WPC | 247.1031 | 3 | 16.55465 | 134166.9 | 738.2874 | oligopeptide |
| 242 | WPC | 370.1511 | 2 | 16.53589 | 394579.1 | 738.2877 | oligopeptide |
| 243 | WPC | 371.6317 | 2 | 18.01268 | 149229.9 | 741.2489 | oligopeptide |
| 244 | WPC | 374.6898 | 2 | 20.78791 | 213939.3 | 747.3651 | oligopeptide |
| 245 | WPC | 376.7003 | 2 | 2.222883 | 192112.4 | 751.3861 | oligopeptide |
| 246 | WPC | 377.6498 | 2 | 21.6631 | 102812.8 | 753.2851 | oligopeptide |
| 247 | WPC | 377.7298 | 2 | 29.30715 | 885810.8 | 753.445 | oligopeptide |
| 248 | WPC | 754.4523 | 1 | 29.29283 | 2322458 | 753.4451 | oligopeptide |
| 249 | WPC | 378.1371 | 2 | 16.53589 | 143219.4 | 754.2597 | oligopeptide |

| | | | | | | | |
|---|---|---|---|---|---|---|---|
| 250 | WPC | 765.1816 | 1 | 1.697417 | 100309.2 | 764.1744 | oligopeptide |
| 251 | WPC | 384.6574 | 2 | 22.66756 | 128519.1 | 767.3002 | oligopeptide |
| 252 | WPC | 769.4105 | 1 | 17.81463 | 498139.5 | 768.4032 | oligopeptide |
| 253 | WPC | 770.3676 | 1 | 2.18985 | 160709.5 | 769.3603 | oligopeptide |
| 254 | WPC | 385.6876 | 2 | 2.222883 | 304576.2 | 769.3606 | oligopeptide |
| 255 | WPC | 778.4486 | 1 | 23.56193 | 121264 | 777.4413 | oligopeptide |
| 256 | WPC | 389.728 | 2 | 23.56193 | 379864.2 | 777.4414 | oligopeptide |
| 257 | WPC | 392.6952 | 2 | 2.21185 | 196456.1 | 783.3758 | oligopeptide |
| 258 | WPC | 802.9646 | 1 | 1.595175 | 215114.1 | 801.9573 | oligopeptide |
| 259 | WPC | 804.3277 | 1 | 13.70475 | 2317512 | 803.3204 | oligopeptide |
| 260 | WPC | 402.6675 | 2 | 13.71081 | 202118.7 | 803.3205 | oligopeptide |
| 261 | WPC | 406.7113 | 2 | 14.1609 | 290124.1 | 811.408 | oligopeptide |
| 262 | WPC | 406.7115 | 2 | 8.7537 | 113522.6 | 811.4085 | oligopeptide |
| 263 | WPC | 408.2304 | 2 | 14.73601 | 117665.3 | 814.4462 | oligopeptide |
| 264 | WPC | 276.1786 | 3 | 15.63843 | 276222.5 | 825.5138 | oligopeptide |
| 265 | WPC | 414.1595 | 2 | 14.04863 | 124023.7 | 826.3044 | oligopeptide |
| 266 | WPC | 414.6465 | 2 | 13.70475 | 808464.5 | 827.2784 | oligopeptide |
| 267 | WPC | 419.7279 | 2 | 25.05807 | 550561.2 | 837.4413 | oligopeptide |
| 268 | WPC | 838.4487 | 1 | 25.05044 | 134848.3 | 837.4414 | oligopeptide |
| 269 | WPC | 421.6409 | 2 | 13.70475 | 187017.1 | 841.2673 | oligopeptide |
| 270 | WPC | 281.4298 | 3 | 13.73775 | 179417.3 | 841.2676 | oligopeptide |
| 271 | WPC | 422.1383 | 2 | 2.482167 | 105587.4 | 842.2621 | oligopeptide |
| 272 | WPC | 424.1478 | 2 | 2.435583 | 127803.2 | 846.281 | oligopeptide |
| 273 | WPC | 853.4408 | 1 | 2.234092 | 101980.4 | 852.4335 | oligopeptide |
| 274 | WPC | 427.2241 | 2 | 2.279217 | 1398397 | 852.4337 | oligopeptide |
| 275 | WPC | 428.6348 | 2 | 12.11498 | 228419.1 | 855.255 | oligopeptide |
| 276 | WPC | 870.9519 | 1 | 1.606508 | 146408.4 | 869.9446 | oligopeptide |
| 277 | WPC | 438.1863 | 2 | 15.55887 | 378076.6 | 874.358 | oligopeptide |
| 278 | WPC | 875.3653 | 1 | 15.55887 | 463579.5 | 874.358 | oligopeptide |
| 279 | WPC | 442.2209 | 2 | 16.27213 | 2648898 | 882.4272 | oligopeptide |
| 280 | WPC | 442.2492 | 2 | 21.53531 | 151686.5 | 882.4839 | oligopeptide |
| 281 | WPC | 445.2459 | 2 | 30.22242 | 715547.7 | 888.4773 | oligopeptide |
| 282 | WPC | 889.485 | 1 | 30.21413 | 374161.1 | 888.4778 | oligopeptide |
| 283 | WPC | 448.2114 | 2 | 2.168392 | 204759.5 | 894.4083 | oligopeptide |
| 284 | WPC | 449.2378 | 2 | 2.904842 | 255080.1 | 896.4611 | oligopeptide |
| 285 | WPC | 450.1653 | 2 | 15.55887 | 262727.3 | 898.316 | oligopeptide |
| 286 | WPC | 450.6574 | 2 | 18.57114 | 206451.7 | 899.3003 | oligopeptide |
| 287 | WPC | 457.6376 | 2 | 13.04848 | 336701.2 | 913.2607 | oligopeptide |
| 288 | WPC | 457.6654 | 2 | 19.98653 | 243660.1 | 913.3162 | oligopeptide |
| 289 | WPC | 921.3709 | 1 | 18.95907 | 412430.1 | 920.3636 | oligopeptide |
| 290 | WPC | 462.2141 | 2 | 1.432192 | 179743.4 | 922.4136 | oligopeptide |
| 291 | WPC | 466.7143 | 2 | 2.168392 | 184586.6 | 931.4141 | oligopeptide |
| 292 | WPC | 938.9391 | 1 | 1.606508 | 102819.6 | 937.9318 | oligopeptide |
| 293 | WPC | 471.1978 | 2 | 22.17535 | 124159.9 | 940.3811 | oligopeptide |
| 294 | WPC | 473.168 | 2 | 18.95907 | 146478.8 | 944.3214 | oligopeptide |
| 295 | WPC | 485.2353 | 2 | 2.21185 | 118105 | 968.456 | oligopeptide |
| 296 | WPC | 974.8145 | 1 | 1.617742 | 207403.6 | 973.8072 | oligopeptide |
| 297 | WPC | 990.789 | 1 | 1.628758 | 177840.7 | 989.7817 | oligopeptide |
| 298 | WPC | 499.2518 | 2 | 16.08898 | 201061 | 996.4891 | oligopeptide |
| 299 | WPC | 507.1721 | 2 | 17.6531 | 105729.4 | 1012.33 | oligopeptide |

| | | | | | | | |
|---|---|---|---|---|---|---|---|
| 300 | WPC | 514.958 | 2 | 1.595175 | 105251.2 | 1027.901 | oligopeptide |
| 301 | WPC | 521.2777 | 2 | 17.21345 | 1825116 | 1040.541 | oligopeptide |
| 302 | WPC | 1042.802 | 1 | 1.617742 | 112199.1 | 1041.794 | oligopeptide |
| 303 | WPC | 523.7867 | 2 | 23.17221 | 292908.7 | 1045.559 | oligopeptide |
| 304 | WPC | 527.7392 | 2 | 18.99943 | 397963.3 | 1053.464 | oligopeptide |
| 305 | WPC | 529.7732 | 2 | 15.09943 | 570236.9 | 1057.532 | oligopeptide |
| 306 | WPC | 534.282 | 2 | 22.13133 | 281260.2 | 1066.549 | oligopeptide |
| 307 | WPC | 548.9517 | 2 | 1.595175 | 108882.7 | 1095.889 | oligopeptide |
| 308 | WPC | 550.7942 | 2 | 28.70863 | 915629.7 | 1099.574 | oligopeptide |
| 309 | WPC | 1100.581 | 1 | 28.72263 | 112359 | 1099.574 | oligopeptide |
| 310 | WPC | 559.2505 | 2 | 20.96831 | 109252.6 | 1116.486 | oligopeptide |
| 311 | WPC | 380.1813 | 3 | 28.70863 | 107459.7 | 1137.522 | oligopeptide |
| 312 | WPC | 570.2834 | 2 | 14.40415 | 242813.4 | 1138.552 | oligopeptide |
| 313 | WPC | 582.9457 | 2 | 1.595175 | 111749 | 1163.877 | oligopeptide |
| 314 | WPC | 590.9329 | 2 | 1.606508 | 107728 | 1179.851 | oligopeptide |
| 315 | WPC | 611.3502 | 2 | 30.66709 | 247428.1 | 1220.686 | oligopeptide |
| 316 | WPC | 616.9397 | 2 | 1.606508 | 106238.3 | 1231.865 | oligopeptide |
| 317 | WPC | 623.2974 | 2 | 18.85794 | 1793319 | 1244.58 | oligopeptide |
| 318 | WPC | 624.9266 | 2 | 1.606508 | 123266.4 | 1247.839 | oligopeptide |
| 319 | WPC | 423.8535 | 3 | 18.85794 | 117316 | 1268.539 | oligopeptide |
| 320 | WPC | 644.7018 | 2 | 16.91353 | 157499.9 | 1287.389 | oligopeptide |
| 321 | WPC | 650.9333 | 2 | 1.595175 | 110958.2 | 1299.852 | oligopeptide |
| 322 | WPC | 658.9205 | 2 | 1.606508 | 109726.8 | 1315.826 | oligopeptide |
| 323 | WPC | 660.8836 | 2 | 31.72243 | 1153159 | 1319.753 | oligopeptide |
| 324 | WPC | 671.8743 | 2 | 31.72243 | 128154.4 | 1341.734 | oligopeptide |
| 325 | WPC | 453.5738 | 3 | 31.72243 | 111530.4 | 1357.7 | oligopeptide |
| 326 | WPC | 683.3317 | 2 | 16.83296 | 1638555 | 1364.649 | oligopeptide |
| 327 | WPC | 686.2282 | 2 | 16.44577 | 389863.6 | 1370.442 | oligopeptide |
| 328 | WPC | 691.8265 | 2 | 14.43147 | 113440.9 | 1381.638 | oligopeptide |
| 329 | WPC | 692.9142 | 2 | 1.606508 | 122083.3 | 1383.814 | oligopeptide |
| 330 | WPC | 463.8764 | 3 | 16.83296 | 168314.5 | 1388.607 | oligopeptide |
| 331 | WPC | 726.9083 | 2 | 1.606508 | 106652.8 | 1451.802 | oligopeptide |
| 332 | WPC | 503.5076 | 3 | 14.09834 | 393668.2 | 1507.501 | oligopeptide |
| 333 | WPC | 760.9016 | 2 | 1.606508 | 112147.4 | 1519.789 | oligopeptide |
| 334 | WPC | 785.3513 | 2 | 18.36856 | 1290617 | 1568.688 | oligopeptide |
| 335 | WPC | 794.8955 | 2 | 1.606508 | 110887.6 | 1587.776 | oligopeptide |
| 336 | WPC | 531.8893 | 3 | 18.38533 | 218926.1 | 1592.646 | oligopeptide |
| 337 | WPC | 828.8889 | 2 | 1.617742 | 102917.6 | 1655.763 | oligopeptide |

| ID | Protein | mz | charge | retentiont | raw.abund | mass | type | Match |
|---|---|---|---|---|---|---|---|---|
| 2 | AWPC | 231.1708 | 1 | 5.273683 | 1904546 | 230.1635 | di/tripepti | Y |
| 3 | AWPC | 231.1708 | 1 | 9.576117 | 3487293 | 230.1635 | di/tripepti | Y |
| 4 | AWPC | 231.1708 | 1 | 11.98742 | 2068211 | 230.1636 | di/tripepti | Y |
| 5 | AWPC | 233.1497 | 1 | 2.196983 | 149747.6 | 232.1424 | di/tripepti | Y |
| 6 | AWPC | 233.1498 | 1 | 2.737975 | 101116.8 | 232.1426 | di/tripepti | Y |
| 7 | AWPC | 239.1027 | 1 | 2.185483 | 238919.6 | 238.0954 | di/tripepti | Y |
| 8 | AWPC | 239.1029 | 1 | 2.420408 | 158459.9 | 238.0956 | di/tripepti | Y |
| 10 | AWPC | 245.1133 | 1 | 1.9787 | 533983.1 | 244.106 | di/tripepti | Y |
| 11 | AWPC | 245.1863 | 1 | 16.41699 | 1651426 | 244.179 | di/tripepti | Y |
| 12 | AWPC | 245.1865 | 1 | 15.88533 | 416531.8 | 244.1793 | di/tripepti | Y |
| 13 | AWPC | 245.1866 | 1 | 14.54718 | 2284778 | 244.1793 | di/tripepti | Y |
| 14 | AWPC | 245.1866 | 1 | 17.6797 | 234215.9 | 244.1793 | di/tripepti | Y |
| 15 | AWPC | 246.145 | 1 | 2.119825 | 581353.2 | 245.1377 | di/tripepti | Y |
| 16 | AWPC | 246.1451 | 1 | 2.628433 | 335570.8 | 245.1378 | di/tripepti | Y |
| 17 | AWPC | 246.1453 | 1 | 3.307208 | 193706.1 | 245.138 | di/tripepti | Y |
| 19 | AWPC | 247.1113 | 1 | 2.381475 | 315852.3 | 246.1041 | di/tripepti | Y |
| 20 | AWPC | 247.129 | 1 | 2.185483 | 501091.9 | 246.1217 | di/tripepti | Y |
| 21 | AWPC | 247.1291 | 1 | 3.090867 | 1133145 | 246.1218 | di/tripepti | Y |
| 22 | AWPC | 247.1293 | 1 | 4.684425 | 1086499 | 246.122 | di/tripepti | Y |
| 24 | AWPC | 250.178 | 1 | 34.8633 | 727241.7 | 249.1707 | di/tripepti | Y |
| 25 | AWPC | 250.1781 | 1 | 37.30648 | 241072.2 | 249.1708 | di/tripepti | Y |
| 26 | AWPC | 252.9628 | 1 | 4.835875 | 119741.2 | 251.9555 | di/tripepti | Y |
| 28 | AWPC | 253.1188 | 1 | 9.2984 | 153476 | 252.1115 | di/tripepti | Y |
| 30 | AWPC | 260.1607 | 1 | 2.185483 | 1241601 | 259.1534 | di/tripepti | Y |
| 31 | AWPC | 260.1608 | 1 | 2.628433 | 684324 | 259.1535 | di/tripepti | Y |
| 32 | AWPC | 260.161 | 1 | 3.647733 | 540904.4 | 259.1537 | di/tripepti | Y |
| 33 | AWPC | 260.1971 | 1 | 2.185483 | 119775.7 | 259.1898 | di/tripepti | Y |
| 34 | AWPC | 261.1447 | 1 | 2.185483 | 354452.3 | 260.1375 | di/tripepti | Y |
| 35 | AWPC | 261.1449 | 1 | 3.221208 | 570225.9 | 260.1376 | di/tripepti | Y |
| 36 | AWPC | 261.145 | 1 | 5.452467 | 650286.2 | 260.1378 | di/tripepti | Y |
| 37 | AWPC | 262.1399 | 1 | 2.090442 | 198865.8 | 261.1327 | di/tripepti | Y |
| 38 | AWPC | 263.143 | 1 | 11.49235 | 1396537 | 262.1357 | di/tripepti | Y |
| 39 | AWPC | 263.1967 | 1 | 2.185483 | 127844.8 | 262.1895 | di/tripepti | Y |
| 41 | AWPC | 267.1343 | 1 | 2.402025 | 129472.4 | 266.127 | di/tripepti | Y |
| 42 | AWPC | 269.1608 | 1 | 2.150667 | 2107226 | 268.1536 | di/tripepti | Y |
| 43 | AWPC | 274.1875 | 1 | 1.9787 | 241761.8 | 273.1802 | di/tripepti | Y |
| 44 | AWPC | 277.1032 | 1 | 1.857775 | 177563.1 | 276.096 | di/tripepti | Y |
| 45 | AWPC | 281.1134 | 1 | 2.551392 | 715060.2 | 280.1062 | di/tripepti | Y |
| 47 | AWPC | 288.1925 | 1 | 16.07473 | 218052.5 | 287.1852 | di/tripepti | Y |
| 48 | AWPC | 290.1714 | 1 | 2.677217 | 1816774 | 289.1641 | di/tripepti | Y |
| 50 | AWPC | 294.1451 | 1 | 2.324908 | 219033.8 | 293.1379 | di/tripepti | Y |
| 51 | AWPC | 295.1295 | 1 | 3.184558 | 231856.9 | 294.1222 | di/tripepti | Y |
| 52 | AWPC | 295.1657 | 1 | 16.41699 | 422982.9 | 294.1585 | di/tripepti | Y |
| 53 | AWPC | 300.192 | 1 | 2.573417 | 5790506 | 299.1847 | di/tripepti | Y |
| 54 | AWPC | 300.1928 | 1 | 15.12667 | 238374.1 | 299.1855 | di/tripepti | Y |
| 56 | AWPC | 302.2082 | 1 | 15.14083 | 321660.8 | 301.2009 | tri/oligope | Y |
| 57 | AWPC | 302.2084 | 1 | 19.86491 | 120421.9 | 301.2011 | tri/oligope | Y |
| 58 | AWPC | 303.1666 | 1 | 2.196983 | 224317.9 | 302.1594 | tri/oligope | Y |
| 59 | AWPC | 304.1506 | 1 | 2.402025 | 919383.1 | 303.1433 | tri/oligope | Y |

| | | | | | | | | |
|---|---|---|---|---|---|---|---|---|
| 60 | AWPC | 304.1509 | 1 | 4.878617 | 1044883 | 303.1437 | tri/oligope | Y |
| 62 | AWPC | 316.2237 | 1 | 13.87716 | 909218.2 | 315.2164 | tri/oligope | Y |
| 63 | AWPC | 318.1664 | 1 | 2.520325 | 1004240 | 317.1591 | tri/oligope | Y |
| 64 | AWPC | 318.1665 | 1 | 2.324908 | 2330787 | 317.1592 | tri/oligope | Y |
| 66 | AWPC | 318.1821 | 1 | 22.56183 | 1680626 | 317.1748 | tri/oligope | Y |
| 69 | AWPC | 319.1615 | 1 | 2.162533 | 390958 | 318.1542 | tri/oligope | Y |
| 71 | AWPC | 330.2396 | 1 | 18.41157 | 314976.7 | 329.2324 | tri/oligope | Y |
| 72 | AWPC | 331.1613 | 1 | 2.078558 | 675098.2 | 330.1541 | tri/oligope | Y |
| 73 | AWPC | 331.1657 | 1 | 2.795458 | 156774.5 | 330.1584 | tri/oligope | Y |
| 74 | AWPC | 331.2851 | 1 | 38.45992 | 160207.6 | 330.2778 | tri/oligope | Y |
| 75 | AWPC | 332.1818 | 1 | 2.599758 | 3512877 | 331.1745 | tri/oligope | Y |
| 76 | AWPC | 332.2186 | 1 | 3.034567 | 1654703 | 331.2113 | tri/oligope | Y |
| 77 | AWPC | 332.2187 | 1 | 3.361925 | 2609927 | 331.2114 | tri/oligope | Y |
| 80 | AWPC | 334.1612 | 1 | 2.208867 | 123167.6 | 333.1539 | tri/oligope | Y |
| 81 | AWPC | 334.1613 | 1 | 3.184558 | 1265698 | 333.1541 | tri/oligope | Y |
| 82 | AWPC | 342.2399 | 1 | 21.81455 | 1036480 | 341.2326 | tri/oligope | Y |
| 83 | AWPC | 343.1239 | 1 | 1.720125 | 178373.7 | 342.1167 | tri/oligope | Y |
| 84 | AWPC | 344.1826 | 1 | 5.57985 | 156489.5 | 343.1753 | tri/oligope | Y |
| 85 | AWPC | 344.2555 | 1 | 21.54573 | 1062855 | 343.2482 | tri/oligope | Y |
| 86 | AWPC | 344.2556 | 1 | 20.18665 | 107348.8 | 343.2483 | tri/oligope | Y |
| 87 | AWPC | 349.1723 | 1 | 1.857775 | 110858.2 | 348.165 | tri/oligope | Y |
| 88 | AWPC | 350.1197 | 1 | 1.827958 | 484545.1 | 349.1125 | tri/oligope | Y |
| 89 | AWPC | 358.2712 | 1 | 24.02033 | 197413.3 | 357.2639 | tri/oligope | Y |
| 90 | AWPC | 359.2293 | 1 | 2.185483 | 444567.2 | 358.2221 | tri/oligope | Y |
| 91 | AWPC | 360.196 | 1 | 18.72603 | 1823035 | 359.1888 | tri/oligope | Y |
| 92 | AWPC | 360.2134 | 1 | 2.701133 | 145799.1 | 359.2061 | tri/oligope | Y |
| 93 | AWPC | 360.2137 | 1 | 13.63674 | 992379 | 359.2064 | tri/oligope | Y |
| 94 | AWPC | 363.1555 | 1 | 2.185483 | 132566.2 | 362.1482 | tri/oligope | Y |
| 95 | AWPC | 365.1366 | 1 | 37.81322 | 104981.3 | 364.1293 | tri/oligope | Y |
| 96 | AWPC | 366.214 | 1 | 2.185483 | 257472.6 | 365.2067 | tri/oligope | Y |
| 97 | AWPC | 374.2297 | 1 | 19.33054 | 645905 | 373.2224 | tri/oligope | Y |
| 99 | AWPC | 376.1467 | 1 | 1.720125 | 112945.9 | 375.1395 | tri/oligope | Y |
| 100 | AWPC | 376.1719 | 1 | 2.196983 | 591415 | 375.1646 | tri/oligope | Y |
| 102 | AWPC | 381.0801 | 1 | 1.674667 | 122802.6 | 380.0728 | tri/oligope | Y |
| 103 | AWPC | 387.2246 | 1 | 14.00868 | 7860160 | 386.2173 | tri/oligope | Y |
| 104 | AWPC | 389.2306 | 1 | 13.9912 | 194084.4 | 388.2233 | tri/oligope | Y |
| 105 | AWPC | 390.1876 | 1 | 2.313533 | 213222.5 | 389.1803 | tri/oligope | Y |
| 106 | AWPC | 395.0394 | 1 | 1.574158 | 210659 | 394.0321 | tri/oligope | Y |
| 107 | AWPC | 399.2612 | 1 | 18.53898 | 305607.4 | 398.254 | tri/oligope | Y |
| 109 | AWPC | 402.2354 | 1 | 11.35857 | 1794719 | 401.2281 | tri/oligope | Y |
| 111 | AWPC | 405.199 | 1 | 4.148108 | 162586.7 | 404.1918 | tri/oligope | Y |
| 112 | AWPC | 415.2118 | 1 | 36.47435 | 323153.5 | 414.2046 | tri/oligope | Y |
| 113 | AWPC | 417.2352 | 1 | 22.09608 | 549731.4 | 416.2279 | tri/oligope | Y |
| 114 | AWPC | 420.2094 | 1 | 2.456142 | 118240 | 419.2021 | tri/oligope | Y |
| 115 | AWPC | 421.1932 | 1 | 2.196983 | 695644.3 | 420.1859 | tri/oligope | Y |
| 117 | AWPC | 428.287 | 1 | 2.196983 | 156737.2 | 427.2797 | tri/oligope | Y |
| 118 | AWPC | 429.272 | 1 | 18.59953 | 102461.8 | 428.2647 | tri/oligope | Y |
| 119 | AWPC | 431.2332 | 1 | 20.34427 | 11620926 | 430.2259 | tri/oligope | Y |
| 121 | AWPC | 433.1931 | 1 | 2.85885 | 2829224 | 432.1858 | tri/oligope | Y |
| 122 | AWPC | 434.1884 | 1 | 2.045575 | 169781.7 | 433.1812 | tri/oligope | Y |

| | | | | | | | | |
|---|---|---|---|---|---|---|---|---|
| 124 | AWPC | 444.2464 | 1 | 10.45416 | 490691.3 | 443.2392 | tri/oligope | Y |
| 125 | AWPC | 445.2305 | 1 | 14.20219 | 541808.9 | 444.2232 | tri/oligope | Y |
| 126 | AWPC | 445.2662 | 1 | 14.23033 | 1305963 | 444.2589 | tri/oligope | Y |
| 127 | AWPC | 445.3027 | 1 | 18.98931 | 842344.3 | 444.2955 | tri/oligope | Y |
| 128 | AWPC | 223.6254 | 2 | 2.138775 | 140440.6 | 445.2363 | tri/oligope | Y |
| 129 | AWPC | 446.2611 | 1 | 2.196983 | 256960.4 | 445.2538 | tri/oligope | Y |
| 130 | AWPC | 223.6342 | 2 | 2.208867 | 250865.4 | 445.2539 | tri/oligope | Y |
| 131 | AWPC | 449.2078 | 1 | 17.59765 | 115730.9 | 448.2005 | tri/oligope | Y |
| 133 | AWPC | 461.3004 | 1 | 37.31842 | 107679.2 | 460.2932 | tri/oligope | Y |
| 134 | AWPC | 463.0268 | 1 | 1.607808 | 274127.4 | 462.0195 | tri/oligope | Y |
| 136 | AWPC | 236.0739 | 2 | 2.852875 | 102948.4 | 470.1332 | tri/oligope | Y |
| 137 | AWPC | 472.2412 | 1 | 11.76702 | 525032.6 | 471.234 | tri/oligope | Y |
| 138 | AWPC | 476.2906 | 1 | 20.34427 | 135478.9 | 475.2833 | tri/oligope | Y |
| 139 | AWPC | 477.183 | 1 | 2.185483 | 107041.6 | 476.1758 | tri/oligope | Y |
| 140 | AWPC | 478.2307 | 1 | 14.85351 | 422627.8 | 477.2235 | tri/oligope | Y |
| 141 | AWPC | 479.3103 | 1 | 37.13739 | 177468.5 | 478.303 | tri/oligope | Y |
| 144 | AWPC | 488.2517 | 1 | 19.13527 | 134694.2 | 487.2444 | tri/oligope | Y |
| 146 | AWPC | 489.2559 | 1 | 17.17401 | 1500020 | 488.2487 | tri/oligope | Y |
| 147 | AWPC | 491.2873 | 1 | 22.64707 | 149780.7 | 490.28 | tri/oligope | Y |
| 149 | AWPC | 492.2304 | 1 | 2.196983 | 114016.4 | 491.2231 | tri/oligope | Y |
| 154 | AWPC | 513.3046 | 1 | 19.64597 | 164203.6 | 512.2974 | tri/oligope | Y |
| 155 | AWPC | 515.2464 | 1 | 2.558717 | 269343.6 | 514.2391 | tri/oligope | Y |
| 156 | AWPC | 515.2914 | 1 | 37.12387 | 101219.7 | 514.2841 | tri/oligope | Y |
| 159 | AWPC | 520.2781 | 1 | 21.17131 | 104791 | 519.2709 | tri/oligope | Y |
| 161 | AWPC | 264.1748 | 2 | 35.27389 | 115760.8 | 526.335 | tri/oligope | Y |
| 162 | AWPC | 265.1712 | 2 | 2.501658 | 341798.6 | 528.3278 | tri/oligope | Y |
| 164 | AWPC | 265.6687 | 2 | 2.196983 | 138299.5 | 529.3229 | tri/oligope | Y |
| 165 | AWPC | 531.0142 | 1 | 1.596708 | 334776.1 | 530.0069 | tri/oligope | Y |
| 166 | AWPC | 531.3871 | 1 | 37.83449 | 109085.3 | 530.3798 | tri/oligope | Y |
| 167 | AWPC | 532.2251 | 1 | 2.196983 | 700561.9 | 531.2178 | tri/oligope | Y |
| 168 | AWPC | 539.2359 | 1 | 8.930575 | 231457 | 538.2286 | tri/oligope | Y |
| 171 | AWPC | 545.2939 | 1 | 18.70098 | 398680.8 | 544.2866 | tri/oligope | Y |
| 172 | AWPC | 273.1507 | 2 | 18.68868 | 189785.6 | 544.2868 | tri/oligope | Y |
| 173 | AWPC | 546.2783 | 1 | 22.25314 | 102459.5 | 545.271 | tri/oligope | Y |
| 174 | AWPC | 274.1677 | 2 | 35.93865 | 114180.2 | 546.3209 | tri/oligope | Y |
| 176 | AWPC | 553.3361 | 1 | 23.57234 | 1373638 | 552.3288 | tri/oligope | Y |
| 179 | AWPC | 562.2734 | 1 | 11.34012 | 178821.2 | 561.2661 | tri/oligope | Y |
| 180 | AWPC | 565.2106 | 1 | 2.011192 | 191006.5 | 564.2033 | tri/oligope | Y |
| 181 | AWPC | 565.2623 | 1 | 16.59621 | 4189464 | 564.255 | tri/oligope | Y |
| 182 | AWPC | 283.1349 | 2 | 16.60434 | 1205912 | 564.2552 | tri/oligope | Y |
| 184 | AWPC | 286.1582 | 2 | 2.174033 | 514510.7 | 570.3018 | tri/oligope | Y |
| 185 | AWPC | 571.3091 | 1 | 2.185483 | 332368.2 | 570.3018 | tri/oligope | Y |
| 186 | AWPC | 571.3105 | 1 | 16.06048 | 112694.3 | 570.3032 | tri/oligope | Y |
| 188 | AWPC | 578.232 | 1 | 9.07675 | 174934.4 | 577.2247 | tri/oligope | Y |
| 190 | AWPC | 584.303 | 1 | 2.185483 | 117259 | 583.2957 | tri/oligope | Y |
| 191 | AWPC | 295.114 | 2 | 16.60434 | 332666.2 | 588.2135 | tri/oligope | Y |
| 192 | AWPC | 590.2688 | 1 | 19.60093 | 145205.4 | 589.2615 | tri/oligope | Y |
| 194 | AWPC | 599.002 | 1 | 1.607808 | 320494.4 | 597.9947 | tri/oligope | Y |
| 195 | AWPC | 300.1609 | 2 | 1.423325 | 157682.3 | 598.3073 | tri/oligope | Y |
| 196 | AWPC | 302.1085 | 2 | 16.59621 | 295941.2 | 602.2025 | tri/oligope | Y |

| | | | | | | | | |
|---|---|---|---|---|---|---|---|---|
| 197 | AWPC | 607.3834 | 1 | 20.79872 | 1389651 | 606.3761 | tri/oligope | Y |
| 198 | AWPC | 304.1954 | 2 | 20.80518 | 167911.7 | 606.3763 | tri/oligope | Y |
| 201 | AWPC | 310.0947 | 2 | 16.59621 | 116601.4 | 618.1748 | oligopepti( | Y |
| 202 | AWPC | 314.1798 | 2 | 26.72738 | 171703.5 | 626.3451 | oligopepti( | Y |
| 203 | AWPC | 627.3524 | 1 | 26.70038 | 599130 | 626.3451 | oligopepti( | Y |
| 204 | AWPC | 628.3317 | 1 | 19.14271 | 736826.5 | 627.3245 | oligopepti( | Y |
| 205 | AWPC | 314.6695 | 2 | 19.15015 | 186067.7 | 627.3245 | oligopepti( | Y |
| 207 | AWPC | 640.2975 | 1 | 19.93092 | 145322.3 | 639.2902 | oligopepti( | Y |
| 208 | AWPC | 643.3318 | 1 | 20.73205 | 110562.9 | 642.3246 | oligopepti( | Y |
| 210 | AWPC | 646.3048 | 1 | 2.279333 | 200445.1 | 645.2975 | oligopepti( | Y |
| 212 | AWPC | 652.3209 | 1 | 21.66942 | 192740.3 | 651.3136 | oligopepti( | Y |
| 214 | AWPC | 328.1577 | 2 | 15.62716 | 235569 | 654.3008 | oligopepti( | Y |
| 215 | AWPC | 658.3422 | 1 | 20.43847 | 370504.2 | 657.3349 | oligopepti( | Y |
| 216 | AWPC | 659.2895 | 1 | 14.01757 | 183256.3 | 658.2822 | oligopepti( | Y |
| 218 | AWPC | 666.9895 | 1 | 1.607808 | 298406.9 | 665.9822 | oligopepti( | Y |
| 219 | AWPC | 668.3623 | 1 | 20.00098 | 428177.6 | 667.355 | oligopepti( | Y |
| 221 | AWPC | 678.347 | 1 | 22.66536 | 1410889 | 677.3398 | oligopepti( | Y |
| 222 | AWPC | 339.6773 | 2 | 22.67218 | 465712.6 | 677.34 | oligopepti( | Y |
| 224 | AWPC | 348.1777 | 2 | 2.232583 | 173009.7 | 694.3408 | oligopepti( | Y |
| 225 | AWPC | 349.198 | 2 | 12.22493 | 482964.5 | 696.3814 | oligopepti( | Y |
| 228 | AWPC | 701.3478 | 1 | 16.53653 | 555605.9 | 700.3406 | oligopepti( | Y |
| 229 | AWPC | 351.1776 | 2 | 16.52618 | 696887.4 | 700.3407 | oligopepti( | Y |
| 230 | AWPC | 351.6563 | 2 | 22.67218 | 267837 | 701.298 | oligopepti( | Y |
| 233 | AWPC | 356.6847 | 2 | 2.02265 | 332281.8 | 711.3549 | oligopepti( | Y |
| 235 | AWPC | 358.6509 | 2 | 22.67218 | 125669.6 | 715.2872 | oligopepti( | Y |
| 237 | AWPC | 363.1464 | 2 | 1.955233 | 654970.3 | 724.2783 | oligopepti( | Y |
| 238 | AWPC | 363.1565 | 2 | 16.52618 | 306530.2 | 724.2985 | oligopepti( | Y |
| 240 | AWPC | 734.9771 | 1 | 1.607808 | 257477.4 | 733.9699 | oligopepti( | Y |
| 242 | AWPC | 370.1511 | 2 | 16.52618 | 140387.6 | 738.2877 | oligopepti( | Y |
| 244 | AWPC | 374.6898 | 2 | 20.78355 | 265838.7 | 747.3651 | oligopepti( | Y |
| 245 | AWPC | 376.7003 | 2 | 2.220583 | 190545.2 | 751.3861 | oligopepti( | Y |
| 247 | AWPC | 377.7298 | 2 | 29.30544 | 607690.8 | 753.445 | oligopepti( | Y |
| 248 | AWPC | 754.4523 | 1 | 29.29083 | 1577483 | 753.4451 | oligopepti( | Y |
| 252 | AWPC | 769.4105 | 1 | 17.80113 | 247805.7 | 768.4032 | oligopepti( | Y |
| 253 | AWPC | 770.3676 | 1 | 2.185483 | 127025.5 | 769.3603 | oligopepti( | Y |
| 254 | AWPC | 385.6876 | 2 | 2.220583 | 230381.3 | 769.3606 | oligopepti( | Y |
| 256 | AWPC | 389.728 | 2 | 23.55654 | 240793.2 | 777.4414 | oligopepti( | Y |
| 257 | AWPC | 392.6952 | 2 | 2.208867 | 148575.3 | 783.3758 | oligopepti( | Y |
| 258 | AWPC | 802.9646 | 1 | 1.596708 | 222248.6 | 801.9573 | oligopepti( | Y |
| 259 | AWPC | 804.3277 | 1 | 13.694 | 1214111 | 803.3204 | oligopepti( | Y |
| 260 | AWPC | 402.6675 | 2 | 13.7027 | 108278.5 | 803.3205 | oligopepti( | Y |
| 261 | AWPC | 406.7113 | 2 | 14.17374 | 101766.7 | 811.408 | oligopepti( | Y |
| 266 | AWPC | 414.6465 | 2 | 13.694 | 438256 | 827.2784 | oligopepti( | Y |
| 267 | AWPC | 419.7279 | 2 | 25.0595 | 292635.2 | 837.4413 | oligopepti( | Y |
| 274 | AWPC | 427.2241 | 2 | 2.279333 | 962064.3 | 852.4337 | oligopepti( | Y |
| 275 | AWPC | 428.6348 | 2 | 12.05813 | 135454.3 | 855.255 | oligopepti( | Y |
| 276 | AWPC | 870.9519 | 1 | 1.607808 | 154687 | 869.9446 | oligopepti( | Y |
| 277 | AWPC | 438.1863 | 2 | 15.54144 | 229065.9 | 874.358 | oligopepti( | Y |
| 278 | AWPC | 875.3653 | 1 | 15.54144 | 269396 | 874.358 | oligopepti( | Y |
| 279 | AWPC | 442.2209 | 2 | 16.2593 | 1780225 | 882.4272 | oligopepti( | Y |

| | | | | | | | | |
|---|---|---|---|---|---|---|---|---|
| 280 | AWPC | 442.2492 | 2 | 21.52652 | 113515.2 | 882.4839 | oligopepti | Y |
| 281 | AWPC | 445.2459 | 2 | 30.22275 | 959019.8 | 888.4773 | oligopepti | Y |
| 282 | AWPC | 889.485 | 1 | 30.21368 | 506720.4 | 888.4778 | oligopepti | Y |
| 283 | AWPC | 448.2114 | 2 | 2.162533 | 122875 | 894.4083 | oligopepti | Y |
| 285 | AWPC | 450.1653 | 2 | 15.54144 | 159143.8 | 898.316 | oligopepti | Y |
| 287 | AWPC | 457.6376 | 2 | 12.95815 | 254773 | 913.2607 | oligopepti | Y |
| 289 | AWPC | 921.3709 | 1 | 18.95233 | 196472.6 | 920.3636 | oligopepti | Y |
| 290 | AWPC | 462.2141 | 2 | 1.431408 | 168902.6 | 922.4136 | oligopepti | Y |
| 296 | AWPC | 974.8145 | 1 | 1.618625 | 207220.9 | 973.8072 | oligopepti | Y |
| 297 | AWPC | 990.789 | 1 | 1.62945 | 166308.8 | 989.7817 | oligopepti | Y |
| 300 | AWPC | 514.958 | 2 | 1.596708 | 107008.4 | 1027.901 | oligopepti | Y |
| 301 | AWPC | 521.2777 | 2 | 17.20263 | 425634.1 | 1040.541 | oligopepti | Y |
| 302 | AWPC | 1042.802 | 1 | 1.618625 | 111663.7 | 1041.794 | oligopepti | Y |
| 305 | AWPC | 529.7732 | 2 | 15.07979 | 405671.7 | 1057.532 | oligopepti | Y |
| 307 | AWPC | 548.9517 | 2 | 1.596708 | 116785.2 | 1095.889 | oligopepti | Y |
| 308 | AWPC | 550.7942 | 2 | 28.71115 | 245930.1 | 1099.574 | oligopepti | Y |
| 312 | AWPC | 570.2834 | 2 | 14.40222 | 127940.4 | 1138.552 | oligopepti | Y |
| 313 | AWPC | 582.9457 | 2 | 1.596708 | 117309 | 1163.877 | oligopepti | Y |
| 314 | AWPC | 590.9329 | 2 | 1.607808 | 120091.2 | 1179.851 | oligopepti | Y |
| 316 | AWPC | 616.9397 | 2 | 1.607808 | 122188 | 1231.865 | oligopepti | Y |
| 317 | AWPC | 623.2974 | 2 | 18.84983 | 239925.4 | 1244.58 | oligopepti | Y |
| 318 | AWPC | 624.9266 | 2 | 1.607808 | 134932.7 | 1247.839 | oligopepti | Y |
| 321 | AWPC | 650.9333 | 2 | 1.596708 | 110882 | 1299.852 | oligopepti | Y |
| 322 | AWPC | 658.9205 | 2 | 1.607808 | 122920.3 | 1315.826 | oligopepti | Y |
| 323 | AWPC | 660.8836 | 2 | 31.73455 | 624798.9 | 1319.753 | oligopepti | Y |
| 326 | AWPC | 683.3317 | 2 | 16.82833 | 571515.6 | 1364.649 | oligopepti | Y |
| 327 | AWPC | 686.2282 | 2 | 16.43336 | 288667.5 | 1370.442 | oligopepti | Y |
| 329 | AWPC | 692.9142 | 2 | 1.607808 | 120101.2 | 1383.814 | oligopepti | Y |
| 331 | AWPC | 726.9083 | 2 | 1.607808 | 114294.6 | 1451.802 | oligopepti | Y |
| 332 | AWPC | 503.5076 | 3 | 14.11873 | 140108.3 | 1507.501 | oligopepti | Y |
| 333 | AWPC | 760.9016 | 2 | 1.607808 | 118725 | 1519.789 | oligopepti | Y |
| 334 | AWPC | 785.3513 | 2 | 18.35963 | 433492.2 | 1568.688 | oligopepti | Y |
| 335 | AWPC | 794.8955 | 2 | 1.607808 | 108313.6 | 1587.776 | oligopepti | Y |
| 337 | AWPC | 828.8889 | 2 | 1.618625 | 102009.8 | 1655.763 | oligopepti | Y |
| 338 | AWPC | 229.1552 | 1 | 11.91688 | 121366.2 | 228.148 | di/tripepti | N |
| 339 | AWPC | 262.1035 | 1 | 1.720125 | 102998.6 | 261.0963 | di/tripepti | N |
| 340 | AWPC | 269.1134 | 1 | 2.185483 | 119958.8 | 268.1061 | di/tripepti | N |
| 341 | AWPC | 276.1556 | 1 | 2.208867 | 112810.7 | 275.1483 | di/tripepti | N |
| 342 | AWPC | 313.176 | 1 | 2.185483 | 117614.6 | 312.1687 | tri/oligope | N |
| 343 | AWPC | 328.2242 | 1 | 15.87005 | 121495.2 | 327.217 | tri/oligope | N |
| 344 | AWPC | 328.234 | 1 | 15.48093 | 100780.4 | 327.2267 | tri/oligope | N |
| 345 | AWPC | 332.219 | 1 | 19.44008 | 153304.3 | 331.2118 | tri/oligope | N |
| 346 | AWPC | 334.1405 | 1 | 13.78616 | 142047.1 | 333.1333 | tri/oligope | N |
| 347 | AWPC | 342.24 | 1 | 20.11797 | 163910.2 | 341.2327 | tri/oligope | N |
| 348 | AWPC | 344.1826 | 1 | 4.819033 | 135656.2 | 343.1753 | tri/oligope | N |
| 349 | AWPC | 346.1981 | 1 | 9.598242 | 143731.4 | 345.1908 | tri/oligope | N |
| 350 | AWPC | 359.2298 | 1 | 14.24358 | 158602.9 | 358.2225 | tri/oligope | N |
| 351 | AWPC | 360.1882 | 1 | 2.056958 | 103364.7 | 359.1809 | tri/oligope | N |
| 352 | AWPC | 360.2141 | 1 | 20.61665 | 118805.8 | 359.2068 | tri/oligope | N |
| 353 | AWPC | 366.167 | 1 | 11.55632 | 130143.5 | 365.1597 | tri/oligope | N |

| | | | | | | | | |
|---|---|---|---|---|---|---|---|---|
| 354 | AWPC | 398.1561 | 1 | 2.208867 | 109932.8 | 397.1489 | tri/oligope | N |
| 355 | AWPC | 443.2139 | 1 | 2.196983 | 113091.1 | 442.2066 | tri/oligope | N |
| 356 | AWPC | 447.2281 | 1 | 13.71706 | 115276.4 | 446.2208 | tri/oligope | N |
| 357 | AWPC | 226.1462 | 2 | 38.45992 | 105229.5 | 450.2778 | tri/oligope | N |
| 358 | AWPC | 475.2236 | 1 | 20.62424 | 117275.7 | 474.2163 | tri/oligope | N |
| 359 | AWPC | 482.3239 | 1 | 37.18689 | 108301.4 | 481.3166 | tri/oligope | N |
| 360 | AWPC | 489.2719 | 1 | 22.1195 | 265583.6 | 488.2646 | tri/oligope | N |
| 361 | AWPC | 505.3036 | 1 | 24.77722 | 154657.1 | 504.2963 | tri/oligope | N |
| 362 | AWPC | 515.2837 | 1 | 15.3988 | 279580.5 | 514.2764 | tri/oligope | N |
| 363 | AWPC | 524.3719 | 1 | 38.22528 | 100147.7 | 523.3646 | tri/oligope | N |
| 364 | AWPC | 526.3143 | 1 | 36.96188 | 169334.5 | 525.307 | tri/oligope | N |
| 365 | AWPC | 530.299 | 1 | 27.48286 | 240105.7 | 529.2917 | tri/oligope | N |
| 366 | AWPC | 269.6293 | 2 | 2.511142 | 279813.3 | 537.2441 | tri/oligope | N |
| 367 | AWPC | 560.2573 | 1 | 2.812683 | 121798.1 | 559.25 | tri/oligope | N |
| 368 | AWPC | 560.2946 | 1 | 16.79592 | 155692.6 | 559.2873 | tri/oligope | N |
| 369 | AWPC | 288.6796 | 2 | 22.6551 | 145991 | 575.3446 | tri/oligope | N |
| 370 | AWPC | 585.326 | 1 | 18.40532 | 100702.6 | 584.3188 | tri/oligope | N |
| 371 | AWPC | 590.2788 | 1 | 2.587317 | 118720.2 | 589.2716 | tri/oligope | N |
| 372 | AWPC | 318.117 | 2 | 16.59621 | 106652.5 | 634.2194 | oligopepti | N |
| 373 | AWPC | 326.7062 | 2 | 20.80518 | 111565.6 | 651.3979 | oligopepti | N |
| 374 | AWPC | 329.6748 | 2 | 20.44598 | 122488.1 | 657.335 | oligopepti | N |
| 375 | AWPC | 341.654 | 2 | 20.44598 | 102843.9 | 681.2933 | oligopepti | N |
| 376 | AWPC | 342.7006 | 2 | 2.196983 | 167182.1 | 683.3866 | oligopepti | N |
| 377 | AWPC | 343.1083 | 2 | 16.59621 | 104361.2 | 684.2021 | oligopepti | N |
| 378 | AWPC | 345.2224 | 2 | 27.67829 | 131256.2 | 688.4302 | oligopepti | N |
| 379 | AWPC | 379.7094 | 2 | 27.11153 | 109722.3 | 757.4042 | oligopepti | N |
| 380 | AWPC | 407.2225 | 2 | 13.20707 | 372110.4 | 812.4304 | oligopepti | N |
| 381 | AWPC | 867.3724 | 1 | 36.47435 | 101430.5 | 866.3651 | oligopepti | N |
| 382 | AWPC | 457.2248 | 2 | 30.20714 | 114774.5 | 912.4351 | oligopepti | N |
| 383 | AWPC | 464.2196 | 2 | 30.20714 | 105427.2 | 926.4247 | oligopepti | N |
| 384 | AWPC | 485.2676 | 2 | 19.41461 | 142236.9 | 968.5207 | oligopepti | N |
| 385 | AWPC | 503.2729 | 2 | 21.74028 | 105311 | 1004.531 | oligopepti | N |
| 386 | AWPC | 684.9273 | 2 | 1.607808 | 109136.6 | 1367.84 | oligopepti | N |

| ID | Protein | mz | charge | retentiont | raw.abund | mass | type | Match |
|---|---|---|---|---|---|---|---|---|
| 3 | Egg | 231.1708 | 1 | 9.488808 | 647117.7 | 230.1635 | di/tripepti | Y |
| 5 | Egg | 233.1499 | 1 | 2.459775 | 140593.6 | 232.1426 | di/tripepti | Y |
| 10 | Egg | 245.1136 | 1 | 2.113563 | 407009.6 | 244.1063 | di/tripepti | Y |
| 11 | Egg | 245.1864 | 1 | 16.87238 | 354337.9 | 244.1791 | di/tripepti | Y |
| 11 | Egg | 245.1865 | 1 | 16.65326 | 713334.5 | 244.1792 | di/tripepti | Y |
| 12 | Egg | 245.1868 | 1 | 16.14035 | 107853.7 | 244.1796 | di/tripepti | Y |
| 13 | Egg | 245.1865 | 1 | 14.88878 | 555098.5 | 244.1793 | di/tripepti | Y |
| 14 | Egg | 245.1867 | 1 | 17.88473 | 143449.7 | 244.1794 | di/tripepti | Y |
| 15 | Egg | 246.1452 | 1 | 2.351929 | 587220.1 | 245.138 | di/tripepti | Y |
| 17 | Egg | 246.1455 | 1 | 3.112496 | 293727.7 | 245.1382 | di/tripepti | Y |
| 20 | Egg | 247.1293 | 1 | 2.369725 | 394111.2 | 246.122 | di/tripepti | Y |
| 21 | Egg | 247.1297 | 1 | 3.511192 | 126537.4 | 246.1224 | di/tripepti | Y |
| 22 | Egg | 247.1297 | 1 | 4.883296 | 125044.1 | 246.1224 | di/tripepti | Y |
| 24 | Egg | 250.1782 | 1 | 34.98545 | 989152.1 | 249.1709 | di/tripepti | Y |
| 24 | Egg | 250.1784 | 1 | 34.5071 | 118855.7 | 249.1711 | di/tripepti | Y |
| 27 | Egg | 253.1189 | 1 | 2.636454 | 171898.1 | 252.1116 | di/tripepti | Y |
| 31 | Egg | 260.161 | 1 | 2.441813 | 370699.2 | 259.1537 | di/tripepti | Y |
| 31 | Egg | 260.1611 | 1 | 2.851121 | 306393 | 259.1539 | di/tripepti | Y |
| 32 | Egg | 260.1612 | 1 | 3.439021 | 181998 | 259.1539 | di/tripepti | Y |
| 33 | Egg | 260.1973 | 1 | 2.422696 | 255141.3 | 259.19 | di/tripepti | Y |
| 34 | Egg | 261.1451 | 1 | 2.441813 | 164964.7 | 260.1378 | di/tripepti | Y |
| 35 | Egg | 261.1451 | 1 | 3.282875 | 115951.7 | 260.1379 | di/tripepti | Y |
| 37 | Egg | 262.1402 | 1 | 2.184104 | 217444.3 | 261.133 | di/tripepti | Y |
| 40 | Egg | 265.1556 | 1 | 16.49057 | 109540.5 | 264.1484 | di/tripepti | Y |
| 42 | Egg | 269.1612 | 1 | 2.422696 | 211595.8 | 268.154 | di/tripepti | Y |
| 43 | Egg | 274.1878 | 1 | 2.025771 | 153394 | 273.1805 | di/tripepti | Y |
| 44 | Egg | 277.1035 | 1 | 1.948421 | 410373.7 | 276.0962 | di/tripepti | Y |
| 48 | Egg | 290.1717 | 1 | 2.680971 | 104287.9 | 289.1644 | di/tripepti | Y |
| 48 | Egg | 290.1718 | 1 | 3.156933 | 159062.1 | 289.1645 | di/tripepti | Y |
| 52 | Egg | 295.166 | 1 | 16.64235 | 373969.1 | 294.1587 | di/tripepti | Y |
| 56 | Egg | 302.2082 | 1 | 15.45535 | 465214 | 301.201 | tri/oligope | Y |
| 57 | Egg | 302.2081 | 1 | 19.95505 | 1480326 | 301.2009 | tri/oligope | Y |
| 58 | Egg | 303.1669 | 1 | 2.422696 | 453670.7 | 302.1596 | tri/oligope | Y |
| 59 | Egg | 304.1509 | 1 | 2.441813 | 156193.7 | 303.1437 | tri/oligope | Y |
| 63 | Egg | 318.1669 | 1 | 2.680971 | 146912.9 | 317.1596 | tri/oligope | Y |
| 66 | Egg | 318.1822 | 1 | 22.63143 | 402724.2 | 317.175 | tri/oligope | Y |
| 69 | Egg | 319.1619 | 1 | 2.141692 | 121485.3 | 318.1546 | tri/oligope | Y |
| 71 | Egg | 330.2398 | 1 | 18.56749 | 181715 | 329.2326 | tri/oligope | Y |
| 72 | Egg | 331.1617 | 1 | 2.334708 | 120000.9 | 330.1544 | tri/oligope | Y |
| 74 | Egg | 331.2851 | 1 | 38.21228 | 367917.6 | 330.2778 | tri/oligope | Y |
| 78 | Egg | 333.1411 | 1 | 1.97625 | 120045.3 | 332.1338 | tri/oligope | Y |
| 80 | Egg | 334.1615 | 1 | 2.351929 | 708292.8 | 333.1542 | tri/oligope | Y |
| 82 | Egg | 342.2398 | 1 | 21.35965 | 446875.6 | 341.2325 | tri/oligope | Y |
| 95 | Egg | 365.1368 | 1 | 37.61193 | 115452.8 | 364.1296 | tri/oligope | Y |
| 112 | Egg | 415.2122 | 1 | 36.27107 | 204098.5 | 414.2049 | tri/oligope | Y |
| 133 | Egg | 461.3008 | 1 | 37.40978 | 156877.5 | 460.2935 | tri/oligope | Y |
| 141 | Egg | 479.3108 | 1 | 36.86001 | 136703.6 | 478.3035 | tri/oligope | Y |
| 144 | Egg | 488.2515 | 1 | 19.22683 | 192323.6 | 487.2443 | tri/oligope | Y |
| 151 | Egg | 500.2884 | 1 | 25.21228 | 118274.9 | 499.2812 | tri/oligope | Y |

| | | | | | | | | |
|---|---|---|---|---|---|---|---|---|
| 154 | Egg | 513.3045 | 1 | 19.78508 | 245122.1 | 512.2972 | tri/oligope | Y |
| 156 | Egg | 515.2912 | 1 | 36.72567 | 161844.9 | 514.2839 | tri/oligope | Y |
| 159 | Egg | 520.2781 | 1 | 21.22253 | 111620.6 | 519.2708 | tri/oligope | Y |
| 161 | Egg | 264.1749 | 2 | 34.94 | 172257.5 | 526.3352 | tri/oligope | Y |
| 165 | Egg | 531.0146 | 1 | 1.595308 | 131810.9 | 530.0074 | tri/oligope | Y |
| 166 | Egg | 531.3875 | 1 | 37.63103 | 117303.9 | 530.3802 | tri/oligope | Y |
| 174 | Egg | 274.1678 | 2 | 35.72245 | 148287.6 | 546.3211 | tri/oligope | Y |
| 208 | Egg | 643.3319 | 1 | 20.7716 | 166196.6 | 642.3246 | oligopepti | Y |
| 218 | Egg | 666.9901 | 1 | 1.582842 | 120058.7 | 665.9829 | oligopepti | Y |
| 240 | Egg | 734.9776 | 1 | 1.582842 | 105516.7 | 733.9703 | oligopepti | Y |
| 297 | Egg | 990.7898 | 1 | 1.607821 | 238549.1 | 989.7825 | oligopepti | Y |
| 338 | Egg | 223.1081 | 1 | 9.328596 | 122462.4 | 222.1008 | dipeptide | N |
| 339 | Egg | 223.1082 | 1 | 8.756054 | 180595.1 | 222.1009 | dipeptide | N |
| 340 | Egg | 223.1085 | 1 | 6.137638 | 276546.2 | 222.1012 | dipeptide | N |
| 341 | Egg | 226.952 | 1 | 1.539604 | 298007.3 | 225.9447 | di/tripepti | N |
| 342 | Egg | 228.1961 | 1 | 34.98545 | 2436024 | 227.1888 | di/tripepti | N |
| 343 | Egg | 228.1963 | 1 | 34.49176 | 504525.6 | 227.1891 | di/tripepti | N |
| 344 | Egg | 229.1552 | 1 | 7.86515 | 117688.6 | 228.1479 | di/tripepti | N |
| 345 | Egg | 229.1552 | 1 | 8.630525 | 120074.5 | 228.1479 | di/tripepti | N |
| 346 | Egg | 229.1553 | 1 | 12.86783 | 212998.9 | 228.148 | di/tripepti | N |
| 347 | Egg | 229.1554 | 1 | 6.156467 | 201637.8 | 228.1482 | di/tripepti | N |
| 348 | Egg | 231.1709 | 1 | 12.83317 | 1231873 | 230.1636 | di/tripepti | N |
| 349 | Egg | 231.1709 | 1 | 11.22379 | 400662.1 | 230.1637 | di/tripepti | N |
| 350 | Egg | 231.1709 | 1 | 10.71672 | 221939.2 | 230.1637 | di/tripepti | N |
| 351 | Egg | 231.171 | 1 | 13.19696 | 333681.5 | 230.1637 | di/tripepti | N |
| 352 | Egg | 231.1711 | 1 | 6.514313 | 460594.7 | 230.1639 | di/tripepti | N |
| 353 | Egg | 232.1295 | 1 | 2.422696 | 487036.3 | 231.1222 | di/tripepti | N |
| 354 | Egg | 232.1295 | 1 | 1.937763 | 192601.6 | 231.1223 | di/tripepti | N |
| 355 | Egg | 233.1136 | 1 | 2.141692 | 182744.7 | 232.1063 | di/tripepti | N |
| 356 | Egg | 233.1136 | 1 | 2.459775 | 1077728 | 232.1063 | di/tripepti | N |
| 357 | Egg | 237.0908 | 1 | 2.404792 | 122608.3 | 236.0835 | di/tripepti | N |
| 358 | Egg | 245.1516 | 1 | 36.32401 | 101706 | 244.1443 | di/tripepti | N |
| 359 | Egg | 246.1453 | 1 | 5.439663 | 165804.7 | 245.138 | di/tripepti | N |
| 360 | Egg | 246.1816 | 1 | 1.937763 | 353879.6 | 245.1744 | di/tripepti | N |
| 361 | Egg | 247.1296 | 1 | 5.723488 | 204767.8 | 246.1223 | di/tripepti | N |
| 362 | Egg | 248.1245 | 1 | 2.071563 | 146734 | 247.1172 | di/tripepti | N |
| 363 | Egg | 249.1085 | 1 | 1.925296 | 165941.8 | 248.1012 | di/tripepti | N |
| 364 | Egg | 249.1271 | 1 | 7.513367 | 182253.8 | 248.1198 | di/tripepti | N |
| 365 | Egg | 249.1274 | 1 | 3.304954 | 180246.7 | 248.1201 | di/tripepti | N |
| 366 | Egg | 252.963 | 1 | 8.253229 | 133296.2 | 251.9557 | di/tripepti | N |
| 367 | Egg | 253.1188 | 1 | 10.20286 | 233143.5 | 252.1115 | di/tripepti | N |
| 368 | Egg | 255.1456 | 1 | 1.925296 | 125452.5 | 254.1383 | di/tripepti | N |
| 369 | Egg | 260.1609 | 1 | 5.747083 | 145905.8 | 259.1537 | di/tripepti | N |
| 370 | Egg | 260.1611 | 1 | 3.191608 | 170873.6 | 259.1539 | di/tripepti | N |
| 371 | Egg | 261.1448 | 1 | 8.563958 | 108526.4 | 260.1375 | di/tripepti | N |
| 372 | Egg | 261.145 | 1 | 7.744004 | 115965.5 | 260.1377 | di/tripepti | N |
| 373 | Egg | 261.1454 | 1 | 6.447513 | 250603.4 | 260.1381 | di/tripepti | N |
| 374 | Egg | 263.1396 | 1 | 14.97629 | 434492.3 | 262.1323 | di/tripepti | N |
| 375 | Egg | 263.1398 | 1 | 14.78566 | 416013.8 | 262.1325 | di/tripepti | N |
| 376 | Egg | 263.143 | 1 | 12.41498 | 1039331 | 262.1358 | di/tripepti | N |

| | | | | | | | | |
|---|---|---|---|---|---|---|---|---|
| 377 | Egg | 263.1432 | 1 | 13.7246 | 142990.5 | 262.1359 | di/tripepti | N |
| 378 | Egg | 263.1969 | 1 | 2.703242 | 192239.8 | 262.1896 | di/tripepti | N |
| 379 | Egg | 276.1196 | 1 | 1.843308 | 119800.1 | 275.1123 | di/tripepti | N |
| 380 | Egg | 276.1353 | 1 | 17.3745 | 105052.8 | 275.128 | di/tripepti | N |
| 381 | Egg | 276.156 | 1 | 2.441813 | 102746.1 | 275.1487 | di/tripepti | N |
| 382 | Egg | 276.1563 | 1 | 4.487554 | 147229.1 | 275.149 | di/tripepti | N |
| 383 | Egg | 279.1709 | 1 | 20.02722 | 256947.2 | 278.1636 | di/tripepti | N |
| 384 | Egg | 280.1298 | 1 | 10.78588 | 111207.5 | 279.1226 | di/tripepti | N |
| 385 | Egg | 281.1501 | 1 | 10.02999 | 182673 | 280.1429 | di/tripepti | N |
| 386 | Egg | 281.1506 | 1 | 8.721442 | 167735.5 | 280.1434 | di/tripepti | N |
| 387 | Egg | 290.1352 | 1 | 2.441813 | 252845 | 289.1279 | di/tripepti | N |
| 388 | Egg | 290.1717 | 1 | 7.554021 | 100872.7 | 289.1644 | di/tripepti | N |
| 389 | Egg | 290.1719 | 1 | 7.9355 | 108408.5 | 289.1646 | di/tripepti | N |
| 390 | Egg | 295.1292 | 1 | 8.524492 | 132116.7 | 294.1219 | di/tripepti | N |
| 391 | Egg | 295.1294 | 1 | 8.984521 | 122862.2 | 294.1221 | di/tripepti | N |
| 392 | Egg | 295.1296 | 1 | 13.02811 | 306754.3 | 294.1224 | di/tripepti | N |
| 393 | Egg | 295.1299 | 1 | 13.48731 | 100706.1 | 294.1227 | di/tripepti | N |
| 394 | Egg | 295.13 | 1 | 3.776679 | 263360.7 | 294.1227 | di/tripepti | N |
| 395 | Egg | 302.1715 | 1 | 7.657067 | 139564.3 | 301.1642 | tri/oligope | N |
| 396 | Egg | 302.1716 | 1 | 2.441813 | 241226.1 | 301.1643 | tri/oligope | N |
| 397 | Egg | 302.1717 | 1 | 6.207183 | 209817.9 | 301.1644 | tri/oligope | N |
| 398 | Egg | 302.2084 | 1 | 18.89318 | 623114.8 | 301.2011 | tri/oligope | N |
| 399 | Egg | 303.1672 | 1 | 4.251367 | 315897.2 | 302.1599 | tri/oligope | N |
| 400 | Egg | 304.1872 | 1 | 6.137638 | 281816 | 303.1799 | tri/oligope | N |
| 401 | Egg | 304.1873 | 1 | 2.441813 | 362980.3 | 303.18 | tri/oligope | N |
| 402 | Egg | 304.1875 | 1 | 7.366421 | 464493.6 | 303.1802 | tri/oligope | N |
| 403 | Egg | 306.1301 | 1 | 1.97625 | 354910.7 | 305.1228 | tri/oligope | N |
| 404 | Egg | 310.3111 | 1 | 36.70178 | 240558.4 | 309.3038 | tri/oligope | N |
| 405 | Egg | 314.2083 | 1 | 13.7246 | 342465.7 | 313.2011 | tri/oligope | N |
| 406 | Egg | 317.1827 | 1 | 2.587379 | 591801.8 | 316.1754 | tri/oligope | N |
| 407 | Egg | 317.1827 | 1 | 5.395513 | 287398.7 | 316.1754 | tri/oligope | N |
| 408 | Egg | 317.1827 | 1 | 6.502104 | 288005.9 | 316.1754 | tri/oligope | N |
| 409 | Egg | 317.219 | 1 | 4.856913 | 104203.9 | 316.2118 | tri/oligope | N |
| 410 | Egg | 318.149 | 1 | 12.38957 | 2041942 | 317.1418 | tri/oligope | N |
| 411 | Egg | 318.1667 | 1 | 5.618017 | 108897.8 | 317.1594 | tri/oligope | N |
| 412 | Egg | 318.1669 | 1 | 10.51695 | 121197.4 | 317.1596 | tri/oligope | N |
| 413 | Egg | 320.1459 | 1 | 2.351929 | 145514.5 | 319.1386 | tri/oligope | N |
| 414 | Egg | 322.1437 | 1 | 6.104546 | 144827 | 321.1364 | tri/oligope | N |
| 415 | Egg | 322.144 | 1 | 2.680971 | 260315.8 | 321.1367 | tri/oligope | N |
| 416 | Egg | 331.1657 | 1 | 5.730913 | 314918.9 | 330.1584 | tri/oligope | N |
| 417 | Egg | 331.1987 | 1 | 11.90143 | 224286.8 | 330.1914 | tri/oligope | N |
| 418 | Egg | 332.2189 | 1 | 15.40528 | 171466.6 | 331.2116 | tri/oligope | N |
| 419 | Egg | 332.2191 | 1 | 18.77735 | 122374.4 | 331.2118 | tri/oligope | N |
| 420 | Egg | 332.2191 | 1 | 19.60275 | 133504 | 331.2118 | tri/oligope | N |
| 421 | Egg | 333.1775 | 1 | 2.422696 | 547044.5 | 332.1702 | tri/oligope | N |
| 422 | Egg | 333.1777 | 1 | 2.96645 | 334210.3 | 332.1704 | tri/oligope | N |
| 423 | Egg | 334.1406 | 1 | 14.24543 | 804958.7 | 333.1333 | tri/oligope | N |
| 424 | Egg | 334.1804 | 1 | 15.85984 | 351842.5 | 333.1731 | tri/oligope | N |
| 425 | Egg | 334.1805 | 1 | 16.07056 | 142089.7 | 333.1732 | tri/oligope | N |
| 426 | Egg | 335.1567 | 1 | 1.960788 | 190202.8 | 334.1495 | tri/oligope | N |

| | | | | | | | | |
|---|---|---|---|---|---|---|---|---|
| 427 | Egg | 338.3423 | 1 | 36.4099 | 8711581 | 337.3351 | tri/oligope | N |
| 428 | Egg | 338.3427 | 1 | 38.97075 | 3861406 | 337.3354 | tri/oligope | N |
| 429 | Egg | 340.3582 | 1 | 36.4099 | 108478.9 | 339.3509 | tri/oligope | N |
| 430 | Egg | 343.2964 | 1 | 35.28562 | 106005.2 | 342.2892 | tri/oligope | N |
| 431 | Egg | 345.2252 | 1 | 2.254725 | 120071.1 | 344.218 | tri/oligope | N |
| 432 | Egg | 345.2505 | 1 | 2.441813 | 122538.1 | 344.2432 | tri/oligope | N |
| 433 | Egg | 347.1932 | 1 | 9.957804 | 159357.2 | 346.1859 | tri/oligope | N |
| 434 | Egg | 347.1935 | 1 | 6.170188 | 144576.4 | 346.1862 | tri/oligope | N |
| 435 | Egg | 347.1936 | 1 | 4.457533 | 472819.4 | 346.1863 | tri/oligope | N |
| 436 | Egg | 347.2297 | 1 | 2.098179 | 127201.4 | 346.2224 | tri/oligope | N |
| 437 | Egg | 348.1409 | 1 | 2.254725 | 168049.3 | 347.1337 | tri/oligope | N |
| 438 | Egg | 348.1772 | 1 | 5.889554 | 218477.7 | 347.1699 | tri/oligope | N |
| 439 | Egg | 348.1773 | 1 | 6.799275 | 221076.6 | 347.17 | tri/oligope | N |
| 440 | Egg | 348.1774 | 1 | 2.561363 | 441361.6 | 347.1701 | tri/oligope | N |
| 441 | Egg | 348.1777 | 1 | 3.08725 | 100367.1 | 347.1704 | tri/oligope | N |
| 442 | Egg | 352.1515 | 1 | 12.65745 | 133403 | 351.1442 | tri/oligope | N |
| 443 | Egg | 358.1984 | 1 | 15.88562 | 1020130 | 357.1911 | tri/oligope | N |
| 444 | Egg | 359.2298 | 1 | 9.589308 | 170369.7 | 358.2225 | tri/oligope | N |
| 445 | Egg | 359.2301 | 1 | 3.050596 | 147851.1 | 358.2228 | tri/oligope | N |
| 446 | Egg | 360.3243 | 1 | 36.4099 | 244299.3 | 359.317 | tri/oligope | N |
| 447 | Egg | 360.3247 | 1 | 38.97075 | 226332.1 | 359.3175 | tri/oligope | N |
| 448 | Egg | 361.1724 | 1 | 5.114638 | 208020.7 | 360.1652 | tri/oligope | N |
| 449 | Egg | 361.1726 | 1 | 2.441813 | 261223.3 | 360.1653 | tri/oligope | N |
| 450 | Egg | 361.1727 | 1 | 2.808975 | 168548.7 | 360.1654 | tri/oligope | N |
| 451 | Egg | 361.1728 | 1 | 10.22242 | 217711.9 | 360.1656 | tri/oligope | N |
| 452 | Egg | 361.2357 | 1 | 37.11418 | 154780.4 | 360.2284 | tri/oligope | N |
| 453 | Egg | 363.1554 | 1 | 3.021488 | 155745.4 | 362.1481 | tri/oligope | N |
| 454 | Egg | 364.1358 | 1 | 2.141692 | 245839.6 | 363.1285 | tri/oligope | N |
| 455 | Egg | 366.3739 | 1 | 36.71975 | 137774.2 | 365.3666 | tri/oligope | N |
| 456 | Egg | 368.3532 | 1 | 39.04783 | 210097.1 | 367.346 | tri/oligope | N |
| 457 | Egg | 374.204 | 1 | 3.430229 | 144672.5 | 373.1967 | tri/oligope | N |
| 458 | Egg | 374.2041 | 1 | 2.441813 | 206944.4 | 373.1969 | tri/oligope | N |
| 459 | Egg | 374.2297 | 1 | 14.87214 | 191768.9 | 373.2224 | tri/oligope | N |
| 460 | Egg | 376.2244 | 1 | 25.40078 | 264727 | 375.2171 | tri/oligope | N |
| 461 | Egg | 378.17 | 1 | 8.837529 | 126590.6 | 377.1627 | tri/oligope | N |
| 462 | Egg | 378.1705 | 1 | 2.939775 | 135346.2 | 377.1632 | tri/oligope | N |
| 463 | Egg | 380.1828 | 1 | 20.58785 | 250338.7 | 379.1755 | tri/oligope | N |
| 464 | Egg | 383.4005 | 1 | 36.00488 | 262363.1 | 382.3932 | tri/oligope | N |
| 465 | Egg | 384.3116 | 1 | 37.12183 | 114065.6 | 383.3043 | tri/oligope | N |
| 466 | Egg | 390.199 | 1 | 2.441813 | 118680.3 | 389.1917 | tri/oligope | N |
| 467 | Egg | 391.1623 | 1 | 14.86355 | 176056.9 | 390.155 | tri/oligope | N |
| 468 | Egg | 391.183 | 1 | 5.470129 | 141462.8 | 390.1757 | tri/oligope | N |
| 469 | Egg | 391.1836 | 1 | 3.220267 | 138385.8 | 390.1763 | tri/oligope | N |
| 470 | Egg | 391.2023 | 1 | 18.982 | 186990.6 | 390.195 | tri/oligope | N |
| 471 | Egg | 391.2194 | 1 | 7.799954 | 109173.6 | 390.2121 | tri/oligope | N |
| 472 | Egg | 391.2856 | 1 | 0.957508 | 337660.6 | 390.2783 | tri/oligope | N |
| 473 | Egg | 394.1809 | 1 | 23.55302 | 407135.5 | 393.1736 | tri/oligope | N |
| 474 | Egg | 398.2337 | 1 | 39.47645 | 145393.2 | 397.2265 | tri/oligope | N |
| 475 | Egg | 399.225 | 1 | 14.86355 | 153885.2 | 398.2177 | tri/oligope | N |
| 476 | Egg | 403.2307 | 1 | 5.843033 | 146328.3 | 402.2234 | tri/oligope | N |

| | | | | | | | | |
|---|---|---|---|---|---|---|---|---|
| 477 | Egg | 403.2308 | 1 | 2.441813 | 324037.8 | 402.2235 | tri/oligope | N |
| 478 | Egg | 405.2145 | 1 | 19.02448 | 286818.3 | 404.2072 | tri/oligope | N |
| 479 | Egg | 405.2619 | 1 | 37.0692 | 353579.3 | 404.2546 | tri/oligope | N |
| 480 | Egg | 407.1415 | 1 | 2.025771 | 138737.6 | 406.1342 | tri/oligope | N |
| 481 | Egg | 407.178 | 1 | 2.404792 | 159043 | 406.1707 | tri/oligope | N |
| 482 | Egg | 407.2699 | 1 | 35.73519 | 165218.9 | 406.2626 | tri/oligope | N |
| 483 | Egg | 409.1728 | 1 | 12.38957 | 317617.3 | 408.1655 | tri/oligope | N |
| 484 | Egg | 413.267 | 1 | 37.84227 | 898769.7 | 412.2597 | tri/oligope | N |
| 485 | Egg | 413.2676 | 1 | 38.24858 | 513518.7 | 412.2603 | tri/oligope | N |
| 486 | Egg | 417.2356 | 1 | 14.87955 | 112847 | 416.2283 | tri/oligope | N |
| 487 | Egg | 419.2149 | 1 | 12.58989 | 125277.2 | 418.2076 | tri/oligope | N |
| 488 | Egg | 421.213 | 1 | 16.38858 | 115701.4 | 420.2057 | tri/oligope | N |
| 489 | Egg | 423.236 | 1 | 36.00488 | 112415.3 | 422.2288 | tri/oligope | N |
| 490 | Egg | 428.2516 | 1 | 17.0312 | 351348.2 | 427.2443 | tri/oligope | N |
| 491 | Egg | 428.3378 | 1 | 37.0692 | 175869.9 | 427.3305 | tri/oligope | N |
| 492 | Egg | 429.2416 | 1 | 38.30464 | 123396.8 | 428.2343 | tri/oligope | N |
| 493 | Egg | 430.2673 | 1 | 20.64273 | 197912.9 | 429.26 | tri/oligope | N |
| 494 | Egg | 432.2573 | 1 | 2.141692 | 281067.7 | 431.25 | tri/oligope | N |
| 495 | Egg | 435.1888 | 1 | 19.8693 | 125170.1 | 434.1815 | tri/oligope | N |
| 496 | Egg | 440.4109 | 1 | 39.08271 | 156417.8 | 439.4036 | tri/oligope | N |
| 497 | Egg | 441.2981 | 1 | 37.15278 | 153152 | 440.2909 | tri/oligope | N |
| 498 | Egg | 446.2257 | 1 | 15.75597 | 454547.7 | 445.2184 | tri/oligope | N |
| 499 | Egg | 446.2614 | 1 | 11.90143 | 217496.7 | 445.2541 | tri/oligope | N |
| 500 | Egg | 446.262 | 1 | 15.35666 | 629321.4 | 445.2547 | tri/oligope | N |
| 501 | Egg | 446.2626 | 1 | 9.92875 | 131778.6 | 445.2553 | tri/oligope | N |
| 502 | Egg | 447.3467 | 1 | 36.38005 | 209128.6 | 446.3394 | tri/oligope | N |
| 503 | Egg | 449.288 | 1 | 37.0323 | 266715.4 | 448.2808 | tri/oligope | N |
| 504 | Egg | 451.2662 | 1 | 37.15278 | 264204.5 | 450.2589 | tri/oligope | N |
| 505 | Egg | 456.2827 | 1 | 21.93832 | 558774.5 | 455.2755 | tri/oligope | N |
| 506 | Egg | 459.2645 | 1 | 20.85927 | 1013701 | 458.2573 | tri/oligope | N |
| 507 | Egg | 461.2618 | 1 | 20.1633 | 103922.2 | 460.2545 | tri/oligope | N |
| 508 | Egg | 465.2182 | 1 | 24.48658 | 230504.1 | 464.2109 | tri/oligope | N |
| 509 | Egg | 465.2359 | 1 | 21.67026 | 119933.6 | 464.2286 | tri/oligope | N |
| 510 | Egg | 466.3167 | 1 | 33.26533 | 2330749 | 465.3095 | tri/oligope | N |
| 511 | Egg | 467.3486 | 1 | 34.0478 | 106236.9 | 466.3413 | tri/oligope | N |
| 512 | Egg | 468.4418 | 1 | 37.48591 | 262781 | 467.4346 | tri/oligope | N |
| 513 | Egg | 235.5955 | 2 | 15.75597 | 131240.6 | 469.1765 | tri/oligope | N |
| 514 | Egg | 471.2829 | 1 | 24.6175 | 472895.6 | 470.2756 | tri/oligope | N |
| 515 | Egg | 472.364 | 1 | 37.0323 | 115984.4 | 471.3567 | tri/oligope | N |
| 516 | Egg | 473.2253 | 1 | 14.1332 | 333433 | 472.218 | tri/oligope | N |
| 517 | Egg | 476.2518 | 1 | 19.70328 | 304537.2 | 475.2445 | tri/oligope | N |
| 518 | Egg | 477.3666 | 1 | 34.98545 | 124092.4 | 476.3594 | tri/oligope | N |
| 519 | Egg | 480.2464 | 1 | 17.74115 | 583411.4 | 479.2391 | tri/oligope | N |
| 520 | Egg | 242.5901 | 2 | 15.75597 | 115155.2 | 483.1656 | tri/oligope | N |
| 521 | Egg | 484.3142 | 1 | 23.72195 | 489893 | 483.3069 | tri/oligope | N |
| 522 | Egg | 484.3144 | 1 | 20.22308 | 119378.4 | 483.3072 | tri/oligope | N |
| 523 | Egg | 491.2944 | 1 | 37.0602 | 113982.6 | 490.2871 | tri/oligope | N |
| 524 | Egg | 493.3142 | 1 | 36.99063 | 100240.1 | 492.3069 | tri/oligope | N |
| 525 | Egg | 498.2893 | 1 | 33.01051 | 192722.1 | 497.2821 | tri/oligope | N |
| 526 | Egg | 250.1884 | 2 | 38.98545 | 110931.3 | 498.3622 | tri/oligope | N |

| | | | | | | | | |
|---|---|---|---|---|---|---|---|---|
| 527 | Egg | 500.305 | 1 | 32.42 | 715467.5 | 499.2978 | tri/oligope | N |
| 528 | Egg | 251.6661 | 2 | 13.40645 | 220406.2 | 501.3177 | tri/oligope | N |
| 529 | Egg | 503.2836 | 1 | 15.5226 | 121719 | 502.2764 | tri/oligope | N |
| 530 | Egg | 504.2673 | 1 | 17.87733 | 992139.1 | 503.26 | tri/oligope | N |
| 531 | Egg | 252.6427 | 2 | 2.404792 | 121225.8 | 503.2709 | tri/oligope | N |
| 532 | Egg | 504.2781 | 1 | 2.422696 | 128437 | 503.2709 | tri/oligope | N |
| 533 | Egg | 505.2262 | 1 | 2.351929 | 130222.3 | 504.219 | tri/oligope | N |
| 534 | Egg | 505.2674 | 1 | 27.08945 | 100582.4 | 504.2602 | tri/oligope | N |
| 535 | Egg | 513.3043 | 1 | 17.53155 | 117628.3 | 512.297 | tri/oligope | N |
| 536 | Egg | 515.3198 | 1 | 19.39674 | 365826.5 | 514.3125 | tri/oligope | N |
| 537 | Egg | 258.1691 | 2 | 7.649746 | 157780.5 | 514.3237 | tri/oligope | N |
| 538 | Egg | 516.2676 | 1 | 13.98917 | 566958.9 | 515.2603 | tri/oligope | N |
| 539 | Egg | 516.3003 | 1 | 31.51498 | 239272.6 | 515.293 | tri/oligope | N |
| 540 | Egg | 258.6614 | 2 | 17.74902 | 225449.7 | 515.3083 | tri/oligope | N |
| 541 | Egg | 516.3245 | 1 | 36.18276 | 100996.6 | 515.3172 | tri/oligope | N |
| 542 | Egg | 517.2629 | 1 | 16.59175 | 108324.6 | 516.2557 | tri/oligope | N |
| 543 | Egg | 517.2992 | 1 | 19.21868 | 210229.2 | 516.2919 | tri/oligope | N |
| 544 | Egg | 519.4391 | 1 | 37.16177 | 118129 | 518.4318 | tri/oligope | N |
| 545 | Egg | 520.2416 | 1 | 16.9723 | 266532.3 | 519.2343 | tri/oligope | N |
| 546 | Egg | 520.3342 | 1 | 21.53453 | 101054.1 | 519.3269 | tri/oligope | N |
| 547 | Egg | 521.2572 | 1 | 10.66618 | 128232.9 | 520.2499 | tri/oligope | N |
| 548 | Egg | 521.2579 | 1 | 3.08725 | 128464.8 | 520.2506 | tri/oligope | N |
| 549 | Egg | 522.2412 | 1 | 7.701329 | 154630.2 | 521.2339 | tri/oligope | N |
| 550 | Egg | 522.2415 | 1 | 2.516542 | 118105.2 | 521.2342 | tri/oligope | N |
| 551 | Egg | 261.6504 | 2 | 17.51057 | 268021.7 | 521.2862 | tri/oligope | N |
| 552 | Egg | 265.1532 | 2 | 11.84195 | 357926.2 | 528.2919 | tri/oligope | N |
| 553 | Egg | 265.1535 | 2 | 3.059763 | 348810.4 | 528.2925 | tri/oligope | N |
| 554 | Egg | 265.6565 | 2 | 4.933442 | 102853.5 | 529.2984 | tri/oligope | N |
| 555 | Egg | 533.4546 | 1 | 37.15278 | 262247.3 | 532.4473 | tri/oligope | N |
| 556 | Egg | 536.2397 | 1 | 16.91682 | 184967.9 | 535.2325 | tri/oligope | N |
| 557 | Egg | 536.2804 | 1 | 33.26533 | 162763.7 | 535.2731 | tri/oligope | N |
| 558 | Egg | 538.252 | 1 | 32.42 | 142233.1 | 537.2447 | tri/oligope | N |
| 559 | Egg | 542.302 | 1 | 22.05201 | 133293.9 | 541.2948 | tri/oligope | N |
| 560 | Egg | 543.3148 | 1 | 13.67203 | 175121.9 | 542.3076 | tri/oligope | N |
| 561 | Egg | 543.3154 | 1 | 13.07962 | 161642.2 | 542.3082 | tri/oligope | N |
| 562 | Egg | 544.2992 | 1 | 18.76391 | 127291.6 | 543.292 | tri/oligope | N |
| 563 | Egg | 273.1488 | 2 | 33.26533 | 139690.4 | 544.283 | tri/oligope | N |
| 564 | Egg | 547.2016 | 1 | 15.37784 | 326834.9 | 546.1943 | tri/oligope | N |
| 565 | Egg | 557.2949 | 1 | 17.35413 | 242803.1 | 556.2877 | tri/oligope | N |
| 566 | Egg | 559.3097 | 1 | 2.441813 | 198202.3 | 558.3024 | tri/oligope | N |
| 567 | Egg | 280.1586 | 2 | 2.459775 | 656934.5 | 558.3026 | tri/oligope | N |
| 568 | Egg | 280.1588 | 2 | 7.3065 | 503211.3 | 558.3031 | tri/oligope | N |
| 569 | Egg | 564.3609 | 1 | 22.45108 | 106341 | 563.3536 | tri/oligope | N |
| 570 | Egg | 284.1427 | 2 | 10.15414 | 172469.2 | 566.2709 | tri/oligope | N |
| 571 | Egg | 568.3353 | 1 | 22.08695 | 1481416 | 567.328 | tri/oligope | N |
| 572 | Egg | 284.6714 | 2 | 22.08695 | 123487.1 | 567.3283 | tri/oligope | N |
| 573 | Egg | 285.1931 | 2 | 18.83526 | 152499.4 | 568.3716 | tri/oligope | N |
| 574 | Egg | 285.6733 | 2 | 33.26533 | 125595.3 | 569.3321 | tri/oligope | N |
| 575 | Egg | 572.2944 | 1 | 19.75581 | 931701.7 | 571.2871 | tri/oligope | N |
| 576 | Egg | 286.6742 | 2 | 7.716142 | 273353.9 | 571.3338 | tri/oligope | N |

| | | | | | | | | |
|---|---|---|---|---|---|---|---|---|
| 577 | Egg | 286.6743 | 2 | 2.441813 | 335114.1 | 571.334 | tri/oligope | N |
| 578 | Egg | 575.2683 | 1 | 11.82405 | 127492 | 574.2611 | tri/oligope | N |
| 579 | Egg | 575.305 | 1 | 16.74948 | 868334.1 | 574.2977 | tri/oligope | N |
| 580 | Egg | 579.2788 | 1 | 15.42431 | 733518 | 578.2715 | tri/oligope | N |
| 581 | Egg | 293.6622 | 2 | 33.26533 | 228263 | 585.3098 | tri/oligope | N |
| 582 | Egg | 293.6877 | 2 | 2.992263 | 119026.3 | 585.3609 | tri/oligope | N |
| 583 | Egg | 293.6878 | 2 | 2.156538 | 156086.3 | 585.3611 | tri/oligope | N |
| 584 | Egg | 298.1515 | 2 | 2.351929 | 202505.4 | 594.2885 | tri/oligope | N |
| 585 | Egg | 298.1516 | 2 | 2.071563 | 364081 | 594.2886 | tri/oligope | N |
| 586 | Egg | 298.1517 | 2 | 3.021488 | 531673.6 | 594.2888 | tri/oligope | N |
| 587 | Egg | 300.1352 | 2 | 16.70318 | 143783.7 | 598.2558 | tri/oligope | N |
| 588 | Egg | 300.1796 | 2 | 10.59145 | 164199.7 | 598.3447 | tri/oligope | N |
| 589 | Egg | 304.1329 | 2 | 2.025771 | 165420.6 | 606.2512 | tri/oligope | N |
| 590 | Egg | 307.1878 | 2 | 19.32281 | 910238.4 | 612.3611 | tri/oligope | N |
| 591 | Egg | 307.6203 | 2 | 18.06362 | 115795 | 613.2261 | oligopepti | N |
| 592 | Egg | 312.1745 | 2 | 16.87238 | 215749.2 | 622.3344 | oligopepti | N |
| 593 | Egg | 315.1672 | 2 | 13.22776 | 124543 | 628.3198 | oligopepti | N |
| 594 | Egg | 632.3371 | 1 | 2.369725 | 140628.1 | 631.3299 | oligopepti | N |
| 595 | Egg | 316.6724 | 2 | 2.369725 | 426256.4 | 631.3302 | oligopepti | N |
| 596 | Egg | 316.6727 | 2 | 3.905454 | 293071.8 | 631.3309 | oligopepti | N |
| 597 | Egg | 316.6749 | 2 | 21.7642 | 281843.5 | 631.3353 | oligopepti | N |
| 598 | Egg | 635.309 | 1 | 22.49933 | 367774.7 | 634.3017 | oligopepti | N |
| 599 | Egg | 637.3072 | 1 | 37.69898 | 135266.9 | 636.2999 | oligopepti | N |
| 600 | Egg | 319.1882 | 2 | 17.2146 | 106366.5 | 636.3619 | oligopepti | N |
| 601 | Egg | 645.3447 | 1 | 31.4738 | 705616.3 | 644.3374 | oligopepti | N |
| 602 | Egg | 323.1882 | 2 | 2.992263 | 107681.7 | 644.3618 | oligopepti | N |
| 603 | Egg | 335.2371 | 2 | 13.04052 | 119501.1 | 668.4596 | oligopepti | N |
| 604 | Egg | 675.151 | 1 | 36.86903 | 100302.5 | 674.1437 | oligopepti | N |
| 605 | Egg | 338.688 | 2 | 22.05201 | 128615.6 | 675.3615 | oligopepti | N |
| 606 | Egg | 339.1855 | 2 | 19.3695 | 1575424 | 676.3564 | oligopepti | N |
| 607 | Egg | 343.139 | 2 | 17.50146 | 110792.6 | 684.2635 | oligopepti | N |
| 608 | Egg | 685.3791 | 1 | 22.69245 | 228722.3 | 684.3718 | oligopepti | N |
| 609 | Egg | 344.1879 | 2 | 7.898871 | 151151.8 | 686.3612 | oligopepti | N |
| 610 | Egg | 344.1882 | 2 | 2.921871 | 130016.1 | 686.3618 | oligopepti | N |
| 611 | Egg | 344.1882 | 2 | 2.459775 | 113770.7 | 686.3619 | oligopepti | N |
| 612 | Egg | 692.2771 | 1 | 26.96676 | 132420.2 | 691.2698 | oligopepti | N |
| 613 | Egg | 347.6807 | 2 | 14.78566 | 140540.5 | 693.3469 | oligopepti | N |
| 614 | Egg | 697.6587 | 1 | 38.9788 | 154584 | 696.6514 | oligopepti | N |
| 615 | Egg | 349.7146 | 2 | 14.49884 | 580888.2 | 697.4146 | oligopepti | N |
| 616 | Egg | 350.686 | 2 | 14.1332 | 341851 | 699.3574 | oligopepti | N |
| 617 | Egg | 359.6914 | 2 | 2.269517 | 150158.4 | 717.3683 | oligopepti | N |
| 618 | Egg | 741.4268 | 1 | 20.06393 | 124249.8 | 740.4195 | oligopepti | N |
| 619 | Egg | 371.2173 | 2 | 20.06393 | 4673409 | 740.4201 | oligopepti | N |
| 620 | Egg | 377.6669 | 2 | 10.68464 | 255160.9 | 753.3193 | oligopepti | N |
| 621 | Egg | 383.1961 | 2 | 20.06393 | 175847.4 | 764.3777 | oligopepti | N |
| 622 | Egg | 384.65 | 2 | 14.25139 | 108431.6 | 767.2854 | oligopepti | N |
| 623 | Egg | 392.7273 | 2 | 26.6692 | 396587.1 | 783.44 | oligopepti | N |
| 624 | Egg | 784.4474 | 1 | 26.6692 | 1044673 | 783.4401 | oligopepti | N |
| 625 | Egg | 394.2147 | 2 | 14.47958 | 123025.6 | 786.4149 | oligopepti | N |
| 626 | Egg | 797.352 | 1 | 19.23475 | 199749.3 | 796.3448 | oligopepti | N |

| | | | | | | | | |
|---|---|---|---|---|---|---|---|---|
| 627 | Egg | 404.7063 | 2 | 26.6692 | 131692.2 | 807.3981 | oligopepti( | N |
| 628 | Egg | 407.7249 | 2 | 10.38225 | 775374.5 | 813.4353 | oligopepti( | N |
| 629 | Egg | 407.7255 | 2 | 2.459775 | 264847 | 813.4364 | oligopepti( | N |
| 630 | Egg | 424.2203 | 2 | 21.19993 | 354728.3 | 846.4261 | oligopepti( | N |
| 631 | Egg | 428.1659 | 2 | 13.76779 | 233381 | 854.3173 | oligopepti( | N |
| 632 | Egg | 298.8403 | 3 | 10.83385 | 180157.5 | 893.4991 | oligopepti( | N |
| 633 | Egg | 452.749 | 2 | 22.37599 | 978595.2 | 903.4835 | oligopepti( | N |
| 634 | Egg | 303.8065 | 3 | 2.318379 | 126301.4 | 908.3977 | oligopepti( | N |
| 635 | Egg | 460.6668 | 2 | 14.75074 | 178302.9 | 919.319 | oligopepti( | N |
| 636 | Egg | 466.2413 | 2 | 17.52313 | 798509.8 | 930.468 | oligopepti( | N |
| 637 | Egg | 931.4755 | 1 | 17.52313 | 267121.1 | 930.4682 | oligopepti( | N |
| 638 | Egg | 468.2286 | 2 | 21.73675 | 176876.5 | 934.4427 | oligopepti( | N |
| 639 | Egg | 471.1939 | 2 | 14.26156 | 110055 | 940.3733 | oligopepti( | N |
| 640 | Egg | 478.2202 | 2 | 17.53155 | 130352.6 | 954.4259 | oligopepti( | N |
| 641 | Egg | 498.902 | 2 | 1.582842 | 248956.6 | 995.7894 | oligopepti( | N |
| 642 | Egg | 515.2654 | 2 | 15.98212 | 113812.4 | 1028.516 | oligopepti( | N |
| 643 | Egg | 521.2914 | 2 | 17.62739 | 115067.1 | 1040.568 | oligopepti( | N |
| 644 | Egg | 347.8636 | 3 | 17.64046 | 543905 | 1040.569 | oligopepti( | N |
| 645 | Egg | 524.771 | 2 | 26.35684 | 156793.7 | 1047.527 | oligopepti( | N |
| 646 | Egg | 548.8698 | 2 | 1.630292 | 124112.6 | 1095.725 | oligopepti( | N |
| 647 | Egg | 556.7309 | 2 | 17.72337 | 109053.3 | 1111.447 | oligopepti( | N |
| 648 | Egg | 1126.764 | 1 | 1.619417 | 127620.2 | 1125.756 | oligopepti( | N |
| 649 | Egg | 587.2569 | 2 | 16.61947 | 129361.5 | 1172.499 | oligopepti( | N |
| 650 | Egg | 587.2571 | 2 | 18.31643 | 135368.1 | 1172.5 | oligopepti( | N |
| 651 | Egg | 589.8189 | 2 | 21.18589 | 136695.8 | 1177.623 | oligopepti( | N |
| 652 | Egg | 608.8164 | 2 | 24.66908 | 378238.8 | 1215.618 | oligopepti( | N |
| 653 | Egg | 615.8062 | 2 | 27.45297 | 121203.7 | 1229.598 | oligopepti( | N |
| 654 | Egg | 418.988 | 3 | 36.76753 | 107804.4 | 1253.942 | oligopepti( | N |
| 655 | Egg | 439.5545 | 3 | 21.45928 | 353139.6 | 1315.642 | oligopepti( | N |
| 656 | Egg | 663.2923 | 2 | 17.4836 | 115361.2 | 1324.57 | oligopepti( | N |
| 657 | Egg | 664.7945 | 2 | 26.2486 | 1064544 | 1327.574 | oligopepti( | N |
| 658 | Egg | 451.5182 | 3 | 26.2486 | 166497.5 | 1351.533 | oligopepti( | N |
| 659 | Egg | 456.1811 | 3 | 26.2486 | 108505.9 | 1365.521 | oligopepti( | N |
| 660 | Egg | 686.4669 | 2 | 34.08 | 212931.8 | 1370.919 | oligopepti( | N |
| 661 | Egg | 502.9077 | 3 | 17.1089 | 141705 | 1505.701 | oligopepti( | N |
| 662 | Egg | 377.4328 | 4 | 17.1089 | 163403.9 | 1505.702 | oligopepti( | N |
| 663 | Egg | 552.8916 | 3 | 20.29598 | 127708.9 | 1655.653 | oligopepti( | N |
| 664 | Egg | 586.5745 | 3 | 19.79107 | 245005 | 1756.702 | oligopepti( | N |
| 665 | Egg | 1024.782 | 2 | 1.607821 | 130112.8 | 2047.55 | oligopepti( | N |
| 666 | Egg | 1058.777 | 2 | 1.607821 | 197013.4 | 2115.54 | oligopepti( | N |
| 667 | Egg | 1092.77 | 2 | 1.607821 | 116870.8 | 2183.525 | oligopepti( | N |
| 668 | Egg | 1031.87 | 3 | 33.14594 | 1117333 | 3092.589 | oligopepti( | N |
| 669 | Egg | 779.8988 | 4 | 33.13924 | 132849.4 | 3115.566 | oligopepti( | N |
| 670 | Egg | 636.5268 | 5 | 31.70659 | 195539.7 | 3177.598 | oligopepti( | N |
| 671 | Egg | 1069.564 | 3 | 33.711 | 179431.8 | 3205.67 | oligopepti( | N |
| 672 | Egg | 1069.568 | 3 | 34.17494 | 126781.8 | 3205.684 | oligopepti( | N |
| 673 | Egg | 872.8746 | 4 | 12.65745 | 125238.4 | 3487.469 | oligopepti( | N |
| 674 | Egg | 935.2255 | 4 | 33.0191 | 2579990 | 3736.873 | oligopepti( | N |
| 675 | Egg | 748.3819 | 5 | 33.01051 | 7303922 | 3736.873 | oligopepti( | N |
| 676 | Egg | 623.8197 | 6 | 33.01051 | 920190.7 | 3736.874 | oligopepti( | N |

| | | | | | | | | |
|---|---|---|---|---|---|---|---|---|
| 677 | Egg | 748.3827 | 5 | 40.04883 | 128365.7 | 3736.877 | oligopepti | N |
| 678 | Egg | 752.7764 | 5 | 33.0191 | 312760.2 | 3758.846 | oligopepti | N |
| 679 | Egg | 627.4816 | 6 | 33.05928 | 145900.4 | 3758.846 | oligopepti | N |
| 680 | Egg | 630.1441 | 6 | 33.00094 | 217420 | 3774.821 | oligopepti | N |
| 681 | Egg | 755.9715 | 5 | 33.01051 | 339460.3 | 3774.821 | oligopepti | N |
| 682 | Egg | 952.9845 | 4 | 33.01051 | 115642.4 | 3807.909 | oligopepti | N |
| 683 | Egg | 762.5901 | 5 | 33.04512 | 570495.8 | 3807.914 | oligopepti | N |
| 684 | Egg | 970.7438 | 4 | 33.0191 | 2757822 | 3878.946 | oligopepti | N |
| 685 | Egg | 776.7965 | 5 | 33.0191 | 10391855 | 3878.946 | oligopepti | N |
| 686 | Egg | 647.4985 | 6 | 33.01051 | 1932310 | 3878.947 | oligopepti | N |
| 687 | Egg | 776.7979 | 5 | 40.07121 | 193215.7 | 3878.953 | oligopepti | N |
| 688 | Egg | 651.1607 | 6 | 33.04512 | 277217.9 | 3900.921 | oligopepti | N |
| 689 | Egg | 781.3915 | 5 | 33.03768 | 415596.7 | 3901.921 | oligopepti | N |
| 690 | Egg | 653.823 | 6 | 33.01051 | 379093.2 | 3916.894 | oligopepti | N |
| 691 | Egg | 784.388 | 5 | 33.16486 | 485295 | 3916.904 | oligopepti | N |
| 692 | Egg | 839.4286 | 5 | 34.34394 | 225926.9 | 4192.107 | oligopepti | N |
| 693 | Egg | 1142.905 | 5 | 23.49596 | 575832.8 | 5709.487 | oligopepti | N |
| 694 | Egg | 952.5892 | 6 | 23.47803 | 953732.3 | 5709.491 | oligopepti | N |
| 695 | Egg | 955.5912 | 6 | 24.45528 | 433373.7 | 5727.503 | oligopepti | N |
| 696 | Egg | 1146.509 | 5 | 24.53197 | 132042.5 | 5727.511 | oligopepti | N |
| 697 | Egg | 819.3654 | 7 | 24.41442 | 138041.7 | 5728.507 | oligopepti | N |

| ID | Protein | mz | charge | retentiont | raw.abund | mass | type | Match |
|---|---|---|---|---|---|---|---|---|
| 2 | Soy | 231.1707 | 1 | 5.410937 | 1300555 | 230.1635 | di/tripepti | Y |
| 3 | Soy | 231.1708 | 1 | 9.712147 | 732405.9 | 230.1635 | di/tripepti | Y |
| 4 | Soy | 231.1708 | 1 | 12.0754 | 841262.9 | 230.1635 | di/tripepti | Y |
| 5 | Soy | 233.1497 | 1 | 2.19034 | 362064.6 | 232.1424 | di/tripepti | Y |
| 6 | Soy | 233.1498 | 1 | 2.804483 | 211873.9 | 232.1425 | di/tripepti | Y |
| 8 | Soy | 239.1028 | 1 | 2.453827 | 112055.1 | 238.0956 | di/tripepti | Y |
| 11 | Soy | 245.1862 | 1 | 16.43531 | 1401550 | 244.1789 | di/tripepti | Y |
| 12 | Soy | 245.1864 | 1 | 15.90536 | 286577.3 | 244.1791 | di/tripepti | Y |
| 13 | Soy | 245.1862 | 1 | 14.55055 | 755014.6 | 244.1789 | di/tripepti | Y |
| 14 | Soy | 245.1865 | 1 | 17.69164 | 359748.6 | 244.1792 | di/tripepti | Y |
| 15 | Soy | 246.145 | 1 | 2.19034 | 957325.6 | 245.1377 | di/tripepti | Y |
| 15 | Soy | 246.145 | 1 | 1.96049 | 122597.2 | 245.1377 | di/tripepti | Y |
| 16 | Soy | 246.145 | 1 | 2.67671 | 525933.2 | 245.1378 | di/tripepti | Y |
| 17 | Soy | 246.1452 | 1 | 3.407643 | 1168265 | 245.1379 | di/tripepti | Y |
| 20 | Soy | 247.1289 | 1 | 2.134193 | 557202.4 | 246.1217 | di/tripepti | Y |
| 21 | Soy | 247.1293 | 1 | 3.186757 | 292606.6 | 246.122 | di/tripepti | Y |
| 22 | Soy | 247.1293 | 1 | 4.693403 | 376982.4 | 246.122 | di/tripepti | Y |
| 27 | Soy | 253.1185 | 1 | 2.299983 | 360570 | 252.1112 | di/tripepti | Y |
| 27 | Soy | 253.1186 | 1 | 2.510303 | 109239.3 | 252.1113 | di/tripepti | Y |
| 28 | Soy | 253.1188 | 1 | 9.330253 | 189787.8 | 252.1115 | di/tripepti | Y |
| 30 | Soy | 260.1607 | 1 | 2.19034 | 853381.7 | 259.1534 | di/tripepti | Y |
| 31 | Soy | 260.1609 | 1 | 2.68468 | 336825.5 | 259.1536 | di/tripepti | Y |
| 33 | Soy | 260.197 | 1 | 1.857653 | 144687.2 | 259.1897 | di/tripepti | Y |
| 34 | Soy | 261.1449 | 1 | 2.201593 | 532283.9 | 260.1376 | di/tripepti | Y |
| 35 | Soy | 261.145 | 1 | 3.31379 | 201665.5 | 260.1377 | di/tripepti | Y |
| 36 | Soy | 261.145 | 1 | 5.432833 | 513078.5 | 260.1377 | di/tripepti | Y |
| 38 | Soy | 263.1429 | 1 | 11.53434 | 346796.2 | 262.1357 | di/tripepti | Y |
| 40 | Soy | 265.1552 | 1 | 16.28741 | 234461.4 | 264.1479 | di/tripepti | Y |
| 41 | Soy | 267.1343 | 1 | 2.41733 | 161881.6 | 266.127 | di/tripepti | Y |
| 42 | Soy | 269.1609 | 1 | 2.179287 | 242856.1 | 268.1536 | di/tripepti | Y |
| 44 | Soy | 277.1032 | 1 | 1.86743 | 113426.3 | 276.0959 | di/tripepti | Y |
| 47 | Soy | 288.1924 | 1 | 16.08951 | 297451.1 | 287.1851 | di/tripepti | Y |
| 50 | Soy | 294.1451 | 1 | 2.34546 | 242460.4 | 293.1378 | di/tripepti | Y |
| 51 | Soy | 295.1295 | 1 | 3.292053 | 391894.4 | 294.1222 | di/tripepti | Y |
| 52 | Soy | 295.1657 | 1 | 16.43531 | 675524.7 | 294.1584 | di/tripepti | Y |
| 55 | Soy | 302.208 | 1 | 16.4289 | 131435.8 | 301.2007 | tri/oligope | Y |
| 56 | Soy | 302.2081 | 1 | 15.16414 | 549487.3 | 301.2008 | tri/oligope | Y |
| 57 | Soy | 302.2081 | 1 | 19.87241 | 233654.2 | 301.2008 | tri/oligope | Y |
| 58 | Soy | 303.1665 | 1 | 2.19034 | 285780.6 | 302.1593 | tri/oligope | Y |
| 59 | Soy | 304.1507 | 1 | 2.22438 | 358797.1 | 303.1434 | tri/oligope | Y |
| 60 | Soy | 304.151 | 1 | 4.902313 | 205994 | 303.1437 | tri/oligope | Y |
| 61 | Soy | 304.1663 | 1 | 19.00177 | 137380 | 303.159 | tri/oligope | Y |
| 63 | Soy | 318.1665 | 1 | 2.769463 | 204777.1 | 317.1593 | tri/oligope | Y |
| 64 | Soy | 318.1663 | 1 | 2.201593 | 385805.6 | 317.1591 | tri/oligope | Y |
| 65 | Soy | 318.1668 | 1 | 9.139763 | 134922.6 | 317.1596 | tri/oligope | Y |
| 66 | Soy | 318.1821 | 1 | 22.57565 | 271068.8 | 317.1748 | tri/oligope | Y |
| 71 | Soy | 330.2395 | 1 | 18.13061 | 256233.3 | 329.2323 | tri/oligope | Y |
| 72 | Soy | 331.1615 | 1 | 1.902557 | 173004.9 | 330.1542 | tri/oligope | Y |
| 73 | Soy | 331.1657 | 1 | 2.86022 | 299527.6 | 330.1584 | tri/oligope | Y |

| | | | | | | | | |
|---|---|---|---|---|---|---|---|---|
| 74 | Soy | 331.2851 | 1 | 38.46329 | 122344.4 | 330.2778 | tri/oligope | Y |
| 75 | Soy | 332.1822 | 1 | 2.58934 | 213983.8 | 331.1749 | tri/oligope | Y |
| 80 | Soy | 334.1613 | 1 | 2.334193 | 326816.8 | 333.154 | tri/oligope | Y |
| 85 | Soy | 344.2554 | 1 | 21.64955 | 298270.3 | 343.2481 | tri/oligope | Y |
| 85 | Soy | 344.2554 | 1 | 21.3753 | 293471.4 | 343.2482 | tri/oligope | Y |
| 86 | Soy | 344.2555 | 1 | 20.19351 | 178863.5 | 343.2482 | tri/oligope | Y |
| 89 | Soy | 358.2712 | 1 | 23.5953 | 106994.3 | 357.2639 | tri/oligope | Y |
| 90 | Soy | 359.2293 | 1 | 2.22438 | 520378.8 | 358.222 | tri/oligope | Y |
| 92 | Soy | 360.2136 | 1 | 2.769463 | 380848.6 | 359.2063 | tri/oligope | Y |
| 94 | Soy | 363.1555 | 1 | 2.19034 | 131465.9 | 362.1482 | tri/oligope | Y |
| 95 | Soy | 365.1366 | 1 | 37.81734 | 108596.2 | 364.1293 | tri/oligope | Y |
| 96 | Soy | 366.2139 | 1 | 2.201593 | 189125.8 | 365.2066 | tri/oligope | Y |
| 105 | Soy | 390.1876 | 1 | 2.356947 | 144443.4 | 389.1804 | tri/oligope | Y |
| 106 | Soy | 395.0393 | 1 | 1.573627 | 133555.9 | 394.032 | tri/oligope | Y |
| 107 | Soy | 399.2608 | 1 | 18.79712 | 937735.7 | 398.2535 | tri/oligope | Y |
| 112 | Soy | 415.2118 | 1 | 36.47641 | 292091.4 | 414.2045 | tri/oligope | Y |
| 121 | Soy | 433.1935 | 1 | 2.49893 | 284382.9 | 432.1862 | tri/oligope | Y |
| 133 | Soy | 461.3005 | 1 | 37.39671 | 107878.8 | 460.2932 | tri/oligope | Y |
| 134 | Soy | 463.0267 | 1 | 1.58502 | 172385.9 | 462.0194 | tri/oligope | Y |
| 141 | Soy | 479.3102 | 1 | 37.13019 | 150483.5 | 478.303 | tri/oligope | Y |
| 144 | Soy | 488.2516 | 1 | 19.13773 | 247176.3 | 487.2443 | tri/oligope | Y |
| 151 | Soy | 500.2882 | 1 | 25.20616 | 170945.1 | 499.2809 | tri/oligope | Y |
| 156 | Soy | 515.2914 | 1 | 37.09338 | 102154.3 | 514.2842 | tri/oligope | Y |
| 159 | Soy | 520.278 | 1 | 21.18518 | 186427.3 | 519.2707 | tri/oligope | Y |
| 161 | Soy | 264.1747 | 2 | 35.30401 | 105257.5 | 526.3349 | tri/oligope | Y |
| 165 | Soy | 531.014 | 1 | 1.596133 | 219946 | 530.0068 | tri/oligope | Y |
| 166 | Soy | 531.3871 | 1 | 37.83629 | 113701.2 | 530.3798 | tri/oligope | Y |
| 174 | Soy | 274.1676 | 2 | 35.93114 | 112584.8 | 546.3207 | tri/oligope | Y |
| 194 | Soy | 599.0019 | 1 | 1.58502 | 201379.3 | 597.9946 | tri/oligope | Y |
| 208 | Soy | 643.3318 | 1 | 20.74867 | 129363.5 | 642.3245 | oligopepti | Y |
| 218 | Soy | 666.9893 | 1 | 1.596133 | 192468.4 | 665.982 | oligopepti | Y |
| 240 | Soy | 734.9768 | 1 | 1.596133 | 170379.3 | 733.9696 | oligopepti | Y |
| 258 | Soy | 802.9643 | 1 | 1.58502 | 145325.8 | 801.9571 | oligopepti | Y |
| 276 | Soy | 870.9516 | 1 | 1.596133 | 101006.3 | 869.9443 | oligopepti | Y |
| 296 | Soy | 974.814 | 1 | 1.60749 | 254589.3 | 973.8067 | oligopepti | Y |
| 297 | Soy | 990.7886 | 1 | 1.60749 | 184364 | 989.7813 | oligopepti | Y |
| 338 | Soy | 223.1081 | 1 | 5.138677 | 292885.1 | 222.1008 | dipeptide | N |
| 339 | Soy | 223.1081 | 1 | 10.45349 | 152375.2 | 222.1009 | dipeptide | N |
| 340 | Soy | 226.9517 | 1 | 1.54202 | 159697.9 | 225.9444 | di/tripepti | N |
| 341 | Soy | 226.9521 | 1 | 1.664157 | 160373.7 | 225.9448 | di/tripepti | N |
| 342 | Soy | 229.1552 | 1 | 9.020453 | 201268.8 | 228.1479 | di/tripepti | N |
| 343 | Soy | 231.1708 | 1 | 8.981633 | 549444.1 | 230.1635 | di/tripepti | N |
| 344 | Soy | 232.1292 | 1 | 2.19034 | 145900 | 231.122 | di/tripepti | N |
| 345 | Soy | 233.1133 | 1 | 2.19034 | 146559.5 | 232.106 | di/tripepti | N |
| 346 | Soy | 233.15 | 1 | 4.093763 | 381285.4 | 232.1427 | di/tripepti | N |
| 347 | Soy | 237.1238 | 1 | 4.64821 | 230371.4 | 236.1165 | di/tripepti | N |
| 348 | Soy | 237.1238 | 1 | 11.22803 | 287221.3 | 236.1165 | di/tripepti | N |
| 349 | Soy | 246.1453 | 1 | 5.35951 | 483600.5 | 245.138 | di/tripepti | N |
| 350 | Soy | 253.1295 | 1 | 2.19034 | 111593 | 252.1222 | di/tripepti | N |
| 351 | Soy | 255.0658 | 1 | 30.4133 | 244336.4 | 254.0586 | di/tripepti | N |

| | | | | | | | | |
|---|---|---|---|---|---|---|---|---|
| 352 | Soy | 258.1103 | 1 | 1.73406 | 146112.3 | 257.103 | di/tripepti | N |
| 353 | Soy | 263.1398 | 1 | 14.45582 | 265925 | 262.1325 | di/tripepti | N |
| 354 | Soy | 263.143 | 1 | 13.13653 | 136218.1 | 262.1357 | di/tripepti | N |
| 355 | Soy | 265.1551 | 1 | 14.76493 | 265680.2 | 264.1478 | di/tripepti | N |
| 356 | Soy | 269.1134 | 1 | 2.34546 | 135641.7 | 268.1062 | di/tripepti | N |
| 357 | Soy | 271.0602 | 1 | 33.8803 | 167278.8 | 270.053 | di/tripepti | N |
| 358 | Soy | 276.1192 | 1 | 1.803487 | 133444.1 | 275.1119 | di/tripepti | N |
| 359 | Soy | 276.1349 | 1 | 17.19951 | 217938.5 | 275.1276 | di/tripepti | N |
| 360 | Soy | 276.1558 | 1 | 2.201593 | 196380.4 | 275.1485 | di/tripepti | N |
| 361 | Soy | 276.156 | 1 | 3.383567 | 187150.3 | 275.1487 | di/tripepti | N |
| 362 | Soy | 278.1171 | 1 | 2.145577 | 112890.7 | 277.1098 | di/tripepti | N |
| 363 | Soy | 279.1709 | 1 | 19.91176 | 442501.4 | 278.1637 | di/tripepti | N |
| 364 | Soy | 279.171 | 1 | 18.99481 | 126435.1 | 278.1637 | di/tripepti | N |
| 365 | Soy | 279.1711 | 1 | 21.03812 | 154404.5 | 278.1638 | di/tripepti | N |
| 366 | Soy | 279.2321 | 1 | 37.01965 | 588390.2 | 278.2248 | di/tripepti | N |
| 367 | Soy | 280.1294 | 1 | 2.22438 | 120614.4 | 279.1221 | di/tripepti | N |
| 368 | Soy | 280.1299 | 1 | 10.05288 | 105144.5 | 279.1226 | di/tripepti | N |
| 369 | Soy | 281.1137 | 1 | 11.59336 | 397016.3 | 280.1065 | di/tripepti | N |
| 370 | Soy | 281.1502 | 1 | 6.15241 | 263167.2 | 280.143 | di/tripepti | N |
| 371 | Soy | 286.1768 | 1 | 13.45593 | 456719.6 | 285.1695 | di/tripepti | N |
| 372 | Soy | 288.192 | 1 | 2.247413 | 372573.7 | 287.1847 | di/tripepti | N |
| 373 | Soy | 288.2032 | 1 | 1.99523 | 129169.2 | 287.1959 | di/tripepti | N |
| 374 | Soy | 289.1509 | 1 | 2.02972 | 132674.3 | 288.1436 | di/tripepti | N |
| 375 | Soy | 290.1348 | 1 | 2.02972 | 113681.5 | 289.1275 | di/tripepti | N |
| 376 | Soy | 290.1716 | 1 | 2.201593 | 577191.9 | 289.1643 | di/tripepti | N |
| 377 | Soy | 292.1296 | 1 | 16.43531 | 222800.7 | 291.1223 | di/tripepti | N |
| 378 | Soy | 295.1297 | 1 | 12.65641 | 191504 | 294.1224 | di/tripepti | N |
| 379 | Soy | 295.1657 | 1 | 14.61802 | 289237.6 | 294.1585 | di/tripepti | N |
| 380 | Soy | 295.166 | 1 | 13.42547 | 759601.5 | 294.1587 | di/tripepti | N |
| 381 | Soy | 295.2271 | 1 | 36.28755 | 121969.6 | 294.2198 | di/tripepti | N |
| 382 | Soy | 295.2272 | 1 | 37.38631 | 317811.2 | 294.2199 | di/tripepti | N |
| 383 | Soy | 296.1243 | 1 | 2.19034 | 119462.3 | 295.1171 | di/tripepti | N |
| 384 | Soy | 297.1086 | 1 | 2.719777 | 109351.4 | 296.1013 | di/tripepti | N |
| 385 | Soy | 297.2427 | 1 | 36.15032 | 180915.9 | 296.2354 | di/tripepti | N |
| 386 | Soy | 303.1669 | 1 | 5.837807 | 130791 | 302.1597 | tri/oligope | N |
| 387 | Soy | 304.133 | 1 | 2.66673 | 102250.4 | 303.1257 | tri/oligope | N |
| 388 | Soy | 304.1869 | 1 | 2.201593 | 114677.2 | 303.1796 | tri/oligope | N |
| 389 | Soy | 304.1871 | 1 | 2.41733 | 132742.1 | 303.1799 | tri/oligope | N |
| 390 | Soy | 306.1457 | 1 | 10.7608 | 546120.8 | 305.1385 | tri/oligope | N |
| 391 | Soy | 306.1662 | 1 | 2.19034 | 192813.9 | 305.1589 | tri/oligope | N |
| 392 | Soy | 310.1402 | 1 | 2.58934 | 190464.3 | 309.1329 | tri/oligope | N |
| 393 | Soy | 311.1243 | 1 | 2.77728 | 121001 | 310.117 | tri/oligope | N |
| 394 | Soy | 313.2376 | 1 | 35.81307 | 184336.3 | 312.2303 | tri/oligope | N |
| 395 | Soy | 314.2081 | 1 | 13.67871 | 1045387 | 313.2008 | tri/oligope | N |
| 396 | Soy | 315.2533 | 1 | 36.15032 | 389994.9 | 314.246 | tri/oligope | N |
| 397 | Soy | 316.2239 | 1 | 14.83729 | 101454.9 | 315.2167 | tri/oligope | N |
| 398 | Soy | 316.224 | 1 | 19.60118 | 250725.8 | 315.2167 | tri/oligope | N |
| 399 | Soy | 317.1823 | 1 | 2.201593 | 299447.4 | 316.175 | tri/oligope | N |
| 400 | Soy | 317.1826 | 1 | 2.48644 | 723417.7 | 316.1753 | tri/oligope | N |
| 401 | Soy | 317.2091 | 1 | 37.38631 | 194403.6 | 316.2018 | tri/oligope | N |

| | | | | | | | | |
|---|---|---|---|---|---|---|---|---|
| 402 | Soy | 318.149 | 1 | 3.69555 | 144784.2 | 317.1417 | tri/oligope | N |
| 403 | Soy | 318.2027 | 1 | 2.356947 | 190941.7 | 317.1954 | tri/oligope | N |
| 404 | Soy | 318.203 | 1 | 14.61802 | 211426.2 | 317.1957 | tri/oligope | N |
| 405 | Soy | 318.2032 | 1 | 15.91184 | 133152.6 | 317.1959 | tri/oligope | N |
| 406 | Soy | 319.141 | 1 | 8.144863 | 194093 | 318.1337 | tri/oligope | N |
| 407 | Soy | 320.125 | 1 | 17.67373 | 145644.5 | 319.1177 | tri/oligope | N |
| 408 | Soy | 320.1456 | 1 | 2.19034 | 122527.5 | 319.1383 | tri/oligope | N |
| 409 | Soy | 327.0519 | 1 | 1.61864 | 168343.6 | 326.0446 | tri/oligope | N |
| 410 | Soy | 328.224 | 1 | 16.87933 | 146216.7 | 327.2168 | tri/oligope | N |
| 411 | Soy | 328.2339 | 1 | 15.50291 | 102345.1 | 327.2266 | tri/oligope | N |
| 412 | Soy | 330.2033 | 1 | 13.98454 | 106322.9 | 329.196 | tri/oligope | N |
| 413 | Soy | 331.1981 | 1 | 2.41733 | 225651.2 | 330.1908 | tri/oligope | N |
| 414 | Soy | 331.2484 | 1 | 35.42013 | 105464.4 | 330.2412 | tri/oligope | N |
| 415 | Soy | 332.2188 | 1 | 15.84154 | 152830.2 | 331.2115 | tri/oligope | N |
| 416 | Soy | 332.2188 | 1 | 18.60684 | 344494.4 | 331.2115 | tri/oligope | N |
| 417 | Soy | 333.1771 | 1 | 2.212967 | 262662 | 332.1699 | tri/oligope | N |
| 418 | Soy | 334.1248 | 1 | 1.93733 | 116368.2 | 333.1175 | tri/oligope | N |
| 419 | Soy | 334.1406 | 1 | 17.97144 | 313258.7 | 333.1333 | tri/oligope | N |
| 420 | Soy | 334.1616 | 1 | 5.14789 | 106534 | 333.1543 | tri/oligope | N |
| 421 | Soy | 334.1771 | 1 | 18.03891 | 107605.3 | 333.1698 | tri/oligope | N |
| 422 | Soy | 335.2196 | 1 | 36.73052 | 101886.2 | 334.2123 | tri/oligope | N |
| 423 | Soy | 335.2196 | 1 | 36.43542 | 154959.3 | 334.2123 | tri/oligope | N |
| 424 | Soy | 336.1386 | 1 | 21.04663 | 177225.4 | 335.1313 | tri/oligope | N |
| 425 | Soy | 336.1927 | 1 | 23.05415 | 198179.7 | 335.1854 | tri/oligope | N |
| 426 | Soy | 337.2352 | 1 | 36.15032 | 169785.6 | 336.2279 | tri/oligope | N |
| 427 | Soy | 340.1508 | 1 | 2.311637 | 114430.8 | 339.1435 | tri/oligope | N |
| 428 | Soy | 340.3579 | 1 | 37.43446 | 145976.5 | 339.3506 | tri/oligope | N |
| 429 | Soy | 342.2398 | 1 | 19.8871 | 126855.6 | 341.2325 | tri/oligope | N |
| 430 | Soy | 343.1982 | 1 | 2.41733 | 136709.1 | 342.1909 | tri/oligope | N |
| 431 | Soy | 344.1827 | 1 | 12.45655 | 179565.1 | 343.1754 | tri/oligope | N |
| 432 | Soy | 346.1613 | 1 | 2.19034 | 100429.1 | 345.154 | tri/oligope | N |
| 433 | Soy | 346.2345 | 1 | 14.85206 | 177085.4 | 345.2272 | tri/oligope | N |
| 434 | Soy | 346.2346 | 1 | 19.19143 | 156126 | 345.2274 | tri/oligope | N |
| 435 | Soy | 348.177 | 1 | 2.201593 | 274275.9 | 347.1697 | tri/oligope | N |
| 436 | Soy | 348.1772 | 1 | 2.567123 | 127681.7 | 347.1699 | tri/oligope | N |
| 437 | Soy | 350.1721 | 1 | 12.71654 | 150800.9 | 349.1649 | tri/oligope | N |
| 438 | Soy | 352.1513 | 1 | 11.9056 | 210962.2 | 351.144 | tri/oligope | N |
| 439 | Soy | 358.1725 | 1 | 1.92573 | 403893 | 357.1652 | tri/oligope | N |
| 440 | Soy | 358.2955 | 1 | 36.28755 | 151609 | 357.2882 | tri/oligope | N |
| 441 | Soy | 358.2955 | 1 | 36.73052 | 107532 | 357.2882 | tri/oligope | N |
| 442 | Soy | 359.2297 | 1 | 16.18152 | 352539 | 358.2224 | tri/oligope | N |
| 443 | Soy | 359.2298 | 1 | 15.84824 | 309432.8 | 358.2225 | tri/oligope | N |
| 444 | Soy | 359.2298 | 1 | 14.2936 | 340451.7 | 358.2225 | tri/oligope | N |
| 445 | Soy | 359.2299 | 1 | 19.47349 | 166400.9 | 358.2226 | tri/oligope | N |
| 446 | Soy | 360.1929 | 1 | 19.62243 | 119914.8 | 359.1857 | tri/oligope | N |
| 447 | Soy | 360.214 | 1 | 20.62003 | 278899.7 | 359.2067 | tri/oligope | N |
| 448 | Soy | 361.1721 | 1 | 2.201593 | 397165.1 | 360.1648 | tri/oligope | N |
| 449 | Soy | 361.2085 | 1 | 2.201593 | 167876.5 | 360.2012 | tri/oligope | N |
| 450 | Soy | 361.2353 | 1 | 37.32485 | 100079.1 | 360.228 | tri/oligope | N |
| 451 | Soy | 362.1564 | 1 | 2.567123 | 116495.4 | 361.1491 | tri/oligope | N |

| | | | | | | | | |
|---|---|---|---|---|---|---|---|---|
| 452 | Soy | 362.1927 | 1 | 2.286927 | 898104.6 | 361.1854 | tri/oligope | N |
| 453 | Soy | 367.1616 | 1 | 2.273767 | 158443.3 | 366.1543 | tri/oligope | N |
| 454 | Soy | 374.2036 | 1 | 2.19034 | 297330.1 | 373.1963 | tri/oligope | N |
| 455 | Soy | 374.2293 | 1 | 14.3877 | 559318.7 | 373.2221 | tri/oligope | N |
| 456 | Soy | 374.2298 | 1 | 21.19854 | 124354.5 | 373.2225 | tri/oligope | N |
| 457 | Soy | 377.183 | 1 | 17.7734 | 334161.5 | 376.1757 | tri/oligope | N |
| 458 | Soy | 383.4 | 1 | 36.25822 | 101096.4 | 382.3927 | tri/oligope | N |
| 459 | Soy | 387.2243 | 1 | 2.334193 | 150867.7 | 386.217 | tri/oligope | N |
| 460 | Soy | 387.2246 | 1 | 14.80274 | 113448.3 | 386.2173 | tri/oligope | N |
| 461 | Soy | 389.2034 | 1 | 2.201593 | 176955.5 | 388.1961 | tri/oligope | N |
| 462 | Soy | 389.2403 | 1 | 13.26386 | 359099.5 | 388.2331 | tri/oligope | N |
| 463 | Soy | 391.1986 | 1 | 16.68167 | 400316.2 | 390.1913 | tri/oligope | N |
| 464 | Soy | 391.2854 | 1 | 0.017617 | 807060.3 | 390.2781 | tri/oligope | N |
| 465 | Soy | 392.1859 | 1 | 11.57734 | 284448.3 | 391.1786 | tri/oligope | N |
| 466 | Soy | 392.3138 | 1 | 37.0313 | 111502.9 | 391.3065 | tri/oligope | N |
| 467 | Soy | 393.2142 | 1 | 22.46718 | 288010.4 | 392.2069 | tri/oligope | N |
| 468 | Soy | 399.2607 | 1 | 14.52741 | 981027.9 | 398.2535 | tri/oligope | N |
| 469 | Soy | 401.2405 | 1 | 13.81713 | 165767.6 | 400.2332 | tri/oligope | N |
| 470 | Soy | 403.2193 | 1 | 2.48644 | 106255.9 | 402.212 | tri/oligope | N |
| 471 | Soy | 405.1983 | 1 | 2.19034 | 119892.9 | 404.191 | tri/oligope | N |
| 472 | Soy | 405.2614 | 1 | 37.28021 | 233736.8 | 404.2542 | tri/oligope | N |
| 473 | Soy | 407.2138 | 1 | 2.201593 | 106349.9 | 406.2066 | tri/oligope | N |
| 474 | Soy | 413.2665 | 1 | 36.81405 | 392413.7 | 412.2592 | tri/oligope | N |
| 475 | Soy | 413.2673 | 1 | 38.08873 | 531728.3 | 412.26 | tri/oligope | N |
| 476 | Soy | 416.2514 | 1 | 15.70977 | 140963.3 | 415.2441 | tri/oligope | N |
| 477 | Soy | 417.119 | 1 | 22.56133 | 183226.8 | 416.1117 | tri/oligope | N |
| 478 | Soy | 417.2176 | 1 | 19.10422 | 120585.2 | 416.2103 | tri/oligope | N |
| 479 | Soy | 423.2356 | 1 | 36.2179 | 102304.5 | 422.2283 | tri/oligope | N |
| 480 | Soy | 424.183 | 1 | 2.322907 | 229124 | 423.1757 | tri/oligope | N |
| 481 | Soy | 428.3373 | 1 | 37.28021 | 109656.8 | 427.3301 | tri/oligope | N |
| 482 | Soy | 429.2415 | 1 | 38.08873 | 130363.9 | 428.2342 | tri/oligope | N |
| 483 | Soy | 429.3082 | 1 | 21.62759 | 229280.4 | 428.3009 | tri/oligope | N |
| 484 | Soy | 432.2091 | 1 | 2.19034 | 176859.1 | 431.2019 | tri/oligope | N |
| 485 | Soy | 433.1139 | 1 | 26.54081 | 317023.3 | 432.1066 | tri/oligope | N |
| 486 | Soy | 433.1939 | 1 | 10.45349 | 343244.4 | 432.1866 | tri/oligope | N |
| 487 | Soy | 433.1939 | 1 | 11.57734 | 289947 | 432.1867 | tri/oligope | N |
| 488 | Soy | 433.2305 | 1 | 12.88126 | 118103 | 432.2232 | tri/oligope | N |
| 489 | Soy | 434.2251 | 1 | 2.567123 | 141076.8 | 433.2178 | tri/oligope | N |
| 490 | Soy | 435.2248 | 1 | 18.38731 | 441463 | 434.2175 | tri/oligope | N |
| 491 | Soy | 441.2982 | 1 | 37.37668 | 191929.6 | 440.2909 | tri/oligope | N |
| 492 | Soy | 442.2661 | 1 | 2.247413 | 361273 | 441.2588 | tri/oligope | N |
| 493 | Soy | 445.2403 | 1 | 2.19034 | 175364.7 | 444.233 | tri/oligope | N |
| 494 | Soy | 446.2248 | 1 | 2.19034 | 117723.3 | 445.2176 | tri/oligope | N |
| 495 | Soy | 447.246 | 1 | 12.42705 | 204378.3 | 446.2388 | tri/oligope | N |
| 496 | Soy | 448.2405 | 1 | 2.322907 | 144010.1 | 447.2332 | tri/oligope | N |
| 497 | Soy | 449.2043 | 1 | 17.99405 | 302649.6 | 448.197 | tri/oligope | N |
| 498 | Soy | 449.2876 | 1 | 37.2412 | 172281.6 | 448.2803 | tri/oligope | N |
| 499 | Soy | 450.2358 | 1 | 19.51463 | 137181.5 | 449.2285 | tri/oligope | N |
| 500 | Soy | 451.2018 | 1 | 21.67102 | 410648.7 | 450.1946 | tri/oligope | N |
| 501 | Soy | 453.2458 | 1 | 2.19034 | 285198.2 | 452.2385 | tri/oligope | N |

| | | | | | | | | |
|---|---|---|---|---|---|---|---|---|
| 502 | Soy | 456.2826 | 1 | 16.64312 | 342760.5 | 455.2753 | tri/oligope | N |
| 503 | Soy | 456.2827 | 1 | 14.08747 | 390447.1 | 455.2754 | tri/oligope | N |
| 504 | Soy | 228.645 | 2 | 10.50599 | 113324.1 | 455.2755 | tri/oligope | N |
| 505 | Soy | 458.1996 | 1 | 2.156933 | 258021.6 | 457.1923 | tri/oligope | N |
| 506 | Soy | 458.2613 | 1 | 2.393167 | 204806.7 | 457.254 | tri/oligope | N |
| 507 | Soy | 460.2776 | 1 | 12.85695 | 263461.1 | 459.2703 | tri/oligope | N |
| 508 | Soy | 461.2615 | 1 | 13.84445 | 466241.1 | 460.2542 | tri/oligope | N |
| 509 | Soy | 462.1992 | 1 | 15.95769 | 103097.8 | 461.1919 | tri/oligope | N |
| 510 | Soy | 462.2197 | 1 | 2.201593 | 157100.1 | 461.2124 | tri/oligope | N |
| 511 | Soy | 462.2205 | 1 | 12.2992 | 225950.9 | 461.2132 | tri/oligope | N |
| 512 | Soy | 463.2197 | 1 | 16.26486 | 216110.2 | 462.2124 | tri/oligope | N |
| 513 | Soy | 463.2198 | 1 | 15.12985 | 141590.3 | 462.2125 | tri/oligope | N |
| 514 | Soy | 464.3738 | 1 | 37.3567 | 197899.8 | 463.3665 | tri/oligope | N |
| 515 | Soy | 467.2148 | 1 | 18.19269 | 144028.4 | 466.2075 | tri/oligope | N |
| 516 | Soy | 468.4415 | 1 | 37.6771 | 119760 | 467.4342 | tri/oligope | N |
| 517 | Soy | 235.1239 | 2 | 2.19034 | 125070.9 | 468.2333 | tri/oligope | N |
| 518 | Soy | 472.2774 | 1 | 16.0576 | 126734.6 | 471.2701 | tri/oligope | N |
| 519 | Soy | 474.2569 | 1 | 15.74701 | 165848.5 | 473.2496 | tri/oligope | N |
| 520 | Soy | 237.6499 | 2 | 2.311637 | 119506.2 | 473.2853 | tri/oligope | N |
| 521 | Soy | 476.2353 | 1 | 2.19034 | 106308.7 | 475.228 | tri/oligope | N |
| 522 | Soy | 240.1528 | 2 | 14.76493 | 739608 | 478.2911 | tri/oligope | N |
| 523 | Soy | 489.2305 | 1 | 2.247413 | 185123.2 | 488.2232 | tri/oligope | N |
| 524 | Soy | 490.1976 | 1 | 13.43423 | 184135.9 | 489.1904 | tri/oligope | N |
| 525 | Soy | 249.1453 | 2 | 2.02972 | 200106.4 | 496.276 | tri/oligope | N |
| 526 | Soy | 250.1534 | 2 | 2.659453 | 123786.6 | 498.2921 | tri/oligope | N |
| 527 | Soy | 250.6456 | 2 | 10.72989 | 186060.5 | 499.2766 | tri/oligope | N |
| 528 | Soy | 500.3083 | 1 | 18.87005 | 2235992 | 499.3011 | tri/oligope | N |
| 529 | Soy | 501.2306 | 1 | 2.156933 | 164785.3 | 500.2233 | tri/oligope | N |
| 530 | Soy | 503.1197 | 1 | 27.02633 | 129593.9 | 502.1124 | tri/oligope | N |
| 531 | Soy | 504.2302 | 1 | 2.212967 | 109880.6 | 503.2229 | tri/oligope | N |
| 532 | Soy | 504.2669 | 1 | 14.52741 | 116079.8 | 503.2596 | tri/oligope | N |
| 533 | Soy | 504.2671 | 1 | 16.66691 | 162236.9 | 503.2598 | tri/oligope | N |
| 534 | Soy | 504.2824 | 1 | 23.7609 | 766611.4 | 503.2751 | tri/oligope | N |
| 535 | Soy | 510.2566 | 1 | 17.70748 | 147006.4 | 509.2493 | tri/oligope | N |
| 536 | Soy | 512.3447 | 1 | 26.08693 | 1336486 | 511.3374 | tri/oligope | N |
| 537 | Soy | 257.1451 | 2 | 16.68167 | 166009.7 | 512.2757 | tri/oligope | N |
| 538 | Soy | 516.2674 | 1 | 13.32147 | 218364 | 515.2601 | tri/oligope | N |
| 539 | Soy | 258.6507 | 2 | 14.77512 | 132502.6 | 515.2868 | tri/oligope | N |
| 540 | Soy | 518.2465 | 1 | 14.77512 | 636732.5 | 517.2392 | tri/oligope | N |
| 541 | Soy | 519.1146 | 1 | 29.53444 | 322883.2 | 518.1073 | tri/oligope | N |
| 542 | Soy | 531.2784 | 1 | 12.59817 | 350125.9 | 530.2711 | tri/oligope | N |
| 543 | Soy | 533.2204 | 1 | 2.26061 | 132444.3 | 532.2131 | tri/oligope | N |
| 544 | Soy | 538.2518 | 1 | 14.27157 | 160234.2 | 537.2445 | tri/oligope | N |
| 545 | Soy | 544.236 | 1 | 2.19034 | 128038.5 | 543.2287 | tri/oligope | N |
| 546 | Soy | 546.3298 | 1 | 25.88982 | 238637.3 | 545.3225 | tri/oligope | N |
| 547 | Soy | 549.268 | 1 | 17.97144 | 283095.8 | 548.2607 | tri/oligope | N |
| 548 | Soy | 275.6716 | 2 | 18.93769 | 182808.9 | 549.3287 | tri/oligope | N |
| 549 | Soy | 278.1793 | 2 | 19.2791 | 141690.6 | 554.3441 | tri/oligope | N |
| 550 | Soy | 555.3515 | 1 | 19.2791 | 229247.9 | 554.3442 | tri/oligope | N |
| 551 | Soy | 558.2783 | 1 | 15.92095 | 127781.5 | 557.271 | tri/oligope | N |

| | | | | | | | | |
|---|---|---|---|---|---|---|---|---|
| 552 | Soy | 560.2681 | 1 | 2.356947 | 160404.4 | 559.2608 | tri/oligope | N |
| 553 | Soy | 563.2108 | 1 | 16.50597 | 132138.5 | 562.2035 | tri/oligope | N |
| 554 | Soy | 566.2471 | 1 | 16.11219 | 212987 | 565.2398 | tri/oligope | N |
| 555 | Soy | 569.3675 | 1 | 20.84264 | 116682.5 | 568.3602 | tri/oligope | N |
| 556 | Soy | 575.2635 | 1 | 30.02411 | 142309 | 574.2562 | tri/oligope | N |
| 557 | Soy | 289.6873 | 2 | 23.36966 | 224559.8 | 577.36 | tri/oligope | N |
| 558 | Soy | 292.6562 | 2 | 13.76391 | 115389.9 | 583.2979 | tri/oligope | N |
| 559 | Soy | 293.6635 | 2 | 2.179287 | 147953.9 | 585.3124 | tri/oligope | N |
| 560 | Soy | 294.1617 | 2 | 12.047 | 125124.6 | 586.3088 | tri/oligope | N |
| 561 | Soy | 588.2892 | 1 | 13.99992 | 127536.8 | 587.2819 | tri/oligope | N |
| 562 | Soy | 295.1511 | 2 | 2.632187 | 338823 | 588.2877 | tri/oligope | N |
| 563 | Soy | 591.2624 | 1 | 2.19034 | 170070.5 | 590.2552 | tri/oligope | N |
| 564 | Soy | 591.2636 | 1 | 11.15505 | 159559.8 | 590.2563 | tri/oligope | N |
| 565 | Soy | 594.2786 | 1 | 17.15513 | 114437.6 | 593.2713 | tri/oligope | N |
| 566 | Soy | 605.2689 | 1 | 18.87751 | 183191.5 | 604.2616 | tri/oligope | N |
| 567 | Soy | 607.274 | 1 | 18.06415 | 116464.8 | 606.2667 | tri/oligope | N |
| 568 | Soy | 611.2325 | 1 | 14.11505 | 120093.8 | 610.2252 | tri/oligope | N |
| 569 | Soy | 307.1771 | 2 | 19.22292 | 658131.1 | 612.3397 | tri/oligope | N |
| 570 | Soy | 619.2948 | 1 | 15.82979 | 154375.6 | 618.2876 | oligopepti | N |
| 571 | Soy | 310.1643 | 2 | 15.70977 | 173655.7 | 618.3141 | oligopepti | N |
| 572 | Soy | 626.284 | 1 | 20.48573 | 126675.3 | 625.2768 | oligopepti | N |
| 573 | Soy | 315.6954 | 2 | 2.659453 | 658334 | 629.3762 | oligopepti | N |
| 574 | Soy | 632.2902 | 1 | 15.15773 | 207929.8 | 631.2829 | oligopepti | N |
| 575 | Soy | 644.363 | 1 | 17.35972 | 112491.3 | 643.3558 | oligopepti | N |
| 576 | Soy | 322.6852 | 2 | 17.35111 | 152611.8 | 643.3559 | oligopepti | N |
| 577 | Soy | 323.1723 | 2 | 15.95175 | 181219.8 | 644.33 | oligopepti | N |
| 578 | Soy | 646.2982 | 1 | 2.19034 | 119220.6 | 645.2909 | oligopepti | N |
| 579 | Soy | 323.6529 | 2 | 2.19034 | 342555.7 | 645.2913 | oligopepti | N |
| 580 | Soy | 325.1459 | 2 | 15.97963 | 187684.6 | 648.2772 | oligopepti | N |
| 581 | Soy | 328.6777 | 2 | 2.006527 | 123850.5 | 655.3409 | oligopepti | N |
| 582 | Soy | 329.1815 | 2 | 2.212967 | 256298.4 | 656.3484 | oligopepti | N |
| 583 | Soy | 672.3212 | 1 | 16.68854 | 481535.7 | 671.3139 | oligopepti | N |
| 584 | Soy | 336.6644 | 2 | 16.68854 | 122183.8 | 671.3142 | oligopepti | N |
| 585 | Soy | 672.3327 | 1 | 15.74701 | 102210 | 671.3254 | oligopepti | N |
| 586 | Soy | 678.4204 | 1 | 23.16424 | 767609.8 | 677.4132 | oligopepti | N |
| 587 | Soy | 339.714 | 2 | 23.17124 | 869239.5 | 677.4134 | oligopepti | N |
| 588 | Soy | 679.332 | 1 | 24.3967 | 190237.6 | 678.3248 | oligopepti | N |
| 589 | Soy | 682.3788 | 1 | 16.85962 | 204501.4 | 681.3715 | oligopepti | N |
| 590 | Soy | 341.6932 | 2 | 16.85121 | 255372.7 | 681.3719 | oligopepti | N |
| 591 | Soy | 348.6434 | 2 | 16.68854 | 168099.8 | 695.2722 | oligopepti | N |
| 592 | Soy | 704.3481 | 1 | 18.20143 | 118567.9 | 703.3409 | oligopepti | N |
| 593 | Soy | 356.1569 | 2 | 2.099537 | 157231.6 | 710.2992 | oligopepti | N |
| 594 | Soy | 357.2199 | 2 | 16.84442 | 114286.2 | 712.4252 | oligopepti | N |
| 595 | Soy | 358.6955 | 2 | 2.212967 | 141567.5 | 715.3764 | oligopepti | N |
| 596 | Soy | 362.7218 | 2 | 2.273767 | 164424.6 | 723.429 | oligopepti | N |
| 597 | Soy | 365.2172 | 2 | 16.6275 | 206718.9 | 728.4199 | oligopepti | N |
| 598 | Soy | 365.6779 | 2 | 2.12261 | 137861.5 | 729.3413 | oligopepti | N |
| 599 | Soy | 731.3222 | 1 | 13.28589 | 118213.4 | 730.3149 | oligopepti | N |
| 600 | Soy | 368.686 | 2 | 19.13773 | 377577.8 | 735.3574 | oligopepti | N |
| 601 | Soy | 371.2067 | 2 | 17.236 | 220506.3 | 740.3989 | oligopepti | N |

| | | | | | | | | |
|---|---|---|---|---|---|---|---|---|
| 602 | Soy | 769.4106 | 1 | 16.14967 | 612208.9 | 768.4034 | oligopepti( | N |
| 603 | Soy | 385.209 | 2 | 16.13606 | 758456 | 768.4035 | oligopepti( | N |
| 604 | Soy | 386.2037 | 2 | 2.168163 | 195971.6 | 770.3929 | oligopepti( | N |
| 605 | Soy | 788.344 | 1 | 13.742 | 127096.5 | 787.3367 | oligopepti( | N |
| 606 | Soy | 396.2254 | 2 | 20.99711 | 108812.2 | 790.4363 | oligopepti( | N |
| 607 | Soy | 397.1884 | 2 | 16.14281 | 112376.4 | 792.3622 | oligopepti( | N |
| 608 | Soy | 398.1863 | 2 | 13.02295 | 479441.3 | 794.358 | oligopepti( | N |
| 609 | Soy | 402.7173 | 2 | 14.01437 | 213871.5 | 803.42 | oligopepti( | N |
| 610 | Soy | 403.6898 | 2 | 2.356947 | 318453.5 | 805.365 | oligopepti( | N |
| 611 | Soy | 409.1783 | 2 | 16.60027 | 117503.2 | 816.342 | oligopepti( | N |
| 612 | Soy | 415.741 | 2 | 16.7314 | 253593.5 | 829.4674 | oligopepti( | N |
| 613 | Soy | 419.7332 | 2 | 19.34892 | 258548.6 | 837.4518 | oligopepti( | N |
| 614 | Soy | 283.158 | 3 | 17.1861 | 131216.6 | 846.452 | oligopepti( | N |
| 615 | Soy | 427.723 | 2 | 15.7727 | 135753 | 853.4315 | oligopepti( | N |
| 616 | Soy | 429.2385 | 2 | 27.05431 | 170344.6 | 856.4625 | oligopepti( | N |
| 617 | Soy | 430.2181 | 2 | 13.69893 | 186728.4 | 858.4216 | oligopepti( | N |
| 618 | Soy | 435.2176 | 2 | 16.49981 | 271066.6 | 868.4207 | oligopepti( | N |
| 619 | Soy | 449.1968 | 2 | 14.52741 | 260827.5 | 896.3791 | oligopepti( | N |
| 620 | Soy | 913.5166 | 1 | 35.22043 | 329495.8 | 912.5093 | oligopepti( | N |
| 621 | Soy | 460.6665 | 2 | 14.27896 | 199116.5 | 919.3185 | oligopepti( | N |
| 622 | Soy | 943.5273 | 1 | 34.96775 | 1035471 | 942.5201 | oligopepti( | N |
| 623 | Soy | 476.2074 | 2 | 17.26986 | 238983.3 | 950.4002 | oligopepti( | N |
| 624 | Soy | 478.7365 | 2 | 26.97019 | 118655.1 | 955.4584 | oligopepti( | N |
| 625 | Soy | 959.5215 | 1 | 34.81903 | 110070.8 | 958.5143 | oligopepti( | N |
| 626 | Soy | 487.2237 | 2 | 20.67789 | 171989 | 972.4328 | oligopepti( | N |
| 627 | Soy | 493.2648 | 2 | 17.72728 | 118773.4 | 984.515 | oligopepti( | N |
| 628 | Soy | 493.268 | 2 | 27.34778 | 148044.3 | 984.5214 | oligopepti( | N |
| 629 | Soy | 332.1811 | 3 | 23.50908 | 381719.1 | 993.5215 | oligopepti( | N |
| 630 | Soy | 332.1917 | 3 | 16.21439 | 284855.6 | 993.5533 | oligopepti( | N |
| 631 | Soy | 501.7313 | 2 | 17.51042 | 172569.1 | 1001.448 | oligopepti( | N |
| 632 | Soy | 503.7652 | 2 | 34.96118 | 123913.1 | 1005.516 | oligopepti( | N |
| 633 | Soy | 511.754 | 2 | 34.96118 | 265755 | 1021.493 | oligopepti( | N |
| 634 | Soy | 513.7778 | 2 | 16.12065 | 302429.8 | 1025.541 | oligopepti( | N |
| 635 | Soy | 554.7502 | 2 | 16.26486 | 133913.6 | 1107.486 | oligopepti( | N |
| 636 | Soy | 557.2945 | 2 | 17.33522 | 614842.3 | 1112.574 | oligopepti( | N |
| 637 | Soy | 374.8675 | 3 | 24.06618 | 449052.1 | 1121.581 | oligopepti( | N |
| 638 | Soy | 590.2686 | 2 | 16.41307 | 176978.4 | 1178.523 | oligopepti( | N |
| 639 | Soy | 417.5537 | 3 | 23.77648 | 105552 | 1249.639 | oligopepti( | N |
| 640 | Soy | 646.811 | 2 | 18.76811 | 141789.4 | 1291.608 | oligopepti( | N |
| 641 | Soy | 678.8014 | 2 | 17.59616 | 119280.3 | 1355.588 | oligopepti( | N |
| 642 | Soy | 580.9133 | 3 | 19.10422 | 115189 | 1739.718 | oligopepti( | N |

| ID | Protein | mz | charge | retentiont | raw.abund | mass | type | Match |
|---|---|---|---|---|---|---|---|---|
| 2 | NPP | 231.1707 | 1 | 5.4642 | 1515101 | 230.1635 | di/tripepti | Y |
| 3 | NPP | 231.1708 | 1 | 9.72845 | 736192.6 | 230.1635 | di/tripepti | Y |
| 4 | NPP | 231.1708 | 1 | 12.05358 | 651257.5 | 230.1635 | di/tripepti | Y |
| 5 | NPP | 233.1497 | 1 | 2.184133 | 341008.1 | 232.1424 | di/tripepti | Y |
| 6 | NPP | 233.1498 | 1 | 2.78445 | 410421.4 | 232.1425 | di/tripepti | Y |
| 11 | NPP | 245.1862 | 1 | 16.42353 | 1479840 | 244.1789 | di/tripepti | Y |
| 12 | NPP | 245.1864 | 1 | 15.85914 | 252562.9 | 244.1791 | di/tripepti | Y |
| 13 | NPP | 245.1862 | 1 | 14.57899 | 348032.5 | 244.1789 | di/tripepti | Y |
| 14 | NPP | 245.1865 | 1 | 17.75511 | 233386.4 | 244.1792 | di/tripepti | Y |
| 15 | NPP | 246.145 | 1 | 2.184133 | 472728.6 | 245.1377 | di/tripepti | Y |
| 15 | NPP | 246.145 | 1 | 1.945975 | 181632.8 | 245.1377 | di/tripepti | Y |
| 16 | NPP | 246.145 | 1 | 2.661525 | 666372.7 | 245.1378 | di/tripepti | Y |
| 17 | NPP | 246.1452 | 1 | 3.381733 | 717288.2 | 245.1379 | di/tripepti | Y |
| 20 | NPP | 247.1289 | 1 | 2.125975 | 652483.5 | 246.1217 | di/tripepti | Y |
| 21 | NPP | 247.1293 | 1 | 3.156692 | 278038.8 | 246.122 | di/tripepti | Y |
| 22 | NPP | 247.1293 | 1 | 4.7487 | 647111.3 | 246.122 | di/tripepti | Y |
| 27 | NPP | 253.1185 | 1 | 2.282258 | 591297.9 | 252.1112 | di/tripepti | Y |
| 27 | NPP | 253.1186 | 1 | 2.530733 | 110511.6 | 252.1113 | di/tripepti | Y |
| 30 | NPP | 260.1607 | 1 | 2.184133 | 659091.1 | 259.1534 | di/tripepti | Y |
| 31 | NPP | 260.1609 | 1 | 2.668133 | 205801.9 | 259.1536 | di/tripepti | Y |
| 33 | NPP | 260.197 | 1 | 1.848725 | 129979.6 | 259.1897 | di/tripepti | Y |
| 34 | NPP | 261.1449 | 1 | 2.195508 | 717756.2 | 260.1376 | di/tripepti | Y |
| 35 | NPP | 261.145 | 1 | 3.285542 | 296214.5 | 260.1377 | di/tripepti | Y |
| 36 | NPP | 261.145 | 1 | 5.48795 | 424495.7 | 260.1377 | di/tripepti | Y |
| 40 | NPP | 265.1552 | 1 | 16.2552 | 138595.1 | 264.1479 | di/tripepti | Y |
| 41 | NPP | 267.1343 | 1 | 2.423283 | 157992.1 | 266.127 | di/tripepti | Y |
| 42 | NPP | 269.1609 | 1 | 2.172858 | 243514.7 | 268.1536 | di/tripepti | Y |
| 44 | NPP | 277.1032 | 1 | 1.857208 | 168710.5 | 276.0959 | di/tripepti | Y |
| 45 | NPP | 281.1135 | 1 | 2.588825 | 389729.8 | 280.1063 | di/tripepti | Y |
| 47 | NPP | 288.1924 | 1 | 16.0521 | 144145.5 | 287.1851 | di/tripepti | Y |
| 50 | NPP | 294.1451 | 1 | 2.327933 | 265958.5 | 293.1378 | di/tripepti | Y |
| 51 | NPP | 295.1295 | 1 | 3.263533 | 679257.6 | 294.1222 | di/tripepti | Y |
| 52 | NPP | 295.1657 | 1 | 16.42353 | 342459.1 | 294.1584 | di/tripepti | Y |
| 53 | NPP | 300.1923 | 1 | 2.628217 | 122520.2 | 299.1851 | di/tripepti | Y |
| 56 | NPP | 302.2081 | 1 | 15.14889 | 486336 | 301.2008 | tri/oligope | Y |
| 57 | NPP | 302.2081 | 1 | 19.86926 | 202738.4 | 301.2008 | tri/oligope | Y |
| 58 | NPP | 303.1665 | 1 | 2.053983 | 114727.5 | 302.1593 | tri/oligope | Y |
| 58 | NPP | 303.1665 | 1 | 2.184133 | 379655.4 | 302.1593 | tri/oligope | Y |
| 59 | NPP | 304.1507 | 1 | 2.2177 | 190934.8 | 303.1434 | tri/oligope | Y |
| 63 | NPP | 318.1665 | 1 | 2.749983 | 181141 | 317.1593 | tri/oligope | Y |
| 64 | NPP | 318.1663 | 1 | 2.195508 | 283953.6 | 317.1591 | tri/oligope | Y |
| 65 | NPP | 318.1668 | 1 | 9.168917 | 201801.8 | 317.1596 | tri/oligope | Y |
| 71 | NPP | 330.2395 | 1 | 18.12856 | 280206.2 | 329.2323 | tri/oligope | Y |
| 73 | NPP | 331.1657 | 1 | 2.841792 | 234401.9 | 330.1584 | tri/oligope | Y |
| 75 | NPP | 332.1822 | 1 | 2.588825 | 184379.5 | 331.1749 | tri/oligope | Y |
| 78 | NPP | 333.1407 | 1 | 1.889925 | 163405.8 | 332.1334 | tri/oligope | Y |
| 80 | NPP | 334.1613 | 1 | 2.315167 | 212238.7 | 333.154 | tri/oligope | Y |
| 85 | NPP | 344.2554 | 1 | 21.62794 | 178490.3 | 343.2481 | tri/oligope | Y |
| 89 | NPP | 358.2713 | 1 | 24.02246 | 124911.2 | 357.264 | tri/oligope | Y |

| | | | | | | | | |
|---|---|---|---|---|---|---|---|---|
| 92 | NPP | 360.2136 | 1 | 2.749983 | 125244.3 | 359.2063 | tri/oligope | Y |
| 94 | NPP | 363.1555 | 1 | 2.184133 | 109533.2 | 362.1482 | tri/oligope | Y |
| 96 | NPP | 366.2139 | 1 | 2.195508 | 114099.8 | 365.2066 | tri/oligope | Y |
| 105 | NPP | 390.1876 | 1 | 2.341375 | 223645.6 | 389.1804 | tri/oligope | Y |
| 106 | NPP | 395.0393 | 1 | 1.597967 | 157673.8 | 394.032 | tri/oligope | Y |
| 107 | NPP | 399.2608 | 1 | 18.79728 | 533289.8 | 398.2535 | tri/oligope | Y |
| 110 | NPP | 405.1619 | 1 | 2.029983 | 113014.7 | 404.1546 | tri/oligope | Y |
| 112 | NPP | 415.2118 | 1 | 36.47671 | 268253.1 | 414.2045 | tri/oligope | Y |
| 133 | NPP | 461.3005 | 1 | 37.3895 | 108434.6 | 460.2932 | tri/oligope | Y |
| 134 | NPP | 463.0267 | 1 | 1.607792 | 202332.2 | 462.0194 | tri/oligope | Y |
| 141 | NPP | 479.3102 | 1 | 37.13064 | 159392.8 | 478.303 | tri/oligope | Y |
| 144 | NPP | 488.2516 | 1 | 19.10416 | 183986.7 | 487.2443 | tri/oligope | Y |
| 151 | NPP | 500.2882 | 1 | 25.1948 | 106577.9 | 499.2809 | tri/oligope | Y |
| 156 | NPP | 515.2914 | 1 | 37.0834 | 107918.8 | 514.2842 | tri/oligope | Y |
| 159 | NPP | 520.278 | 1 | 21.17941 | 104174.9 | 519.2707 | tri/oligope | Y |
| 161 | NPP | 264.1747 | 2 | 35.30205 | 105218.5 | 526.3349 | tri/oligope | Y |
| 165 | NPP | 531.014 | 1 | 1.61735 | 258427.7 | 530.0068 | tri/oligope | Y |
| 166 | NPP | 531.3871 | 1 | 37.8434 | 103832.9 | 530.3798 | tri/oligope | Y |
| 174 | NPP | 274.1676 | 2 | 35.92205 | 115058.8 | 546.3207 | tri/oligope | Y |
| 194 | NPP | 599.0019 | 1 | 1.607792 | 239198.1 | 597.9946 | tri/oligope | Y |
| 218 | NPP | 666.9893 | 1 | 1.61735 | 224842.4 | 665.982 | oligopepti | Y |
| 240 | NPP | 734.9768 | 1 | 1.61735 | 197464.7 | 733.9696 | oligopepti | Y |
| 258 | NPP | 802.9643 | 1 | 1.607792 | 172291.1 | 801.9571 | oligopepti | Y |
| 276 | NPP | 870.9516 | 1 | 1.61735 | 119727.8 | 869.9443 | oligopepti | Y |
| 296 | NPP | 974.814 | 1 | 1.62715 | 251111.4 | 973.8067 | oligopepti | Y |
| 297 | NPP | 990.7886 | 1 | 1.62715 | 155436.1 | 989.7813 | oligopepti | Y |
| 307 | NPP | 548.9516 | 2 | 1.61735 | 100103.1 | 1095.889 | oligopepti | Y |
| 316 | NPP | 616.9395 | 2 | 1.61735 | 101630.3 | 1231.864 | oligopepti | Y |
| 318 | NPP | 624.9264 | 2 | 1.62715 | 100080.2 | 1247.838 | oligopepti | Y |
| 321 | NPP | 650.9331 | 2 | 1.61735 | 102783.4 | 1299.852 | oligopepti | Y |
| 338 | NPP | 223.1081 | 1 | 5.189033 | 574594.9 | 222.1008 | dipeptide | N |
| 339 | NPP | 223.1081 | 1 | 10.47455 | 107394.7 | 222.1009 | dipeptide | N |
| 340 | NPP | 226.9517 | 1 | 1.568358 | 393883.3 | 225.9444 | di/tripepti | N |
| 341 | NPP | 229.1551 | 1 | 5.037183 | 182056 | 228.1479 | di/tripepti | N |
| 342 | NPP | 229.1552 | 1 | 9.049017 | 271821.6 | 228.1479 | di/tripepti | N |
| 343 | NPP | 229.1552 | 1 | 12.00996 | 172618.5 | 228.1479 | di/tripepti | N |
| 344 | NPP | 231.1708 | 1 | 9.010442 | 352441.2 | 230.1635 | di/tripepti | N |
| 345 | NPP | 232.1292 | 1 | 2.184133 | 197152.8 | 231.122 | di/tripepti | N |
| 346 | NPP | 233.1133 | 1 | 2.184133 | 146504 | 232.106 | di/tripepti | N |
| 347 | NPP | 233.1133 | 1 | 1.993883 | 144206.4 | 232.106 | di/tripepti | N |
| 348 | NPP | 233.15 | 1 | 4.118817 | 160356.5 | 232.1427 | di/tripepti | N |
| 349 | NPP | 237.1238 | 1 | 4.701592 | 210455.1 | 236.1165 | di/tripepti | N |
| 350 | NPP | 237.1238 | 1 | 11.24586 | 163343.2 | 236.1165 | di/tripepti | N |
| 351 | NPP | 246.1453 | 1 | 5.404925 | 185955.7 | 245.138 | di/tripepti | N |
| 352 | NPP | 258.1103 | 1 | 1.739042 | 130248.4 | 257.103 | di/tripepti | N |
| 353 | NPP | 263.1398 | 1 | 14.4688 | 237641.9 | 262.1325 | di/tripepti | N |
| 354 | NPP | 265.1551 | 1 | 14.76754 | 283251 | 264.1478 | di/tripepti | N |
| 355 | NPP | 272.1605 | 1 | 2.282258 | 125230.9 | 271.1533 | di/tripepti | N |
| 356 | NPP | 276.1192 | 1 | 1.79985 | 133843.8 | 275.1119 | di/tripepti | N |
| 357 | NPP | 276.1558 | 1 | 2.195508 | 205415.8 | 275.1485 | di/tripepti | N |

| | | | | | | | | |
|---|---|---|---|---|---|---|---|---|
| 358 | NPP | 279.1709 | 1 | 19.90808 | 1356473 | 278.1637 | di/tripepti | N |
| 359 | NPP | 279.171 | 1 | 18.99346 | 155435.9 | 278.1637 | di/tripepti | N |
| 360 | NPP | 279.1711 | 1 | 21.0258 | 111122.1 | 278.1638 | di/tripepti | N |
| 361 | NPP | 279.2321 | 1 | 37.01214 | 207627.2 | 278.2248 | di/tripepti | N |
| 362 | NPP | 281.1137 | 1 | 11.59373 | 308329.8 | 280.1065 | di/tripepti | N |
| 363 | NPP | 281.1502 | 1 | 6.208517 | 332837.1 | 280.143 | di/tripepti | N |
| 364 | NPP | 286.1768 | 1 | 13.46579 | 368354.5 | 285.1695 | di/tripepti | N |
| 365 | NPP | 286.1768 | 1 | 10.93262 | 171071.1 | 285.1696 | di/tripepti | N |
| 366 | NPP | 288.192 | 1 | 2.239233 | 177076.9 | 287.1847 | di/tripepti | N |
| 367 | NPP | 288.1923 | 1 | 14.68307 | 186736.7 | 287.185 | di/tripepti | N |
| 368 | NPP | 288.2032 | 1 | 1.98205 | 100580.4 | 287.1959 | di/tripepti | N |
| 369 | NPP | 290.1716 | 1 | 2.195508 | 257118.6 | 289.1643 | di/tripepti | N |
| 370 | NPP | 295.1297 | 1 | 12.65775 | 100061.4 | 294.1224 | di/tripepti | N |
| 371 | NPP | 295.1657 | 1 | 14.63573 | 158996.1 | 294.1585 | di/tripepti | N |
| 372 | NPP | 295.166 | 1 | 13.43193 | 133661.2 | 294.1587 | di/tripepti | N |
| 373 | NPP | 295.2272 | 1 | 37.37733 | 154952.2 | 294.2199 | di/tripepti | N |
| 374 | NPP | 296.1243 | 1 | 2.184133 | 121368.9 | 295.1171 | di/tripepti | N |
| 375 | NPP | 297.1086 | 1 | 2.69675 | 100004.8 | 296.1013 | di/tripepti | N |
| 376 | NPP | 297.2427 | 1 | 36.14295 | 156280.3 | 296.2354 | di/tripepti | N |
| 377 | NPP | 304.1869 | 1 | 2.195508 | 136789.1 | 303.1796 | tri/oligope | N |
| 378 | NPP | 304.1871 | 1 | 2.423283 | 144656.7 | 303.1799 | tri/oligope | N |
| 379 | NPP | 306.1297 | 1 | 1.857208 | 129126.7 | 305.1224 | tri/oligope | N |
| 380 | NPP | 306.1457 | 1 | 10.77991 | 330152 | 305.1385 | tri/oligope | N |
| 381 | NPP | 306.1662 | 1 | 2.184133 | 124763.2 | 305.1589 | tri/oligope | N |
| 382 | NPP | 310.1402 | 1 | 2.588825 | 158520 | 309.1329 | tri/oligope | N |
| 383 | NPP | 311.1239 | 1 | 2.184133 | 137107.8 | 310.1167 | tri/oligope | N |
| 384 | NPP | 311.1243 | 1 | 2.759667 | 120526.6 | 310.117 | tri/oligope | N |
| 385 | NPP | 313.2376 | 1 | 35.81866 | 235696.7 | 312.2303 | tri/oligope | N |
| 386 | NPP | 314.2081 | 1 | 13.69944 | 107042.8 | 313.2008 | tri/oligope | N |
| 387 | NPP | 315.2533 | 1 | 36.14295 | 394626.6 | 314.246 | tri/oligope | N |
| 388 | NPP | 316.224 | 1 | 19.62116 | 368080.9 | 315.2167 | tri/oligope | N |
| 389 | NPP | 317.1823 | 1 | 2.195508 | 276406.6 | 316.175 | tri/oligope | N |
| 390 | NPP | 317.1826 | 1 | 2.508375 | 118050.1 | 316.1753 | tri/oligope | N |
| 391 | NPP | 318.2027 | 1 | 2.341375 | 180588.4 | 317.1954 | tri/oligope | N |
| 392 | NPP | 320.1623 | 1 | 15.69466 | 122220.3 | 319.155 | tri/oligope | N |
| 393 | NPP | 327.0519 | 1 | 1.636808 | 178573.7 | 326.0446 | tri/oligope | N |
| 394 | NPP | 328.224 | 1 | 16.90748 | 195549.5 | 327.2168 | tri/oligope | N |
| 395 | NPP | 328.2241 | 1 | 18.06979 | 155212.7 | 327.2169 | tri/oligope | N |
| 396 | NPP | 328.2339 | 1 | 15.46508 | 110643.2 | 327.2266 | tri/oligope | N |
| 397 | NPP | 330.2394 | 1 | 17.32916 | 185466.6 | 329.2322 | tri/oligope | N |
| 398 | NPP | 330.2397 | 1 | 17.52208 | 227371.4 | 329.2324 | tri/oligope | N |
| 399 | NPP | 331.1984 | 1 | 3.57105 | 249362.9 | 330.1911 | tri/oligope | N |
| 400 | NPP | 331.2484 | 1 | 35.42086 | 126079.6 | 330.2412 | tri/oligope | N |
| 401 | NPP | 332.2188 | 1 | 18.61123 | 140842.2 | 331.2115 | tri/oligope | N |
| 402 | NPP | 332.2189 | 1 | 19.45903 | 169364.4 | 331.2116 | tri/oligope | N |
| 403 | NPP | 333.1771 | 1 | 2.206958 | 269154.8 | 332.1699 | tri/oligope | N |
| 404 | NPP | 334.1248 | 1 | 1.9232 | 107261.8 | 333.1175 | tri/oligope | N |
| 405 | NPP | 334.1406 | 1 | 18.00792 | 118081.3 | 333.1333 | tri/oligope | N |
| 406 | NPP | 334.1406 | 1 | 13.77534 | 161143.5 | 333.1333 | tri/oligope | N |
| 407 | NPP | 334.1616 | 1 | 5.19585 | 286155.7 | 333.1543 | tri/oligope | N |

| | | | | | | | | |
|---|---|---|---|---|---|---|---|---|
| 408 | NPP | 337.2352 | 1 | 36.14295 | 153814.6 | 336.2279 | tri/oligope | N |
| 409 | NPP | 338.1356 | 1 | 12.45395 | 297986.8 | 337.1283 | tri/oligope | N |
| 410 | NPP | 342.2399 | 1 | 21.27672 | 185564.8 | 341.2327 | tri/oligope | N |
| 411 | NPP | 343.1982 | 1 | 2.423283 | 387160.8 | 342.1909 | tri/oligope | N |
| 412 | NPP | 343.1986 | 1 | 11.17171 | 311730 | 342.1913 | tri/oligope | N |
| 413 | NPP | 345.1773 | 1 | 2.172858 | 120625.8 | 344.17 | tri/oligope | N |
| 414 | NPP | 345.2137 | 1 | 2.195508 | 136541.2 | 344.2065 | tri/oligope | N |
| 415 | NPP | 345.214 | 1 | 14.63121 | 126040.9 | 344.2067 | tri/oligope | N |
| 416 | NPP | 345.2141 | 1 | 14.79063 | 309375.5 | 344.2068 | tri/oligope | N |
| 417 | NPP | 345.2142 | 1 | 5.302083 | 188462.2 | 344.2069 | tri/oligope | N |
| 418 | NPP | 346.1613 | 1 | 2.184133 | 134265.2 | 345.154 | tri/oligope | N |
| 419 | NPP | 346.1979 | 1 | 2.487525 | 258575.8 | 345.1906 | tri/oligope | N |
| 420 | NPP | 346.1982 | 1 | 4.339742 | 536472.3 | 345.1909 | tri/oligope | N |
| 421 | NPP | 348.177 | 1 | 2.195508 | 335248.3 | 347.1697 | tri/oligope | N |
| 422 | NPP | 348.1775 | 1 | 8.369717 | 163892.2 | 347.1703 | tri/oligope | N |
| 423 | NPP | 357.2138 | 1 | 2.553883 | 594314.5 | 356.2065 | tri/oligope | N |
| 424 | NPP | 358.1984 | 1 | 12.67498 | 111568.9 | 357.1911 | tri/oligope | N |
| 425 | NPP | 358.2956 | 1 | 35.81866 | 110466.1 | 357.2884 | tri/oligope | N |
| 426 | NPP | 359.2298 | 1 | 15.79431 | 418962.8 | 358.2225 | tri/oligope | N |
| 427 | NPP | 359.2299 | 1 | 19.48459 | 132123.9 | 358.2226 | tri/oligope | N |
| 428 | NPP | 359.23 | 1 | 17.11709 | 214692.4 | 358.2227 | tri/oligope | N |
| 429 | NPP | 360.2138 | 1 | 19.86926 | 183715.4 | 359.2065 | tri/oligope | N |
| 430 | NPP | 360.2139 | 1 | 9.2388 | 141480 | 359.2066 | tri/oligope | N |
| 431 | NPP | 360.214 | 1 | 20.60992 | 396200.7 | 359.2067 | tri/oligope | N |
| 432 | NPP | 360.2141 | 1 | 21.26312 | 144711.6 | 359.2069 | tri/oligope | N |
| 433 | NPP | 361.1721 | 1 | 2.195508 | 310527.2 | 360.1648 | tri/oligope | N |
| 434 | NPP | 362.1564 | 1 | 2.571208 | 156423.6 | 361.1491 | tri/oligope | N |
| 435 | NPP | 362.1927 | 1 | 2.271408 | 325006 | 361.1854 | tri/oligope | N |
| 436 | NPP | 363.1514 | 1 | 1.857208 | 102268.5 | 362.1441 | tri/oligope | N |
| 437 | NPP | 364.1354 | 1 | 1.9232 | 106777.9 | 363.1281 | tri/oligope | N |
| 438 | NPP | 373.2086 | 1 | 2.596392 | 123569.9 | 372.2013 | tri/oligope | N |
| 439 | NPP | 374.2036 | 1 | 2.184133 | 112829.7 | 373.1963 | tri/oligope | N |
| 440 | NPP | 374.2293 | 1 | 14.40792 | 237718.2 | 373.2221 | tri/oligope | N |
| 441 | NPP | 375.1884 | 1 | 8.813642 | 100631 | 374.1812 | tri/oligope | N |
| 442 | NPP | 377.183 | 1 | 18.24747 | 152547.6 | 376.1757 | tri/oligope | N |
| 443 | NPP | 384.3479 | 1 | 37.72038 | 102555.8 | 383.3406 | tri/oligope | N |
| 444 | NPP | 385.2454 | 1 | 2.822575 | 140298.3 | 384.2381 | tri/oligope | N |
| 445 | NPP | 389.2034 | 1 | 2.195508 | 175661.5 | 388.1961 | tri/oligope | N |
| 446 | NPP | 391.1986 | 1 | 16.68108 | 137912.5 | 390.1913 | tri/oligope | N |
| 447 | NPP | 391.2854 | 1 | 0.022608 | 831216.5 | 390.2781 | tri/oligope | N |
| 448 | NPP | 397.1728 | 1 | 8.493083 | 153384.5 | 396.1655 | tri/oligope | N |
| 449 | NPP | 399.2607 | 1 | 14.54808 | 951788.5 | 398.2535 | tri/oligope | N |
| 450 | NPP | 399.2612 | 1 | 14.93016 | 440848.2 | 398.2539 | tri/oligope | N |
| 451 | NPP | 405.1983 | 1 | 2.184133 | 164728.3 | 404.191 | tri/oligope | N |
| 452 | NPP | 405.2614 | 1 | 37.27447 | 191094.5 | 404.2542 | tri/oligope | N |
| 453 | NPP | 407.1935 | 1 | 18.20371 | 100563.3 | 406.1862 | tri/oligope | N |
| 454 | NPP | 408.2137 | 1 | 17.47438 | 200597.9 | 407.2064 | tri/oligope | N |
| 455 | NPP | 408.2141 | 1 | 17.65273 | 442805.9 | 407.2068 | tri/oligope | N |
| 456 | NPP | 409.1722 | 1 | 2.250008 | 139746.2 | 408.1649 | tri/oligope | N |
| 457 | NPP | 409.2196 | 1 | 2.195508 | 292296.6 | 408.2124 | tri/oligope | N |

| | | | | | | | | |
|---|---|---|---|---|---|---|---|---|
| 458 | NPP | 413.2665 | 1 | 36.81423 | 617482.2 | 412.2592 | tri/oligope | N |
| 459 | NPP | 413.2673 | 1 | 38.0882 | 355736.6 | 412.26 | tri/oligope | N |
| 460 | NPP | 415.2354 | 1 | 25.40925 | 121804.5 | 414.2281 | tri/oligope | N |
| 461 | NPP | 416.2514 | 1 | 14.95028 | 164039.1 | 415.2442 | tri/oligope | N |
| 462 | NPP | 417.2351 | 1 | 14.63915 | 178808.7 | 416.2279 | tri/oligope | N |
| 463 | NPP | 418.1933 | 1 | 2.195508 | 191484.5 | 417.186 | tri/oligope | N |
| 464 | NPP | 427.2927 | 1 | 20.13854 | 368253.3 | 426.2854 | tri/oligope | N |
| 465 | NPP | 428.3373 | 1 | 37.27447 | 105769 | 427.3301 | tri/oligope | N |
| 466 | NPP | 431.1886 | 1 | 2.149642 | 111537 | 430.1813 | tri/oligope | N |
| 467 | NPP | 432.2091 | 1 | 2.184133 | 212902.2 | 431.2019 | tri/oligope | N |
| 468 | NPP | 432.2462 | 1 | 9.822708 | 120898.7 | 431.239 | tri/oligope | N |
| 469 | NPP | 432.2464 | 1 | 10.37308 | 206739.6 | 431.2391 | tri/oligope | N |
| 470 | NPP | 441.2982 | 1 | 37.36793 | 212518.1 | 440.2909 | tri/oligope | N |
| 471 | NPP | 447.2097 | 1 | 9.831692 | 122394.5 | 446.2024 | tri/oligope | N |
| 472 | NPP | 449.2876 | 1 | 37.23938 | 137064.7 | 448.2803 | tri/oligope | N |
| 473 | NPP | 450.2358 | 1 | 19.52118 | 311948.6 | 449.2285 | tri/oligope | N |
| 474 | NPP | 456.2826 | 1 | 16.64333 | 120297.6 | 455.2753 | tri/oligope | N |
| 475 | NPP | 457.2418 | 1 | 15.01508 | 184557.7 | 456.2345 | tri/oligope | N |
| 476 | NPP | 458.262 | 1 | 13.38733 | 481768.4 | 457.2547 | tri/oligope | N |
| 477 | NPP | 459.2454 | 1 | 2.580925 | 163969.7 | 458.2382 | tri/oligope | N |
| 478 | NPP | 462.2197 | 1 | 2.195508 | 136565 | 461.2124 | tri/oligope | N |
| 479 | NPP | 464.3738 | 1 | 37.34974 | 223209.9 | 463.3665 | tri/oligope | N |
| 480 | NPP | 467.1784 | 1 | 13.05894 | 180456.2 | 466.1711 | tri/oligope | N |
| 481 | NPP | 469.3294 | 1 | 37.10913 | 112459.7 | 468.3221 | tri/oligope | N |
| 482 | NPP | 470.2622 | 1 | 13.66411 | 225688 | 469.2549 | tri/oligope | N |
| 483 | NPP | 240.1528 | 2 | 14.76754 | 222726 | 478.2911 | tri/oligope | N |
| 484 | NPP | 483.2564 | 1 | 2.184133 | 163514 | 482.2491 | tri/oligope | N |
| 485 | NPP | 486.2936 | 1 | 17.81638 | 100673.5 | 485.2863 | tri/oligope | N |
| 486 | NPP | 487.2523 | 1 | 11.17171 | 340992.3 | 486.245 | tri/oligope | N |
| 487 | NPP | 488.2726 | 1 | 18.3599 | 278324 | 487.2653 | tri/oligope | N |
| 488 | NPP | 490.215 | 1 | 2.530733 | 244347.9 | 489.2077 | tri/oligope | N |
| 489 | NPP | 249.1314 | 2 | 13.1199 | 115325.9 | 496.2482 | tri/oligope | N |
| 490 | NPP | 502.2675 | 1 | 27.85682 | 139030.2 | 501.2603 | tri/oligope | N |
| 491 | NPP | 251.6532 | 2 | 2.282258 | 117664.9 | 501.2918 | tri/oligope | N |
| 492 | NPP | 504.2824 | 1 | 23.76166 | 1011572 | 503.2751 | tri/oligope | N |
| 493 | NPP | 504.2829 | 1 | 18.98219 | 117496.1 | 503.2756 | tri/oligope | N |
| 494 | NPP | 505.2673 | 1 | 23.12218 | 165403.7 | 504.26 | tri/oligope | N |
| 495 | NPP | 510.3299 | 1 | 22.55671 | 110105.8 | 509.3226 | tri/oligope | N |
| 496 | NPP | 512.3447 | 1 | 26.08698 | 2171451 | 511.3374 | tri/oligope | N |
| 497 | NPP | 512.3454 | 1 | 21.66851 | 534476.5 | 511.3381 | tri/oligope | N |
| 498 | NPP | 512.3456 | 1 | 28.23679 | 458661.4 | 511.3384 | tri/oligope | N |
| 499 | NPP | 518.2461 | 1 | 2.553883 | 258739.7 | 517.2388 | tri/oligope | N |
| 500 | NPP | 518.2465 | 1 | 14.77656 | 529331.1 | 517.2392 | tri/oligope | N |
| 501 | NPP | 532.2628 | 1 | 14.20099 | 170742.1 | 531.2555 | tri/oligope | N |
| 502 | NPP | 267.1503 | 2 | 2.066 | 168742.3 | 532.2861 | tri/oligope | N |
| 503 | NPP | 534.2932 | 1 | 23.85441 | 978422.5 | 533.2859 | tri/oligope | N |
| 504 | NPP | 536.2729 | 1 | 17.02177 | 107130.8 | 535.2656 | tri/oligope | N |
| 505 | NPP | 540.3768 | 1 | 27.65166 | 394720.5 | 539.3696 | tri/oligope | N |
| 506 | NPP | 541.3351 | 1 | 16.44775 | 515671.2 | 540.3278 | tri/oligope | N |
| 507 | NPP | 271.1713 | 2 | 16.44775 | 273344.4 | 540.3281 | tri/oligope | N |

| | | | | | | | | |
|---|---|---|---|---|---|---|---|---|
| 508 | NPP | 271.177 | 2 | 14.66273 | 269569.2 | 540.3394 | tri/oligope | N |
| 509 | NPP | 271.1898 | 2 | 19.28232 | 143268.8 | 540.365 | tri/oligope | N |
| 510 | NPP | 542.3564 | 1 | 26.24859 | 171292.3 | 541.3491 | tri/oligope | N |
| 511 | NPP | 546.3298 | 1 | 25.8821 | 1168667 | 545.3225 | tri/oligope | N |
| 512 | NPP | 546.3304 | 1 | 25.6061 | 237678.1 | 545.3231 | tri/oligope | N |
| 513 | NPP | 279.169 | 2 | 18.06213 | 274914.2 | 556.3234 | tri/oligope | N |
| 514 | NPP | 557.331 | 1 | 18.07555 | 284061.3 | 556.3237 | tri/oligope | N |
| 515 | NPP | 559.0208 | 1 | 1.646933 | 109179.1 | 558.0135 | tri/oligope | N |
| 516 | NPP | 570.3271 | 1 | 22.47389 | 241538.2 | 569.3199 | tri/oligope | N |
| 517 | NPP | 286.1512 | 2 | 1.945975 | 170102.5 | 570.2879 | tri/oligope | N |
| 518 | NPP | 571.3101 | 1 | 17.45358 | 124404.5 | 570.3028 | tri/oligope | N |
| 519 | NPP | 571.3104 | 1 | 17.62652 | 127075.6 | 570.3031 | tri/oligope | N |
| 520 | NPP | 573.3255 | 1 | 14.72428 | 160427.3 | 572.3183 | tri/oligope | N |
| 521 | NPP | 577.2847 | 1 | 14.04796 | 119907 | 576.2774 | tri/oligope | N |
| 522 | NPP | 583.3471 | 1 | 22.23929 | 199022.7 | 582.3398 | tri/oligope | N |
| 523 | NPP | 292.6922 | 2 | 2.195508 | 410639.2 | 583.3698 | tri/oligope | N |
| 524 | NPP | 585.3626 | 1 | 21.47653 | 183282.9 | 584.3554 | tri/oligope | N |
| 525 | NPP | 587.3051 | 1 | 18.30503 | 589360.8 | 586.2978 | tri/oligope | N |
| 526 | NPP | 588.2892 | 1 | 14.00581 | 362246.5 | 587.2819 | tri/oligope | N |
| 527 | NPP | 299.7007 | 2 | 19.12557 | 133885.7 | 597.3869 | tri/oligope | N |
| 528 | NPP | 300.1796 | 2 | 2.303717 | 392470.3 | 598.3446 | tri/oligope | N |
| 529 | NPP | 601.3209 | 1 | 18.4743 | 336503.1 | 600.3137 | tri/oligope | N |
| 530 | NPP | 306.6954 | 2 | 2.69675 | 388507.2 | 611.3763 | tri/oligope | N |
| 531 | NPP | 306.6959 | 2 | 11.55517 | 381356.8 | 611.3772 | tri/oligope | N |
| 532 | NPP | 308.6875 | 2 | 14.66273 | 105805.9 | 615.3605 | oligopepti | N |
| 533 | NPP | 623.2687 | 1 | 19.33034 | 107927.7 | 622.2614 | oligopepti | N |
| 534 | NPP | 312.1802 | 2 | 20.46632 | 188490.6 | 622.3459 | oligopepti | N |
| 535 | NPP | 314.6878 | 2 | 18.94098 | 104881.6 | 627.361 | oligopepti | N |
| 536 | NPP | 633.3259 | 1 | 19.30169 | 124143.7 | 632.3186 | oligopepti | N |
| 537 | NPP | 320.724 | 2 | 19.03958 | 460035.9 | 639.4335 | oligopepti | N |
| 538 | NPP | 649.3216 | 1 | 24.80442 | 115866.5 | 648.3144 | oligopepti | N |
| 539 | NPP | 655.368 | 1 | 19.03165 | 226703.3 | 654.3608 | oligopepti | N |
| 540 | NPP | 330.1621 | 2 | 14.18043 | 124655.5 | 658.3097 | oligopepti | N |
| 541 | NPP | 331.6524 | 2 | 2.149642 | 118343.1 | 661.2902 | oligopepti | N |
| 542 | NPP | 667.2745 | 1 | 23.93479 | 128607.2 | 666.2672 | oligopepti | N |
| 543 | NPP | 334.7273 | 2 | 25.84584 | 372780.4 | 667.4401 | oligopepti | N |
| 544 | NPP | 341.1642 | 2 | 2.228358 | 126116.4 | 680.3138 | oligopepti | N |
| 545 | NPP | 348.1911 | 2 | 15.26294 | 126270.3 | 694.3676 | oligopepti | N |
| 546 | NPP | 350.2245 | 2 | 19.41308 | 1439744 | 698.4344 | oligopepti | N |
| 547 | NPP | 358.1753 | 2 | 16.30568 | 197058.1 | 714.3361 | oligopepti | N |
| 548 | NPP | 742.4007 | 1 | 21.42936 | 141286.2 | 741.3935 | oligopepti | N |
| 549 | NPP | 382.2407 | 2 | 26.11503 | 104086.4 | 762.4669 | oligopepti | N |
| 550 | NPP | 389.2492 | 2 | 27.78298 | 159026.5 | 776.4839 | oligopepti | N |
| 551 | NPP | 434.7078 | 2 | 14.09174 | 704584.5 | 867.401 | oligopepti | N |
| 552 | NPP | 460.6665 | 2 | 14.30288 | 141859.7 | 919.3185 | oligopepti | N |
| 553 | NPP | 943.5273 | 1 | 34.96544 | 259495.4 | 942.5201 | oligopepti | N |
| 554 | NPP | 476.2074 | 2 | 17.35534 | 107899.9 | 950.4002 | oligopepti | N |

| ID | Protein | mz | charge | retentiont | raw.abund | mass | type | Match |
|---|---|---|---|---|---|---|---|---|
| 2 | BP | 231.171 | 1 | 5.378708 | 625083.8 | 230.1637 | di/tripepti | Y |
| 3 | BP | 231.1711 | 1 | 9.670667 | 1063120 | 230.1638 | di/tripepti | Y |
| 4 | BP | 231.1711 | 1 | 12.04521 | 1229779 | 230.1638 | di/tripepti | Y |
| 5 | BP | 233.15 | 1 | 2.201208 | 521386 | 232.1427 | di/tripepti | Y |
| 6 | BP | 233.1502 | 1 | 2.786958 | 209835.1 | 232.143 | di/tripepti | Y |
| 7 | BP | 239.103 | 1 | 2.190508 | 134390.2 | 238.0957 | di/tripepti | Y |
| 11 | BP | 245.1867 | 1 | 16.42313 | 1676322 | 244.1794 | di/tripepti | Y |
| 12 | BP | 245.1868 | 1 | 15.8879 | 245695.2 | 244.1795 | di/tripepti | Y |
| 13 | BP | 245.1868 | 1 | 14.51483 | 446650.9 | 244.1795 | di/tripepti | Y |
| 14 | BP | 245.1868 | 1 | 17.67486 | 582972.2 | 244.1795 | di/tripepti | Y |
| 15 | BP | 246.1453 | 1 | 2.201208 | 370495.7 | 245.138 | di/tripepti | Y |
| 16 | BP | 246.1455 | 1 | 2.631358 | 184281.4 | 245.1382 | di/tripepti | Y |
| 17 | BP | 246.1456 | 1 | 3.405142 | 485877.8 | 245.1384 | di/tripepti | Y |
| 20 | BP | 247.1293 | 1 | 2.126558 | 1128987 | 246.122 | di/tripepti | Y |
| 22 | BP | 247.1296 | 1 | 4.721192 | 589508.2 | 246.1223 | di/tripepti | Y |
| 23 | BP | 249.1275 | 1 | 2.725833 | 279589 | 248.1202 | di/tripepti | Y |
| 24 | BP | 250.1784 | 1 | 34.86318 | 770470.4 | 249.1711 | di/tripepti | Y |
| 25 | BP | 250.1784 | 1 | 37.2597 | 212118.8 | 249.1711 | di/tripepti | Y |
| 27 | BP | 253.1188 | 1 | 2.2919 | 984714.8 | 252.1115 | di/tripepti | Y |
| 30 | BP | 260.161 | 1 | 2.201208 | 503863 | 259.1538 | di/tripepti | Y |
| 31 | BP | 260.1611 | 1 | 2.551008 | 257285.7 | 259.1539 | di/tripepti | Y |
| 33 | BP | 260.1974 | 1 | 1.844667 | 139829.3 | 259.1901 | di/tripepti | Y |
| 34 | BP | 261.145 | 1 | 2.190508 | 476441.1 | 260.1378 | di/tripepti | Y |
| 36 | BP | 261.1453 | 1 | 5.476758 | 393311.7 | 260.1381 | di/tripepti | Y |
| 38 | BP | 263.1433 | 1 | 11.5175 | 259251.8 | 262.136 | di/tripepti | Y |
| 40 | BP | 265.1556 | 1 | 16.25883 | 148965.3 | 264.1483 | di/tripepti | Y |
| 41 | BP | 267.1345 | 1 | 2.401483 | 824444.5 | 266.1273 | di/tripepti | Y |
| 42 | BP | 269.1613 | 1 | 2.190508 | 342425.3 | 268.1541 | di/tripepti | Y |
| 45 | BP | 281.1139 | 1 | 2.558442 | 250075.5 | 280.1066 | di/tripepti | Y |
| 47 | BP | 288.1928 | 1 | 15.93067 | 102285.5 | 287.1855 | di/tripepti | Y |
| 50 | BP | 294.1455 | 1 | 2.317767 | 236668.6 | 293.1382 | di/tripepti | Y |
| 51 | BP | 295.13 | 1 | 3.348983 | 145668.1 | 294.1227 | di/tripepti | Y |
| 52 | BP | 295.1661 | 1 | 16.40204 | 480260.4 | 294.1588 | di/tripepti | Y |
| 57 | BP | 302.2086 | 1 | 19.86203 | 215639.2 | 301.2013 | tri/oligope | Y |
| 59 | BP | 304.151 | 1 | 2.222792 | 154050 | 303.1437 | tri/oligope | Y |
| 60 | BP | 304.1513 | 1 | 4.90775 | 899493.5 | 303.144 | tri/oligope | Y |
| 61 | BP | 304.1667 | 1 | 18.98487 | 326614.3 | 303.1595 | tri/oligope | Y |
| 64 | BP | 318.1668 | 1 | 2.244567 | 690080.8 | 317.1595 | tri/oligope | Y |
| 65 | BP | 318.1671 | 1 | 9.108958 | 829494.7 | 317.1598 | tri/oligope | Y |
| 67 | BP | 318.1827 | 1 | 24.7141 | 123280.9 | 317.1754 | tri/oligope | Y |
| 69 | BP | 319.1619 | 1 | 1.959117 | 158112.7 | 318.1546 | tri/oligope | Y |
| 73 | BP | 331.1663 | 1 | 2.770708 | 262811.4 | 330.159 | tri/oligope | Y |
| 75 | BP | 332.1825 | 1 | 2.576742 | 482236.1 | 331.1752 | tri/oligope | Y |
| 76 | BP | 332.2192 | 1 | 2.818683 | 117212.3 | 331.2119 | tri/oligope | Y |
| 79 | BP | 333.157 | 1 | 11.04885 | 494257 | 332.1497 | tri/oligope | Y |
| 80 | BP | 334.1615 | 1 | 2.201208 | 412958.1 | 333.1543 | tri/oligope | Y |
| 92 | BP | 360.2142 | 1 | 2.643 | 181142.9 | 359.2069 | tri/oligope | Y |
| 94 | BP | 363.1559 | 1 | 2.190508 | 127436.2 | 362.1486 | tri/oligope | Y |
| 112 | BP | 415.2123 | 1 | 36.47223 | 217662.5 | 414.2051 | tri/oligope | Y |

| | | | | | | | | |
|---|---|---|---|---|---|---|---|---|
| 114 | BP | 420.2096 | 1 | 2.2919 | 242467.8 | 419.2024 | tri/oligope | Y |
| 141 | BP | 479.3109 | 1 | 36.99012 | 150898.5 | 478.3036 | tri/oligope | Y |
| 144 | BP | 488.2522 | 1 | 19.13919 | 190180.2 | 487.245 | tri/oligope | Y |
| 148 | BP | 491.2879 | 1 | 23.13454 | 953820 | 490.2807 | tri/oligope | Y |
| 151 | BP | 500.2889 | 1 | 25.19403 | 140298.3 | 499.2816 | tri/oligope | Y |
| 153 | BP | 513.2685 | 1 | 13.73743 | 138772.4 | 512.2612 | tri/oligope | Y |
| 159 | BP | 520.2785 | 1 | 21.16339 | 139516.9 | 519.2712 | tri/oligope | Y |
| 161 | BP | 264.1752 | 2 | 35.28835 | 106889 | 526.3358 | tri/oligope | Y |
| 165 | BP | 531.0146 | 1 | 1.596683 | 282829 | 530.0074 | tri/oligope | Y |
| 174 | BP | 274.1681 | 2 | 35.94048 | 110173.1 | 546.3217 | tri/oligope | Y |
| 187 | BP | 572.2772 | 1 | 19.16912 | 734901 | 571.2699 | tri/oligope | Y |
| 194 | BP | 599.0026 | 1 | 1.596683 | 248062.2 | 597.9953 | tri/oligope | Y |
| 218 | BP | 666.9901 | 1 | 1.608117 | 259936.9 | 665.9828 | oligopepti | Y |
| 231 | BP | 702.37 | 1 | 18.61288 | 173453 | 701.3627 | oligopepti | Y |
| 240 | BP | 734.9776 | 1 | 1.596683 | 218189.9 | 733.9704 | oligopepti | Y |
| 258 | BP | 802.9652 | 1 | 1.585458 | 193038.1 | 801.958 | oligopepti | Y |
| 264 | BP | 276.1789 | 3 | 15.52703 | 1343296 | 825.5148 | oligopepti | Y |
| 276 | BP | 870.9526 | 1 | 1.585458 | 147370.5 | 869.9453 | oligopepti | Y |
| 297 | BP | 990.79 | 1 | 1.596683 | 151081.8 | 989.7827 | oligopepti | Y |
| 300 | BP | 514.9584 | 2 | 1.596683 | 112449.6 | 1027.902 | oligopepti | Y |
| 303 | BP | 523.7873 | 2 | 23.1578 | 294597.8 | 1045.56 | oligopepti | Y |
| 307 | BP | 548.9522 | 2 | 1.608117 | 124133.1 | 1095.89 | oligopepti | Y |
| 313 | BP | 582.9463 | 2 | 1.596683 | 122831.9 | 1163.878 | oligopepti | Y |
| 314 | BP | 590.9332 | 2 | 1.608117 | 100215.8 | 1179.852 | oligopepti | Y |
| 316 | BP | 616.9403 | 2 | 1.608117 | 123697.1 | 1231.866 | oligopepti | Y |
| 318 | BP | 624.9269 | 2 | 1.596683 | 111869 | 1247.839 | oligopepti | Y |
| 321 | BP | 650.9337 | 2 | 1.608117 | 128075.4 | 1299.853 | oligopepti | Y |
| 322 | BP | 658.9207 | 2 | 1.608117 | 108972 | 1315.827 | oligopepti | Y |
| 329 | BP | 692.9147 | 2 | 1.596683 | 110335.3 | 1383.815 | oligopepti | Y |
| 333 | BP | 760.902 | 2 | 1.596683 | 101012.9 | 1519.789 | oligopepti | Y |
| 335 | BP | 794.896 | 2 | 1.608117 | 101060.5 | 1587.777 | oligopepti | Y |
| 338 | BP | 223.1084 | 1 | 10.38173 | 165635.6 | 222.1011 | dipeptide | N |
| 339 | BP | 226.9525 | 1 | 1.677442 | 166223.7 | 225.9452 | di/tripepti | N |
| 340 | BP | 229.1191 | 1 | 10.25577 | 151794.7 | 228.1118 | di/tripepti | N |
| 341 | BP | 229.1554 | 1 | 8.98885 | 227353.2 | 228.1481 | di/tripepti | N |
| 342 | BP | 233.1136 | 1 | 1.994208 | 144599.5 | 232.1063 | di/tripepti | N |
| 343 | BP | 233.1504 | 1 | 4.1048 | 496718.7 | 232.1431 | di/tripepti | N |
| 344 | BP | 237.0908 | 1 | 2.190508 | 120608.2 | 236.0835 | di/tripepti | N |
| 345 | BP | 237.1241 | 1 | 4.611867 | 110915.9 | 236.1168 | di/tripepti | N |
| 346 | BP | 237.1241 | 1 | 11.19204 | 327781.6 | 236.1168 | di/tripepti | N |
| 347 | BP | 244.1296 | 1 | 2.1048 | 147406.2 | 243.1223 | di/tripepti | N |
| 348 | BP | 252.9631 | 1 | 9.934192 | 152778.4 | 251.9558 | di/tripepti | N |
| 349 | BP | 255.1456 | 1 | 1.783425 | 205768.8 | 254.1383 | di/tripepti | N |
| 350 | BP | 262.1195 | 1 | 14.4752 | 580140 | 261.1122 | di/tripepti | N |
| 351 | BP | 263.1433 | 1 | 13.09528 | 173396.8 | 262.136 | di/tripepti | N |
| 352 | BP | 265.1556 | 1 | 14.74059 | 265548.2 | 264.1483 | di/tripepti | N |
| 353 | BP | 269.1138 | 1 | 2.190508 | 513803.3 | 268.1065 | di/tripepti | N |
| 354 | BP | 276.1353 | 1 | 14.32788 | 116733.3 | 275.128 | di/tripepti | N |
| 355 | BP | 276.1559 | 1 | 2.0604 | 372575.9 | 275.1487 | di/tripepti | N |
| 356 | BP | 276.156 | 1 | 2.190508 | 699997.4 | 275.1487 | di/tripepti | N |

| | | | | | | | | |
|---|---|---|---|---|---|---|---|---|
| 357 | BP | 276.156 | 1 | 2.389858 | 785000.6 | 275.1487 | di/tripepti | N |
| 358 | BP | 279.1714 | 1 | 19.89973 | 394200.4 | 278.1641 | di/tripepti | N |
| 359 | BP | 279.1715 | 1 | 21.03043 | 219766.6 | 278.1642 | di/tripepti | N |
| 360 | BP | 279.2324 | 1 | 37.0111 | 165050.5 | 278.2252 | di/tripepti | N |
| 361 | BP | 281.1142 | 1 | 11.55735 | 160981.5 | 280.1069 | di/tripepti | N |
| 362 | BP | 281.1505 | 1 | 5.510233 | 138349.7 | 280.1432 | di/tripepti | N |
| 363 | BP | 281.1506 | 1 | 6.053717 | 300770.9 | 280.1433 | di/tripepti | N |
| 364 | BP | 283.1293 | 1 | 2.201208 | 145615.9 | 282.1221 | di/tripepti | N |
| 365 | BP | 288.1925 | 1 | 2.666167 | 426513.1 | 287.1853 | di/tripepti | N |
| 366 | BP | 289.1513 | 1 | 2.190508 | 321953.2 | 288.144 | di/tripepti | N |
| 367 | BP | 290.1716 | 1 | 2.201208 | 385068.5 | 289.1644 | di/tripepti | N |
| 368 | BP | 292.1301 | 1 | 16.41637 | 466011.2 | 291.1228 | di/tripepti | N |
| 369 | BP | 292.1301 | 1 | 9.455833 | 366045.7 | 291.1228 | di/tripepti | N |
| 370 | BP | 292.1508 | 1 | 2.038392 | 214025.1 | 291.1435 | di/tripepti | N |
| 371 | BP | 295.13 | 1 | 12.62893 | 167975.7 | 294.1227 | di/tripepti | N |
| 372 | BP | 295.1663 | 1 | 14.5967 | 143747.1 | 294.1591 | di/tripepti | N |
| 373 | BP | 295.1664 | 1 | 13.3899 | 340748.7 | 294.1591 | di/tripepti | N |
| 374 | BP | 295.2275 | 1 | 37.38036 | 110241.7 | 294.2202 | di/tripepti | N |
| 375 | BP | 302.1717 | 1 | 2.222792 | 127980.7 | 301.1644 | tri/oligope | N |
| 376 | BP | 302.2086 | 1 | 11.18408 | 118989.6 | 301.2013 | tri/oligope | N |
| 377 | BP | 304.1668 | 1 | 19.9139 | 173946.5 | 303.1595 | tri/oligope | N |
| 378 | BP | 304.1874 | 1 | 2.190508 | 590437.2 | 303.1801 | tri/oligope | N |
| 379 | BP | 304.1879 | 1 | 11.34206 | 130644.5 | 303.1806 | tri/oligope | N |
| 380 | BP | 306.1302 | 1 | 1.856525 | 101759.4 | 305.1229 | tri/oligope | N |
| 381 | BP | 306.1668 | 1 | 2.190508 | 780396.8 | 305.1595 | tri/oligope | N |
| 382 | BP | 311.1244 | 1 | 2.190508 | 478811 | 310.1171 | tri/oligope | N |
| 383 | BP | 316.1875 | 1 | 2.355467 | 810914.2 | 315.1802 | tri/oligope | N |
| 384 | BP | 316.188 | 1 | 12.97118 | 243580.9 | 315.1807 | tri/oligope | N |
| 385 | BP | 316.2244 | 1 | 15.40928 | 119463.5 | 315.2171 | tri/oligope | N |
| 386 | BP | 316.2244 | 1 | 14.80839 | 111623.3 | 315.2171 | tri/oligope | N |
| 387 | BP | 318.2031 | 1 | 2.317767 | 117046.6 | 317.1958 | tri/oligope | N |
| 388 | BP | 318.2037 | 1 | 12.38566 | 162850.9 | 317.1964 | tri/oligope | N |
| 389 | BP | 319.1413 | 1 | 16.81207 | 100676 | 318.134 | tri/oligope | N |
| 390 | BP | 320.1253 | 1 | 11.79946 | 119970.6 | 319.1181 | tri/oligope | N |
| 391 | BP | 320.1254 | 1 | 17.65223 | 166494.3 | 319.1181 | tri/oligope | N |
| 392 | BP | 320.1459 | 1 | 1.925067 | 153394.6 | 319.1387 | tri/oligope | N |
| 393 | BP | 324.1298 | 1 | 2.190508 | 569218 | 323.1225 | tri/oligope | N |
| 394 | BP | 328.2245 | 1 | 17.09425 | 500267.3 | 327.2172 | tri/oligope | N |
| 395 | BP | 328.2343 | 1 | 15.48075 | 138814.4 | 327.227 | tri/oligope | N |
| 396 | BP | 331.1984 | 1 | 2.201208 | 149211 | 330.1911 | tri/oligope | N |
| 397 | BP | 332.146 | 1 | 2.038392 | 151662.4 | 331.1387 | tri/oligope | N |
| 398 | BP | 332.2192 | 1 | 4.856167 | 582582.7 | 331.2119 | tri/oligope | N |
| 399 | BP | 332.2193 | 1 | 8.98005 | 532498.8 | 331.212 | tri/oligope | N |
| 400 | BP | 332.2193 | 1 | 15.89443 | 103267.4 | 331.212 | tri/oligope | N |
| 401 | BP | 332.2195 | 1 | 19.44302 | 129266.8 | 331.2122 | tri/oligope | N |
| 402 | BP | 333.1776 | 1 | 2.201208 | 237375.1 | 332.1704 | tri/oligope | N |
| 403 | BP | 334.1252 | 1 | 1.936058 | 151593.4 | 333.1179 | tri/oligope | N |
| 404 | BP | 334.1409 | 1 | 13.73743 | 119216.2 | 333.1336 | tri/oligope | N |
| 405 | BP | 334.1412 | 1 | 17.95638 | 174562.5 | 333.1339 | tri/oligope | N |
| 406 | BP | 334.1771 | 1 | 16.38625 | 143868.8 | 333.1698 | tri/oligope | N |

| | | | | | | | | |
|---|---|---|---|---|---|---|---|---|
| 407 | BP | 334.1775 | 1 | 18.02637 | 101370.8 | 333.1702 | tri/oligope | N |
| 408 | BP | 334.1979 | 1 | 2.201208 | 153225.3 | 333.1906 | tri/oligope | N |
| 409 | BP | 345.2146 | 1 | 12.60298 | 304639.9 | 344.2074 | tri/oligope | N |
| 410 | BP | 346.1986 | 1 | 12.74334 | 641039.9 | 345.1914 | tri/oligope | N |
| 411 | BP | 346.2353 | 1 | 19.18507 | 103314.3 | 345.228 | tri/oligope | N |
| 412 | BP | 347.1569 | 1 | 2.038392 | 119756.8 | 346.1496 | tri/oligope | N |
| 413 | BP | 348.1774 | 1 | 2.201208 | 370341.1 | 347.1701 | tri/oligope | N |
| 414 | BP | 350.1752 | 1 | 10.06618 | 124806.7 | 349.1679 | tri/oligope | N |
| 415 | BP | 352.1517 | 1 | 11.26378 | 956510.1 | 351.1444 | tri/oligope | N |
| 416 | BP | 357.2141 | 1 | 2.190508 | 425379.9 | 356.2069 | tri/oligope | N |
| 417 | BP | 360.2144 | 1 | 8.789833 | 503433.4 | 359.2071 | tri/oligope | N |
| 418 | BP | 361.1726 | 1 | 2.201208 | 391053.9 | 360.1653 | tri/oligope | N |
| 419 | BP | 362.1931 | 1 | 2.244567 | 1206515 | 361.1858 | tri/oligope | N |
| 420 | BP | 363.1518 | 1 | 1.891675 | 136193.7 | 362.1445 | tri/oligope | N |
| 421 | BP | 372.1887 | 1 | 2.169517 | 199660.8 | 371.1814 | tri/oligope | N |
| 422 | BP | 373.1729 | 1 | 2.071533 | 287741.2 | 372.1657 | tri/oligope | N |
| 423 | BP | 374.109 | 1 | 2.190508 | 447175.6 | 373.1017 | tri/oligope | N |
| 424 | BP | 375.2253 | 1 | 12.79243 | 139692.6 | 374.218 | tri/oligope | N |
| 425 | BP | 378.2036 | 1 | 15.08798 | 564407.1 | 377.1964 | tri/oligope | N |
| 426 | BP | 382.1619 | 1 | 2.522775 | 347352.5 | 381.1547 | tri/oligope | N |
| 427 | BP | 389.2039 | 1 | 2.201208 | 105874.6 | 388.1966 | tri/oligope | N |
| 428 | BP | 393.1988 | 1 | 2.190508 | 131623.9 | 392.1915 | tri/oligope | N |
| 429 | BP | 399.2617 | 1 | 17.65223 | 233847.6 | 398.2544 | tri/oligope | N |
| 430 | BP | 405.2352 | 1 | 2.212083 | 375180.7 | 404.228 | tri/oligope | N |
| 431 | BP | 409.0201 | 1 | 2.201208 | 207427.5 | 408.0129 | tri/oligope | N |
| 432 | BP | 409.1886 | 1 | 23.70576 | 439877.6 | 408.1814 | tri/oligope | N |
| 433 | BP | 413.2673 | 1 | 36.71189 | 936003.1 | 412.26 | tri/oligope | N |
| 434 | BP | 419.2152 | 1 | 2.978092 | 643183.1 | 418.2079 | tri/oligope | N |
| 435 | BP | 419.2513 | 1 | 14.38211 | 639094.7 | 418.244 | tri/oligope | N |
| 436 | BP | 424.2093 | 1 | 14.50823 | 158932.6 | 423.202 | tri/oligope | N |
| 437 | BP | 429.2726 | 1 | 20.71539 | 142548.4 | 428.2653 | tri/oligope | N |
| 438 | BP | 430.2676 | 1 | 14.89067 | 107990.1 | 429.2603 | tri/oligope | N |
| 439 | BP | 431.2517 | 1 | 14.26715 | 138983.2 | 430.2444 | tri/oligope | N |
| 440 | BP | 432.2096 | 1 | 2.027433 | 154209.9 | 431.2024 | tri/oligope | N |
| 441 | BP | 433.2675 | 1 | 19.19297 | 103897.9 | 432.2602 | tri/oligope | N |
| 442 | BP | 435.2256 | 1 | 17.70002 | 519512.3 | 434.2183 | tri/oligope | N |
| 443 | BP | 436.3438 | 1 | 1.366417 | 284646.2 | 435.3365 | tri/oligope | N |
| 444 | BP | 439.2935 | 1 | 23.49402 | 121088.3 | 438.2862 | tri/oligope | N |
| 445 | BP | 440.252 | 1 | 15.60512 | 119187.1 | 439.2448 | tri/oligope | N |
| 446 | BP | 447.2003 | 1 | 13.90281 | 269554.4 | 446.1931 | tri/oligope | N |
| 447 | BP | 447.2256 | 1 | 16.43152 | 174362.3 | 446.2183 | tri/oligope | N |
| 448 | BP | 447.2461 | 1 | 15.45893 | 956666 | 446.2388 | tri/oligope | N |
| 449 | BP | 448.1841 | 1 | 18.69215 | 500190.7 | 447.1769 | tri/oligope | N |
| 450 | BP | 455.3248 | 1 | 25.64467 | 182157.9 | 454.3176 | tri/oligope | N |
| 451 | BP | 230.1322 | 2 | 1.959117 | 102921.1 | 458.2498 | tri/oligope | N |
| 452 | BP | 462.2202 | 1 | 2.179992 | 135761.7 | 461.213 | tri/oligope | N |
| 453 | BP | 471.2933 | 1 | 2.190508 | 140971.5 | 470.286 | tri/oligope | N |
| 454 | BP | 236.1504 | 2 | 2.201208 | 203556.6 | 470.2862 | tri/oligope | N |
| 455 | BP | 472.2782 | 1 | 15.58155 | 288057.5 | 471.2709 | tri/oligope | N |
| 456 | BP | 474.2403 | 1 | 19.4263 | 162394 | 473.233 | tri/oligope | N |

| | | | | | | | | |
|---|---|---|---|---|---|---|---|---|
| 457 | BP | 479.2151 | 1 | 2.676858 | 475729.1 | 478.2078 | tri/oligope | N |
| 458 | BP | 486.2569 | 1 | 13.51828 | 281074.5 | 485.2497 | tri/oligope | N |
| 459 | BP | 486.2938 | 1 | 16.25883 | 407640.2 | 485.2865 | tri/oligope | N |
| 460 | BP | 487.2892 | 1 | 12.74992 | 201596 | 486.2819 | tri/oligope | N |
| 461 | BP | 488.2734 | 1 | 13.92928 | 211040.2 | 487.2661 | tri/oligope | N |
| 462 | BP | 249.6745 | 2 | 16.84591 | 190639.2 | 497.3345 | tri/oligope | N |
| 463 | BP | 504.2668 | 1 | 2.190508 | 167640.1 | 503.2595 | tri/oligope | N |
| 464 | BP | 253.6528 | 2 | 2.190508 | 187111.1 | 505.2911 | tri/oligope | N |
| 465 | BP | 255.1642 | 2 | 1.389167 | 256353.9 | 508.3139 | tri/oligope | N |
| 466 | BP | 522.2577 | 1 | 17.08049 | 108857.1 | 521.2504 | tri/oligope | N |
| 467 | BP | 261.6508 | 2 | 17.34193 | 443166.2 | 521.2871 | tri/oligope | N |
| 468 | BP | 530.2996 | 1 | 26.96137 | 137845.2 | 529.2924 | tri/oligope | N |
| 469 | BP | 531.2617 | 1 | 18.36313 | 155544.9 | 530.2544 | tri/oligope | N |
| 470 | BP | 543.2779 | 1 | 2.190508 | 157185 | 542.2706 | tri/oligope | N |
| 471 | BP | 544.2641 | 1 | 15.62132 | 428683.4 | 543.2568 | tri/oligope | N |
| 472 | BP | 545.2578 | 1 | 2.607225 | 122451.4 | 544.2505 | tri/oligope | N |
| 473 | BP | 553.3365 | 1 | 24.37516 | 2044654 | 552.3292 | tri/oligope | N |
| 474 | BP | 279.1877 | 2 | 18.42107 | 147591.5 | 556.3609 | tri/oligope | N |
| 475 | BP | 285.6853 | 2 | 13.73743 | 148085.4 | 569.356 | tri/oligope | N |
| 476 | BP | 286.6461 | 2 | 15.20957 | 102092.4 | 571.2777 | tri/oligope | N |
| 477 | BP | 573.2537 | 1 | 12.05268 | 246675.7 | 572.2464 | tri/oligope | N |
| 478 | BP | 587.3057 | 1 | 12.07098 | 102389.9 | 586.2984 | tri/oligope | N |
| 479 | BP | 592.3006 | 1 | 21.88585 | 300123.6 | 591.2933 | tri/oligope | N |
| 480 | BP | 300.0945 | 2 | 1.959117 | 175317.6 | 598.1745 | tri/oligope | N |
| 481 | BP | 600.3359 | 1 | 2.179992 | 178292.6 | 599.3287 | tri/oligope | N |
| 482 | BP | 300.6718 | 2 | 2.190508 | 424340.8 | 599.329 | tri/oligope | N |
| 483 | BP | 302.6694 | 2 | 2.049308 | 259618.9 | 603.3242 | tri/oligope | N |
| 484 | BP | 304.177 | 2 | 2.389858 | 667040.6 | 606.3394 | tri/oligope | N |
| 485 | BP | 623.3054 | 1 | 16.43152 | 413242.9 | 622.2981 | oligopepti | N |
| 486 | BP | 316.6543 | 2 | 2.027433 | 163041 | 631.2941 | oligopepti | N |
| 487 | BP | 328.1257 | 2 | 2.622217 | 136347.5 | 654.2369 | oligopepti | N |
| 488 | BP | 330.1314 | 2 | 17.51987 | 145801.2 | 658.2483 | oligopepti | N |
| 489 | BP | 342.1902 | 2 | 2.212083 | 279651.1 | 682.3658 | oligopepti | N |
| 490 | BP | 349.7165 | 2 | 19.68709 | 129234.2 | 697.4185 | oligopepti | N |
| 491 | BP | 716.3856 | 1 | 18.95381 | 407255.8 | 715.3783 | oligopepti | N |
| 492 | BP | 387.1884 | 2 | 2.179992 | 167695.7 | 772.3623 | oligopepti | N |
| 493 | BP | 395.1865 | 2 | 14.72253 | 103120.1 | 788.3585 | oligopepti | N |
| 494 | BP | 798.3708 | 1 | 24.10095 | 106066.9 | 797.3635 | oligopepti | N |
| 495 | BP | 412.2077 | 2 | 2.169517 | 111907.7 | 822.4009 | oligopepti | N |
| 496 | BP | 413.7646 | 2 | 16.43152 | 308034.3 | 825.5146 | oligopepti | N |
| 497 | BP | 419.2104 | 2 | 14.89067 | 389338.6 | 836.4063 | oligopepti | N |
| 498 | BP | 460.6672 | 2 | 14.26049 | 109760.8 | 919.3198 | oligopepti | N |
| 499 | BP | 463.747 | 2 | 16.09563 | 200564.7 | 925.4794 | oligopepti | N |
| 500 | BP | 478.7759 | 2 | 16.53578 | 265114.6 | 955.5373 | oligopepti | N |
| 501 | BP | 325.4923 | 3 | 15.3074 | 115392.1 | 973.455 | oligopepti | N |
| 502 | BP | 338.8432 | 3 | 13.47607 | 331977.2 | 1013.508 | oligopepti | N |
| 503 | BP | 510.2557 | 2 | 16.24487 | 137621.1 | 1018.497 | oligopepti | N |
| 504 | BP | 520.2896 | 2 | 21.83461 | 184232.1 | 1038.565 | oligopepti | N |
| 505 | BP | 542.7745 | 2 | 15.65519 | 106576.6 | 1083.534 | oligopepti | N |
| 506 | BP | 684.9277 | 2 | 1.608117 | 120038.9 | 1367.841 | oligopepti | N |

| | | | | | | | | |
|---|---|---|---|---|---|---|---|---|
| 507 | BP | 718.9215 | 2 | 1.608117 | 119749.2 | 1435.828 | oligopepti | N |
| 508 | BP | 752.9154 | 2 | 1.608117 | 102845 | 1503.816 | oligopepti | N |
| 509 | BP | 786.9083 | 2 | 1.608117 | 102210.9 | 1571.802 | oligopepti | N |
| 510 | BP | 820.903 | 2 | 1.596683 | 101779 | 1639.791 | oligopepti | N |
| 511 | BP | 922.802 | 2 | 1.608117 | 216151.5 | 1843.589 | oligopepti | N |
| 512 | BP | 1058.777 | 2 | 1.608117 | 110486.7 | 2115.54 | oligopepti | N |

| ID | Protein | mz | charge | retentionti | raw.abund | mass | type | Match |
|---|---|---|---|---|---|---|---|---|
| 15 | Peptan | 246.1453 | 1 | 2.173233 | 192720.5 | 245.138 | di/tripepti | Y |
| 22 | Peptan | 247.1296 | 1 | 4.710933 | 145785 | 246.1223 | di/tripepti | Y |
| 24 | Peptan | 250.1784 | 1 | 34.8638 | 719351.8 | 249.1711 | di/tripepti | Y |
| 25 | Peptan | 250.1784 | 1 | 37.26694 | 206849.7 | 249.1711 | di/tripepti | Y |
| 30 | Peptan | 260.161 | 1 | 2.173233 | 276150.4 | 259.1538 | di/tripepti | Y |
| 31 | Peptan | 260.1611 | 1 | 2.509258 | 348136.1 | 259.1539 | di/tripepti | Y |
| 59 | Peptan | 304.151 | 1 | 2.19435 | 360622.1 | 303.1437 | tri/oligope | Y |
| 64 | Peptan | 318.1668 | 1 | 2.2156 | 107753 | 317.1595 | tri/oligope | Y |
| 72 | Peptan | 331.1619 | 1 | 2.119542 | 147552.8 | 330.1546 | tri/oligope | Y |
| 112 | Peptan | 415.2123 | 1 | 36.47483 | 250209.1 | 414.2051 | tri/oligope | Y |
| 133 | Peptan | 461.301 | 1 | 37.42613 | 101667.5 | 460.2937 | tri/oligope | Y |
| 141 | Peptan | 479.3109 | 1 | 36.99213 | 156959.8 | 478.3036 | tri/oligope | Y |
| 156 | Peptan | 515.292 | 1 | 37.11854 | 103130.9 | 514.2847 | tri/oligope | Y |
| 161 | Peptan | 264.1752 | 2 | 35.28911 | 107658.7 | 526.3358 | tri/oligope | Y |
| 165 | Peptan | 531.0146 | 1 | 1.581025 | 183538.5 | 530.0074 | tri/oligope | Y |
| 174 | Peptan | 274.1681 | 2 | 35.93957 | 110090 | 546.3217 | tri/oligope | Y |
| 190 | Peptan | 584.3056 | 1 | 2.2263 | 359131 | 583.2983 | tri/oligope | Y |
| 194 | Peptan | 599.0026 | 1 | 1.581025 | 162461.9 | 597.9953 | tri/oligope | Y |
| 218 | Peptan | 666.9901 | 1 | 1.593725 | 162240 | 665.9828 | oligopepti | Y |
| 240 | Peptan | 734.9776 | 1 | 1.581025 | 142238.8 | 733.9704 | oligopepti | Y |
| 258 | Peptan | 802.9652 | 1 | 1.569142 | 121084.7 | 801.958 | oligopepti | Y |
| 297 | Peptan | 990.79 | 1 | 1.581025 | 208202.4 | 989.7827 | oligopepti | Y |
| 338 | Peptan | 223.1084 | 1 | 10.46356 | 431339.6 | 222.1011 | dipeptide | N |
| 339 | Peptan | 226.9522 | 1 | 1.520492 | 147888.7 | 225.945 | di/tripepti | N |
| 340 | Peptan | 226.9525 | 1 | 1.667042 | 427105.9 | 225.9452 | di/tripepti | N |
| 341 | Peptan | 242.9266 | 1 | 1.667042 | 130247.8 | 241.9193 | di/tripepti | N |
| 342 | Peptan | 244.1296 | 1 | 2.075283 | 730378.6 | 243.1223 | di/tripepti | N |
| 343 | Peptan | 252.9631 | 1 | 10.01325 | 131336.5 | 251.9558 | di/tripepti | N |
| 344 | Peptan | 262.1039 | 1 | 1.877517 | 172495 | 261.0966 | di/tripepti | N |
| 345 | Peptan | 272.161 | 1 | 2.282258 | 522589.9 | 271.1538 | di/tripepti | N |
| 346 | Peptan | 276.1195 | 1 | 1.877517 | 251645.4 | 275.1122 | di/tripepti | N |
| 347 | Peptan | 286.1771 | 1 | 10.93833 | 370508 | 285.1698 | di/tripepti | N |
| 348 | Peptan | 290.1352 | 1 | 2.19435 | 245617.3 | 289.128 | di/tripepti | N |
| 349 | Peptan | 294.9405 | 1 | 1.667042 | 174204.7 | 293.9332 | di/tripepti | N |
| 350 | Peptan | 301.1512 | 1 | 2.097825 | 269156.6 | 300.144 | di/tripepti | N |
| 351 | Peptan | 302.1717 | 1 | 2.19435 | 139941.5 | 301.1644 | tri/oligope | N |
| 352 | Peptan | 304.1513 | 1 | 8.269217 | 149441.1 | 303.144 | tri/oligope | N |
| 353 | Peptan | 315.1669 | 1 | 2.19435 | 572662.5 | 314.1596 | tri/oligope | N |
| 354 | Peptan | 317.1462 | 1 | 1.855583 | 118774.5 | 316.139 | tri/oligope | N |
| 355 | Peptan | 317.1827 | 1 | 2.19435 | 127270.9 | 316.1754 | tri/oligope | N |
| 356 | Peptan | 318.1302 | 1 | 1.918883 | 129756.1 | 317.123 | tri/oligope | N |
| 357 | Peptan | 328.2343 | 1 | 15.48488 | 112942.9 | 327.227 | tri/oligope | N |
| 358 | Peptan | 329.1941 | 1 | 1.7503 | 153476.7 | 328.1868 | tri/oligope | N |
| 359 | Peptan | 331.1986 | 1 | 2.663867 | 158001.9 | 330.1913 | tri/oligope | N |
| 360 | Peptan | 343.1987 | 1 | 2.259725 | 742589.9 | 342.1914 | tri/oligope | N |
| 361 | Peptan | 343.1989 | 1 | 13.68614 | 109833.7 | 342.1917 | tri/oligope | N |
| 362 | Peptan | 345.1777 | 1 | 2.19435 | 383970.5 | 344.1704 | tri/oligope | N |
| 363 | Peptan | 357.2146 | 1 | 12.53406 | 250950.2 | 356.2073 | tri/oligope | N |
| 364 | Peptan | 359.157 | 1 | 2.19435 | 141461.4 | 358.1497 | tri/oligope | N |

| | | | | | | | | |
|---|---|---|---|---|---|---|---|---|
| 365 | Peptan | 359.1933 | 1 | 2.534867 | 1979714 | 358.1861 | tri/oligope | N |
| 366 | Peptan | 374.2044 | 1 | 2.582608 | 130314.9 | 373.1971 | tri/oligope | N |
| 367 | Peptan | 375.1518 | 1 | 1.907467 | 150953.7 | 374.1445 | tri/oligope | N |
| 368 | Peptan | 385.2461 | 1 | 17.78398 | 211270.1 | 384.2388 | tri/oligope | N |
| 369 | Peptan | 388.1838 | 1 | 2.183825 | 355345.2 | 387.1765 | tri/oligope | N |
| 370 | Peptan | 393.1783 | 1 | 12.84225 | 187525.3 | 392.171 | tri/oligope | N |
| 371 | Peptan | 393.181 | 1 | 2.19435 | 621497.5 | 392.1737 | tri/oligope | N |
| 372 | Peptan | 400.2202 | 1 | 2.4053 | 158853.8 | 399.2129 | tri/oligope | N |
| 373 | Peptan | 401.2044 | 1 | 8.483533 | 114881.4 | 400.1971 | tri/oligope | N |
| 374 | Peptan | 405.2617 | 1 | 37.28453 | 142718.3 | 404.2544 | tri/oligope | N |
| 375 | Peptan | 413.2673 | 1 | 36.71656 | 978885.2 | 412.26 | tri/oligope | N |
| 376 | Peptan | 414.2357 | 1 | 2.552767 | 646775.2 | 413.2284 | tri/oligope | N |
| 377 | Peptan | 420.1733 | 1 | 2.097825 | 373386.6 | 419.166 | tri/oligope | N |
| 378 | Peptan | 430.194 | 1 | 2.19435 | 756565.4 | 429.1867 | tri/oligope | N |
| 379 | Peptan | 430.231 | 1 | 2.978375 | 419766.5 | 429.2237 | tri/oligope | N |
| 380 | Peptan | 430.2311 | 1 | 13.83522 | 887748.8 | 429.2238 | tri/oligope | N |
| 381 | Peptan | 432.1732 | 1 | 2.009375 | 457393.8 | 431.1659 | tri/oligope | N |
| 382 | Peptan | 436.3438 | 1 | 1.350842 | 256414.5 | 435.3365 | tri/oligope | N |
| 383 | Peptan | 443.2256 | 1 | 2.19435 | 273234.1 | 442.2183 | tri/oligope | N |
| 384 | Peptan | 446.1888 | 1 | 2.020658 | 133835.1 | 445.1816 | tri/oligope | N |
| 385 | Peptan | 446.189 | 1 | 2.19435 | 277523.6 | 445.1817 | tri/oligope | N |
| 386 | Peptan | 446.2258 | 1 | 2.789092 | 776291.4 | 445.2186 | tri/oligope | N |
| 387 | Peptan | 449.2878 | 1 | 37.23838 | 106847.8 | 448.2806 | tri/oligope | N |
| 388 | Peptan | 450.1997 | 1 | 13.63806 | 355891 | 449.1924 | tri/oligope | N |
| 389 | Peptan | 458.2255 | 1 | 2.552767 | 119466.6 | 457.2182 | tri/oligope | N |
| 390 | Peptan | 458.2624 | 1 | 15.79448 | 610472.2 | 457.2552 | tri/oligope | N |
| 391 | Peptan | 458.2624 | 1 | 19.21448 | 551908.1 | 457.2552 | tri/oligope | N |
| 392 | Peptan | 460.2415 | 1 | 14.50872 | 1148174 | 459.2342 | tri/oligope | N |
| 393 | Peptan | 462.1838 | 1 | 2.031942 | 211890.4 | 461.1765 | tri/oligope | N |
| 394 | Peptan | 464.2156 | 1 | 16.85857 | 500541.3 | 463.2083 | tri/oligope | N |
| 395 | Peptan | 471.2572 | 1 | 2.2156 | 155177.1 | 470.2499 | tri/oligope | N |
| 396 | Peptan | 471.2576 | 1 | 3.840367 | 185450.4 | 470.2503 | tri/oligope | N |
| 397 | Peptan | 471.2576 | 1 | 9.174592 | 232146 | 470.2504 | tri/oligope | N |
| 398 | Peptan | 472.2416 | 1 | 15.9233 | 270997.8 | 471.2343 | tri/oligope | N |
| 399 | Peptan | 473.236 | 1 | 2.19435 | 294020.6 | 472.2287 | tri/oligope | N |
| 400 | Peptan | 238.6182 | 2 | 2.009375 | 298913.2 | 475.2219 | tri/oligope | N |
| 401 | Peptan | 485.2363 | 1 | 2.19435 | 486540.5 | 484.229 | tri/oligope | N |
| 402 | Peptan | 487.2154 | 1 | 2.19435 | 177247.1 | 486.2082 | tri/oligope | N |
| 403 | Peptan | 488.2363 | 1 | 13.10863 | 1719919 | 487.229 | tri/oligope | N |
| 404 | Peptan | 488.2367 | 1 | 12.19323 | 566408.7 | 487.2294 | tri/oligope | N |
| 405 | Peptan | 495.2572 | 1 | 2.41845 | 294763.8 | 494.2499 | tri/oligope | N |
| 406 | Peptan | 499.2521 | 1 | 2.19435 | 265676.6 | 498.2448 | tri/oligope | N |
| 407 | Peptan | 499.2884 | 1 | 2.489342 | 313494.7 | 498.2812 | tri/oligope | N |
| 408 | Peptan | 500.273 | 1 | 12.52728 | 197977.2 | 499.2657 | tri/oligope | N |
| 409 | Peptan | 501.2311 | 1 | 2.19435 | 473175.8 | 500.2238 | tri/oligope | N |
| 410 | Peptan | 501.2681 | 1 | 13.67786 | 148357.1 | 500.2608 | tri/oligope | N |
| 411 | Peptan | 501.2681 | 1 | 13.3276 | 123584.1 | 500.2609 | tri/oligope | N |
| 412 | Peptan | 501.2683 | 1 | 11.84976 | 109217.7 | 500.261 | tri/oligope | N |
| 413 | Peptan | 511.2519 | 1 | 2.19435 | 104664.7 | 510.2446 | tri/oligope | N |
| 414 | Peptan | 257.1614 | 2 | 2.19435 | 135827.8 | 512.3082 | tri/oligope | N |

| | | | | | | | | |
|---|---|---|---|---|---|---|---|---|
| 415 | Peptan | 516.242 | 1 | 2.19435 | 521926.4 | 515.2347 | tri/oligope | N |
| 416 | Peptan | 517.226 | 1 | 2.19435 | 187104.1 | 516.2188 | tri/oligope | N |
| 417 | Peptan | 519.2053 | 1 | 2.151417 | 310570.1 | 518.198 | tri/oligope | N |
| 418 | Peptan | 522.1873 | 1 | 2.19435 | 309975.1 | 521.18 | tri/oligope | N |
| 419 | Peptan | 522.2214 | 1 | 16.36224 | 944899.5 | 521.2141 | tri/oligope | N |
| 420 | Peptan | 527.2839 | 1 | 15.90679 | 682689.2 | 526.2767 | tri/oligope | N |
| 421 | Peptan | 264.1458 | 2 | 15.89866 | 187400.8 | 526.277 | tri/oligope | N |
| 422 | Peptan | 264.6612 | 2 | 2.19435 | 183872.6 | 527.3079 | tri/oligope | N |
| 423 | Peptan | 529.2632 | 1 | 11.16034 | 100220.2 | 528.256 | tri/oligope | N |
| 424 | Peptan | 529.2996 | 1 | 18.52648 | 216362.5 | 528.2924 | tri/oligope | N |
| 425 | Peptan | 529.2999 | 1 | 20.04718 | 313331.8 | 528.2926 | tri/oligope | N |
| 426 | Peptan | 531.2416 | 1 | 2.19435 | 390523.3 | 530.2344 | tri/oligope | N |
| 427 | Peptan | 531.2787 | 1 | 14.02315 | 368265 | 530.2714 | tri/oligope | N |
| 428 | Peptan | 535.2527 | 1 | 16.5157 | 577423.4 | 534.2454 | tri/oligope | N |
| 429 | Peptan | 537.3049 | 1 | 18.63977 | 167008.2 | 536.2976 | tri/oligope | N |
| 430 | Peptan | 539.3198 | 1 | 20.53473 | 1368667 | 538.3125 | tri/oligope | N |
| 431 | Peptan | 542.2575 | 1 | 2.19435 | 187149.5 | 541.2502 | tri/oligope | N |
| 432 | Peptan | 543.2418 | 1 | 2.19435 | 714348.8 | 542.2345 | tri/oligope | N |
| 433 | Peptan | 272.6405 | 2 | 1.975583 | 451345.6 | 543.2664 | tri/oligope | N |
| 434 | Peptan | 555.3158 | 1 | 18.75917 | 218094.3 | 554.3085 | tri/oligope | N |
| 435 | Peptan | 556.3213 | 1 | 2.183825 | 113748.2 | 555.314 | tri/oligope | N |
| 436 | Peptan | 278.6644 | 2 | 2.19435 | 440759.6 | 555.3142 | tri/oligope | N |
| 437 | Peptan | 557.2954 | 1 | 20.14087 | 121956.4 | 556.2881 | tri/oligope | N |
| 438 | Peptan | 557.3318 | 1 | 18.81414 | 288308.9 | 556.3245 | tri/oligope | N |
| 439 | Peptan | 559.3109 | 1 | 18.91823 | 167669.4 | 558.3036 | tri/oligope | N |
| 440 | Peptan | 568.2737 | 1 | 2.19435 | 333537.7 | 567.2664 | tri/oligope | N |
| 441 | Peptan | 285.6539 | 2 | 2.19435 | 337252.2 | 569.2933 | tri/oligope | N |
| 442 | Peptan | 574.2478 | 1 | 2.19435 | 524010.5 | 573.2405 | tri/oligope | N |
| 443 | Peptan | 579.2431 | 1 | 15.93778 | 205410.2 | 578.2358 | tri/oligope | N |
| 444 | Peptan | 579.2453 | 1 | 2.19435 | 182733.9 | 578.238 | tri/oligope | N |
| 445 | Peptan | 582.3278 | 1 | 17.23125 | 108949.4 | 581.3205 | tri/oligope | N |
| 446 | Peptan | 292.6743 | 2 | 2.2263 | 148248.7 | 583.3341 | tri/oligope | N |
| 447 | Peptan | 586.2845 | 1 | 2.183825 | 266234 | 585.2772 | tri/oligope | N |
| 448 | Peptan | 588.2645 | 1 | 2.19435 | 224927.6 | 587.2572 | tri/oligope | N |
| 449 | Peptan | 295.1605 | 2 | 2.19435 | 158698.1 | 588.3065 | tri/oligope | N |
| 450 | Peptan | 592.2746 | 1 | 16.24601 | 669490.9 | 591.2674 | tri/oligope | N |
| 451 | Peptan | 596.3428 | 1 | 19.58732 | 126738.6 | 595.3355 | tri/oligope | N |
| 452 | Peptan | 299.1721 | 2 | 2.19435 | 179496.5 | 596.3296 | tri/oligope | N |
| 453 | Peptan | 598.3218 | 1 | 15.11926 | 340932.4 | 597.3145 | tri/oligope | N |
| 454 | Peptan | 599.2797 | 1 | 2.183825 | 113623.1 | 598.2724 | tri/oligope | N |
| 455 | Peptan | 600.2629 | 1 | 2.183825 | 146870.6 | 599.2557 | tri/oligope | N |
| 456 | Peptan | 600.3375 | 1 | 19.33392 | 110860.1 | 599.3303 | tri/oligope | N |
| 457 | Peptan | 602.2801 | 1 | 13.31931 | 131868.2 | 601.2728 | tri/oligope | N |
| 458 | Peptan | 602.2802 | 1 | 12.59656 | 175580.5 | 601.273 | tri/oligope | N |
| 459 | Peptan | 301.6696 | 2 | 16.15702 | 215899 | 601.3245 | tri/oligope | N |
| 460 | Peptan | 608.342 | 1 | 18.576 | 432048.9 | 607.3348 | tri/oligope | N |
| 461 | Peptan | 610.2844 | 1 | 2.19435 | 579525 | 609.2771 | tri/oligope | N |
| 462 | Peptan | 611.3166 | 1 | 2.645558 | 124350.6 | 610.3093 | tri/oligope | N |
| 463 | Peptan | 612.3005 | 1 | 2.205033 | 270210.1 | 611.2932 | tri/oligope | N |
| 464 | Peptan | 612.337 | 1 | 18.56745 | 343667.9 | 611.3298 | tri/oligope | N |

| | | | | | | | | |
|---|---|---|---|---|---|---|---|---|
| 465 | Peptan | 612.3376 | 1 | 17.97792 | 142989.4 | 611.3303 | tri/oligope | N |
| 466 | Peptan | 307.6489 | 2 | 2.141275 | 183514.6 | 613.2833 | oligopepti | N |
| 467 | Peptan | 614.3534 | 1 | 20.22931 | 111193.9 | 613.3461 | oligopepti | N |
| 468 | Peptan | 615.3118 | 1 | 13.20163 | 238197.6 | 614.3045 | oligopepti | N |
| 469 | Peptan | 308.62 | 2 | 16.24601 | 144086.5 | 615.2254 | oligopepti | N |
| 470 | Peptan | 616.2955 | 1 | 13.79248 | 152923.4 | 615.2882 | oligopepti | N |
| 471 | Peptan | 618.2743 | 1 | 2.19435 | 118047.9 | 617.267 | oligopepti | N |
| 472 | Peptan | 624.3376 | 1 | 17.49828 | 109342.2 | 623.3303 | oligopepti | N |
| 473 | Peptan | 629.2901 | 1 | 2.183825 | 296983.7 | 628.2829 | oligopepti | N |
| 474 | Peptan | 629.2905 | 1 | 2.552767 | 157274.6 | 628.2832 | oligopepti | N |
| 475 | Peptan | 315.1489 | 2 | 2.552767 | 143646.1 | 628.2833 | oligopepti | N |
| 476 | Peptan | 630.2721 | 1 | 2.19435 | 141897 | 629.2648 | oligopepti | N |
| 477 | Peptan | 634.2853 | 1 | 16.3534 | 103200.1 | 633.278 | oligopepti | N |
| 478 | Peptan | 317.6646 | 2 | 13.10863 | 347925.4 | 633.3146 | oligopepti | N |
| 479 | Peptan | 321.1726 | 2 | 2.19435 | 1140137 | 640.3307 | oligopepti | N |
| 480 | Peptan | 641.3387 | 1 | 2.19435 | 233258.1 | 640.3314 | oligopepti | N |
| 481 | Peptan | 644.3276 | 1 | 17.28434 | 204689.5 | 643.3203 | oligopepti | N |
| 482 | Peptan | 644.329 | 1 | 20.51143 | 130085.6 | 643.3217 | oligopepti | N |
| 483 | Peptan | 655.3433 | 1 | 16.50208 | 198194.6 | 654.336 | oligopepti | N |
| 484 | Peptan | 329.1638 | 2 | 2.183825 | 148471.1 | 656.3131 | oligopepti | N |
| 485 | Peptan | 329.1827 | 2 | 2.19435 | 217249.3 | 656.3509 | oligopepti | N |
| 486 | Peptan | 669.3215 | 1 | 2.183825 | 173620.1 | 668.3143 | oligopepti | N |
| 487 | Peptan | 339.6654 | 2 | 14.49849 | 246522 | 677.3162 | oligopepti | N |
| 488 | Peptan | 342.1902 | 2 | 2.183825 | 113788.4 | 682.3658 | oligopepti | N |
| 489 | Peptan | 683.3748 | 1 | 18.90156 | 171739 | 682.3676 | oligopepti | N |
| 490 | Peptan | 342.1913 | 2 | 18.89199 | 111561.2 | 682.368 | oligopepti | N |
| 491 | Peptan | 686.3749 | 1 | 19.66499 | 165073.1 | 685.3676 | oligopepti | N |
| 492 | Peptan | 687.2952 | 1 | 2.183825 | 105754.2 | 686.2879 | oligopepti | N |
| 493 | Peptan | 689.275 | 1 | 2.19435 | 141832 | 688.2677 | oligopepti | N |
| 494 | Peptan | 701.3115 | 1 | 2.19435 | 109879.3 | 700.3042 | oligopepti | N |
| 495 | Peptan | 352.6651 | 2 | 2.054608 | 155487.1 | 703.3156 | oligopepti | N |
| 496 | Peptan | 705.3227 | 1 | 18.54459 | 781857.2 | 704.3155 | oligopepti | N |
| 497 | Peptan | 353.1652 | 2 | 18.55301 | 200011 | 704.3159 | oligopepti | N |
| 498 | Peptan | 358.6876 | 2 | 19.41008 | 110945.1 | 715.3606 | oligopepti | N |
| 499 | Peptan | 360.689 | 2 | 15.27584 | 114598.4 | 719.3634 | oligopepti | N |
| 500 | Peptan | 721.3177 | 1 | 15.98494 | 179717.7 | 720.3105 | oligopepti | N |
| 501 | Peptan | 362.7095 | 2 | 2.205033 | 333002.6 | 723.4045 | oligopepti | N |
| 502 | Peptan | 365.1441 | 2 | 18.54459 | 175315.7 | 728.2736 | oligopepti | N |
| 503 | Peptan | 366.2096 | 2 | 19.28442 | 341454.4 | 730.4046 | oligopepti | N |
| 504 | Peptan | 735.3339 | 1 | 21.19383 | 246976.8 | 734.3266 | oligopepti | N |
| 505 | Peptan | 736.3649 | 1 | 13.20163 | 146647.9 | 735.3577 | oligopepti | N |
| 506 | Peptan | 368.6863 | 2 | 13.16188 | 243450.7 | 735.358 | oligopepti | N |
| 507 | Peptan | 744.3553 | 1 | 14.19074 | 128410 | 743.3481 | oligopepti | N |
| 508 | Peptan | 755.3598 | 1 | 16.80773 | 211687.3 | 754.3526 | oligopepti | N |
| 509 | Peptan | 378.1837 | 2 | 16.78675 | 475024.9 | 754.3528 | oligopepti | N |
| 510 | Peptan | 378.1885 | 2 | 2.020658 | 116676.3 | 754.3625 | oligopepti | N |
| 511 | Peptan | 756.355 | 1 | 12.43064 | 103233.6 | 755.3477 | oligopepti | N |
| 512 | Peptan | 378.7051 | 2 | 12.63975 | 170538.2 | 755.3955 | oligopepti | N |
| 513 | Peptan | 764.4334 | 1 | 22.33383 | 176564.1 | 763.4261 | oligopepti | N |
| 514 | Peptan | 384.207 | 2 | 2.19435 | 519927.3 | 766.3994 | oligopepti | N |

| | | | | | | | | | |
|---|---|---|---|---|---|---|---|---|---|
| 515 | Peptan | 767.4067 | | 1 | 2.19435 | 105256.9 | 766.3994 | oligopepti | N |
| 516 | Peptan | 385.1962 | | 2 | 2.19435 | 271912 | 768.3778 | oligopepti | N |
| 517 | Peptan | 387.1884 | | 2 | 2.151417 | 272091.9 | 772.3623 | oligopepti | N |
| 518 | Peptan | 390.1626 | | 2 | 16.76308 | 108439.8 | 778.3106 | oligopepti | N |
| 519 | Peptan | 390.2125 | | 2 | 2.183825 | 292430.9 | 778.4105 | oligopepti | N |
| 520 | Peptan | 391.2023 | | 2 | 2.19435 | 379589.5 | 780.39 | oligopepti | N |
| 521 | Peptan | 392.2229 | | 2 | 13.89069 | 1723643 | 782.4312 | oligopepti | N |
| 522 | Peptan | 398.2104 | | 2 | 2.19435 | 202081.9 | 794.4062 | oligopepti | N |
| 523 | Peptan | 797.3808 | | 1 | 2.2156 | 108466.9 | 796.3735 | oligopepti | N |
| 524 | Peptan | 404.1724 | | 2 | 17.50731 | 156210.7 | 806.3302 | oligopepti | N |
| 525 | Peptan | 405.2129 | | 2 | 16.8015 | 125147.4 | 808.4112 | oligopepti | N |
| 526 | Peptan | 810.3648 | | 1 | 2.248483 | 113621.5 | 809.3575 | oligopepti | N |
| 527 | Peptan | 405.6861 | | 2 | 2.350883 | 518761.9 | 809.3577 | oligopepti | N |
| 528 | Peptan | 811.397 | | 1 | 13.72504 | 187092.2 | 810.3897 | oligopepti | N |
| 529 | Peptan | 406.2076 | | 2 | 2.19435 | 417788.4 | 810.4006 | oligopepti | N |
| 530 | Peptan | 411.2366 | | 2 | 16.61754 | 243252.3 | 820.4586 | oligopepti | N |
| 531 | Peptan | 414.2237 | | 2 | 12.6025 | 113616.7 | 826.4328 | oligopepti | N |
| 532 | Peptan | 417.6651 | | 2 | 2.350883 | 178014.3 | 833.3155 | oligopepti | N |
| 533 | Peptan | 418.7232 | | 2 | 2.19435 | 716155.3 | 835.4318 | oligopepti | N |
| 534 | Peptan | 419.2144 | | 2 | 2.2156 | 117241.7 | 836.4143 | oligopepti | N |
| 535 | Peptan | 424.2025 | | 2 | 17.69002 | 115569 | 846.3905 | oligopepti | N |
| 536 | Peptan | 427.2314 | | 2 | 14.88533 | 1247744 | 852.4483 | oligopepti | N |
| 537 | Peptan | 427.7103 | | 2 | 2.19435 | 476992.5 | 853.406 | oligopepti | N |
| 538 | Peptan | 439.7285 | | 2 | 2.19435 | 212269.7 | 877.4424 | oligopepti | N |
| 539 | Peptan | 440.2337 | | 2 | 18.6336 | 213419.7 | 878.4528 | oligopepti | N |
| 540 | Peptan | 441.7392 | | 2 | 16.99826 | 145301.9 | 881.4639 | oligopepti | N |
| 541 | Peptan | 448.2043 | | 2 | 14.57429 | 161881.3 | 894.394 | oligopepti | N |
| 542 | Peptan | 453.721 | | 2 | 14.21893 | 176388.2 | 905.4274 | oligopepti | N |
| 543 | Peptan | 458.2174 | | 2 | 14.89539 | 116681.1 | 914.4202 | oligopepti | N |
| 544 | Peptan | 462.2573 | | 2 | 2.19435 | 115100.6 | 922.5 | oligopepti | N |
| 545 | Peptan | 466.2316 | | 2 | 22.88008 | 140673 | 930.4486 | oligopepti | N |
| 546 | Peptan | 931.4559 | | 1 | 22.86679 | 137522.3 | 930.4487 | oligopepti | N |
| 547 | Peptan | 472.2289 | | 2 | 2.205033 | 112778.6 | 942.4433 | oligopepti | N |
| 548 | Peptan | 947.4505 | | 1 | 21.39534 | 148314.1 | 946.4432 | oligopepti | N |
| 549 | Peptan | 474.2289 | | 2 | 21.41241 | 160573.5 | 946.4433 | oligopepti | N |
| 550 | Peptan | 483.7364 | | 2 | 2.205033 | 151617.6 | 965.4582 | oligopepti | N |
| 551 | Peptan | 484.2475 | | 2 | 16.50208 | 188629 | 966.4804 | oligopepti | N |
| 552 | Peptan | 490.7396 | | 2 | 17.98568 | 180272.6 | 979.4646 | oligopepti | N |
| 553 | Peptan | 496.2662 | | 2 | 19.67261 | 285733.1 | 990.5178 | oligopepti | N |
| 554 | Peptan | 496.7759 | | 2 | 21.38751 | 687699.7 | 991.5372 | oligopepti | N |
| 555 | Peptan | 504.2631 | | 2 | 18.61848 | 467836.8 | 1006.512 | oligopepti | N |
| 556 | Peptan | 339.5059 | | 3 | 21.38147 | 122766.3 | 1015.496 | oligopepti | N |
| 557 | Peptan | 517.2712 | | 2 | 19.86586 | 137550.9 | 1032.528 | oligopepti | N |
| 558 | Peptan | 524.763 | | 2 | 2.205033 | 177180.3 | 1047.512 | oligopepti | N |
| 559 | Peptan | 524.7816 | | 2 | 2.534867 | 227953.4 | 1047.549 | oligopepti | N |
| 560 | Peptan | 524.7819 | | 2 | 13.56627 | 185409.4 | 1047.549 | oligopepti | N |
| 561 | Peptan | 526.2574 | | 2 | 2.183825 | 411992.2 | 1050.5 | oligopepti | N |
| 562 | Peptan | 528.7708 | | 2 | 2.282258 | 102308.2 | 1055.527 | oligopepti | N |
| 563 | Peptan | 538.2877 | | 2 | 15.23566 | 135271.9 | 1074.561 | oligopepti | N |
| 564 | Peptan | 538.774 | | 2 | 18.80704 | 273179.7 | 1075.533 | oligopepti | N |

| | | | | | | | | |
|---|---|---|---|---|---|---|---|---|
| 565 | Peptan | 542.7745 | 2 | 15.65792 | 142697.5 | 1083.534 | oligopepti( | N |
| 566 | Peptan | 544.7738 | 2 | 18.55301 | 565250.1 | 1087.533 | oligopepti( | N |
| 567 | Peptan | 544.774 | 2 | 18.96904 | 232217.3 | 1087.533 | oligopepti( | N |
| 568 | Peptan | 546.7715 | 2 | 17.67602 | 591132.4 | 1091.528 | oligopepti( | N |
| 569 | Peptan | 552.2564 | 2 | 21.61837 | 327806.3 | 1102.498 | oligopepti( | N |
| 570 | Peptan | 553.2874 | 2 | 19.44127 | 128949.5 | 1104.56 | oligopepti( | N |
| 571 | Peptan | 553.2928 | 2 | 14.76489 | 997932 | 1104.571 | oligopepti( | N |
| 572 | Peptan | 553.785 | 2 | 15.47732 | 253368.9 | 1105.555 | oligopepti( | N |
| 573 | Peptan | 554.7695 | 2 | 18.33628 | 116911.4 | 1107.524 | oligopepti( | N |
| 574 | Peptan | 558.253 | 2 | 2.19435 | 292871.7 | 1114.491 | oligopepti( | N |
| 575 | Peptan | 577.787 | 2 | 15.98494 | 281444.9 | 1153.559 | oligopepti( | N |
| 576 | Peptan | 581.8169 | 2 | 18.90156 | 281946.9 | 1161.619 | oligopepti( | N |
| 577 | Peptan | 390.1738 | 3 | 2.19435 | 128946.9 | 1167.5 | oligopepti( | N |
| 578 | Peptan | 600.7829 | 2 | 24.44879 | 265575.8 | 1199.551 | oligopepti( | N |
| 579 | Peptan | 601.2986 | 2 | 19.85294 | 258730.6 | 1200.583 | oligopepti( | N |
| 580 | Peptan | 601.7906 | 2 | 20.28936 | 290048.6 | 1201.567 | oligopepti( | N |
| 581 | Peptan | 608.7804 | 2 | 23.55998 | 133776.6 | 1215.546 | oligopepti( | N |
| 582 | Peptan | 634.3295 | 2 | 19.86586 | 176545.5 | 1266.644 | oligopepti( | N |
| 583 | Peptan | 638.8331 | 2 | 24.10134 | 144711.2 | 1275.652 | oligopepti( | N |
| 584 | Peptan | 638.8384 | 2 | 18.99985 | 252228.4 | 1275.662 | oligopepti( | N |
| 585 | Peptan | 639.3306 | 2 | 19.55983 | 233092.8 | 1276.647 | oligopepti( | N |
| 586 | Peptan | 666.8323 | 2 | 2.205033 | 442765.4 | 1331.65 | oligopepti( | N |
| 587 | Peptan | 667.3249 | 2 | 14.59383 | 124226.6 | 1332.635 | oligopepti( | N |
| 588 | Peptan | 695.3443 | 2 | 16.45014 | 118178.2 | 1388.674 | oligopepti( | N |
| 589 | Peptan | 715.8759 | 2 | 19.96082 | 107797.6 | 1429.737 | oligopepti( | N |
| 590 | Peptan | 731.8551 | 2 | 23.3877 | 239104.8 | 1461.696 | oligopepti( | N |
| 591 | Peptan | 749.3736 | 2 | 22.9781 | 100094.5 | 1496.733 | oligopepti( | N |
| 592 | Peptan | 764.3584 | 2 | 16.61754 | 257337.9 | 1526.702 | oligopepti( | N |
| 593 | Peptan | 767.3737 | 2 | 23.94483 | 189770.8 | 1532.733 | oligopepti( | N |
| 594 | Peptan | 781.8979 | 2 | 22.81726 | 152390.1 | 1561.781 | oligopepti( | N |
| 595 | Peptan | 843.3949 | 2 | 18.576 | 136310.5 | 1684.775 | oligopepti( | N |
| 596 | Peptan | 600.6066 | 3 | 15.31625 | 100602.6 | 1798.798 | oligopepti( | N |
| 597 | Peptan | 922.802 | 2 | 1.593725 | 253244.3 | 1843.589 | oligopepti( | N |
| 598 | Peptan | 1058.777 | 2 | 1.593725 | 148045.5 | 2115.54 | oligopepti( | N |

| ID | Protein | mz | charge | retentiont | raw.abund | mass | type | Match |
|---|---|---|---|---|---|---|---|---|
| 2 | LM | 231.171 | 1 | 5.400792 | 790372.3 | 230.1637 | di/tripepti | Y |
| 3 | LM | 231.171 | 1 | 9.743442 | 481015.2 | 230.1637 | di/tripepti | Y |
| 4 | LM | 231.1711 | 1 | 12.08171 | 762393.3 | 230.1638 | di/tripepti | Y |
| 5 | LM | 233.1499 | 1 | 2.187992 | 825371.5 | 232.1426 | di/tripepti | Y |
| 6 | LM | 233.1501 | 1 | 2.806692 | 282953.3 | 232.1428 | di/tripepti | Y |
| 7 | LM | 239.1029 | 1 | 2.187992 | 392966.5 | 238.0956 | di/tripepti | Y |
| 8 | LM | 239.103 | 1 | 2.447792 | 597528.1 | 238.0958 | di/tripepti | Y |
| 11 | LM | 245.1867 | 1 | 16.43414 | 2109532 | 244.1794 | di/tripepti | Y |
| 12 | LM | 245.1868 | 1 | 15.91003 | 330362.2 | 244.1795 | di/tripepti | Y |
| 13 | LM | 245.1867 | 1 | 14.55766 | 430262.7 | 244.1794 | di/tripepti | Y |
| 14 | LM | 245.1868 | 1 | 17.69159 | 349448 | 244.1795 | di/tripepti | Y |
| 15 | LM | 246.1452 | 1 | 2.187992 | 674200.9 | 245.1379 | di/tripepti | Y |
| 15 | LM | 246.1452 | 1 | 1.959917 | 251259.4 | 245.1379 | di/tripepti | Y |
| 16 | LM | 246.1454 | 1 | 2.66985 | 340039.6 | 245.1381 | di/tripepti | Y |
| 17 | LM | 246.1455 | 1 | 3.434625 | 647613.3 | 245.1382 | di/tripepti | Y |
| 20 | LM | 247.1292 | 1 | 2.187992 | 962733.7 | 246.1219 | di/tripepti | Y |
| 22 | LM | 247.1296 | 1 | 4.767342 | 934677.6 | 246.1223 | di/tripepti | Y |
| 23 | LM | 249.1274 | 1 | 2.822408 | 320545.5 | 248.1201 | di/tripepti | Y |
| 24 | LM | 250.1783 | 1 | 34.86093 | 708119.4 | 249.1711 | di/tripepti | Y |
| 25 | LM | 250.1784 | 1 | 37.25158 | 203109.1 | 249.1711 | di/tripepti | Y |
| 27 | LM | 253.1187 | 1 | 2.198958 | 777943.1 | 252.1114 | di/tripepti | Y |
| 27 | LM | 253.1187 | 1 | 2.513058 | 721125.8 | 252.1115 | di/tripepti | Y |
| 28 | LM | 253.119 | 1 | 9.369508 | 325801.9 | 252.1117 | di/tripepti | Y |
| 30 | LM | 260.1609 | 1 | 2.198958 | 900233 | 259.1536 | di/tripepti | Y |
| 31 | LM | 260.1611 | 1 | 2.679042 | 401502.1 | 259.1539 | di/tripepti | Y |
| 33 | LM | 260.1973 | 1 | 2.187992 | 177406.3 | 259.19 | di/tripepti | Y |
| 33 | LM | 260.1973 | 1 | 1.815142 | 143541.9 | 259.19 | di/tripepti | Y |
| 34 | LM | 261.1449 | 1 | 2.198958 | 877421.8 | 260.1377 | di/tripepti | Y |
| 35 | LM | 261.1453 | 1 | 3.349892 | 369742.4 | 260.138 | di/tripepti | Y |
| 36 | LM | 261.1453 | 1 | 5.518942 | 689477.6 | 260.138 | di/tripepti | Y |
| 38 | LM | 263.1432 | 1 | 11.57159 | 428758.1 | 262.1359 | di/tripepti | Y |
| 40 | LM | 265.1556 | 1 | 16.28167 | 745277.5 | 264.1484 | di/tripepti | Y |
| 41 | LM | 267.1345 | 1 | 2.424883 | 476588.9 | 266.1272 | di/tripepti | Y |
| 42 | LM | 269.1612 | 1 | 2.187992 | 541533.6 | 268.1539 | di/tripepti | Y |
| 44 | LM | 277.1035 | 1 | 1.863258 | 124110.4 | 276.0962 | di/tripepti | Y |
| 45 | LM | 281.1138 | 1 | 2.586833 | 381695.1 | 280.1065 | di/tripepti | Y |
| 46 | LM | 281.1505 | 1 | 7.14775 | 515517 | 280.1432 | di/tripepti | Y |
| 50 | LM | 294.1454 | 1 | 2.345083 | 345050.9 | 293.1381 | di/tripepti | Y |
| 51 | LM | 295.1298 | 1 | 3.385825 | 716262.6 | 294.1225 | di/tripepti | Y |
| 52 | LM | 295.1661 | 1 | 16.43414 | 927847.8 | 294.1588 | di/tripepti | Y |
| 57 | LM | 302.2084 | 1 | 19.87893 | 491743 | 301.2011 | tri/oligope | Y |
| 58 | LM | 303.1669 | 1 | 2.198958 | 164850.2 | 302.1596 | tri/oligope | Y |
| 58 | LM | 303.1673 | 1 | 2.649925 | 167910.7 | 302.16 | tri/oligope | Y |
| 59 | LM | 304.1509 | 1 | 2.210133 | 388629.5 | 303.1436 | tri/oligope | Y |
| 63 | LM | 318.1669 | 1 | 2.66985 | 165596.4 | 317.1596 | tri/oligope | Y |
| 64 | LM | 318.1666 | 1 | 2.210133 | 516146.7 | 317.1594 | tri/oligope | Y |
| 65 | LM | 318.167 | 1 | 9.162492 | 467533.8 | 317.1597 | tri/oligope | Y |
| 66 | LM | 318.1826 | 1 | 22.57528 | 349225.5 | 317.1753 | tri/oligope | Y |
| 66 | LM | 318.1826 | 1 | 22.90584 | 120259.8 | 317.1753 | tri/oligope | Y |

| | | | | | | | | |
|---|---|---|---|---|---|---|---|---|
| 73 | LM | 331.1662 | 1 | 2.85575 | 156018.8 | 330.1589 | tri/oligope | Y |
| 75 | LM | 332.1825 | 1 | 2.598342 | 215947.3 | 331.1752 | tri/oligope | Y |
| 76 | LM | 332.2191 | 1 | 3.113358 | 708843.7 | 331.2118 | tri/oligope | Y |
| 79 | LM | 333.1569 | 1 | 11.09138 | 125614.9 | 332.1496 | tri/oligope | Y |
| 80 | LM | 334.1615 | 1 | 2.198958 | 239286.5 | 333.1542 | tri/oligope | Y |
| 92 | LM | 360.2139 | 1 | 2.791033 | 161325.4 | 359.2066 | tri/oligope | Y |
| 94 | LM | 363.1558 | 1 | 2.187992 | 215902.2 | 362.1486 | tri/oligope | Y |
| 100 | LM | 376.1722 | 1 | 2.210133 | 105742.6 | 375.1649 | tri/oligope | Y |
| 106 | LM | 395.0398 | 1 | 1.598842 | 146851.5 | 394.0325 | tri/oligope | Y |
| 112 | LM | 415.2123 | 1 | 36.4673 | 258007.9 | 414.205 | tri/oligope | Y |
| 118 | LM | 429.2723 | 1 | 18.85342 | 170343.8 | 428.2651 | tri/oligope | Y |
| 134 | LM | 463.0273 | 1 | 1.621117 | 198866 | 462.02 | tri/oligope | Y |
| 141 | LM | 479.3109 | 1 | 37.16047 | 159411.4 | 478.3036 | tri/oligope | Y |
| 144 | LM | 488.2522 | 1 | 19.14933 | 138432.2 | 487.2449 | tri/oligope | Y |
| 151 | LM | 500.2888 | 1 | 25.19285 | 139476.2 | 499.2815 | tri/oligope | Y |
| 156 | LM | 515.292 | 1 | 37.08576 | 115294.3 | 514.2847 | tri/oligope | Y |
| 161 | LM | 264.1751 | 2 | 35.26698 | 108830.5 | 526.3357 | tri/oligope | Y |
| 165 | LM | 531.0146 | 1 | 1.610017 | 248139.9 | 530.0073 | tri/oligope | Y |
| 166 | LM | 531.3877 | 1 | 37.84633 | 100068 | 530.3804 | tri/oligope | Y |
| 174 | LM | 274.1681 | 2 | 35.92648 | 111895.7 | 546.3216 | tri/oligope | Y |
| 194 | LM | 599.0024 | 1 | 1.610017 | 226038.2 | 597.9952 | tri/oligope | Y |
| 218 | LM | 666.99 | 1 | 1.610017 | 219741.3 | 665.9827 | oligopepti | Y |
| 240 | LM | 734.9775 | 1 | 1.610017 | 196535.2 | 733.9702 | oligopepti | Y |
| 258 | LM | 802.9651 | 1 | 1.610017 | 173428.3 | 801.9578 | oligopepti | Y |
| 276 | LM | 870.9528 | 1 | 1.598842 | 118476.4 | 869.9455 | oligopepti | Y |
| 296 | LM | 974.8153 | 1 | 1.621117 | 266358 | 973.808 | oligopepti | Y |
| 297 | LM | 990.7896 | 1 | 1.621117 | 202825.3 | 989.7823 | oligopepti | Y |
| 302 | LM | 1042.802 | 1 | 1.621117 | 156478.3 | 1041.795 | oligopepti | Y |
| 313 | LM | 582.9462 | 2 | 1.610017 | 101013.2 | 1163.878 | oligopepti | Y |
| 316 | LM | 616.9401 | 2 | 1.621117 | 103805.2 | 1231.866 | oligopepti | Y |
| 318 | LM | 624.9268 | 2 | 1.610017 | 106833.9 | 1247.839 | oligopepti | Y |
| 329 | LM | 692.9145 | 2 | 1.621117 | 103839.1 | 1383.814 | oligopepti | Y |
| 338 | LM | 223.1083 | 1 | 5.106542 | 483354.7 | 222.1011 | dipeptide | N |
| 339 | LM | 223.1084 | 1 | 10.47817 | 431659.3 | 222.1011 | dipeptide | N |
| 340 | LM | 229.1554 | 1 | 5.049692 | 153906.4 | 228.1481 | di/tripepti | N |
| 341 | LM | 231.171 | 1 | 9.006225 | 291605.9 | 230.1637 | di/tripepti | N |
| 342 | LM | 233.1135 | 1 | 2.198958 | 120423.1 | 232.1062 | di/tripepti | N |
| 343 | LM | 233.1136 | 1 | 2.005575 | 198786.9 | 232.1063 | di/tripepti | N |
| 344 | LM | 233.1503 | 1 | 4.133825 | 378099.2 | 232.143 | di/tripepti | N |
| 345 | LM | 237.124 | 1 | 11.23058 | 456998.2 | 236.1167 | di/tripepti | N |
| 346 | LM | 237.124 | 1 | 4.676475 | 235116.5 | 236.1168 | di/tripepti | N |
| 347 | LM | 246.1456 | 1 | 5.425042 | 235592.3 | 245.1383 | di/tripepti | N |
| 348 | LM | 252.963 | 1 | 8.455267 | 138554 | 251.9558 | di/tripepti | N |
| 349 | LM | 258.1108 | 1 | 1.724358 | 134427.6 | 257.1036 | di/tripepti | N |
| 350 | LM | 262.1196 | 1 | 17.00185 | 100435.3 | 261.1124 | di/tripepti | N |
| 351 | LM | 263.1399 | 1 | 14.47133 | 166826.9 | 262.1326 | di/tripepti | N |
| 352 | LM | 263.1433 | 1 | 13.14811 | 178151.8 | 262.136 | di/tripepti | N |
| 353 | LM | 265.1556 | 1 | 14.76838 | 368743.1 | 264.1483 | di/tripepti | N |
| 354 | LM | 269.0884 | 1 | 2.198958 | 186821.6 | 268.0811 | di/tripepti | N |
| 355 | LM | 269.1135 | 1 | 2.187992 | 386596 | 268.1062 | di/tripepti | N |

| | | | | | | | | |
|---|---|---|---|---|---|---|---|---|
| 356 | LM | 269.1137 | 1 | 2.345083 | 382219.4 | 268.1065 | di/tripepti | N |
| 357 | LM | 274.1401 | 1 | 2.187992 | 109126.7 | 273.1328 | di/tripepti | N |
| 358 | LM | 274.1766 | 1 | 2.27715 | 178221.5 | 273.1693 | di/tripepti | N |
| 359 | LM | 274.1767 | 1 | 2.501892 | 133789.1 | 273.1694 | di/tripepti | N |
| 360 | LM | 276.1353 | 1 | 14.35843 | 164860.9 | 275.128 | di/tripepti | N |
| 361 | LM | 276.1354 | 1 | 17.19731 | 115912.2 | 275.1281 | di/tripepti | N |
| 362 | LM | 276.1559 | 1 | 2.198958 | 249873.7 | 275.1486 | di/tripepti | N |
| 363 | LM | 276.1561 | 1 | 2.413558 | 148626.7 | 275.1488 | di/tripepti | N |
| 364 | LM | 276.1563 | 1 | 5.971225 | 130143.3 | 275.1491 | di/tripepti | N |
| 365 | LM | 277.1194 | 1 | 20.44814 | 296245.7 | 276.1121 | di/tripepti | N |
| 366 | LM | 279.1348 | 1 | 9.507417 | 148050.9 | 278.1276 | di/tripepti | N |
| 367 | LM | 279.1348 | 1 | 3.796183 | 265994.3 | 278.1276 | di/tripepti | N |
| 368 | LM | 279.1713 | 1 | 19.91518 | 944712.8 | 278.164 | di/tripepti | N |
| 369 | LM | 279.1714 | 1 | 19.00699 | 140545.5 | 278.1641 | di/tripepti | N |
| 370 | LM | 279.1714 | 1 | 21.04403 | 229202.6 | 278.1641 | di/tripepti | N |
| 371 | LM | 279.2325 | 1 | 37.01129 | 233685.7 | 278.2252 | di/tripepti | N |
| 372 | LM | 280.1296 | 1 | 2.221225 | 220320.3 | 279.1223 | di/tripepti | N |
| 373 | LM | 280.1302 | 1 | 10.05645 | 105074.3 | 279.1229 | di/tripepti | N |
| 374 | LM | 281.1141 | 1 | 11.60508 | 401405.3 | 280.1068 | di/tripepti | N |
| 375 | LM | 281.1505 | 1 | 6.1342 | 623305.1 | 280.1432 | di/tripepti | N |
| 376 | LM | 283.1293 | 1 | 2.198958 | 277670.6 | 282.122 | di/tripepti | N |
| 377 | LM | 286.1771 | 1 | 13.53491 | 103690 | 285.1699 | di/tripepti | N |
| 378 | LM | 288.1557 | 1 | 2.17705 | 125772.7 | 287.1485 | di/tripepti | N |
| 379 | LM | 288.1922 | 1 | 2.243617 | 206569.4 | 287.185 | di/tripepti | N |
| 380 | LM | 288.1924 | 1 | 2.402158 | 122764.8 | 287.1851 | di/tripepti | N |
| 381 | LM | 288.1926 | 1 | 9.516258 | 243417.8 | 287.1854 | di/tripepti | N |
| 382 | LM | 288.1927 | 1 | 10.74696 | 116072.6 | 287.1854 | di/tripepti | N |
| 383 | LM | 288.2035 | 1 | 1.9941 | 102758.4 | 287.1962 | di/tripepti | N |
| 384 | LM | 290.1351 | 1 | 2.039967 | 114042.1 | 289.1278 | di/tripepti | N |
| 385 | LM | 290.1715 | 1 | 2.198958 | 572393.6 | 289.1643 | di/tripepti | N |
| 386 | LM | 292.1301 | 1 | 16.43414 | 110485.1 | 291.1228 | di/tripepti | N |
| 387 | LM | 295.1299 | 1 | 12.67196 | 381692.4 | 294.1227 | di/tripepti | N |
| 388 | LM | 295.1662 | 1 | 14.63307 | 409113.1 | 294.159 | di/tripepti | N |
| 389 | LM | 295.1663 | 1 | 13.44093 | 918679.1 | 294.159 | di/tripepti | N |
| 390 | LM | 295.2275 | 1 | 37.37008 | 124380.9 | 294.2202 | di/tripepti | N |
| 391 | LM | 296.1246 | 1 | 2.198958 | 278559 | 295.1174 | di/tripepti | N |
| 392 | LM | 296.1248 | 1 | 2.413558 | 128894.9 | 295.1176 | di/tripepti | N |
| 393 | LM | 297.1086 | 1 | 2.198958 | 140735.9 | 296.1013 | di/tripepti | N |
| 394 | LM | 297.1089 | 1 | 2.717092 | 383503.5 | 296.1016 | di/tripepti | N |
| 395 | LM | 300.1928 | 1 | 4.074342 | 122548.7 | 299.1855 | di/tripepti | N |
| 396 | LM | 300.1929 | 1 | 13.41257 | 380115.2 | 299.1856 | di/tripepti | N |
| 397 | LM | 300.1929 | 1 | 12.08773 | 122686.8 | 299.1856 | di/tripepti | N |
| 398 | LM | 300.1929 | 1 | 13.69647 | 188393.1 | 299.1857 | di/tripepti | N |
| 399 | LM | 302.2084 | 1 | 11.18603 | 130543 | 301.2012 | tri/oligope | N |
| 400 | LM | 302.2084 | 1 | 13.14811 | 125786.2 | 301.2012 | tri/oligope | N |
| 401 | LM | 302.2085 | 1 | 6.428992 | 163427.7 | 301.2012 | tri/oligope | N |
| 402 | LM | 302.2086 | 1 | 15.83846 | 138480.4 | 301.2013 | tri/oligope | N |
| 403 | LM | 304.1665 | 1 | 19.92085 | 484931.5 | 303.1592 | tri/oligope | N |
| 404 | LM | 304.1872 | 1 | 2.198958 | 141633.7 | 303.1799 | tri/oligope | N |
| 405 | LM | 304.1874 | 1 | 2.424883 | 104670.7 | 303.1801 | tri/oligope | N |

| | | | | | | | | |
|---|---|---|---|---|---|---|---|---|
| 406 | LM | 304.1877 | 1 | 11.40626 | 128839 | 303.1804 | tri/oligope | N |
| 407 | LM | 306.1462 | 1 | 10.8124 | 103856.2 | 305.139 | tri/oligope | N |
| 408 | LM | 306.1665 | 1 | 2.187992 | 174350.2 | 305.1592 | tri/oligope | N |
| 409 | LM | 310.1402 | 1 | 2.210133 | 673439.5 | 309.133 | tri/oligope | N |
| 410 | LM | 310.1404 | 1 | 2.447792 | 157134.1 | 309.1332 | tri/oligope | N |
| 411 | LM | 310.1408 | 1 | 9.923342 | 119633.8 | 309.1335 | tri/oligope | N |
| 412 | LM | 310.1767 | 1 | 1.97135 | 101008.2 | 309.1694 | tri/oligope | N |
| 413 | LM | 311.1242 | 1 | 2.198958 | 363040.9 | 310.117 | tri/oligope | N |
| 414 | LM | 311.1246 | 1 | 2.799667 | 310704.6 | 310.1173 | tri/oligope | N |
| 415 | LM | 313.1227 | 1 | 11.82952 | 284120.4 | 312.1154 | tri/oligope | N |
| 416 | LM | 314.2085 | 1 | 13.68121 | 242073.4 | 313.2012 | tri/oligope | N |
| 417 | LM | 314.2085 | 1 | 13.03123 | 125922.9 | 313.2012 | tri/oligope | N |
| 418 | LM | 316.1879 | 1 | 13.06332 | 139702.9 | 315.1807 | tri/oligope | N |
| 419 | LM | 316.188 | 1 | 14.58649 | 176512 | 315.1807 | tri/oligope | N |
| 420 | LM | 316.2243 | 1 | 14.84319 | 143387.7 | 315.2171 | tri/oligope | N |
| 421 | LM | 316.2244 | 1 | 19.60503 | 194943.7 | 315.2171 | tri/oligope | N |
| 422 | LM | 317.1825 | 1 | 2.198958 | 222620.1 | 316.1753 | tri/oligope | N |
| 423 | LM | 317.1831 | 1 | 3.529967 | 212051.2 | 316.1758 | tri/oligope | N |
| 424 | LM | 318.1671 | 1 | 5.5851 | 182424 | 317.1598 | tri/oligope | N |
| 425 | LM | 318.2029 | 1 | 2.210133 | 225729 | 317.1956 | tri/oligope | N |
| 426 | LM | 318.2035 | 1 | 8.1336 | 133853.8 | 317.1963 | tri/oligope | N |
| 427 | LM | 318.2037 | 1 | 16.1012 | 107512.2 | 317.1964 | tri/oligope | N |
| 428 | LM | 319.1414 | 1 | 16.83336 | 156950.2 | 318.1341 | tri/oligope | N |
| 429 | LM | 320.1254 | 1 | 17.66704 | 116637.4 | 319.1181 | tri/oligope | N |
| 430 | LM | 320.1458 | 1 | 2.039967 | 130254.6 | 319.1386 | tri/oligope | N |
| 431 | LM | 320.1619 | 1 | 18.3665 | 194554.8 | 319.1546 | tri/oligope | N |
| 432 | LM | 328.2244 | 1 | 16.88842 | 137668.4 | 327.2172 | tri/oligope | N |
| 433 | LM | 328.2244 | 1 | 17.11261 | 189201 | 327.2172 | tri/oligope | N |
| 434 | LM | 328.2245 | 1 | 17.80768 | 102801.6 | 327.2172 | tri/oligope | N |
| 435 | LM | 329.1825 | 1 | 2.198958 | 627822.8 | 328.1752 | tri/oligope | N |
| 436 | LM | 331.1982 | 1 | 2.198958 | 159919.4 | 330.191 | tri/oligope | N |
| 437 | LM | 332.1828 | 1 | 11.64338 | 108197.6 | 331.1755 | tri/oligope | N |
| 438 | LM | 333.1774 | 1 | 2.187992 | 382225.8 | 332.1701 | tri/oligope | N |
| 439 | LM | 333.1778 | 1 | 2.658675 | 388816.9 | 332.1705 | tri/oligope | N |
| 440 | LM | 334.141 | 1 | 13.77377 | 213315.8 | 333.1337 | tri/oligope | N |
| 441 | LM | 334.162 | 1 | 5.175725 | 116668.8 | 333.1547 | tri/oligope | N |
| 442 | LM | 334.1775 | 1 | 18.0527 | 204287.5 | 333.1702 | tri/oligope | N |
| 443 | LM | 336.1566 | 1 | 12.89045 | 131672.7 | 335.1493 | tri/oligope | N |
| 444 | LM | 338.1719 | 1 | 2.490767 | 106418.2 | 337.1646 | tri/oligope | N |
| 445 | LM | 340.151 | 1 | 2.27715 | 108871.3 | 339.1438 | tri/oligope | N |
| 446 | LM | 342.2403 | 1 | 21.30493 | 121678.6 | 341.233 | tri/oligope | N |
| 447 | LM | 342.2403 | 1 | 20.75707 | 112204.1 | 341.233 | tri/oligope | N |
| 448 | LM | 343.1985 | 1 | 2.221225 | 235806.6 | 342.1912 | tri/oligope | N |
| 449 | LM | 345.1459 | 1 | 14.0599 | 112968.2 | 344.1386 | tri/oligope | N |
| 450 | LM | 345.214 | 1 | 2.198958 | 108736 | 344.2068 | tri/oligope | N |
| 451 | LM | 346.1808 | 1 | 14.51786 | 128588.1 | 345.1735 | tri/oligope | N |
| 452 | LM | 346.1982 | 1 | 2.479592 | 135395.3 | 345.191 | tri/oligope | N |
| 453 | LM | 346.2349 | 1 | 14.60074 | 132960.4 | 345.2276 | tri/oligope | N |
| 454 | LM | 346.2351 | 1 | 13.57153 | 128729.3 | 345.2278 | tri/oligope | N |
| 455 | LM | 348.1772 | 1 | 2.198958 | 392067.3 | 347.17 | tri/oligope | N |

| | | | | | | | | |
|---|---|---|---|---|---|---|---|---|
| 456 | LM | 350.1723 | 1 | 12.41783 | 397774.7 | 349.165 | tri/oligope | N |
| 457 | LM | 352.1879 | 1 | 11.18603 | 112614.8 | 351.1807 | tri/oligope | N |
| 458 | LM | 357.2141 | 1 | 2.288542 | 738446 | 356.2068 | tri/oligope | N |
| 459 | LM | 357.2142 | 1 | 2.791033 | 2133665 | 356.2069 | tri/oligope | N |
| 460 | LM | 357.2145 | 1 | 11.2235 | 194576.1 | 356.2072 | tri/oligope | N |
| 461 | LM | 357.2146 | 1 | 12.91559 | 112820.5 | 356.2073 | tri/oligope | N |
| 462 | LM | 358.296 | 1 | 36.28283 | 224752.3 | 357.2887 | tri/oligope | N |
| 463 | LM | 359.2302 | 1 | 7.407167 | 157784.5 | 358.223 | tri/oligope | N |
| 464 | LM | 359.2303 | 1 | 16.17923 | 126130.2 | 358.2231 | tri/oligope | N |
| 465 | LM | 360.2145 | 1 | 20.46971 | 147907.7 | 359.2072 | tri/oligope | N |
| 466 | LM | 361.1725 | 1 | 2.187992 | 334741.3 | 360.1652 | tri/oligope | N |
| 467 | LM | 361.2088 | 1 | 2.210133 | 117349.3 | 360.2015 | tri/oligope | N |
| 468 | LM | 362.193 | 1 | 2.27715 | 357634.7 | 361.1858 | tri/oligope | N |
| 469 | LM | 362.209 | 1 | 21.35968 | 265200.1 | 361.2017 | tri/oligope | N |
| 470 | LM | 364.066 | 1 | 2.210133 | 152598.2 | 363.0587 | tri/oligope | N |
| 471 | LM | 364.1883 | 1 | 18.27964 | 107759.6 | 363.181 | tri/oligope | N |
| 472 | LM | 371.2301 | 1 | 11.47663 | 599971.6 | 370.2229 | tri/oligope | N |
| 473 | LM | 371.2301 | 1 | 14.64038 | 482636.6 | 370.2229 | tri/oligope | N |
| 474 | LM | 371.2302 | 1 | 14.208 | 383515.1 | 370.2229 | tri/oligope | N |
| 475 | LM | 371.2302 | 1 | 12.52073 | 486803.2 | 370.223 | tri/oligope | N |
| 476 | LM | 373.2461 | 1 | 18.83558 | 161186.7 | 372.2389 | tri/oligope | N |
| 477 | LM | 374.2039 | 1 | 2.187992 | 144360.8 | 373.1966 | tri/oligope | N |
| 478 | LM | 377.2038 | 1 | 2.243617 | 114531.1 | 376.1965 | tri/oligope | N |
| 479 | LM | 378.2039 | 1 | 15.12353 | 176720.4 | 377.1966 | tri/oligope | N |
| 480 | LM | 385.2461 | 1 | 15.47318 | 121649.8 | 384.2388 | tri/oligope | N |
| 481 | LM | 387.2247 | 1 | 2.775042 | 102517.1 | 386.2174 | tri/oligope | N |
| 482 | LM | 389.2037 | 1 | 2.198958 | 171793.5 | 388.1965 | tri/oligope | N |
| 483 | LM | 391.1999 | 1 | 14.98866 | 161478.2 | 390.1927 | tri/oligope | N |
| 484 | LM | 396.1775 | 1 | 2.66985 | 200787.6 | 395.1702 | tri/oligope | N |
| 485 | LM | 397.1726 | 1 | 2.345083 | 105775.7 | 396.1653 | tri/oligope | N |
| 486 | LM | 399.2612 | 1 | 16.42042 | 139763.6 | 398.2539 | tri/oligope | N |
| 487 | LM | 399.2617 | 1 | 16.76551 | 116559.9 | 398.2545 | tri/oligope | N |
| 488 | LM | 400.2197 | 1 | 2.187992 | 153912.8 | 399.2124 | tri/oligope | N |
| 489 | LM | 401.2408 | 1 | 10.94893 | 147305.7 | 400.2336 | tri/oligope | N |
| 490 | LM | 402.1673 | 1 | 12.61625 | 105480.4 | 401.1601 | tri/oligope | N |
| 491 | LM | 403.2567 | 1 | 16.91779 | 147257.6 | 402.2494 | tri/oligope | N |
| 492 | LM | 403.2567 | 1 | 18.06443 | 187279.4 | 402.2494 | tri/oligope | N |
| 493 | LM | 405.1987 | 1 | 2.187992 | 108081.6 | 404.1914 | tri/oligope | N |
| 494 | LM | 408.1779 | 1 | 11.54782 | 155927 | 407.1706 | tri/oligope | N |
| 495 | LM | 412.1723 | 1 | 2.311033 | 110393.8 | 411.165 | tri/oligope | N |
| 496 | LM | 414.236 | 1 | 11.06041 | 173266.3 | 413.2287 | tri/oligope | N |
| 497 | LM | 415.2202 | 1 | 14.49839 | 110830 | 414.2129 | tri/oligope | N |
| 498 | LM | 419.1779 | 1 | 2.198958 | 102442.4 | 418.1706 | tri/oligope | N |
| 499 | LM | 421.2095 | 1 | 14.79268 | 129399.4 | 420.2023 | tri/oligope | N |
| 500 | LM | 421.2096 | 1 | 13.92144 | 130000.2 | 420.2024 | tri/oligope | N |
| 501 | LM | 423.2356 | 1 | 2.187992 | 166871.9 | 422.2283 | tri/oligope | N |
| 502 | LM | 426.204 | 1 | 23.88436 | 232113.6 | 425.1968 | tri/oligope | N |
| 503 | LM | 428.2513 | 1 | 2.557658 | 678014 | 427.244 | tri/oligope | N |
| 504 | LM | 428.2517 | 1 | 11.47663 | 231734.8 | 427.2444 | tri/oligope | N |
| 505 | LM | 428.2518 | 1 | 13.63509 | 204905.2 | 427.2445 | tri/oligope | N |

| | | | | | | | | |
|---|---|---|---|---|---|---|---|---|
| 506 | LM | 436.3433 | 1 | 35.94393 | 457356.2 | 435.336 | tri/oligope | N |
| 507 | LM | 436.3437 | 1 | 0.020917 | 262032 | 435.3364 | tri/oligope | N |
| 508 | LM | 437.2045 | 1 | 12.55126 | 116842.6 | 436.1973 | tri/oligope | N |
| 509 | LM | 439.268 | 1 | 14.29792 | 125572 | 438.2607 | tri/oligope | N |
| 510 | LM | 439.2934 | 1 | 22.64638 | 190586.6 | 438.2861 | tri/oligope | N |
| 511 | LM | 442.1989 | 1 | 17.61672 | 326654 | 441.1916 | tri/oligope | N |
| 512 | LM | 442.2673 | 1 | 13.37642 | 334991.6 | 441.2601 | tri/oligope | N |
| 513 | LM | 442.2674 | 1 | 14.4488 | 311248.1 | 441.2602 | tri/oligope | N |
| 514 | LM | 442.2675 | 1 | 15.86203 | 123991.1 | 441.2602 | tri/oligope | N |
| 515 | LM | 444.2461 | 1 | 2.221225 | 185431 | 443.2388 | tri/oligope | N |
| 516 | LM | 446.2624 | 1 | 16.72344 | 199736.1 | 445.2552 | tri/oligope | N |
| 517 | LM | 450.1626 | 1 | 2.187992 | 259521.9 | 449.1553 | tri/oligope | N |
| 518 | LM | 456.2829 | 1 | 15.94793 | 575865.5 | 455.2756 | tri/oligope | N |
| 519 | LM | 458.2623 | 1 | 11.12248 | 100197.8 | 457.2551 | tri/oligope | N |
| 520 | LM | 463.2989 | 1 | 37.55269 | 106520.1 | 462.2916 | tri/oligope | N |
| 521 | LM | 472.2781 | 1 | 16.56067 | 500892.6 | 471.2708 | tri/oligope | N |
| 522 | LM | 478.231 | 1 | 11.19344 | 184813.5 | 477.2238 | tri/oligope | N |
| 523 | LM | 491.2516 | 1 | 19.99671 | 320056.8 | 490.2443 | tri/oligope | N |
| 524 | LM | 491.2519 | 1 | 24.47185 | 110392 | 490.2447 | tri/oligope | N |
| 525 | LM | 492.2468 | 1 | 13.71196 | 227828.6 | 491.2395 | tri/oligope | N |
| 526 | LM | 494.2726 | 1 | 2.198958 | 551168.6 | 493.2653 | tri/oligope | N |
| 527 | LM | 247.64 | 2 | 2.187992 | 204242.6 | 493.2655 | tri/oligope | N |
| 528 | LM | 496.2413 | 1 | 16.38859 | 148665.9 | 495.234 | tri/oligope | N |
| 529 | LM | 498.902 | 1 | 1.610017 | 598312.4 | 497.8947 | tri/oligope | N |
| 530 | LM | 502.2712 | 1 | 18.72871 | 135789.4 | 501.2639 | tri/oligope | N |
| 531 | LM | 504.2835 | 1 | 21.433 | 134856.5 | 503.2762 | tri/oligope | N |
| 532 | LM | 508.2892 | 1 | 11.17733 | 239454.9 | 507.282 | tri/oligope | N |
| 533 | LM | 254.6484 | 2 | 11.2078 | 155440.3 | 507.2822 | tri/oligope | N |
| 534 | LM | 257.6537 | 2 | 11.85296 | 109268.5 | 513.2928 | tri/oligope | N |
| 535 | LM | 520.2784 | 1 | 18.28602 | 272917.5 | 519.2711 | tri/oligope | N |
| 536 | LM | 526.3256 | 1 | 23.384 | 173721.8 | 525.3183 | tri/oligope | N |
| 537 | LM | 554.332 | 1 | 15.341 | 164807.8 | 553.3247 | tri/oligope | N |
| 538 | LM | 283.1586 | 2 | 2.198958 | 123969.8 | 564.3027 | tri/oligope | N |
| 539 | LM | 566.3316 | 1 | 16.3722 | 143936.8 | 565.3243 | tri/oligope | N |
| 540 | LM | 283.6694 | 2 | 16.38113 | 151513.6 | 565.3243 | tri/oligope | N |
| 541 | LM | 595.2901 | 1 | 24.75063 | 128487.8 | 594.2828 | tri/oligope | N |
| 542 | LM | 316.1875 | 2 | 2.379283 | 104989.3 | 630.3604 | oligopepti | N |
| 543 | LM | 639.3738 | 1 | 19.38297 | 161934.5 | 638.3665 | oligopepti | N |
| 544 | LM | 320.1907 | 2 | 19.36935 | 109235.1 | 638.3668 | oligopepti | N |
| 545 | LM | 336.6725 | 2 | 15.77554 | 129132.9 | 671.3305 | oligopepti | N |
| 546 | LM | 384.1789 | 2 | 17.32999 | 107188.3 | 766.3433 | oligopepti | N |
| 547 | LM | 384.7172 | 2 | 2.557658 | 156029.5 | 767.4198 | oligopepti | N |
| 548 | LM | 820.8202 | 1 | 1.632142 | 116464.2 | 819.8129 | oligopepti | N |
| 549 | LM | 439.226 | 2 | 14.84319 | 207156.7 | 876.4375 | oligopepti | N |
| 550 | LM | 460.6672 | 2 | 14.28904 | 111083.6 | 919.3199 | oligopepti | N |
| 551 | LM | 475.7421 | 2 | 23.89188 | 170404.7 | 949.4696 | oligopepti | N |
| 552 | LM | 1058.777 | 2 | 1.621117 | 139747.5 | 2115.539 | oligopepti | N |

| ID | Protein | mz | charge | retentiont | raw.abund | mass | type | Match |
|---|---|---|---|---|---|---|---|---|
| 2 | LMC2 | 231.171 | 1 | 5.308483 | 162965.6 | 230.1637 | di/tripepti | Y |
| 5 | LMC2 | 233.1499 | 1 | 2.178042 | 171408.1 | 232.1426 | di/tripepti | Y |
| 8 | LMC2 | 239.103 | 1 | 2.447683 | 121182.2 | 238.0958 | di/tripepti | Y |
| 11 | LMC2 | 245.1867 | 1 | 16.4214 | 109168.8 | 244.1794 | di/tripepti | Y |
| 15 | LMC2 | 246.1452 | 1 | 2.178042 | 182636.1 | 245.1379 | di/tripepti | Y |
| 20 | LMC2 | 247.1292 | 1 | 2.178042 | 414327.4 | 246.1219 | di/tripepti | Y |
| 22 | LMC2 | 247.1296 | 1 | 4.667183 | 218239.9 | 246.1223 | di/tripepti | Y |
| 24 | LMC2 | 250.1783 | 1 | 34.86174 | 753175.9 | 249.1711 | di/tripepti | Y |
| 25 | LMC2 | 250.1784 | 1 | 37.25502 | 204283.7 | 249.1711 | di/tripepti | Y |
| 30 | LMC2 | 260.1609 | 1 | 2.189583 | 161266.1 | 259.1536 | di/tripepti | Y |
| 34 | LMC2 | 261.1449 | 1 | 2.189583 | 331013.5 | 260.1377 | di/tripepti | Y |
| 36 | LMC2 | 261.1453 | 1 | 5.425975 | 114040.5 | 260.138 | di/tripepti | Y |
| 44 | LMC2 | 277.1035 | 1 | 1.8456 | 115215.8 | 276.0962 | di/tripepti | Y |
| 45 | LMC2 | 281.1138 | 1 | 2.592508 | 112078.4 | 280.1065 | di/tripepti | Y |
| 51 | LMC2 | 295.1298 | 1 | 3.259108 | 148779.1 | 294.1225 | di/tripepti | Y |
| 57 | LMC2 | 302.2084 | 1 | 19.87918 | 276345.1 | 301.2011 | tri/oligope | Y |
| 59 | LMC2 | 304.1509 | 1 | 2.201183 | 208868.4 | 303.1436 | tri/oligope | Y |
| 64 | LMC2 | 318.1666 | 1 | 2.201183 | 244948.6 | 317.1594 | tri/oligope | Y |
| 65 | LMC2 | 318.167 | 1 | 9.080292 | 216788.3 | 317.1597 | tri/oligope | Y |
| 75 | LMC2 | 332.1825 | 1 | 2.602717 | 133828.6 | 331.1752 | tri/oligope | Y |
| 76 | LMC2 | 332.2191 | 1 | 3.025442 | 104777.8 | 331.2118 | tri/oligope | Y |
| 80 | LMC2 | 334.1615 | 1 | 2.189583 | 113988.3 | 333.1542 | tri/oligope | Y |
| 105 | LMC2 | 390.188 | 1 | 2.340608 | 105600.9 | 389.1807 | tri/oligope | Y |
| 106 | LMC2 | 395.0398 | 1 | 1.577917 | 121743.3 | 394.0325 | tri/oligope | Y |
| 112 | LMC2 | 415.2123 | 1 | 36.46694 | 291695.3 | 414.205 | tri/oligope | Y |
| 134 | LMC2 | 463.0273 | 1 | 1.59865 | 151387.5 | 462.02 | tri/oligope | Y |
| 141 | LMC2 | 479.3109 | 1 | 37.15848 | 157109.3 | 478.3036 | tri/oligope | Y |
| 156 | LMC2 | 515.292 | 1 | 37.08083 | 114310.3 | 514.2847 | tri/oligope | Y |
| 161 | LMC2 | 264.1751 | 2 | 35.26117 | 112157.7 | 526.3357 | tri/oligope | Y |
| 165 | LMC2 | 531.0146 | 1 | 1.588292 | 179670.1 | 530.0073 | tri/oligope | Y |
| 166 | LMC2 | 531.3877 | 1 | 37.83429 | 100276.1 | 530.3804 | tri/oligope | Y |
| 174 | LMC2 | 274.1681 | 2 | 35.92845 | 113243.4 | 546.3216 | tri/oligope | Y |
| 194 | LMC2 | 599.0024 | 1 | 1.588292 | 162989.7 | 597.9952 | tri/oligope | Y |
| 218 | LMC2 | 666.99 | 1 | 1.588292 | 154789 | 665.9827 | oligopepti | Y |
| 240 | LMC2 | 734.9775 | 1 | 1.588292 | 130103.4 | 733.9702 | oligopepti | Y |
| 296 | LMC2 | 974.8153 | 1 | 1.59865 | 133878.8 | 973.808 | oligopepti | Y |
| 297 | LMC2 | 990.7896 | 1 | 1.59865 | 138053.7 | 989.7823 | oligopepti | Y |
| 338 | LMC2 | 223.1084 | 1 | 10.40951 | 106602.9 | 222.1011 | dipeptide | N |
| 339 | LMC2 | 226.9526 | 1 | 1.650042 | 195448.4 | 225.9453 | di/tripepti | N |
| 340 | LMC2 | 229.1554 | 1 | 11.88037 | 238370.6 | 228.1482 | di/tripepti | N |
| 341 | LMC2 | 235.1654 | 1 | 1.7832 | 368201.7 | 234.1581 | di/tripepti | N |
| 342 | LMC2 | 252.963 | 1 | 8.379542 | 146697.2 | 251.9558 | di/tripepti | N |
| 343 | LMC2 | 261.1454 | 1 | 10.77103 | 140602.1 | 260.1381 | di/tripepti | N |
| 344 | LMC2 | 263.14 | 1 | 16.82774 | 194641.6 | 262.1328 | di/tripepti | N |
| 345 | LMC2 | 268.1408 | 1 | 2.189583 | 100466.3 | 267.1335 | di/tripepti | N |
| 346 | LMC2 | 269.0884 | 1 | 2.189583 | 293937.8 | 268.0811 | di/tripepti | N |
| 347 | LMC2 | 274.1299 | 1 | 2.189583 | 184041.3 | 273.1227 | di/tripepti | N |
| 348 | LMC2 | 279.1348 | 1 | 9.432442 | 301840.2 | 278.1276 | di/tripepti | N |
| 349 | LMC2 | 290.1351 | 1 | 2.028083 | 117307.3 | 289.1278 | di/tripepti | N |

| | | | | | | | | |
|---|---|---|---|---|---|---|---|---|
| 350 | LMC2 | 290.1715 | 1 | 2.189583 | 149052 | 289.1643 | di/tripepti | N |
| 351 | LMC2 | 294.9405 | 1 | 1.650042 | 120607.9 | 293.9333 | di/tripepti | N |
| 352 | LMC2 | 295.0609 | 1 | 2.154708 | 118228.2 | 294.0536 | di/tripepti | N |
| 353 | LMC2 | 311.1242 | 1 | 2.189583 | 162549.5 | 310.117 | tri/oligope | N |
| 354 | LMC2 | 318.1203 | 1 | 2.189583 | 102168.1 | 317.113 | tri/oligope | N |
| 355 | LMC2 | 348.1772 | 1 | 2.189583 | 181931.6 | 347.17 | tri/oligope | N |
| 356 | LMC2 | 361.1725 | 1 | 2.178042 | 147074 | 360.1652 | tri/oligope | N |
| 357 | LMC2 | 362.193 | 1 | 2.272383 | 109095.4 | 361.1858 | tri/oligope | N |
| 358 | LMC2 | 406.1728 | 1 | 2.201183 | 128053 | 405.1656 | tri/oligope | N |
| 359 | LMC2 | 433.2464 | 1 | 22.54095 | 223442.7 | 432.2391 | tri/oligope | N |
| 360 | LMC2 | 436.3433 | 1 | 35.9459 | 450263.6 | 435.336 | tri/oligope | N |
| 361 | LMC2 | 436.3437 | 1 | 0.020983 | 248379.4 | 435.3364 | tri/oligope | N |
| 362 | LMC2 | 450.1626 | 1 | 2.178042 | 113922.9 | 449.1553 | tri/oligope | N |
| 363 | LMC2 | 498.902 | 1 | 1.588292 | 243353.3 | 497.8947 | tri/oligope | N |
| 364 | LMC2 | 498.9028 | 1 | 1.650042 | 126213.5 | 497.8955 | tri/oligope | N |
| 365 | LMC2 | 598.838 | 1 | 1.619058 | 108531.4 | 597.8308 | tri/oligope | N |

| ID | Protein | mz | charge | retentiont | raw.abund | mass | type | Match |
|---|---|---|---|---|---|---|---|---|
| 2 | TPP1 | 231.1712 | 1 | 4.993583 | 1087549 | 230.1639 | di/tripepti | Y |
| 3 | TPP1 | 231.1712 | 1 | 9.508383 | 1138185 | 230.1639 | di/tripepti | Y |
| 4 | TPP1 | 231.1712 | 1 | 11.98594 | 860501.2 | 230.1639 | di/tripepti | Y |
| 5 | TPP1 | 233.1502 | 1 | 2.194608 | 290728.5 | 232.1429 | di/tripepti | Y |
| 6 | TPP1 | 233.1504 | 1 | 2.719742 | 536815.3 | 232.1431 | di/tripepti | Y |
| 8 | TPP1 | 239.1034 | 1 | 2.397617 | 216850.3 | 238.0961 | di/tripepti | Y |
| 11 | TPP1 | 245.1868 | 1 | 16.41001 | 1196334 | 244.1795 | di/tripepti | Y |
| 12 | TPP1 | 245.1869 | 1 | 15.86598 | 221307.8 | 244.1796 | di/tripepti | Y |
| 13 | TPP1 | 245.1869 | 1 | 14.49528 | 1021074 | 244.1796 | di/tripepti | Y |
| 14 | TPP1 | 245.1868 | 1 | 17.66004 | 1634066 | 244.1796 | di/tripepti | Y |
| 15 | TPP1 | 246.1455 | 1 | 2.183042 | 620093.7 | 245.1382 | di/tripepti | Y |
| 16 | TPP1 | 246.1456 | 1 | 2.606783 | 533341.4 | 245.1384 | di/tripepti | Y |
| 20 | TPP1 | 247.1296 | 1 | 2.183042 | 312523.4 | 246.1223 | di/tripepti | Y |
| 21 | TPP1 | 247.1298 | 1 | 3.0391 | 322793.5 | 246.1225 | di/tripepti | Y |
| 22 | TPP1 | 247.1297 | 1 | 4.590683 | 499708.3 | 246.1224 | di/tripepti | Y |
| 23 | TPP1 | 249.1285 | 1 | 2.730717 | 239298.4 | 248.1213 | di/tripepti | Y |
| 27 | TPP1 | 253.119 | 1 | 2.27365 | 249564.1 | 252.1117 | di/tripepti | Y |
| 27 | TPP1 | 253.1191 | 1 | 2.455017 | 121967.1 | 252.1118 | di/tripepti | Y |
| 28 | TPP1 | 253.1191 | 1 | 9.2369 | 309478.6 | 252.1119 | di/tripepti | Y |
| 30 | TPP1 | 260.1613 | 1 | 2.194608 | 665058.6 | 259.154 | di/tripepti | Y |
| 31 | TPP1 | 260.1615 | 1 | 2.618408 | 141134.3 | 259.1542 | di/tripepti | Y |
| 33 | TPP1 | 260.1976 | 1 | 2.194608 | 314673.4 | 259.1904 | di/tripepti | Y |
| 33 | TPP1 | 260.1976 | 1 | 1.860925 | 130507.8 | 259.1904 | di/tripepti | Y |
| 34 | TPP1 | 261.1453 | 1 | 2.194608 | 225404.4 | 260.138 | di/tripepti | Y |
| 35 | TPP1 | 261.1456 | 1 | 3.151317 | 129631.5 | 260.1383 | di/tripepti | Y |
| 36 | TPP1 | 261.1455 | 1 | 5.290575 | 547601.2 | 260.1382 | di/tripepti | Y |
| 38 | TPP1 | 263.1434 | 1 | 11.43265 | 165336.1 | 262.1362 | di/tripepti | Y |
| 40 | TPP1 | 265.1557 | 1 | 16.25203 | 920959.6 | 264.1484 | di/tripepti | Y |
| 41 | TPP1 | 267.1349 | 1 | 2.374933 | 183594.6 | 266.1276 | di/tripepti | Y |
| 42 | TPP1 | 269.1616 | 1 | 2.183042 | 334475.7 | 268.1543 | di/tripepti | Y |
| 45 | TPP1 | 281.1142 | 1 | 2.5323 | 284112.9 | 280.1069 | di/tripepti | Y |
| 46 | TPP1 | 281.1506 | 1 | 6.639933 | 402843.7 | 280.1434 | di/tripepti | Y |
| 47 | TPP1 | 288.1929 | 1 | 15.91368 | 175040.6 | 287.1856 | di/tripepti | Y |
| 50 | TPP1 | 294.1456 | 1 | 2.307033 | 236906.7 | 293.1383 | di/tripepti | Y |
| 51 | TPP1 | 295.1301 | 1 | 3.078567 | 249463.5 | 294.1228 | di/tripepti | Y |
| 52 | TPP1 | 295.1662 | 1 | 16.41001 | 629202.1 | 294.1589 | di/tripepti | Y |
| 55 | TPP1 | 302.2084 | 1 | 16.39781 | 114550.2 | 301.2012 | tri/oligope | Y |
| 57 | TPP1 | 302.2087 | 1 | 19.85245 | 332471.6 | 301.2014 | tri/oligope | Y |
| 58 | TPP1 | 303.1673 | 1 | 2.194608 | 144609.3 | 302.1601 | tri/oligope | Y |
| 59 | TPP1 | 304.1513 | 1 | 2.194608 | 159978.2 | 303.144 | tri/oligope | Y |
| 64 | TPP1 | 318.1671 | 1 | 2.183042 | 920231.2 | 317.1598 | tri/oligope | Y |
| 65 | TPP1 | 318.1673 | 1 | 8.99935 | 141819.4 | 317.16 | tri/oligope | Y |
| 66 | TPP1 | 318.1827 | 1 | 22.88427 | 206560.8 | 317.1754 | tri/oligope | Y |
| 66 | TPP1 | 318.1828 | 1 | 22.561 | 521525.6 | 317.1755 | tri/oligope | Y |
| 73 | TPP1 | 331.1663 | 1 | 2.78065 | 295841.2 | 330.159 | tri/oligope | Y |
| 75 | TPP1 | 332.1829 | 1 | 2.5323 | 114262 | 331.1756 | tri/oligope | Y |
| 80 | TPP1 | 334.1619 | 1 | 2.194608 | 794391.5 | 333.1547 | tri/oligope | Y |
| 89 | TPP1 | 358.2715 | 1 | 24.00949 | 619761.4 | 357.2642 | tri/oligope | Y |
| 90 | TPP1 | 359.2301 | 1 | 2.229358 | 121687.7 | 358.2228 | tri/oligope | Y |

| | | | | | | | | |
|---|---|---|---|---|---|---|---|---|
| 94 | TPP1 | 363.1563 | 1 | 2.183042 | 210459.1 | 362.149 | tri/oligope | Y |
| 98 | TPP1 | 375.2248 | 1 | 2.183042 | 178762.3 | 374.2175 | tri/oligope | Y |
| 100 | TPP1 | 376.1729 | 1 | 2.420517 | 121769.4 | 375.1656 | tri/oligope | Y |
| 101 | TPP1 | 376.2255 | 1 | 21.16854 | 240797.6 | 375.2182 | tri/oligope | Y |
| 106 | TPP1 | 395.0399 | 1 | 1.596875 | 176177.1 | 394.0327 | tri/oligope | Y |
| 112 | TPP1 | 415.2127 | 1 | 36.47063 | 125494.4 | 414.2054 | tri/oligope | Y |
| 144 | TPP1 | 488.2524 | 1 | 19.12113 | 236100.8 | 487.2451 | tri/oligope | Y |
| 151 | TPP1 | 500.2889 | 1 | 25.19041 | 167677.9 | 499.2817 | tri/oligope | Y |
| 159 | TPP1 | 520.2785 | 1 | 21.16854 | 142919.4 | 519.2712 | tri/oligope | Y |
| 165 | TPP1 | 531.0149 | 1 | 1.596875 | 293853.4 | 530.0076 | tri/oligope | Y |
| 194 | TPP1 | 599.0028 | 1 | 1.596875 | 264403.2 | 597.9955 | tri/oligope | Y |
| 218 | TPP1 | 666.9904 | 1 | 1.596875 | 266644.2 | 665.9831 | oligopepti | Y |
| 258 | TPP1 | 802.9655 | 1 | 1.596875 | 200713.7 | 801.9582 | oligopepti | Y |
| 276 | TPP1 | 870.953 | 1 | 1.585625 | 141502.7 | 869.9457 | oligopepti | Y |
| 300 | TPP1 | 514.9588 | 2 | 1.607775 | 113160.6 | 1027.903 | oligopepti | Y |
| 307 | TPP1 | 548.9524 | 2 | 1.596875 | 116258.5 | 1095.89 | oligopepti | Y |
| 313 | TPP1 | 582.9463 | 2 | 1.596875 | 133610.7 | 1163.878 | oligopepti | Y |
| 314 | TPP1 | 590.9336 | 2 | 1.596875 | 104025.5 | 1179.853 | oligopepti | Y |
| 316 | TPP1 | 616.9405 | 2 | 1.596875 | 127818.5 | 1231.866 | oligopepti | Y |
| 318 | TPP1 | 624.9273 | 2 | 1.607775 | 116638.9 | 1247.84 | oligopepti | Y |
| 321 | TPP1 | 650.9336 | 2 | 1.607775 | 115941.6 | 1299.853 | oligopepti | Y |
| 322 | TPP1 | 658.9208 | 2 | 1.596875 | 114580.1 | 1315.827 | oligopepti | Y |
| 329 | TPP1 | 692.9146 | 2 | 1.596875 | 114866.5 | 1383.815 | oligopepti | Y |
| 331 | TPP1 | 726.9088 | 2 | 1.607775 | 100252.8 | 1451.803 | oligopepti | Y |
| 333 | TPP1 | 760.9023 | 2 | 1.607775 | 109802.8 | 1519.79 | oligopepti | Y |
| 335 | TPP1 | 794.8962 | 2 | 1.618825 | 105588.5 | 1587.778 | oligopepti | Y |
| 338 | TPP1 | 223.1085 | 1 | 4.728142 | 281913 | 222.1012 | dipeptide | N |
| 339 | TPP1 | 223.1085 | 1 | 10.36156 | 112491.1 | 222.1012 | dipeptide | N |
| 340 | TPP1 | 231.1712 | 1 | 8.785917 | 298049.9 | 230.1639 | di/tripepti | N |
| 341 | TPP1 | 232.1298 | 1 | 2.1716 | 219129.1 | 231.1225 | di/tripepti | N |
| 342 | TPP1 | 232.1298 | 1 | 1.902508 | 110235.1 | 231.1225 | di/tripepti | N |
| 343 | TPP1 | 233.1139 | 1 | 2.194608 | 214165.9 | 232.1066 | di/tripepti | N |
| 344 | TPP1 | 233.1505 | 1 | 3.9329 | 316939.6 | 232.1432 | di/tripepti | N |
| 345 | TPP1 | 237.1242 | 1 | 4.278833 | 885188.7 | 236.1169 | di/tripepti | N |
| 346 | TPP1 | 237.1242 | 1 | 11.13268 | 141139.1 | 236.1169 | di/tripepti | N |
| 347 | TPP1 | 251.1066 | 1 | 2.183042 | 183472 | 250.0994 | di/tripepti | N |
| 348 | TPP1 | 260.1615 | 1 | 6.117333 | 400916.2 | 259.1542 | di/tripepti | N |
| 349 | TPP1 | 262.1196 | 1 | 14.46567 | 103324.5 | 261.1123 | di/tripepti | N |
| 350 | TPP1 | 263.14 | 1 | 14.41424 | 170894.9 | 262.1327 | di/tripepti | N |
| 351 | TPP1 | 263.1434 | 1 | 13.05917 | 256073.9 | 262.1361 | di/tripepti | N |
| 352 | TPP1 | 265.1557 | 1 | 14.71689 | 645284.2 | 264.1484 | di/tripepti | N |
| 353 | TPP1 | 269.1139 | 1 | 2.307033 | 156469.8 | 268.1067 | di/tripepti | N |
| 354 | TPP1 | 274.1771 | 1 | 10.33156 | 506105.6 | 273.1698 | di/tripepti | N |
| 355 | TPP1 | 276.1563 | 1 | 2.194608 | 433463.1 | 275.149 | di/tripepti | N |
| 356 | TPP1 | 277.2169 | 1 | 36.75047 | 537316.4 | 276.2096 | di/tripepti | N |
| 357 | TPP1 | 278.1177 | 1 | 2.194608 | 241226.9 | 277.1104 | di/tripepti | N |
| 358 | TPP1 | 279.1714 | 1 | 21.01918 | 519795 | 278.1641 | di/tripepti | N |
| 359 | TPP1 | 279.1714 | 1 | 19.89113 | 1210250 | 278.1641 | di/tripepti | N |
| 360 | TPP1 | 279.1715 | 1 | 18.96737 | 140174 | 278.1642 | di/tripepti | N |
| 361 | TPP1 | 279.2326 | 1 | 37.01238 | 988600 | 278.2253 | di/tripepti | N |

| | | | | | | | | |
|---|---|---|---|---|---|---|---|---|
| 362 | TPP1 | 281.1143 | 1 | 11.51086 | 181335.1 | 280.107 | di/tripepti | N |
| 363 | TPP1 | 281.1506 | 1 | 5.606542 | 141636.6 | 280.1434 | di/tripepti | N |
| 364 | TPP1 | 286.1772 | 1 | 13.43262 | 1104791 | 285.1699 | di/tripepti | N |
| 365 | TPP1 | 290.1354 | 1 | 2.183042 | 149517.3 | 289.1282 | di/tripepti | N |
| 366 | TPP1 | 290.1719 | 1 | 2.194608 | 770765.1 | 289.1646 | di/tripepti | N |
| 367 | TPP1 | 292.1302 | 1 | 16.41001 | 109017.2 | 291.123 | di/tripepti | N |
| 368 | TPP1 | 294.9403 | 1 | 1.675225 | 220749 | 293.933 | di/tripepti | N |
| 369 | TPP1 | 295.13 | 1 | 12.58608 | 915728.5 | 294.1228 | di/tripepti | N |
| 370 | TPP1 | 295.1664 | 1 | 14.56998 | 286538.5 | 294.1592 | di/tripepti | N |
| 371 | TPP1 | 295.1665 | 1 | 13.35489 | 164127.1 | 294.1592 | di/tripepti | N |
| 372 | TPP1 | 295.2277 | 1 | 37.377 | 527949 | 294.2204 | di/tripepti | N |
| 373 | TPP1 | 297.1092 | 1 | 2.636492 | 317476.2 | 296.1019 | di/tripepti | N |
| 374 | TPP1 | 297.2423 | 1 | 37.76648 | 118587 | 296.235 | di/tripepti | N |
| 375 | TPP1 | 304.1336 | 1 | 2.606783 | 335980.6 | 303.1264 | tri/oligope | N |
| 376 | TPP1 | 304.1667 | 1 | 19.90886 | 538756.4 | 303.1594 | tri/oligope | N |
| 377 | TPP1 | 304.1878 | 1 | 2.576733 | 173810.5 | 303.1805 | tri/oligope | N |
| 378 | TPP1 | 306.1461 | 1 | 10.63 | 113663.2 | 305.1388 | tri/oligope | N |
| 379 | TPP1 | 306.1669 | 1 | 2.194608 | 291205.6 | 305.1597 | tri/oligope | N |
| 380 | TPP1 | 306.167 | 1 | 2.618408 | 121176.2 | 305.1597 | tri/oligope | N |
| 381 | TPP1 | 313.1767 | 1 | 2.183042 | 107032.1 | 312.1694 | tri/oligope | N |
| 382 | TPP1 | 316.1878 | 1 | 2.32975 | 107982.3 | 315.1806 | tri/oligope | N |
| 383 | TPP1 | 316.2245 | 1 | 14.89203 | 165784.8 | 315.2172 | tri/oligope | N |
| 384 | TPP1 | 317.2096 | 1 | 37.377 | 240170.9 | 316.2023 | tri/oligope | N |
| 385 | TPP1 | 318.2034 | 1 | 2.194608 | 213587.4 | 317.1961 | tri/oligope | N |
| 386 | TPP1 | 318.2037 | 1 | 12.28745 | 669389 | 317.1964 | tri/oligope | N |
| 387 | TPP1 | 318.2037 | 1 | 14.56098 | 149961.9 | 317.1964 | tri/oligope | N |
| 388 | TPP1 | 319.2254 | 1 | 37.01238 | 103592.2 | 318.2181 | tri/oligope | N |
| 389 | TPP1 | 320.1464 | 1 | 2.116342 | 170700.5 | 319.1391 | tri/oligope | N |
| 390 | TPP1 | 322.144 | 1 | 2.295758 | 320150.5 | 321.1368 | tri/oligope | N |
| 391 | TPP1 | 324.1566 | 1 | 2.509608 | 151630.1 | 323.1494 | tri/oligope | N |
| 392 | TPP1 | 328.2245 | 1 | 15.41343 | 219502.2 | 327.2172 | tri/oligope | N |
| 393 | TPP1 | 331.1988 | 1 | 2.194608 | 259363.8 | 330.1915 | tri/oligope | N |
| 394 | TPP1 | 331.2349 | 1 | 2.194608 | 109864 | 330.2276 | tri/oligope | N |
| 395 | TPP1 | 332.2193 | 1 | 4.9847 | 300802.9 | 331.212 | tri/oligope | N |
| 396 | TPP1 | 332.2193 | 1 | 9.5437 | 313352.6 | 331.212 | tri/oligope | N |
| 397 | TPP1 | 332.2194 | 1 | 19.42713 | 108549.1 | 331.2121 | tri/oligope | N |
| 398 | TPP1 | 333.1571 | 1 | 17.05924 | 453391.8 | 332.1498 | tri/oligope | N |
| 399 | TPP1 | 333.1779 | 1 | 2.194608 | 280643.1 | 332.1706 | tri/oligope | N |
| 400 | TPP1 | 336.1236 | 1 | 2.262583 | 135855 | 335.1163 | tri/oligope | N |
| 401 | TPP1 | 336.139 | 1 | 21.01918 | 214250.6 | 335.1317 | tri/oligope | N |
| 402 | TPP1 | 336.1933 | 1 | 23.03313 | 177137.8 | 335.1861 | tri/oligope | N |
| 403 | TPP1 | 340.2858 | 1 | 37.377 | 142454.5 | 339.2785 | tri/oligope | N |
| 404 | TPP1 | 342.2402 | 1 | 19.86153 | 225628.4 | 341.2329 | tri/oligope | N |
| 405 | TPP1 | 342.2402 | 1 | 20.73378 | 1035432 | 341.233 | tri/oligope | N |
| 406 | TPP1 | 343.1991 | 1 | 13.31236 | 889164.8 | 342.1918 | tri/oligope | N |
| 407 | TPP1 | 345.2147 | 1 | 15.7986 | 117227.4 | 344.2075 | tri/oligope | N |
| 408 | TPP1 | 346.2352 | 1 | 19.17043 | 396054.6 | 345.2279 | tri/oligope | N |
| 409 | TPP1 | 347.1573 | 1 | 2.194608 | 219918.5 | 346.15 | tri/oligope | N |
| 410 | TPP1 | 348.1414 | 1 | 2.116342 | 267898.4 | 347.1342 | tri/oligope | N |
| 411 | TPP1 | 348.1778 | 1 | 2.262583 | 609053.7 | 347.1705 | tri/oligope | N |

| | | | | | | | | |
|---|---|---|---|---|---|---|---|---|
| 412 | TPP1 | 348.178 | 1 | 9.995033 | 281076.6 | 347.1708 | tri/oligope | N |
| 413 | TPP1 | 354.1307 | 1 | 2.194608 | 218057.1 | 353.1234 | tri/oligope | N |
| 414 | TPP1 | 358.2962 | 1 | 36.42645 | 160834.5 | 357.2889 | tri/oligope | N |
| 415 | TPP1 | 359.1573 | 1 | 2.194608 | 152782.7 | 358.15 | tri/oligope | N |
| 416 | TPP1 | 359.1934 | 1 | 2.183042 | 293574.8 | 358.1861 | tri/oligope | N |
| 417 | TPP1 | 359.2303 | 1 | 16.81008 | 115106.2 | 358.223 | tri/oligope | N |
| 418 | TPP1 | 359.2663 | 1 | 2.2403 | 129574.8 | 358.259 | tri/oligope | N |
| 419 | TPP1 | 360.2146 | 1 | 20.4456 | 112145.6 | 359.2073 | tri/oligope | N |
| 420 | TPP1 | 361.173 | 1 | 2.183042 | 197819.5 | 360.1657 | tri/oligope | N |
| 421 | TPP1 | 361.1732 | 1 | 2.819 | 128837.8 | 360.166 | tri/oligope | N |
| 422 | TPP1 | 362.2088 | 1 | 21.3447 | 145341.1 | 361.2016 | tri/oligope | N |
| 423 | TPP1 | 363.1886 | 1 | 2.5545 | 102944 | 362.1813 | tri/oligope | N |
| 424 | TPP1 | 366.134 | 1 | 2.194608 | 225905.9 | 365.1268 | tri/oligope | N |
| 425 | TPP1 | 368.1468 | 1 | 8.852467 | 108384.5 | 367.1395 | tri/oligope | N |
| 426 | TPP1 | 371.2304 | 1 | 14.11968 | 666059.9 | 370.2231 | tri/oligope | N |
| 427 | TPP1 | 373.2092 | 1 | 2.576733 | 2070175 | 372.2019 | tri/oligope | N |
| 428 | TPP1 | 373.2459 | 1 | 16.47554 | 992004.9 | 372.2386 | tri/oligope | N |
| 429 | TPP1 | 373.2462 | 1 | 18.80608 | 124517.1 | 372.2389 | tri/oligope | N |
| 430 | TPP1 | 374.2045 | 1 | 2.183042 | 581604.1 | 373.1972 | tri/oligope | N |
| 431 | TPP1 | 375.189 | 1 | 8.367283 | 115244.1 | 374.1817 | tri/oligope | N |
| 432 | TPP1 | 376.2252 | 1 | 23.89678 | 353426.4 | 375.2179 | tri/oligope | N |
| 433 | TPP1 | 377.2042 | 1 | 2.194608 | 184855 | 376.1969 | tri/oligope | N |
| 434 | TPP1 | 389.2048 | 1 | 10.81154 | 276222 | 388.1975 | tri/oligope | N |
| 435 | TPP1 | 391.2198 | 1 | 2.21825 | 107812.6 | 390.2125 | tri/oligope | N |
| 436 | TPP1 | 401.2412 | 1 | 15.06663 | 158445.6 | 400.2339 | tri/oligope | N |
| 437 | TPP1 | 404.2152 | 1 | 2.475942 | 424661.2 | 403.2079 | tri/oligope | N |
| 438 | TPP1 | 405.1992 | 1 | 2.206367 | 131816.3 | 404.192 | tri/oligope | N |
| 439 | TPP1 | 408.1449 | 1 | 2.5323 | 112779.5 | 407.1376 | tri/oligope | N |
| 440 | TPP1 | 408.2145 | 1 | 17.54296 | 152335.9 | 407.2073 | tri/oligope | N |
| 441 | TPP1 | 419.2148 | 1 | 2.27365 | 1126873 | 418.2075 | tri/oligope | N |
| 442 | TPP1 | 419.2304 | 1 | 24.53998 | 1536043 | 418.2232 | tri/oligope | N |
| 443 | TPP1 | 423.2255 | 1 | 15.8946 | 179781.4 | 422.2182 | tri/oligope | N |
| 444 | TPP1 | 429.2725 | 1 | 23.96479 | 246307.3 | 428.2652 | tri/oligope | N |
| 445 | TPP1 | 431.1942 | 1 | 19.96136 | 199353.9 | 430.1869 | tri/oligope | N |
| 446 | TPP1 | 431.2517 | 1 | 17.46008 | 150160.9 | 430.2445 | tri/oligope | N |
| 447 | TPP1 | 432.2463 | 1 | 2.183042 | 147062.7 | 431.239 | tri/oligope | N |
| 448 | TPP1 | 433.2672 | 1 | 12.17632 | 926242.1 | 432.26 | tri/oligope | N |
| 449 | TPP1 | 434.2263 | 1 | 10.68503 | 404689.4 | 433.219 | tri/oligope | N |
| 450 | TPP1 | 443.2515 | 1 | 19.61331 | 739667.1 | 442.2443 | tri/oligope | N |
| 451 | TPP1 | 444.2467 | 1 | 2.5323 | 724110.8 | 443.2395 | tri/oligope | N |
| 452 | TPP1 | 222.627 | 2 | 2.5545 | 110728.3 | 443.2395 | tri/oligope | N |
| 453 | TPP1 | 444.247 | 1 | 13.44047 | 493649.5 | 443.2397 | tri/oligope | N |
| 454 | TPP1 | 445.2308 | 1 | 16.27382 | 431831.6 | 444.2235 | tri/oligope | N |
| 455 | TPP1 | 224.6321 | 2 | 2.5323 | 280258.3 | 447.2496 | tri/oligope | N |
| 456 | TPP1 | 450.2001 | 1 | 13.51618 | 656103.1 | 449.1928 | tri/oligope | N |
| 457 | TPP1 | 450.2204 | 1 | 2.194608 | 232753 | 449.2131 | tri/oligope | N |
| 458 | TPP1 | 456.2835 | 1 | 20.99268 | 317198.1 | 455.2762 | tri/oligope | N |
| 459 | TPP1 | 457.2423 | 1 | 14.17213 | 124050.4 | 456.235 | tri/oligope | N |
| 460 | TPP1 | 458.2257 | 1 | 2.194608 | 137205.2 | 457.2184 | tri/oligope | N |
| 461 | TPP1 | 458.2263 | 1 | 14.87784 | 235457.1 | 457.219 | tri/oligope | N |

| | | | | | | | | |
|---|---|---|---|---|---|---|---|---|
| 462 | TPP1 | 460.2417 | 1 | 2.645608 | 119808 | 459.2344 | tri/oligope | N |
| 463 | TPP1 | 461.2621 | 1 | 15.59184 | 187757.4 | 460.2549 | tri/oligope | N |
| 464 | TPP1 | 467.1437 | 1 | 23.24796 | 410426.1 | 466.1364 | tri/oligope | N |
| 465 | TPP1 | 470.2987 | 1 | 21.43569 | 2544861 | 469.2914 | tri/oligope | N |
| 466 | TPP1 | 236.1324 | 2 | 2.082508 | 128431 | 470.2503 | tri/oligope | N |
| 467 | TPP1 | 472.2569 | 1 | 27.18286 | 1135490 | 471.2496 | tri/oligope | N |
| 468 | TPP1 | 473.2735 | 1 | 13.07853 | 134950.7 | 472.2662 | tri/oligope | N |
| 469 | TPP1 | 473.2735 | 1 | 14.8516 | 385024.8 | 472.2663 | tri/oligope | N |
| 470 | TPP1 | 474.2316 | 1 | 2.183042 | 396415.1 | 473.2243 | tri/oligope | N |
| 471 | TPP1 | 475.2778 | 1 | 17.69141 | 256807 | 474.2705 | tri/oligope | N |
| 472 | TPP1 | 475.2927 | 1 | 26.25843 | 442448 | 474.2854 | tri/oligope | N |
| 473 | TPP1 | 238.64 | 2 | 2.262583 | 276071.2 | 475.2655 | tri/oligope | N |
| 474 | TPP1 | 478.2518 | 1 | 2.194608 | 105857.4 | 477.2446 | tri/oligope | N |
| 475 | TPP1 | 481.231 | 1 | 24.36167 | 303312.7 | 480.2237 | tri/oligope | N |
| 476 | TPP1 | 483.2464 | 1 | 12.64023 | 161178.3 | 482.2392 | tri/oligope | N |
| 477 | TPP1 | 486.294 | 1 | 18.44501 | 401282.4 | 485.2867 | tri/oligope | N |
| 478 | TPP1 | 487.289 | 1 | 16.27382 | 166294.7 | 486.2817 | tri/oligope | N |
| 479 | TPP1 | 487.2894 | 1 | 16.56902 | 119307.2 | 486.2821 | tri/oligope | N |
| 480 | TPP1 | 487.3142 | 1 | 21.45246 | 145912.5 | 486.307 | tri/oligope | N |
| 481 | TPP1 | 488.2734 | 1 | 15.25283 | 154767.3 | 487.2661 | tri/oligope | N |
| 482 | TPP1 | 489.3091 | 1 | 27.88488 | 1169345 | 488.3018 | tri/oligope | N |
| 483 | TPP1 | 489.358 | 1 | 36.4528 | 100684.9 | 488.3507 | tri/oligope | N |
| 484 | TPP1 | 491.2472 | 1 | 2.386317 | 202749.8 | 490.24 | tri/oligope | N |
| 485 | TPP1 | 497.1898 | 1 | 13.61781 | 276881.4 | 496.1825 | tri/oligope | N |
| 486 | TPP1 | 502.3253 | 1 | 21.57278 | 461516.7 | 501.318 | tri/oligope | N |
| 487 | TPP1 | 503.2845 | 1 | 15.18874 | 101914.8 | 502.2772 | tri/oligope | N |
| 488 | TPP1 | 504.2424 | 1 | 2.194608 | 141417.8 | 503.2352 | tri/oligope | N |
| 489 | TPP1 | 504.2681 | 1 | 15.51249 | 439063 | 503.2608 | tri/oligope | N |
| 490 | TPP1 | 506.1546 | 1 | 25.23488 | 326865.5 | 505.1473 | tri/oligope | N |
| 491 | TPP1 | 506.2296 | 1 | 20.4456 | 173914.1 | 505.2223 | tri/oligope | N |
| 492 | TPP1 | 506.266 | 1 | 17.27622 | 457793 | 505.2588 | tri/oligope | N |
| 493 | TPP1 | 517.2627 | 1 | 2.183042 | 119857.6 | 516.2554 | tri/oligope | N |
| 494 | TPP1 | 519.2421 | 1 | 2.183042 | 177645.2 | 518.2348 | tri/oligope | N |
| 495 | TPP1 | 522.2576 | 1 | 17.89618 | 368137.7 | 521.2503 | tri/oligope | N |
| 496 | TPP1 | 262.6643 | 2 | 13.98698 | 100722 | 523.314 | tri/oligope | N |
| 497 | TPP1 | 531.2792 | 1 | 3.180542 | 169647.2 | 530.2719 | tri/oligope | N |
| 498 | TPP1 | 535.274 | 1 | 2.956392 | 191172.9 | 534.2667 | tri/oligope | N |
| 499 | TPP1 | 545.2944 | 1 | 2.465533 | 324481.3 | 544.2871 | tri/oligope | N |
| 500 | TPP1 | 547.2897 | 1 | 23.46378 | 105415.2 | 546.2825 | tri/oligope | N |
| 501 | TPP1 | 548.2207 | 1 | 2.206367 | 162265.1 | 547.2134 | tri/oligope | N |
| 502 | TPP1 | 550.256 | 1 | 18.96737 | 268721 | 549.2487 | tri/oligope | N |
| 503 | TPP1 | 556.3372 | 1 | 22.40913 | 189092.5 | 555.3299 | tri/oligope | N |
| 504 | TPP1 | 557.332 | 1 | 23.30199 | 286429.1 | 556.3247 | tri/oligope | N |
| 505 | TPP1 | 558.326 | 1 | 2.194608 | 239764.8 | 557.3187 | tri/oligope | N |
| 506 | TPP1 | 279.6667 | 2 | 2.183042 | 576478.5 | 557.3188 | tri/oligope | N |
| 507 | TPP1 | 559.2742 | 1 | 2.618408 | 147979.1 | 558.2669 | tri/oligope | N |
| 508 | TPP1 | 559.2746 | 1 | 11.78493 | 228333.5 | 558.2673 | tri/oligope | N |
| 509 | TPP1 | 560.269 | 1 | 2.194608 | 105902.1 | 559.2617 | tri/oligope | N |
| 510 | TPP1 | 561.29 | 1 | 14.92539 | 446654.5 | 560.2828 | tri/oligope | N |
| 511 | TPP1 | 574.3263 | 1 | 26.24413 | 100211.7 | 573.319 | tri/oligope | N |

| | | | | | | | | |
|---|---|---|---|---|---|---|---|---|
| 512 | TPP1 | 288.1477 | 2 | 18.86806 | 100228.6 | 574.2809 | tri/oligope | N |
| 513 | TPP1 | 289.6511 | 2 | 2.262583 | 238065.7 | 577.2877 | tri/oligope | N |
| 514 | TPP1 | 580.2288 | 1 | 28.67756 | 190994.8 | 579.2215 | tri/oligope | N |
| 515 | TPP1 | 580.2749 | 1 | 15.68346 | 134701.2 | 579.2676 | tri/oligope | N |
| 516 | TPP1 | 586.2857 | 1 | 16.66209 | 161728.2 | 585.2784 | tri/oligope | N |
| 517 | TPP1 | 588.3006 | 1 | 18.00173 | 377762.4 | 587.2933 | tri/oligope | N |
| 518 | TPP1 | 588.3011 | 1 | 17.35949 | 206899.4 | 587.2939 | tri/oligope | N |
| 519 | TPP1 | 590.3208 | 1 | 28.20753 | 912592.7 | 589.3136 | tri/oligope | N |
| 520 | TPP1 | 597.2903 | 1 | 15.48333 | 207511.5 | 596.283 | tri/oligope | N |
| 521 | TPP1 | 305.167 | 2 | 18.14001 | 123063.8 | 608.3194 | tri/oligope | N |
| 522 | TPP1 | 618.312 | 1 | 18.49333 | 280044.1 | 617.3047 | oligopepti | N |
| 523 | TPP1 | 315.1672 | 2 | 2.21825 | 345843.7 | 628.3199 | oligopepti | N |
| 524 | TPP1 | 631.2957 | 1 | 16.45283 | 333451.8 | 630.2884 | oligopepti | N |
| 525 | TPP1 | 631.3118 | 1 | 25.7057 | 140803 | 630.3045 | oligopepti | N |
| 526 | TPP1 | 632.3275 | 1 | 12.47783 | 269865.6 | 631.3202 | oligopepti | N |
| 527 | TPP1 | 634.3065 | 1 | 2.78905 | 151227.2 | 633.2992 | oligopepti | N |
| 528 | TPP1 | 321.6961 | 2 | 16.81643 | 107085.9 | 641.3777 | oligopepti | N |
| 529 | TPP1 | 322.6858 | 2 | 14.00702 | 213423.2 | 643.357 | oligopepti | N |
| 530 | TPP1 | 648.3589 | 1 | 19.6073 | 239946.4 | 647.3516 | oligopepti | N |
| 531 | TPP1 | 649.317 | 1 | 2.352442 | 206437.7 | 648.3097 | oligopepti | N |
| 532 | TPP1 | 325.1663 | 2 | 14.86041 | 295596.3 | 648.318 | oligopepti | N |
| 533 | TPP1 | 650.3176 | 1 | 21.86493 | 103292.1 | 649.3103 | oligopepti | N |
| 534 | TPP1 | 330.662 | 2 | 17.2042 | 3620718 | 659.3095 | oligopepti | N |
| 535 | TPP1 | 669.4213 | 1 | 28.10895 | 410772.9 | 668.4141 | oligopepti | N |
| 536 | TPP1 | 335.2143 | 2 | 28.10895 | 166065.1 | 668.4141 | oligopepti | N |
| 537 | TPP1 | 672.3592 | 1 | 18.38946 | 203388.3 | 671.352 | oligopepti | N |
| 538 | TPP1 | 676.3357 | 1 | 16.39781 | 199893.1 | 675.3285 | oligopepti | N |
| 539 | TPP1 | 683.4117 | 1 | 22.6982 | 213255.3 | 682.4044 | oligopepti | N |
| 540 | TPP1 | 685.3543 | 1 | 18.65493 | 110560.7 | 684.3471 | oligopepti | N |
| 541 | TPP1 | 349.2356 | 2 | 20.64037 | 215781.7 | 696.4566 | oligopepti | N |
| 542 | TPP1 | 356.6777 | 2 | 16.96288 | 162705.2 | 711.3409 | oligopepti | N |
| 543 | TPP1 | 713.3858 | 1 | 17.81028 | 115652.6 | 712.3785 | oligopepti | N |
| 544 | TPP1 | 720.288 | 1 | 30.38458 | 225304.9 | 719.2808 | oligopepti | N |
| 545 | TPP1 | 362.1647 | 2 | 2.194608 | 187589.7 | 722.3149 | oligopepti | N |
| 546 | TPP1 | 241.7791 | 3 | 2.194608 | 228797.9 | 722.3154 | oligopepti | N |
| 547 | TPP1 | 362.225 | 2 | 16.22791 | 148188.2 | 722.4355 | oligopepti | N |
| 548 | TPP1 | 733.3956 | 1 | 27.89757 | 106209.9 | 732.3883 | oligopepti | N |
| 549 | TPP1 | 373.7073 | 2 | 12.85807 | 1160834 | 745.4001 | oligopepti | N |
| 550 | TPP1 | 747.3912 | 1 | 17.34187 | 119142.5 | 746.384 | oligopepti | N |
| 551 | TPP1 | 758.3345 | 1 | 17.2042 | 151518 | 757.3273 | oligopepti | N |
| 552 | TPP1 | 761.3895 | 1 | 24.26591 | 229415.8 | 760.3822 | oligopepti | N |
| 553 | TPP1 | 381.6863 | 2 | 2.363625 | 131342 | 761.3581 | oligopepti | N |
| 554 | TPP1 | 762.4025 | 1 | 18.76222 | 108034.6 | 761.3952 | oligopepti | N |
| 555 | TPP1 | 385.7253 | 2 | 2.374933 | 237100.4 | 769.436 | oligopepti | N |
| 556 | TPP1 | 390.6871 | 2 | 11.09963 | 223551.1 | 779.3596 | oligopepti | N |
| 557 | TPP1 | 391.179 | 2 | 13.11365 | 137697.8 | 780.3433 | oligopepti | N |
| 558 | TPP1 | 395.2304 | 2 | 25.15031 | 150015.8 | 788.4462 | oligopepti | N |
| 559 | TPP1 | 799.4228 | 1 | 19.72483 | 646522.6 | 798.4156 | oligopepti | N |
| 560 | TPP1 | 400.2152 | 2 | 19.72483 | 151649.3 | 798.4158 | oligopepti | N |
| 561 | TPP1 | 400.7127 | 2 | 15.82025 | 280579.4 | 799.4108 | oligopepti | N |

| | | | | | | | | |
|---|---|---|---|---|---|---|---|---|
| 562 | TPP1 | 412.1941 | 2 | 19.73385 | 178540.3 | 822.3737 | oligopepti | N |
| 563 | TPP1 | 838.3606 | 1 | 14.50453 | 103119.5 | 837.3533 | oligopepti | N |
| 564 | TPP1 | 425.205 | 2 | 24.15548 | 229580.4 | 848.3955 | oligopepti | N |
| 565 | TPP1 | 425.205 | 2 | 23.21603 | 146244.9 | 848.3955 | oligopepti | N |
| 566 | TPP1 | 426.1697 | 2 | 12.75703 | 153879.5 | 850.3249 | oligopepti | N |
| 567 | TPP1 | 426.7467 | 2 | 17.39143 | 209550.8 | 851.4789 | oligopepti | N |
| 568 | TPP1 | 429.7128 | 2 | 20.27536 | 183846.2 | 857.411 | oligopepti | N |
| 569 | TPP1 | 433.7546 | 2 | 19.67143 | 195430.8 | 865.4947 | oligopepti | N |
| 570 | TPP1 | 435.7309 | 2 | 23.58158 | 582166 | 869.4472 | oligopepti | N |
| 571 | TPP1 | 436.2368 | 2 | 25.44717 | 103211.5 | 870.4591 | oligopepti | N |
| 572 | TPP1 | 438.2289 | 2 | 16.84127 | 161612.4 | 874.4433 | oligopepti | N |
| 573 | TPP1 | 296.1342 | 3 | 13.93757 | 229416.8 | 885.3807 | oligopepti | N |
| 574 | TPP1 | 890.3927 | 1 | 32.64786 | 132452.7 | 889.3854 | oligopepti | N |
| 575 | TPP1 | 460.6674 | 2 | 14.16565 | 119015.8 | 919.3202 | oligopepti | N |
| 576 | TPP1 | 461.6877 | 2 | 2.262583 | 110845.7 | 921.3609 | oligopepti | N |
| 577 | TPP1 | 487.7631 | 2 | 16.19805 | 154598 | 973.5116 | oligopepti | N |
| 578 | TPP1 | 488.7527 | 2 | 22.72034 | 103466.6 | 975.4908 | oligopepti | N |
| 579 | TPP1 | 515.2477 | 2 | 18.05342 | 174117.5 | 1028.481 | oligopepti | N |
| 580 | TPP1 | 578.2939 | 2 | 15.25283 | 421008.5 | 1154.573 | oligopepti | N |
| 581 | TPP1 | 596.7805 | 2 | 20.16584 | 180004 | 1191.547 | oligopepti | N |
| 582 | TPP1 | 643.8141 | 2 | 17.72069 | 196992.9 | 1285.614 | oligopepti | N |
| 583 | TPP1 | 684.9278 | 2 | 1.596875 | 122133.4 | 1367.841 | oligopepti | N |
| 584 | TPP1 | 718.9214 | 2 | 1.607775 | 110720.2 | 1435.828 | oligopepti | N |
| 585 | TPP1 | 752.9153 | 2 | 1.607775 | 105421.8 | 1503.816 | oligopepti | N |

| ID | Protein | mz | charge | retentiont | raw.abund | mass | type | Match |
|---|---|---|---|---|---|---|---|---|
| 2 | TPP2 | 231.1712 | 1 | 4.997817 | 135424.5 | 230.1639 | di/tripepti | Y |
| 3 | TPP2 | 231.1712 | 1 | 9.505967 | 386322.7 | 230.1639 | di/tripepti | Y |
| 4 | TPP2 | 231.1712 | 1 | 11.9964 | 416855 | 230.1639 | di/tripepti | Y |
| 5 | TPP2 | 233.1502 | 1 | 2.1884 | 100058.4 | 232.1429 | di/tripepti | Y |
| 11 | TPP2 | 245.1868 | 1 | 16.40949 | 500797.7 | 244.1795 | di/tripepti | Y |
| 13 | TPP2 | 245.1869 | 1 | 14.5004 | 115043.7 | 244.1796 | di/tripepti | Y |
| 14 | TPP2 | 245.1868 | 1 | 17.66532 | 391293.2 | 244.1796 | di/tripepti | Y |
| 15 | TPP2 | 246.1455 | 1 | 2.177583 | 200113.4 | 245.1382 | di/tripepti | Y |
| 20 | TPP2 | 247.1296 | 1 | 2.177583 | 191821.1 | 246.1223 | di/tripepti | Y |
| 22 | TPP2 | 247.1297 | 1 | 4.587892 | 125962 | 246.1224 | di/tripepti | Y |
| 23 | TPP2 | 249.1285 | 1 | 2.734183 | 106288.5 | 248.1213 | di/tripepti | Y |
| 30 | TPP2 | 260.1613 | 1 | 2.1884 | 289570.5 | 259.154 | di/tripepti | Y |
| 33 | TPP2 | 260.1976 | 1 | 2.1884 | 336150.9 | 259.1904 | di/tripepti | Y |
| 34 | TPP2 | 261.1453 | 1 | 2.1884 | 166961.9 | 260.138 | di/tripepti | Y |
| 36 | TPP2 | 261.1455 | 1 | 5.296858 | 213905 | 260.1382 | di/tripepti | Y |
| 40 | TPP2 | 265.1557 | 1 | 16.26598 | 766152.8 | 264.1484 | di/tripepti | Y |
| 45 | TPP2 | 281.1142 | 1 | 2.536775 | 128186.6 | 280.1069 | di/tripepti | Y |
| 46 | TPP2 | 281.1506 | 1 | 6.647933 | 153884.2 | 280.1434 | di/tripepti | Y |
| 51 | TPP2 | 295.1301 | 1 | 3.0816 | 184850.2 | 294.1228 | di/tripepti | Y |
| 52 | TPP2 | 295.1662 | 1 | 16.40949 | 254863.8 | 294.1589 | di/tripepti | Y |
| 55 | TPP2 | 302.2086 | 1 | 16.80439 | 238821.2 | 301.2014 | tri/oligope | Y |
| 55 | TPP2 | 302.2087 | 1 | 16.0749 | 141453.4 | 301.2014 | tri/oligope | Y |
| 57 | TPP2 | 302.2087 | 1 | 19.8542 | 134713.9 | 301.2014 | tri/oligope | Y |
| 64 | TPP2 | 318.1671 | 1 | 2.177583 | 300332 | 317.1598 | tri/oligope | Y |
| 65 | TPP2 | 318.1673 | 1 | 9.003108 | 154665.8 | 317.16 | tri/oligope | Y |
| 66 | TPP2 | 318.1828 | 1 | 22.55677 | 153273.6 | 317.1755 | tri/oligope | Y |
| 73 | TPP2 | 331.1663 | 1 | 2.787617 | 218811 | 330.159 | tri/oligope | Y |
| 80 | TPP2 | 334.1619 | 1 | 2.1884 | 921797.6 | 333.1547 | tri/oligope | Y |
| 85 | TPP2 | 344.2559 | 1 | 21.35241 | 115673.4 | 343.2486 | tri/oligope | Y |
| 89 | TPP2 | 358.2715 | 1 | 24.00614 | 694179.2 | 357.2642 | tri/oligope | Y |
| 90 | TPP2 | 359.2301 | 1 | 2.222067 | 148663.8 | 358.2228 | tri/oligope | Y |
| 94 | TPP2 | 363.1563 | 1 | 2.177583 | 114539.5 | 362.149 | tri/oligope | Y |
| 105 | TPP2 | 390.1883 | 1 | 2.313217 | 105142.4 | 389.1811 | tri/oligope | Y |
| 112 | TPP2 | 415.2127 | 1 | 36.47674 | 171492.8 | 414.2054 | tri/oligope | Y |
| 338 | TPP2 | 237.1242 | 1 | 4.276717 | 502257.7 | 236.1169 | di/tripepti | N |
| 339 | TPP2 | 244.0936 | 1 | 1.898758 | 156335.2 | 243.0863 | di/tripepti | N |
| 340 | TPP2 | 263.1434 | 1 | 13.07238 | 142370.9 | 262.1361 | di/tripepti | N |
| 341 | TPP2 | 265.1557 | 1 | 14.72633 | 208975 | 264.1484 | di/tripepti | N |
| 342 | TPP2 | 272.9448 | 1 | 1.5495 | 118289.5 | 271.9376 | di/tripepti | N |
| 343 | TPP2 | 276.1563 | 1 | 2.1884 | 473827.7 | 275.149 | di/tripepti | N |
| 344 | TPP2 | 279.1714 | 1 | 21.02306 | 170124.9 | 278.1641 | di/tripepti | N |
| 345 | TPP2 | 279.1714 | 1 | 19.89648 | 230375 | 278.1641 | di/tripepti | N |
| 346 | TPP2 | 279.2326 | 1 | 37.00563 | 113489.6 | 278.2253 | di/tripepti | N |
| 347 | TPP2 | 280.1299 | 1 | 2.199483 | 108212.2 | 279.1227 | di/tripepti | N |
| 348 | TPP2 | 286.1772 | 1 | 13.45209 | 618239.2 | 285.1699 | di/tripepti | N |
| 349 | TPP2 | 288.2037 | 1 | 2.1884 | 105253.5 | 287.1965 | di/tripepti | N |
| 350 | TPP2 | 290.1719 | 1 | 2.1884 | 254514.9 | 289.1646 | di/tripepti | N |
| 351 | TPP2 | 290.1721 | 1 | 4.884233 | 114203.5 | 289.1648 | di/tripepti | N |
| 352 | TPP2 | 294.9403 | 1 | 1.669308 | 162880.5 | 293.933 | di/tripepti | N |

| | | | | | | | | |
|---|---|---|---|---|---|---|---|---|
| 353 | TPP2 | 295.13 | 1 | 12.59933 | 316486.1 | 294.1228 | di/tripepti | N |
| 354 | TPP2 | 295.1664 | 1 | 14.57049 | 124749.8 | 294.1592 | di/tripepti | N |
| 355 | TPP2 | 295.1665 | 1 | 13.37085 | 102192.4 | 294.1592 | di/tripepti | N |
| 356 | TPP2 | 295.2277 | 1 | 37.37093 | 120066.3 | 294.2204 | di/tripepti | N |
| 357 | TPP2 | 297.1092 | 1 | 2.634942 | 124889.3 | 296.1019 | di/tripepti | N |
| 358 | TPP2 | 300.1929 | 1 | 14.46062 | 293045.3 | 299.1856 | di/tripepti | N |
| 359 | TPP2 | 302.2086 | 1 | 11.27234 | 323010.1 | 301.2013 | tri/oligope | N |
| 360 | TPP2 | 304.1878 | 1 | 2.576392 | 149723.3 | 303.1805 | tri/oligope | N |
| 361 | TPP2 | 306.1669 | 1 | 2.1884 | 671211.7 | 305.1597 | tri/oligope | N |
| 362 | TPP2 | 314.2087 | 1 | 13.6132 | 154489.3 | 313.2014 | tri/oligope | N |
| 363 | TPP2 | 316.2244 | 1 | 21.19288 | 193751.9 | 315.2171 | tri/oligope | N |
| 364 | TPP2 | 316.2244 | 1 | 19.58301 | 159317.7 | 315.2172 | tri/oligope | N |
| 365 | TPP2 | 316.2244 | 1 | 20.82078 | 130570.8 | 315.2172 | tri/oligope | N |
| 366 | TPP2 | 316.2245 | 1 | 20.29017 | 174884.9 | 315.2172 | tri/oligope | N |
| 367 | TPP2 | 317.1833 | 1 | 6.438217 | 160153.5 | 316.176 | tri/oligope | N |
| 368 | TPP2 | 318.2037 | 1 | 12.29903 | 2201521 | 317.1964 | tri/oligope | N |
| 369 | TPP2 | 318.2037 | 1 | 16.70711 | 128254 | 317.1964 | tri/oligope | N |
| 370 | TPP2 | 322.144 | 1 | 2.290142 | 1307407 | 321.1368 | tri/oligope | N |
| 371 | TPP2 | 324.1299 | 1 | 2.1884 | 381735.8 | 323.1226 | tri/oligope | N |
| 372 | TPP2 | 328.2245 | 1 | 15.41029 | 148634.4 | 327.2172 | tri/oligope | N |
| 373 | TPP2 | 328.2245 | 1 | 18.28107 | 199573 | 327.2172 | tri/oligope | N |
| 374 | TPP2 | 330.2402 | 1 | 19.17391 | 158810.1 | 329.2329 | tri/oligope | N |
| 375 | TPP2 | 331.1988 | 1 | 2.1884 | 131297.8 | 330.1915 | tri/oligope | N |
| 376 | TPP2 | 331.2349 | 1 | 2.1884 | 225837.9 | 330.2276 | tri/oligope | N |
| 377 | TPP2 | 332.1829 | 1 | 11.14161 | 850968.6 | 331.1756 | tri/oligope | N |
| 378 | TPP2 | 332.2193 | 1 | 21.21669 | 2022052 | 331.212 | tri/oligope | N |
| 379 | TPP2 | 332.2194 | 1 | 15.88318 | 1064036 | 331.2121 | tri/oligope | N |
| 380 | TPP2 | 332.2194 | 1 | 20.46273 | 183692.4 | 331.2121 | tri/oligope | N |
| 381 | TPP2 | 332.2194 | 1 | 19.42958 | 131997.7 | 331.2121 | tri/oligope | N |
| 382 | TPP2 | 333.1779 | 1 | 2.1884 | 185614.8 | 332.1706 | tri/oligope | N |
| 383 | TPP2 | 333.1783 | 1 | 2.787617 | 149686.3 | 332.171 | tri/oligope | N |
| 384 | TPP2 | 336.139 | 1 | 21.02306 | 123294.9 | 335.1317 | tri/oligope | N |
| 385 | TPP2 | 340.9328 | 1 | 1.561442 | 110119.7 | 339.9256 | tri/oligope | N |
| 386 | TPP2 | 342.2043 | 1 | 21.74471 | 109611.2 | 341.197 | tri/oligope | N |
| 387 | TPP2 | 342.2402 | 1 | 20.72298 | 107263.3 | 341.233 | tri/oligope | N |
| 388 | TPP2 | 344.2559 | 1 | 23.29111 | 580437 | 343.2486 | tri/oligope | N |
| 389 | TPP2 | 344.256 | 1 | 22.40665 | 289134.7 | 343.2487 | tri/oligope | N |
| 390 | TPP2 | 344.256 | 1 | 22.17358 | 176573.2 | 343.2487 | tri/oligope | N |
| 391 | TPP2 | 345.1459 | 1 | 14.00087 | 128385 | 344.1386 | tri/oligope | N |
| 392 | TPP2 | 346.1987 | 1 | 16.41489 | 107450.7 | 345.1915 | tri/oligope | N |
| 393 | TPP2 | 346.2352 | 1 | 19.16568 | 155850.3 | 345.2279 | tri/oligope | N |
| 394 | TPP2 | 348.1778 | 1 | 2.25585 | 195006.6 | 347.1705 | tri/oligope | N |
| 395 | TPP2 | 348.178 | 1 | 9.999408 | 200735.4 | 347.1708 | tri/oligope | N |
| 396 | TPP2 | 350.1757 | 1 | 2.65355 | 124979.2 | 349.1684 | tri/oligope | N |
| 397 | TPP2 | 358.1986 | 1 | 15.62301 | 792640.9 | 357.1914 | tri/oligope | N |
| 398 | TPP2 | 358.2714 | 1 | 26.26385 | 291023.9 | 357.2642 | tri/oligope | N |
| 399 | TPP2 | 358.2715 | 1 | 25.32757 | 332119.5 | 357.2642 | tri/oligope | N |
| 400 | TPP2 | 358.2715 | 1 | 24.93741 | 403529.1 | 357.2643 | tri/oligope | N |
| 401 | TPP2 | 359.2303 | 1 | 16.81376 | 811984.9 | 358.223 | tri/oligope | N |
| 402 | TPP2 | 359.2303 | 1 | 15.41029 | 2364329 | 358.2231 | tri/oligope | N |

| | | | | | | | | |
|---|---|---|---|---|---|---|---|---|
| 403 | TPP2 | 359.2304 | 1 | 20.53754 | 103776 | 358.2232 | tri/oligope | N |
| 404 | TPP2 | 359.2411 | 1 | 2.1884 | 103627.9 | 358.2339 | tri/oligope | N |
| 405 | TPP2 | 360.2143 | 1 | 17.60134 | 159553.6 | 359.2071 | tri/oligope | N |
| 406 | TPP2 | 360.2144 | 1 | 21.24149 | 220418.2 | 359.2071 | tri/oligope | N |
| 407 | TPP2 | 360.2146 | 1 | 16.06823 | 354432 | 359.2073 | tri/oligope | N |
| 408 | TPP2 | 360.2146 | 1 | 20.4459 | 125092.6 | 359.2073 | tri/oligope | N |
| 409 | TPP2 | 361.2457 | 1 | 2.1884 | 116487.8 | 360.2384 | tri/oligope | N |
| 410 | TPP2 | 362.1937 | 1 | 12.14779 | 116113.1 | 361.1864 | tri/oligope | N |
| 411 | TPP2 | 362.2088 | 1 | 21.33214 | 1342999 | 361.2016 | tri/oligope | N |
| 412 | TPP2 | 366.134 | 1 | 2.1884 | 309980.6 | 365.1268 | tri/oligope | N |
| 413 | TPP2 | 371.2304 | 1 | 14.13379 | 178210.4 | 370.2231 | tri/oligope | N |
| 414 | TPP2 | 371.2305 | 1 | 11.14161 | 104817.3 | 370.2232 | tri/oligope | N |
| 415 | TPP2 | 373.2092 | 1 | 2.576392 | 121970.6 | 372.2019 | tri/oligope | N |
| 416 | TPP2 | 374.1092 | 1 | 2.1884 | 339847.9 | 373.102 | tri/oligope | N |
| 417 | TPP2 | 374.23 | 1 | 12.99975 | 139222.2 | 373.2227 | tri/oligope | N |
| 418 | TPP2 | 374.23 | 1 | 20.64168 | 440443 | 373.2228 | tri/oligope | N |
| 419 | TPP2 | 374.2301 | 1 | 21.18305 | 190482.7 | 373.2228 | tri/oligope | N |
| 420 | TPP2 | 374.2301 | 1 | 22.01933 | 437700.3 | 373.2228 | tri/oligope | N |
| 421 | TPP2 | 374.2302 | 1 | 14.32368 | 143120.8 | 373.2229 | tri/oligope | N |
| 422 | TPP2 | 375.189 | 1 | 8.368875 | 150048.7 | 374.1817 | tri/oligope | N |
| 423 | TPP2 | 377.2042 | 1 | 2.1884 | 254004.3 | 376.1969 | tri/oligope | N |
| 424 | TPP2 | 388.2201 | 1 | 2.1884 | 106019.3 | 387.2128 | tri/oligope | N |
| 425 | TPP2 | 389.241 | 1 | 10.70473 | 131638.8 | 388.2337 | tri/oligope | N |
| 426 | TPP2 | 389.2411 | 1 | 18.8771 | 100361.3 | 388.2338 | tri/oligope | N |
| 427 | TPP2 | 389.2517 | 1 | 2.199483 | 155264 | 388.2445 | tri/oligope | N |
| 428 | TPP2 | 392.1864 | 1 | 20.0029 | 878533.8 | 391.1791 | tri/oligope | N |
| 429 | TPP2 | 393.2148 | 1 | 23.93388 | 112259.4 | 392.2075 | tri/oligope | N |
| 430 | TPP2 | 398.1576 | 1 | 5.231058 | 509581.4 | 397.1503 | tri/oligope | N |
| 431 | TPP2 | 398.3428 | 1 | 34.87829 | 136017.9 | 397.3356 | tri/oligope | N |
| 432 | TPP2 | 402.2358 | 1 | 2.634942 | 127349 | 401.2285 | tri/oligope | N |
| 433 | TPP2 | 405.1992 | 1 | 2.199483 | 131013.8 | 404.192 | tri/oligope | N |
| 434 | TPP2 | 405.236 | 1 | 17.37768 | 150318.9 | 404.2287 | tri/oligope | N |
| 435 | TPP2 | 408.9202 | 1 | 1.561442 | 121875.9 | 407.913 | tri/oligope | N |
| 436 | TPP2 | 409.0204 | 1 | 2.210858 | 127835.9 | 408.0131 | tri/oligope | N |
| 437 | TPP2 | 410.1937 | 1 | 19.54245 | 353671.8 | 409.1864 | tri/oligope | N |
| 438 | TPP2 | 413.2776 | 1 | 21.36654 | 147092.8 | 412.2703 | tri/oligope | N |
| 439 | TPP2 | 419.2148 | 1 | 2.26725 | 145810.9 | 418.2075 | tri/oligope | N |
| 440 | TPP2 | 419.2304 | 1 | 24.53838 | 237200.4 | 418.2232 | tri/oligope | N |
| 441 | TPP2 | 419.2515 | 1 | 17.62796 | 245265 | 418.2442 | tri/oligope | N |
| 442 | TPP2 | 419.2516 | 1 | 13.19333 | 410168.8 | 418.2443 | tri/oligope | N |
| 443 | TPP2 | 424.2095 | 1 | 13.27385 | 146034.8 | 423.2022 | tri/oligope | N |
| 444 | TPP2 | 429.3091 | 1 | 25.69925 | 162239.8 | 428.3018 | tri/oligope | N |
| 445 | TPP2 | 431.2877 | 1 | 20.77187 | 1243182 | 430.2805 | tri/oligope | N |
| 446 | TPP2 | 440.1674 | 1 | 2.222067 | 156825 | 439.1601 | tri/oligope | N |
| 447 | TPP2 | 446.2256 | 1 | 2.1884 | 186565.4 | 445.2184 | tri/oligope | N |
| 448 | TPP2 | 446.2263 | 1 | 10.00723 | 294015.2 | 445.219 | tri/oligope | N |
| 449 | TPP2 | 449.2081 | 1 | 16.65863 | 240668.7 | 448.2009 | tri/oligope | N |
| 450 | TPP2 | 451.2568 | 1 | 18.82551 | 167728.3 | 450.2495 | tri/oligope | N |
| 451 | TPP2 | 458.2257 | 1 | 2.1884 | 484273 | 457.2184 | tri/oligope | N |
| 452 | TPP2 | 459.257 | 1 | 2.1884 | 111382 | 458.2498 | tri/oligope | N |

| | | | | | | | | |
|---|---|---|---|---|---|---|---|---|
| 453 | TPP2 | 230.6482 | 2 | 2.348325 | 137433.7 | 459.2819 | tri/oligope | N |
| 454 | TPP2 | 464.1981 | 1 | 22.38317 | 139024.4 | 463.1908 | tri/oligope | N |
| 455 | TPP2 | 464.218 | 1 | 2.1884 | 107898.5 | 463.2107 | tri/oligope | N |
| 456 | TPP2 | 464.2521 | 1 | 16.96685 | 304705.9 | 463.2448 | tri/oligope | N |
| 457 | TPP2 | 465.1737 | 1 | 2.1884 | 128303.4 | 464.1664 | tri/oligope | N |
| 458 | TPP2 | 467.3488 | 1 | 33.8306 | 104233.7 | 466.3415 | tri/oligope | N |
| 459 | TPP2 | 470.2987 | 1 | 21.43092 | 283038.9 | 469.2914 | tri/oligope | N |
| 460 | TPP2 | 472.2414 | 1 | 2.210858 | 144372.3 | 471.2341 | tri/oligope | N |
| 461 | TPP2 | 472.2569 | 1 | 27.18503 | 1450951 | 471.2496 | tri/oligope | N |
| 462 | TPP2 | 472.3147 | 1 | 23.31904 | 276080.8 | 471.3074 | tri/oligope | N |
| 463 | TPP2 | 475.2927 | 1 | 26.24974 | 3664380 | 474.2854 | tri/oligope | N |
| 464 | TPP2 | 476.273 | 1 | 16.28594 | 165093.6 | 475.2658 | tri/oligope | N |
| 465 | TPP2 | 476.9077 | 1 | 1.561442 | 123538.1 | 475.9004 | tri/oligope | N |
| 466 | TPP2 | 483.2464 | 1 | 12.65471 | 1483696 | 482.2392 | tri/oligope | N |
| 467 | TPP2 | 483.2466 | 1 | 14.38507 | 238287.1 | 482.2393 | tri/oligope | N |
| 468 | TPP2 | 486.294 | 1 | 18.44643 | 392367.9 | 485.2867 | tri/oligope | N |
| 469 | TPP2 | 487.289 | 1 | 16.28594 | 181478 | 486.2817 | tri/oligope | N |
| 470 | TPP2 | 490.2633 | 1 | 2.1884 | 100815.1 | 489.256 | tri/oligope | N |
| 471 | TPP2 | 245.6353 | 2 | 2.199483 | 219784.5 | 489.256 | tri/oligope | N |
| 472 | TPP2 | 246.624 | 2 | 18.75618 | 123195 | 491.2335 | tri/oligope | N |
| 473 | TPP2 | 504.3044 | 1 | 21.21066 | 105244.5 | 503.2972 | tri/oligope | N |
| 474 | TPP2 | 506.2296 | 1 | 20.4459 | 648582.1 | 505.2223 | tri/oligope | N |
| 475 | TPP2 | 509.2779 | 1 | 28.34368 | 578349.6 | 508.2706 | tri/oligope | N |
| 476 | TPP2 | 511.2048 | 1 | 2.360142 | 256324.8 | 510.1975 | tri/oligope | N |
| 477 | TPP2 | 511.3754 | 1 | 32.60342 | 121295.6 | 510.3681 | tri/oligope | N |
| 478 | TPP2 | 257.1473 | 2 | 19.73062 | 108819.3 | 512.28 | tri/oligope | N |
| 479 | TPP2 | 517.2636 | 1 | 15.08162 | 117452.9 | 516.2563 | tri/oligope | N |
| 480 | TPP2 | 518.2827 | 1 | 2.1884 | 109103.2 | 517.2755 | tri/oligope | N |
| 481 | TPP2 | 259.6455 | 2 | 2.177583 | 135074.9 | 517.2764 | tri/oligope | N |
| 482 | TPP2 | 518.3202 | 1 | 24.50025 | 423474.4 | 517.3129 | tri/oligope | N |
| 483 | TPP2 | 522.2576 | 1 | 17.90261 | 119411.4 | 521.2503 | tri/oligope | N |
| 484 | TPP2 | 526.3621 | 1 | 27.36036 | 146501.8 | 525.3548 | tri/oligope | N |
| 485 | TPP2 | 529.1979 | 1 | 15.42055 | 1074035 | 528.1906 | tri/oligope | N |
| 486 | TPP2 | 529.198 | 1 | 12.22855 | 100923.8 | 528.1908 | tri/oligope | N |
| 487 | TPP2 | 531.2899 | 1 | 2.15545 | 430404.1 | 530.2827 | tri/oligope | N |
| 488 | TPP2 | 266.1487 | 2 | 2.144192 | 591018 | 530.2828 | tri/oligope | N |
| 489 | TPP2 | 542.3568 | 1 | 22.3421 | 147563.8 | 541.3496 | tri/oligope | N |
| 490 | TPP2 | 544.8952 | 1 | 1.561442 | 142740.2 | 543.8879 | tri/oligope | N |
| 491 | TPP2 | 550.256 | 1 | 18.97212 | 376966.3 | 549.2487 | tri/oligope | N |
| 492 | TPP2 | 559.2742 | 1 | 2.612817 | 403368.5 | 558.2669 | tri/oligope | N |
| 493 | TPP2 | 559.3105 | 1 | 18.2159 | 1416560 | 558.3032 | tri/oligope | N |
| 494 | TPP2 | 280.1589 | 2 | 18.2159 | 466098.7 | 558.3032 | tri/oligope | N |
| 495 | TPP2 | 281.6642 | 2 | 2.199483 | 1143034 | 561.3138 | tri/oligope | N |
| 496 | TPP2 | 562.3211 | 1 | 2.1884 | 605490.2 | 561.3139 | tri/oligope | N |
| 497 | TPP2 | 562.8071 | 1 | 1.6454 | 113438.9 | 561.7998 | tri/oligope | N |
| 498 | TPP2 | 568.2635 | 1 | 17.72558 | 180515.5 | 567.2563 | tri/oligope | N |
| 499 | TPP2 | 569.2768 | 1 | 2.1884 | 158093 | 568.2696 | tri/oligope | N |
| 500 | TPP2 | 285.1423 | 2 | 2.199483 | 420129.2 | 568.2701 | tri/oligope | N |
| 501 | TPP2 | 571.3109 | 1 | 19.11498 | 333219.4 | 570.3036 | tri/oligope | N |
| 502 | TPP2 | 288.1748 | 2 | 3.037492 | 261617 | 574.335 | tri/oligope | N |

| | | | | | | | | |
|---|---|---|---|---|---|---|---|---|
| 503 | TPP2 | 577.2674 | 1 | 20.46273 | 568620.5 | 576.2601 | tri/oligope | N |
| 504 | TPP2 | 582.3167 | 1 | 18.77168 | 246216.8 | 581.3094 | tri/oligope | N |
| 505 | TPP2 | 292.1379 | 2 | 18.20898 | 195339.5 | 582.2613 | tri/oligope | N |
| 506 | TPP2 | 588.3011 | 1 | 17.36698 | 337762.6 | 587.2939 | tri/oligope | N |
| 507 | TPP2 | 588.3782 | 1 | 30.70853 | 120122 | 587.3709 | tri/oligope | N |
| 508 | TPP2 | 590.3208 | 1 | 28.20951 | 702972.3 | 589.3136 | tri/oligope | N |
| 509 | TPP2 | 296.1618 | 2 | 15.76422 | 232533.1 | 590.309 | tri/oligope | N |
| 510 | TPP2 | 300.6799 | 2 | 24.35998 | 599060.1 | 599.3453 | tri/oligope | N |
| 511 | TPP2 | 604.3369 | 1 | 27.89078 | 264414.7 | 603.3296 | tri/oligope | N |
| 512 | TPP2 | 607.3114 | 1 | 24.23044 | 129052.6 | 606.3041 | tri/oligope | N |
| 513 | TPP2 | 309.696 | 2 | 14.9813 | 414391.5 | 617.3774 | oligopepti | N |
| 514 | TPP2 | 310.6806 | 2 | 2.84185 | 162421.2 | 619.3467 | oligopepti | N |
| 515 | TPP2 | 311.1909 | 2 | 23.5841 | 113029.5 | 620.3672 | oligopepti | N |
| 516 | TPP2 | 314.683 | 2 | 24.32401 | 165250.3 | 627.3514 | oligopepti | N |
| 517 | TPP2 | 630.2747 | 1 | 2.177583 | 179485.2 | 629.2674 | oligopepti | N |
| 518 | TPP2 | 630.3483 | 1 | 17.08539 | 412169.4 | 629.341 | oligopepti | N |
| 519 | TPP2 | 315.6961 | 2 | 14.69896 | 127581.3 | 629.3777 | oligopepti | N |
| 520 | TPP2 | 316.6077 | 2 | 15.65833 | 520010.6 | 631.2008 | oligopepti | N |
| 521 | TPP2 | 632.3275 | 1 | 12.4944 | 1195900 | 631.3202 | oligopepti | N |
| 522 | TPP2 | 316.6675 | 2 | 12.50832 | 322312.8 | 631.3205 | oligopepti | N |
| 523 | TPP2 | 317.7068 | 2 | 2.88095 | 147682 | 633.3991 | oligopepti | N |
| 524 | TPP2 | 322.6858 | 2 | 14.01734 | 4288565 | 643.357 | oligopepti | N |
| 525 | TPP2 | 646.3108 | 1 | 20.39733 | 109521.6 | 645.3035 | oligopepti | N |
| 526 | TPP2 | 323.6992 | 2 | 15.89038 | 131596.2 | 645.3839 | oligopepti | N |
| 527 | TPP2 | 325.1663 | 2 | 14.87298 | 965231.7 | 648.318 | oligopepti | N |
| 528 | TPP2 | 328.6464 | 2 | 12.4944 | 224373.9 | 655.2782 | oligopepti | N |
| 529 | TPP2 | 657.3582 | 1 | 2.1884 | 140223 | 656.3509 | oligopepti | N |
| 530 | TPP2 | 329.183 | 2 | 2.199483 | 511208.5 | 656.3515 | oligopepti | N |
| 531 | TPP2 | 330.1912 | 2 | 2.210858 | 237772.3 | 658.3679 | oligopepti | N |
| 532 | TPP2 | 330.662 | 2 | 17.19422 | 2703659 | 659.3095 | oligopepti | N |
| 533 | TPP2 | 335.6412 | 2 | 12.50083 | 106958.2 | 669.2679 | oligopepti | N |
| 534 | TPP2 | 336.1987 | 2 | 24.82946 | 541674.1 | 670.3829 | oligopepti | N |
| 535 | TPP2 | 336.2095 | 2 | 15.24851 | 156601.6 | 670.4044 | oligopepti | N |
| 536 | TPP2 | 675.3129 | 1 | 23.21875 | 148625.9 | 674.3056 | oligopepti | N |
| 537 | TPP2 | 677.3494 | 1 | 15.12153 | 113689.1 | 676.3422 | oligopepti | N |
| 538 | TPP2 | 341.6848 | 2 | 18.63676 | 436375.2 | 681.3551 | oligopepti | N |
| 539 | TPP2 | 352.7039 | 2 | 24.3723 | 1540538 | 703.3932 | oligopepti | N |
| 540 | TPP2 | 235.8192 | 3 | 13.80561 | 103332.7 | 704.4358 | oligopepti | N |
| 541 | TPP2 | 706.3797 | 1 | 25.2859 | 186193.5 | 705.3724 | oligopepti | N |
| 542 | TPP2 | 353.6937 | 2 | 25.2859 | 215434.9 | 705.3728 | oligopepti | N |
| 543 | TPP2 | 359.7118 | 2 | 25.08268 | 613824.1 | 717.4091 | oligopepti | N |
| 544 | TPP2 | 364.2407 | 2 | 22.27803 | 362345.7 | 726.4668 | oligopepti | N |
| 545 | TPP2 | 729.3809 | 1 | 16.05263 | 147660.2 | 728.3737 | oligopepti | N |
| 546 | TPP2 | 366.7175 | 2 | 13.78144 | 265841.1 | 731.4204 | oligopepti | N |
| 547 | TPP2 | 367.2088 | 2 | 15.74576 | 474945.4 | 732.4031 | oligopepti | N |
| 548 | TPP2 | 735.37 | 1 | 25.08268 | 164027.5 | 734.3627 | oligopepti | N |
| 549 | TPP2 | 371.7176 | 2 | 24.99795 | 2414164 | 741.4207 | oligopepti | N |
| 550 | TPP2 | 373.1499 | 2 | 19.8353 | 228849 | 744.2852 | oligopepti | N |
| 551 | TPP2 | 745.4117 | 1 | 18.00378 | 135831.2 | 744.4044 | oligopepti | N |
| 552 | TPP2 | 373.2097 | 2 | 17.94768 | 106266.7 | 744.4048 | oligopepti | N |

| | | | | | | | | |
|---|---|---|---|---|---|---|---|---|
| 553 | TPP2 | 373.7073 | 2 | 12.86345 | 346472.2 | 745.4001 | oligopepti( | N |
| 554 | TPP2 | 373.7151 | 2 | 23.68373 | 113670.8 | 745.4156 | oligopepti( | N |
| 555 | TPP2 | 375.2072 | 2 | 14.40998 | 626744.1 | 748.3998 | oligopepti( | N |
| 556 | TPP2 | 377.6781 | 2 | 2.199483 | 115584.8 | 753.3416 | oligopepti( | N |
| 557 | TPP2 | 378.1825 | 2 | 1.4173 | 118779.1 | 754.3504 | oligopepti( | N |
| 558 | TPP2 | 758.4078 | 1 | 21.11267 | 107950.5 | 757.4005 | oligopepti( | N |
| 559 | TPP2 | 380.2038 | 2 | 27.11254 | 132292 | 758.393 | oligopepti( | N |
| 560 | TPP2 | 380.7206 | 2 | 14.90358 | 160914.6 | 759.4266 | oligopepti( | N |
| 561 | TPP2 | 386.7333 | 2 | 17.88735 | 499411.8 | 771.452 | oligopepti( | N |
| 562 | TPP2 | 386.7334 | 2 | 13.91943 | 160673.6 | 771.4523 | oligopepti( | N |
| 563 | TPP2 | 392.189 | 2 | 18.02023 | 226019.1 | 782.3635 | oligopepti( | N |
| 564 | TPP2 | 395.2304 | 2 | 25.14417 | 815112.3 | 788.4462 | oligopepti( | N |
| 565 | TPP2 | 799.4228 | 1 | 19.7228 | 150267 | 798.4156 | oligopepti( | N |
| 566 | TPP2 | 400.7127 | 2 | 15.83037 | 227359.2 | 799.4108 | oligopepti( | N |
| 567 | TPP2 | 403.1991 | 2 | 14.59439 | 190671.8 | 804.3836 | oligopepti( | N |
| 568 | TPP2 | 406.7389 | 2 | 27.92788 | 245560.7 | 811.4632 | oligopepti( | N |
| 569 | TPP2 | 412.2231 | 2 | 26.89203 | 133382.1 | 822.4316 | oligopepti( | N |
| 570 | TPP2 | 418.2153 | 2 | 27.08448 | 113306 | 834.416 | oligopepti( | N |
| 571 | TPP2 | 418.7077 | 2 | 17.20163 | 129305.9 | 835.4009 | oligopepti( | N |
| 572 | TPP2 | 279.4743 | 3 | 17.20163 | 349664.3 | 835.4011 | oligopepti( | N |
| 573 | TPP2 | 429.7128 | 2 | 20.27624 | 164365.9 | 857.411 | oligopepti( | N |
| 574 | TPP2 | 859.4555 | 1 | 20.87527 | 198767.9 | 858.4483 | oligopepti( | N |
| 575 | TPP2 | 430.7492 | 2 | 25.39282 | 338840.6 | 859.4839 | oligopepti( | N |
| 576 | TPP2 | 868.5095 | 1 | 29.96952 | 179414.6 | 867.5022 | oligopepti( | N |
| 577 | TPP2 | 435.7309 | 2 | 23.5841 | 2333700 | 869.4472 | oligopepti( | N |
| 578 | TPP2 | 436.2368 | 2 | 25.44233 | 1294525 | 870.4591 | oligopepti( | N |
| 579 | TPP2 | 438.7176 | 2 | 16.70711 | 182872.2 | 875.4207 | oligopepti( | N |
| 580 | TPP2 | 884.8336 | 1 | 1.573417 | 100001.1 | 883.8263 | oligopepti( | N |
| 581 | TPP2 | 447.7411 | 2 | 27.07788 | 138304.3 | 893.4677 | oligopepti( | N |
| 582 | TPP2 | 454.2237 | 2 | 16.68283 | 250922.1 | 906.4329 | oligopepti( | N |
| 583 | TPP2 | 907.4403 | 1 | 16.68283 | 193707.2 | 906.433 | oligopepti( | N |
| 584 | TPP2 | 304.4935 | 3 | 15.66727 | 315265.3 | 910.4587 | oligopepti( | N |
| 585 | TPP2 | 304.8687 | 3 | 15.53763 | 105409.6 | 911.5842 | oligopepti( | N |
| 586 | TPP2 | 457.7701 | 2 | 2.199483 | 183452 | 913.5256 | oligopepti( | N |
| 587 | TPP2 | 305.516 | 3 | 2.210858 | 182323.8 | 913.5261 | oligopepti( | N |
| 588 | TPP2 | 465.2316 | 2 | 20.72298 | 101446.2 | 928.4487 | oligopepti( | N |
| 589 | TPP2 | 931.6278 | 1 | 32.61258 | 119629.2 | 930.6205 | oligopepti( | N |
| 590 | TPP2 | 471.2494 | 2 | 23.69223 | 294638.4 | 940.4843 | oligopepti( | N |
| 591 | TPP2 | 477.3021 | 2 | 32.61258 | 111224.7 | 952.5897 | oligopepti( | N |
| 592 | TPP2 | 478.7154 | 2 | 2.348325 | 125372 | 955.4163 | oligopepti( | N |
| 593 | TPP2 | 487.7289 | 2 | 16.5951 | 102209.7 | 973.4432 | oligopepti( | N |
| 594 | TPP2 | 488.7527 | 2 | 22.71693 | 856062.8 | 975.4908 | oligopepti( | N |
| 595 | TPP2 | 976.4985 | 1 | 22.71693 | 751153.2 | 975.4912 | oligopepti( | N |
| 596 | TPP2 | 489.7421 | 2 | 19.43706 | 843621.7 | 977.4697 | oligopepti( | N |
| 597 | TPP2 | 978.4775 | 1 | 19.43706 | 173654.5 | 977.4702 | oligopepti( | N |
| 598 | TPP2 | 496.7581 | 2 | 25.98807 | 114114.7 | 991.5017 | oligopepti( | N |
| 599 | TPP2 | 500.7317 | 2 | 22.71693 | 223680.5 | 999.4489 | oligopepti( | N |
| 600 | TPP2 | 334.157 | 3 | 22.71693 | 252686.7 | 999.449 | oligopepti( | N |
| 601 | TPP2 | 500.7499 | 2 | 21.03171 | 191217.4 | 999.4852 | oligopepti( | N |
| 602 | TPP2 | 501.7211 | 2 | 19.43706 | 104089 | 1001.428 | oligopepti( | N |

| | | | | | | | | |
|---|---|---|---|---|---|---|---|---|
| 603 | TPP2 | 334.8166 | 3 | 19.44806 | 198565 | 1001.428 | oligopepti( | N |
| 604 | TPP2 | 506.7347 | 2 | 22.09797 | 190637.3 | 1011.455 | oligopepti( | N |
| 605 | TPP2 | 506.7681 | 2 | 24.02238 | 363455.7 | 1011.522 | oligopepti( | N |
| 606 | TPP2 | 507.7263 | 2 | 22.71693 | 151024.3 | 1013.438 | oligopepti( | N |
| 607 | TPP2 | 338.8203 | 3 | 22.71693 | 179328.5 | 1013.439 | oligopepti( | N |
| 608 | TPP2 | 339.4799 | 3 | 19.43706 | 118235.2 | 1015.418 | oligopepti( | N |
| 609 | TPP2 | 512.1956 | 2 | 24.62472 | 213821.9 | 1022.377 | oligopepti( | N |
| 610 | TPP2 | 514.3124 | 2 | 18.05154 | 385412.4 | 1026.61 | oligopepti( | N |
| 611 | TPP2 | 343.2111 | 3 | 18.05154 | 1310039 | 1026.611 | oligopepti( | N |
| 612 | TPP2 | 515.3028 | 2 | 25.69925 | 112058.8 | 1028.591 | oligopepti( | N |
| 613 | TPP2 | 516.7345 | 2 | 2.166583 | 189218.8 | 1031.454 | oligopepti( | N |
| 614 | TPP2 | 519.7441 | 2 | 21.30231 | 316798.1 | 1037.474 | oligopepti( | N |
| 615 | TPP2 | 1038.481 | 1 | 21.31288 | 134323.5 | 1037.474 | oligopepti( | N |
| 616 | TPP2 | 525.7499 | 2 | 14.22944 | 247002.7 | 1049.485 | oligopepti( | N |
| 617 | TPP2 | 540.7757 | 2 | 23.11044 | 115493.7 | 1079.537 | oligopepti( | N |
| 618 | TPP2 | 542.2865 | 2 | 24.37846 | 139234 | 1082.558 | oligopepti( | N |
| 619 | TPP2 | 545.2947 | 2 | 27.21258 | 552011.1 | 1088.575 | oligopepti( | N |
| 620 | TPP2 | 1089.582 | 1 | 27.22069 | 227187.3 | 1088.575 | oligopepti( | N |
| 621 | TPP2 | 549.7557 | 2 | 22.24138 | 255316.6 | 1097.497 | oligopepti( | N |
| 622 | TPP2 | 366.8398 | 3 | 22.24138 | 967351 | 1097.497 | oligopepti( | N |
| 623 | TPP2 | 371.8518 | 3 | 27.22069 | 133461.5 | 1112.534 | oligopepti( | N |
| 624 | TPP2 | 560.7695 | 2 | 17.53639 | 155707.4 | 1119.524 | oligopepti( | N |
| 625 | TPP2 | 561.2689 | 2 | 16.65863 | 214792.5 | 1120.523 | oligopepti( | N |
| 626 | TPP2 | 563.2771 | 2 | 26.13627 | 969749 | 1124.54 | oligopepti( | N |
| 627 | TPP2 | 563.8299 | 2 | 26.89203 | 200381.9 | 1125.645 | oligopepti( | N |
| 628 | TPP2 | 383.8396 | 3 | 26.1419 | 180621.5 | 1148.497 | oligopepti( | N |
| 629 | TPP2 | 578.2939 | 2 | 15.26479 | 103295.5 | 1154.573 | oligopepti( | N |
| 630 | TPP2 | 388.5028 | 3 | 26.1419 | 140965.5 | 1162.487 | oligopepti( | N |
| 631 | TPP2 | 588.8118 | 2 | 23.49098 | 201467.4 | 1175.609 | oligopepti( | N |
| 632 | TPP2 | 595.8199 | 2 | 25.75923 | 115551.4 | 1189.625 | oligopepti( | N |
| 633 | TPP2 | 599.3483 | 2 | 25.91261 | 807402.2 | 1196.682 | oligopepti( | N |
| 634 | TPP2 | 618.283 | 2 | 16.34216 | 128646.4 | 1234.551 | oligopepti( | N |
| 635 | TPP2 | 625.2908 | 2 | 17.23412 | 705621.6 | 1248.567 | oligopepti( | N |
| 636 | TPP2 | 630.3092 | 2 | 21.68146 | 229687.9 | 1258.604 | oligopepti( | N |
| 637 | TPP2 | 636.3271 | 2 | 24.66195 | 473545.3 | 1270.64 | oligopepti( | N |
| 638 | TPP2 | 649.8724 | 2 | 25.57779 | 257887.6 | 1297.73 | oligopepti( | N |
| 639 | TPP2 | 651.8283 | 2 | 20.888 | 473567.9 | 1301.642 | oligopepti( | N |
| 640 | TPP2 | 655.8907 | 2 | 29.23105 | 192938.9 | 1309.767 | oligopepti( | N |
| 641 | TPP2 | 670.3548 | 2 | 18.33473 | 254607.8 | 1338.695 | oligopepti( | N |
| 642 | TPP2 | 674.8312 | 2 | 18.90288 | 157804.8 | 1347.648 | oligopepti( | N |
| 643 | TPP2 | 687.347 | 2 | 20.65529 | 525009.9 | 1372.679 | oligopepti( | N |
| 644 | TPP2 | 694.3347 | 2 | 24.80748 | 108919.6 | 1386.655 | oligopepti( | N |
| 645 | TPP2 | 705.8735 | 2 | 18.64552 | 171050.3 | 1409.733 | oligopepti( | N |
| 646 | TPP2 | 472.2317 | 3 | 24.81514 | 101790.3 | 1413.673 | oligopepti( | N |
| 647 | TPP2 | 711.841 | 2 | 23.28468 | 121497.1 | 1421.667 | oligopepti( | N |
| 648 | TPP2 | 712.4326 | 2 | 30.734 | 202858.5 | 1422.851 | oligopepti( | N |
| 649 | TPP2 | 717.8593 | 2 | 26.12399 | 123869.6 | 1433.704 | oligopepti( | N |
| 650 | TPP2 | 722.8657 | 2 | 21.04044 | 373114.9 | 1443.717 | oligopepti( | N |
| 651 | TPP2 | 755.9494 | 2 | 30.09504 | 126121.7 | 1509.884 | oligopepti( | N |
| 652 | TPP2 | 758.3836 | 2 | 21.31288 | 223561 | 1514.753 | oligopepti( | N |

| | | | | | | | | |
|---|---|---|---|---|---|---|---|---|
| 653 | TPP2 | 520.5717 | 3 | 21.39325 | 107799.1 | 1558.693 | oligopepti | N |
| 654 | TPP2 | 435.4843 | 4 | 17.48428 | 162278.5 | 1737.908 | oligopepti | N |
| 655 | TPP2 | 580.3102 | 3 | 17.49148 | 114463.1 | 1737.909 | oligopepti | N |
| 656 | TPP2 | 586.9664 | 3 | 18.92811 | 188928.6 | 1757.877 | oligopepti | N |
| 657 | TPP2 | 739.0584 | 3 | 28.37018 | 180560.4 | 2214.153 | oligopepti | N |

| ID | Protein | mz | charge | retentionti | raw.abund | mass | type | Match |
|---|---|---|---|---|---|---|---|---|
| 2 | Q | 231.1711 | 1 | 5.21135 | 728053.5 | 230.1638 | di/tripepti | Y |
| 3 | Q | 231.1711 | 1 | 9.658717 | 137511.5 | 230.1639 | di/tripepti | Y |
| 4 | Q | 231.1711 | 1 | 12.02454 | 144395.5 | 230.1638 | di/tripepti | Y |
| 5 | Q | 233.15 | 1 | 2.18805 | 418005.5 | 232.1427 | di/tripepti | Y |
| 6 | Q | 233.1502 | 1 | 2.753658 | 179755.5 | 232.143 | di/tripepti | Y |
| 8 | Q | 239.1032 | 1 | 2.42835 | 153963.7 | 238.0959 | di/tripepti | Y |
| 11 | Q | 245.1866 | 1 | 16.44068 | 370144.7 | 244.1794 | di/tripepti | Y |
| 13 | Q | 245.1867 | 1 | 14.53846 | 314807.1 | 244.1794 | di/tripepti | Y |
| 15 | Q | 246.1453 | 1 | 2.18805 | 395316.6 | 245.1381 | di/tripepti | Y |
| 16 | Q | 246.1455 | 1 | 2.635392 | 160203.4 | 245.1382 | di/tripepti | Y |
| 20 | Q | 247.1294 | 1 | 2.18805 | 616068.5 | 246.1221 | di/tripepti | Y |
| 21 | Q | 247.1297 | 1 | 3.080392 | 287335.6 | 246.1224 | di/tripepti | Y |
| 22 | Q | 247.1297 | 1 | 4.681792 | 764079.8 | 246.1224 | di/tripepti | Y |
| 27 | Q | 253.1189 | 1 | 2.292533 | 154937 | 252.1117 | di/tripepti | Y |
| 30 | Q | 260.1611 | 1 | 2.18805 | 354691.1 | 259.1538 | di/tripepti | Y |
| 31 | Q | 260.1613 | 1 | 2.42835 | 217312.7 | 259.154 | di/tripepti | Y |
| 34 | Q | 261.1452 | 1 | 2.18805 | 457723.6 | 260.1379 | di/tripepti | Y |
| 35 | Q | 261.1455 | 1 | 3.245683 | 272568.2 | 260.1382 | di/tripepti | Y |
| 36 | Q | 261.1454 | 1 | 5.392858 | 305189.4 | 260.1381 | di/tripepti | Y |
| 37 | Q | 262.1404 | 1 | 1.965242 | 102662.2 | 261.1331 | di/tripepti | Y |
| 38 | Q | 263.1434 | 1 | 11.50638 | 133078.8 | 262.1361 | di/tripepti | Y |
| 41 | Q | 267.1346 | 1 | 2.4006 | 175573.1 | 266.1274 | di/tripepti | Y |
| 42 | Q | 269.1613 | 1 | 2.176683 | 125615.7 | 268.154 | di/tripepti | Y |
| 45 | Q | 281.114 | 1 | 2.557267 | 180888.3 | 280.1068 | di/tripepti | Y |
| 51 | Q | 295.13 | 1 | 3.238992 | 248941.3 | 294.1228 | di/tripepti | Y |
| 56 | Q | 302.2086 | 1 | 15.13646 | 146504.9 | 301.2013 | tri/oligope | Y |
| 57 | Q | 302.2083 | 1 | 19.85459 | 279609.6 | 301.201 | tri/oligope | Y |
| 58 | Q | 303.1671 | 1 | 2.18805 | 181954.1 | 302.1598 | tri/oligope | Y |
| 59 | Q | 304.1511 | 1 | 2.18805 | 498753.9 | 303.1438 | tri/oligope | Y |
| 63 | Q | 318.167 | 1 | 2.664417 | 196735.3 | 317.1598 | tri/oligope | Y |
| 64 | Q | 318.1669 | 1 | 2.199558 | 484767 | 317.1597 | tri/oligope | Y |
| 65 | Q | 318.1672 | 1 | 8.984167 | 172802.6 | 317.1599 | tri/oligope | Y |
| 73 | Q | 331.1661 | 1 | 2.806017 | 151408.2 | 330.1589 | tri/oligope | Y |
| 74 | Q | 331.2852 | 1 | 38.47418 | 104876.4 | 330.2779 | tri/oligope | Y |
| 75 | Q | 332.1827 | 1 | 2.557267 | 189767.4 | 331.1754 | tri/oligope | Y |
| 76 | Q | 332.2193 | 1 | 3.073592 | 133430.3 | 331.212 | tri/oligope | Y |
| 80 | Q | 334.1617 | 1 | 2.18805 | 262212.4 | 333.1544 | tri/oligope | Y |
| 82 | Q | 342.2403 | 1 | 21.77958 | 144714.9 | 341.233 | tri/oligope | Y |
| 94 | Q | 363.1563 | 1 | 2.18805 | 128936.6 | 362.149 | tri/oligope | Y |
| 100 | Q | 376.1724 | 1 | 2.165425 | 105204.1 | 375.1651 | tri/oligope | Y |
| 106 | Q | 395.0398 | 1 | 1.600225 | 125752.6 | 394.0325 | tri/oligope | Y |
| 112 | Q | 415.2123 | 1 | 36.47458 | 142327.1 | 414.205 | tri/oligope | Y |
| 134 | Q | 463.0272 | 1 | 1.600225 | 161615.3 | 462.02 | tri/oligope | Y |
| 141 | Q | 479.3109 | 1 | 37.11703 | 137564.4 | 478.3036 | tri/oligope | Y |
| 156 | Q | 515.2921 | 1 | 37.11703 | 110832.6 | 514.2848 | tri/oligope | Y |
| 161 | Q | 264.175 | 2 | 35.27874 | 101924 | 526.3355 | tri/oligope | Y |
| 165 | Q | 531.0147 | 1 | 1.600225 | 204076.3 | 530.0074 | tri/oligope | Y |
| 174 | Q | 274.168 | 2 | 35.94152 | 106009 | 546.3214 | tri/oligope | Y |
| 194 | Q | 599.0026 | 1 | 1.600225 | 181625.2 | 597.9953 | tri/oligope | Y |

| | | | | | | | | |
|---|---|---|---|---|---|---|---|---|
| 218 | Q | 666.9901 | 1 | 1.600225 | 185240.2 | 665.9828 | oligopepti | Y |
| 240 | Q | 734.9777 | 1 | 1.600225 | 156251 | 733.9704 | oligopepti | Y |
| 258 | Q | 802.9652 | 1 | 1.600225 | 138766.2 | 801.958 | oligopepti | Y |
| 296 | Q | 974.8154 | 1 | 1.600225 | 240696.7 | 973.8082 | oligopepti | Y |
| 297 | Q | 990.7897 | 1 | 1.611008 | 144750.1 | 989.7825 | oligopepti | Y |
| 338 | Q | 223.1084 | 1 | 10.38069 | 188166.8 | 222.1011 | dipeptide | N |
| 339 | Q | 223.1085 | 1 | 4.891675 | 118922.8 | 222.1012 | dipeptide | N |
| 340 | Q | 226.9521 | 1 | 1.543025 | 315868 | 225.9448 | di/tripepti | N |
| 341 | Q | 229.1555 | 1 | 8.956642 | 323117.9 | 228.1482 | di/tripepti | N |
| 342 | Q | 229.1555 | 1 | 12.04725 | 110059.5 | 228.1482 | di/tripepti | N |
| 343 | Q | 229.1555 | 1 | 4.911475 | 135928.8 | 228.1482 | di/tripepti | N |
| 344 | Q | 231.1711 | 1 | 8.888058 | 257914 | 230.1638 | di/tripepti | N |
| 345 | Q | 232.1296 | 1 | 2.18805 | 170234.6 | 231.1223 | di/tripepti | N |
| 346 | Q | 233.1137 | 1 | 2.18805 | 196613.4 | 232.1064 | di/tripepti | N |
| 347 | Q | 237.1241 | 1 | 11.19039 | 103955.8 | 236.1169 | di/tripepti | N |
| 348 | Q | 239.2009 | 1 | 36.01476 | 121683 | 238.1936 | di/tripepti | N |
| 349 | Q | 263.14 | 1 | 14.44434 | 135878.1 | 262.1327 | di/tripepti | N |
| 350 | Q | 269.0887 | 1 | 2.18805 | 100651.6 | 268.0814 | di/tripepti | N |
| 351 | Q | 272.1612 | 1 | 2.280725 | 106808.4 | 271.1539 | di/tripepti | N |
| 352 | Q | 276.1561 | 1 | 2.18805 | 301068.4 | 275.1488 | di/tripepti | N |
| 353 | Q | 279.2325 | 1 | 37.01757 | 152621.9 | 278.2252 | di/tripepti | N |
| 354 | Q | 281.1142 | 1 | 11.55701 | 311625 | 280.1069 | di/tripepti | N |
| 355 | Q | 281.1507 | 1 | 5.8785 | 107740.1 | 280.1434 | di/tripepti | N |
| 356 | Q | 284.2591 | 1 | 36.02242 | 215580.1 | 283.2518 | di/tripepti | N |
| 357 | Q | 288.1929 | 1 | 14.6512 | 105315.8 | 287.1856 | di/tripepti | N |
| 358 | Q | 290.1352 | 1 | 2.03365 | 112886.5 | 289.128 | di/tripepti | N |
| 359 | Q | 290.1717 | 1 | 2.18805 | 388015.3 | 289.1644 | di/tripepti | N |
| 360 | Q | 292.1301 | 1 | 16.44068 | 130928.8 | 291.1228 | di/tripepti | N |
| 361 | Q | 294.9403 | 1 | 1.676875 | 422979.1 | 293.9331 | di/tripepti | N |
| 362 | Q | 297.1091 | 1 | 2.664417 | 147524.8 | 296.1018 | di/tripepti | N |
| 363 | Q | 300.1929 | 1 | 13.37495 | 104527.9 | 299.1857 | di/tripepti | N |
| 364 | Q | 302.1719 | 1 | 2.18805 | 204389 | 301.1647 | tri/oligope | N |
| 365 | Q | 304.1874 | 1 | 2.18805 | 194104.4 | 303.1801 | tri/oligope | N |
| 366 | Q | 306.1302 | 1 | 1.944867 | 127688.4 | 305.1229 | tri/oligope | N |
| 367 | Q | 314.2087 | 1 | 12.99117 | 161690.7 | 313.2014 | tri/oligope | N |
| 368 | Q | 316.188 | 1 | 13.04861 | 113796.7 | 315.1807 | tri/oligope | N |
| 369 | Q | 317.1828 | 1 | 2.245708 | 184771 | 316.1755 | tri/oligope | N |
| 370 | Q | 318.2037 | 1 | 14.59879 | 137011.8 | 317.1964 | tri/oligope | N |
| 371 | Q | 320.1254 | 1 | 17.67406 | 106195.7 | 319.1182 | tri/oligope | N |
| 372 | Q | 320.146 | 1 | 2.176683 | 111242.8 | 319.1387 | tri/oligope | N |
| 373 | Q | 328.2245 | 1 | 18.05055 | 258696.2 | 327.2172 | tri/oligope | N |
| 374 | Q | 328.2245 | 1 | 16.88998 | 231561.1 | 327.2172 | tri/oligope | N |
| 375 | Q | 330.2037 | 1 | 11.44704 | 102079.5 | 329.1964 | tri/oligope | N |
| 376 | Q | 332.1829 | 1 | 11.55701 | 101081.8 | 331.1756 | tri/oligope | N |
| 377 | Q | 333.1776 | 1 | 2.18805 | 180087.7 | 332.1703 | tri/oligope | N |
| 378 | Q | 334.1411 | 1 | 13.74723 | 131397.3 | 333.1339 | tri/oligope | N |
| 379 | Q | 334.1775 | 1 | 18.0365 | 134889.1 | 333.1702 | tri/oligope | N |
| 380 | Q | 338.3429 | 1 | 39.28503 | 181364.7 | 337.3356 | tri/oligope | N |
| 381 | Q | 342.24 | 1 | 21.25604 | 119712.3 | 341.2327 | tri/oligope | N |
| 382 | Q | 342.2403 | 1 | 20.73765 | 110676 | 341.233 | tri/oligope | N |

| | | | | | | | | |
|---|---|---|---|---|---|---|---|---|
| 383 | Q | 343.1986 | 1 | 2.18805 | 134521.6 | 342.1913 | tri/oligope | N |
| 384 | Q | 346.1987 | 1 | 17.06241 | 108670.6 | 345.1914 | tri/oligope | N |
| 385 | Q | 347.1571 | 1 | 2.03365 | 100337.2 | 346.1498 | tri/oligope | N |
| 386 | Q | 348.1411 | 1 | 2.022317 | 100446.8 | 347.1339 | tri/oligope | N |
| 387 | Q | 348.1776 | 1 | 2.222733 | 384782.5 | 347.1703 | tri/oligope | N |
| 388 | Q | 350.1727 | 1 | 17.50589 | 111000.6 | 349.1654 | tri/oligope | N |
| 389 | Q | 360.2144 | 1 | 16.85426 | 123735.6 | 359.2072 | tri/oligope | N |
| 390 | Q | 360.2145 | 1 | 20.45567 | 200705.7 | 359.2072 | tri/oligope | N |
| 391 | Q | 361.1726 | 1 | 2.18805 | 234145.8 | 360.1654 | tri/oligope | N |
| 392 | Q | 362.1934 | 1 | 2.257225 | 226954.2 | 361.1861 | tri/oligope | N |
| 393 | Q | 362.2091 | 1 | 21.36968 | 113933.4 | 361.2019 | tri/oligope | N |
| 394 | Q | 364.1883 | 1 | 16.67554 | 105968.5 | 363.181 | tri/oligope | N |
| 395 | Q | 389.2041 | 1 | 2.199558 | 128851.7 | 388.1968 | tri/oligope | N |
| 396 | Q | 404.2147 | 1 | 2.18805 | 104664.2 | 403.2075 | tri/oligope | N |
| 397 | Q | 405.1991 | 1 | 2.18805 | 185159.9 | 404.1919 | tri/oligope | N |
| 398 | Q | 413.267 | 1 | 36.75835 | 489794.6 | 412.2597 | tri/oligope | N |
| 399 | Q | 413.2676 | 1 | 38.05748 | 312830.3 | 412.2603 | tri/oligope | N |
| 400 | Q | 425.1776 | 1 | 1.852758 | 161036.4 | 424.1703 | tri/oligope | N |
| 401 | Q | 432.2099 | 1 | 2.199558 | 109190.1 | 431.2026 | tri/oligope | N |
| 402 | Q | 436.3433 | 1 | 35.94152 | 557059.9 | 435.336 | tri/oligope | N |
| 403 | Q | 436.3438 | 1 | 0.019108 | 257556.2 | 435.3365 | tri/oligope | N |
| 404 | Q | 441.2988 | 1 | 37.35428 | 173848.4 | 440.2915 | tri/oligope | N |
| 405 | Q | 446.2254 | 1 | 2.18805 | 127273.9 | 445.2181 | tri/oligope | N |
| 406 | Q | 450.1631 | 1 | 2.18805 | 112960.4 | 449.1558 | tri/oligope | N |
| 407 | Q | 463.298 | 1 | 37.55515 | 132748.2 | 462.2907 | tri/oligope | N |
| 408 | Q | 469.3297 | 1 | 37.19694 | 103915.7 | 468.3225 | tri/oligope | N |
| 409 | Q | 488.8733 | 1 | 1.632492 | 162513.1 | 487.866 | tri/oligope | N |
| 410 | Q | 533.455 | 1 | 37.39097 | 109600.9 | 532.4477 | tri/oligope | N |
| 411 | Q | 395.2053 | 2 | 16.56945 | 100381.6 | 788.396 | oligopeptic | N |

| ID | Protein | mz | charge | retentiont | raw.abund | mass | type | Match |
|---|---|---|---|---|---|---|---|---|
| 2 | QrtoE | 231.1711 | 1 | 5.220383 | 986951.6 | 230.1638 | di/tripepti | Y |
| 3 | QrtoE | 231.1711 | 1 | 9.671092 | 323583.1 | 230.1639 | di/tripepti | Y |
| 4 | QrtoE | 231.1711 | 1 | 12.02031 | 330636.8 | 230.1638 | di/tripepti | Y |
| 5 | QrtoE | 233.15 | 1 | 2.186908 | 404703.4 | 232.1427 | di/tripepti | Y |
| 6 | QrtoE | 233.1502 | 1 | 2.757242 | 241287.1 | 232.143 | di/tripepti | Y |
| 8 | QrtoE | 239.1032 | 1 | 2.414267 | 151311.8 | 238.0959 | di/tripepti | Y |
| 10 | QrtoE | 245.1137 | 1 | 1.963575 | 101505.6 | 244.1064 | di/tripepti | Y |
| 11 | QrtoE | 245.1866 | 1 | 16.42108 | 823777.3 | 244.1794 | di/tripepti | Y |
| 12 | QrtoE | 245.1869 | 1 | 15.88466 | 127010.1 | 244.1796 | di/tripepti | Y |
| 13 | QrtoE | 245.1867 | 1 | 14.52336 | 595998.5 | 244.1794 | di/tripepti | Y |
| 15 | QrtoE | 246.1453 | 1 | 2.186908 | 454986.8 | 245.1381 | di/tripepti | Y |
| 16 | QrtoE | 246.1455 | 1 | 2.640517 | 269079.6 | 245.1382 | di/tripepti | Y |
| 20 | QrtoE | 247.1294 | 1 | 2.186908 | 608208.8 | 246.1221 | di/tripepti | Y |
| 21 | QrtoE | 247.1297 | 1 | 3.08275 | 316341.4 | 246.1224 | di/tripepti | Y |
| 22 | QrtoE | 247.1297 | 1 | 4.689058 | 751819.1 | 246.1224 | di/tripepti | Y |
| 23 | QrtoE | 249.1275 | 1 | 2.771358 | 136513.1 | 248.1202 | di/tripepti | Y |
| 27 | QrtoE | 253.1189 | 1 | 2.285392 | 262205.9 | 252.1117 | di/tripepti | Y |
| 28 | QrtoE | 253.1192 | 1 | 9.3319 | 114916.7 | 252.1119 | di/tripepti | Y |
| 30 | QrtoE | 260.1611 | 1 | 2.186908 | 425273 | 259.1538 | di/tripepti | Y |
| 30 | QrtoE | 260.1613 | 1 | 2.414267 | 336702 | 259.154 | di/tripepti | Y |
| 33 | QrtoE | 260.1975 | 1 | 1.809775 | 102922.7 | 259.1902 | di/tripepti | Y |
| 34 | QrtoE | 261.1452 | 1 | 2.186908 | 498519.1 | 260.1379 | di/tripepti | Y |
| 35 | QrtoE | 261.1455 | 1 | 3.2497 | 264382.5 | 260.1382 | di/tripepti | Y |
| 36 | QrtoE | 261.1454 | 1 | 5.402433 | 410028.5 | 260.1381 | di/tripepti | Y |
| 37 | QrtoE | 262.1404 | 1 | 1.963575 | 127956.1 | 261.1331 | di/tripepti | Y |
| 38 | QrtoE | 263.1434 | 1 | 11.50003 | 277303.9 | 262.1361 | di/tripepti | Y |
| 41 | QrtoE | 267.1346 | 1 | 2.386992 | 198076.6 | 266.1274 | di/tripepti | Y |
| 42 | QrtoE | 269.1613 | 1 | 2.176208 | 204018.9 | 268.154 | di/tripepti | Y |
| 45 | QrtoE | 281.114 | 1 | 2.553442 | 171609 | 280.1068 | di/tripepti | Y |
| 47 | QrtoE | 288.1929 | 1 | 15.92202 | 111615.2 | 287.1856 | di/tripepti | Y |
| 48 | QrtoE | 290.1719 | 1 | 2.656808 | 109784 | 289.1646 | di/tripepti | Y |
| 51 | QrtoE | 295.13 | 1 | 3.2417 | 316304.3 | 294.1228 | di/tripepti | Y |
| 56 | QrtoE | 302.2086 | 1 | 15.12705 | 181077.3 | 301.2013 | tri/oligope | Y |
| 57 | QrtoE | 302.2083 | 1 | 19.84419 | 375966.9 | 301.201 | tri/oligope | Y |
| 58 | QrtoE | 303.1671 | 1 | 2.186908 | 285122.8 | 302.1598 | tri/oligope | Y |
| 59 | QrtoE | 304.1511 | 1 | 2.186908 | 487485 | 303.1438 | tri/oligope | Y |
| 63 | QrtoE | 318.167 | 1 | 2.667942 | 194647.1 | 317.1598 | tri/oligope | Y |
| 64 | QrtoE | 318.1669 | 1 | 2.1978 | 536495.2 | 317.1597 | tri/oligope | Y |
| 65 | QrtoE | 318.1672 | 1 | 8.998808 | 199752 | 317.1599 | tri/oligope | Y |
| 66 | QrtoE | 318.1824 | 1 | 22.57593 | 158030.6 | 317.1752 | tri/oligope | Y |
| 70 | QrtoE | 325.1137 | 1 | 1.73225 | 122376.7 | 324.1064 | tri/oligope | Y |
| 73 | QrtoE | 331.1661 | 1 | 2.806233 | 172415.4 | 330.1589 | tri/oligope | Y |
| 74 | QrtoE | 331.2852 | 1 | 38.45228 | 122142.8 | 330.2779 | tri/oligope | Y |
| 75 | QrtoE | 332.1826 | 1 | 2.296467 | 115312.5 | 331.1753 | tri/oligope | Y |
| 75 | QrtoE | 332.1827 | 1 | 2.553442 | 214580.6 | 331.1754 | tri/oligope | Y |
| 76 | QrtoE | 332.2193 | 1 | 3.075067 | 174775.1 | 331.212 | tri/oligope | Y |
| 80 | QrtoE | 334.1617 | 1 | 2.186908 | 421787.9 | 333.1544 | tri/oligope | Y |
| 82 | QrtoE | 342.2403 | 1 | 21.78168 | 140627.7 | 341.233 | tri/oligope | Y |
| 90 | QrtoE | 359.2301 | 1 | 2.481033 | 100917.2 | 358.2228 | tri/oligope | Y |

| | | | | | | | | |
|---|---|---|---|---|---|---|---|---|
| 94 | QrtoE | 363.1563 | 1 | 2.186908 | 105259.9 | 362.149 | tri/oligope | Y |
| 100 | QrtoE | 376.1724 | 1 | 2.165617 | 125082.6 | 375.1651 | tri/oligope | Y |
| 106 | QrtoE | 395.0398 | 1 | 1.606858 | 111804.3 | 394.0325 | tri/oligope | Y |
| 112 | QrtoE | 415.2123 | 1 | 36.463 | 163203.2 | 414.205 | tri/oligope | Y |
| 134 | QrtoE | 463.0272 | 1 | 1.606858 | 142717.4 | 462.02 | tri/oligope | Y |
| 141 | QrtoE | 479.3109 | 1 | 37.12123 | 132510.4 | 478.3036 | tri/oligope | Y |
| 144 | QrtoE | 488.252 | 1 | 19.10149 | 105005 | 487.2447 | tri/oligope | Y |
| 156 | QrtoE | 515.2921 | 1 | 37.12123 | 108499.7 | 514.2848 | tri/oligope | Y |
| 159 | QrtoE | 520.2784 | 1 | 21.12864 | 116750.3 | 519.2711 | tri/oligope | Y |
| 161 | QrtoE | 264.175 | 2 | 35.27298 | 100975.5 | 526.3355 | tri/oligope | Y |
| 165 | QrtoE | 531.0147 | 1 | 1.606858 | 177922 | 530.0074 | tri/oligope | Y |
| 166 | QrtoE | 531.3876 | 1 | 37.84973 | 100998.8 | 530.3804 | tri/oligope | Y |
| 174 | QrtoE | 274.168 | 2 | 35.93383 | 105220.2 | 546.3214 | tri/oligope | Y |
| 194 | QrtoE | 599.0026 | 1 | 1.606858 | 157075.4 | 597.9953 | tri/oligope | Y |
| 208 | QrtoE | 643.3321 | 1 | 20.74675 | 129722 | 642.3248 | oligopepti | Y |
| 218 | QrtoE | 666.9901 | 1 | 1.606858 | 156421.6 | 665.9828 | oligopepti | Y |
| 240 | QrtoE | 734.9777 | 1 | 1.606858 | 135488.6 | 733.9704 | oligopepti | Y |
| 258 | QrtoE | 802.9652 | 1 | 1.606858 | 117071.7 | 801.958 | oligopepti | Y |
| 296 | QrtoE | 974.8154 | 1 | 1.606858 | 203612.6 | 973.8082 | oligopepti | Y |
| 297 | QrtoE | 990.7897 | 1 | 1.617158 | 128964 | 989.7825 | oligopepti | Y |
| 338 | QrtoE | 223.1084 | 1 | 10.38584 | 169888.3 | 222.1011 | dipeptide | N |
| 339 | QrtoE | 223.1085 | 1 | 4.895408 | 239736.9 | 222.1012 | dipeptide | N |
| 340 | QrtoE | 226.9521 | 1 | 1.555258 | 381308.2 | 225.9448 | di/tripepti | N |
| 341 | QrtoE | 229.1555 | 1 | 8.972033 | 298663.2 | 228.1482 | di/tripepti | N |
| 342 | QrtoE | 229.1555 | 1 | 12.04122 | 123630.9 | 228.1482 | di/tripepti | N |
| 343 | QrtoE | 229.1555 | 1 | 4.916442 | 207513.6 | 228.1482 | di/tripepti | N |
| 344 | QrtoE | 231.1711 | 1 | 8.903342 | 316963.6 | 230.1638 | di/tripepti | N |
| 345 | QrtoE | 232.1296 | 1 | 2.186908 | 252498.3 | 231.1223 | di/tripepti | N |
| 346 | QrtoE | 233.1137 | 1 | 2.186908 | 292497.6 | 232.1064 | di/tripepti | N |
| 347 | QrtoE | 233.1504 | 1 | 4.028742 | 100192 | 232.1431 | di/tripepti | N |
| 348 | QrtoE | 237.1241 | 1 | 11.19468 | 141152 | 236.1169 | di/tripepti | N |
| 349 | QrtoE | 258.1106 | 1 | 1.755825 | 269123 | 257.1033 | di/tripepti | N |
| 350 | QrtoE | 263.14 | 1 | 14.43127 | 417894.5 | 262.1327 | di/tripepti | N |
| 351 | QrtoE | 276.1196 | 1 | 1.893792 | 107920.7 | 275.1123 | di/tripepti | N |
| 352 | QrtoE | 276.1353 | 1 | 17.17783 | 105673.8 | 275.1281 | di/tripepti | N |
| 353 | QrtoE | 276.1561 | 1 | 2.186908 | 375068.2 | 275.1488 | di/tripepti | N |
| 354 | QrtoE | 279.1712 | 1 | 19.89998 | 181279 | 278.1639 | di/tripepti | N |
| 355 | QrtoE | 281.1142 | 1 | 11.54666 | 274394.5 | 280.1069 | di/tripepti | N |
| 356 | QrtoE | 281.1507 | 1 | 5.890842 | 174932.8 | 280.1434 | di/tripepti | N |
| 357 | QrtoE | 288.1925 | 1 | 2.370658 | 120522.2 | 287.1853 | di/tripepti | N |
| 358 | QrtoE | 290.1352 | 1 | 2.186908 | 124080.6 | 289.1279 | di/tripepti | N |
| 359 | QrtoE | 290.1352 | 1 | 2.031792 | 121327.7 | 289.128 | di/tripepti | N |
| 360 | QrtoE | 290.1717 | 1 | 2.186908 | 449165.6 | 289.1644 | di/tripepti | N |
| 361 | QrtoE | 292.1301 | 1 | 16.42108 | 138839.2 | 291.1228 | di/tripepti | N |
| 362 | QrtoE | 294.9403 | 1 | 1.679075 | 345628.8 | 293.9331 | di/tripepti | N |
| 363 | QrtoE | 295.1301 | 1 | 12.61662 | 130893.8 | 294.1228 | di/tripepti | N |
| 364 | QrtoE | 295.1665 | 1 | 13.38839 | 142009.7 | 294.1592 | di/tripepti | N |
| 365 | QrtoE | 297.1091 | 1 | 2.667942 | 131528 | 296.1018 | di/tripepti | N |
| 366 | QrtoE | 302.1719 | 1 | 2.186908 | 248722.7 | 301.1647 | tri/oligope | N |
| 367 | QrtoE | 304.1874 | 1 | 2.186908 | 289764 | 303.1801 | tri/oligope | N |

| | | | | | | | | |
|---|---|---|---|---|---|---|---|---|
| 368 | QrtoE | 306.1302 | 1 | 1.944675 | 155628.5 | 305.1229 | tri/oligope | N |
| 369 | QrtoE | 306.1465 | 1 | 10.72147 | 105555.8 | 305.1392 | tri/oligope | N |
| 370 | QrtoE | 306.1667 | 1 | 2.186908 | 116134 | 305.1594 | tri/oligope | N |
| 371 | QrtoE | 311.1247 | 1 | 2.731008 | 116586.9 | 310.1174 | tri/oligope | N |
| 372 | QrtoE | 314.2087 | 1 | 12.96946 | 202064.4 | 313.2014 | tri/oligope | N |
| 373 | QrtoE | 316.188 | 1 | 13.03022 | 120041.9 | 315.1807 | tri/oligope | N |
| 374 | QrtoE | 316.2244 | 1 | 19.59053 | 103719.1 | 315.2171 | tri/oligope | N |
| 375 | QrtoE | 317.1828 | 1 | 2.241467 | 292327.9 | 316.1755 | tri/oligope | N |
| 376 | QrtoE | 318.2032 | 1 | 2.186908 | 129908.9 | 317.1959 | tri/oligope | N |
| 377 | QrtoE | 318.2037 | 1 | 14.58613 | 152682 | 317.1964 | tri/oligope | N |
| 378 | QrtoE | 320.146 | 1 | 2.176208 | 130559.9 | 319.1387 | tri/oligope | N |
| 379 | QrtoE | 328.2245 | 1 | 18.04731 | 245469.3 | 327.2172 | tri/oligope | N |
| 380 | QrtoE | 328.2245 | 1 | 16.8836 | 234405.1 | 327.2172 | tri/oligope | N |
| 381 | QrtoE | 329.1829 | 1 | 2.219667 | 100008.5 | 328.1756 | tri/oligope | N |
| 382 | QrtoE | 332.1829 | 1 | 11.54666 | 104661.8 | 331.1756 | tri/oligope | N |
| 383 | QrtoE | 333.1776 | 1 | 2.186908 | 263529.3 | 332.1703 | tri/oligope | N |
| 384 | QrtoE | 334.1411 | 1 | 13.73191 | 302490.2 | 333.1339 | tri/oligope | N |
| 385 | QrtoE | 334.1775 | 1 | 18.03326 | 153881.3 | 333.1702 | tri/oligope | N |
| 386 | QrtoE | 338.3429 | 1 | 39.26554 | 195080.8 | 337.3356 | tri/oligope | N |
| 387 | QrtoE | 342.24 | 1 | 21.24108 | 192859.8 | 341.2327 | tri/oligope | N |
| 388 | QrtoE | 342.2403 | 1 | 20.7391 | 109340.3 | 341.233 | tri/oligope | N |
| 389 | QrtoE | 343.1986 | 1 | 2.186908 | 195990.7 | 342.1913 | tri/oligope | N |
| 390 | QrtoE | 346.1987 | 1 | 17.05514 | 127823 | 345.1914 | tri/oligope | N |
| 391 | QrtoE | 347.1571 | 1 | 2.031792 | 107028.5 | 346.1498 | tri/oligope | N |
| 392 | QrtoE | 348.1411 | 1 | 2.020667 | 131345.5 | 347.1339 | tri/oligope | N |
| 393 | QrtoE | 348.1776 | 1 | 2.219667 | 554659.6 | 347.1703 | tri/oligope | N |
| 394 | QrtoE | 350.1727 | 1 | 17.49933 | 110356.7 | 349.1654 | tri/oligope | N |
| 395 | QrtoE | 360.2144 | 1 | 16.84708 | 137556.9 | 359.2072 | tri/oligope | N |
| 396 | QrtoE | 360.2145 | 1 | 20.45028 | 220723.1 | 359.2072 | tri/oligope | N |
| 397 | QrtoE | 361.1726 | 1 | 2.186908 | 286082.5 | 360.1654 | tri/oligope | N |
| 398 | QrtoE | 361.2091 | 1 | 2.1978 | 128564.8 | 360.2018 | tri/oligope | N |
| 399 | QrtoE | 362.1934 | 1 | 2.252433 | 277846.9 | 361.1861 | tri/oligope | N |
| 400 | QrtoE | 364.136 | 1 | 1.9346 | 119552.7 | 363.1287 | tri/oligope | N |
| 401 | QrtoE | 374.2043 | 1 | 2.186908 | 113500.6 | 373.197 | tri/oligope | N |
| 402 | QrtoE | 374.2301 | 1 | 14.33977 | 118061.8 | 373.2228 | tri/oligope | N |
| 403 | QrtoE | 389.2041 | 1 | 2.1978 | 156087.8 | 388.1968 | tri/oligope | N |
| 404 | QrtoE | 392.2196 | 1 | 20.16593 | 158439.9 | 391.2124 | tri/oligope | N |
| 405 | QrtoE | 403.2204 | 1 | 14.96088 | 107361.2 | 402.2131 | tri/oligope | N |
| 406 | QrtoE | 404.2147 | 1 | 2.186908 | 124783 | 403.2075 | tri/oligope | N |
| 407 | QrtoE | 405.1991 | 1 | 2.186908 | 220434.5 | 404.1919 | tri/oligope | N |
| 408 | QrtoE | 405.2619 | 1 | 37.26945 | 220961.6 | 404.2546 | tri/oligope | N |
| 409 | QrtoE | 413.267 | 1 | 36.76018 | 474236.7 | 412.2597 | tri/oligope | N |
| 410 | QrtoE | 413.2676 | 1 | 38.058 | 309688.2 | 412.2603 | tri/oligope | N |
| 411 | QrtoE | 425.1776 | 1 | 1.857008 | 249793.5 | 424.1703 | tri/oligope | N |
| 412 | QrtoE | 428.3378 | 1 | 37.27921 | 142941.9 | 427.3305 | tri/oligope | N |
| 413 | QrtoE | 432.2099 | 1 | 2.1978 | 125030.2 | 431.2026 | tri/oligope | N |
| 414 | QrtoE | 436.3433 | 1 | 35.93383 | 565948.6 | 435.336 | tri/oligope | N |
| 415 | QrtoE | 436.3438 | 1 | 0.022758 | 251182.7 | 435.3365 | tri/oligope | N |
| 416 | QrtoE | 441.2988 | 1 | 37.36306 | 174956 | 440.2915 | tri/oligope | N |
| 417 | QrtoE | 446.2254 | 1 | 2.186908 | 161051.9 | 445.2181 | tri/oligope | N |

| | | | | | | | | |
|---|---|---|---|---|---|---|---|---|
| 418 | QrtoE | 446.2626 | 1 | 14.91627 | 112016.4 | 445.2553 | tri/oligope | N |
| 419 | QrtoE | 449.288 | 1 | 37.23882 | 157316.5 | 448.2808 | tri/oligope | N |
| 420 | QrtoE | 462.2203 | 1 | 2.176208 | 114986.4 | 461.213 | tri/oligope | N |
| 421 | QrtoE | 463.298 | 1 | 37.55037 | 124958.7 | 462.2907 | tri/oligope | N |
| 422 | QrtoE | 469.3297 | 1 | 37.19075 | 105420.9 | 468.3225 | tri/oligope | N |
| 423 | QrtoE | 473.2259 | 1 | 13.4364 | 102122 | 472.2186 | tri/oligope | N |
| 424 | QrtoE | 475.2519 | 1 | 2.186908 | 112753.5 | 474.2447 | tri/oligope | N |
| 425 | QrtoE | 480.2467 | 1 | 17.4917 | 124179.4 | 479.2394 | tri/oligope | N |
| 426 | QrtoE | 488.8733 | 1 | 1.6376 | 156457.9 | 487.866 | tri/oligope | N |
| 427 | QrtoE | 252.1404 | 2 | 15.80574 | 100104.3 | 502.2662 | tri/oligope | N |
| 428 | QrtoE | 265.1535 | 2 | 2.533567 | 151443.1 | 528.2924 | tri/oligope | N |
| 429 | QrtoE | 531.2792 | 1 | 10.39986 | 103791.4 | 530.272 | tri/oligope | N |
| 430 | QrtoE | 533.455 | 1 | 37.39205 | 109131 | 532.4477 | tri/oligope | N |
| 431 | QrtoE | 572.2947 | 1 | 19.65016 | 157831 | 571.2874 | tri/oligope | N |
| 432 | QrtoE | 585.3268 | 1 | 22.37813 | 217780.6 | 584.3195 | tri/oligope | N |
| 433 | QrtoE | 307.1881 | 2 | 19.22523 | 269361.7 | 612.3616 | tri/oligope | N |
| 434 | QrtoE | 339.1857 | 2 | 19.30112 | 368105.5 | 676.3569 | oligopepti | N |
| 435 | QrtoE | 371.2175 | 2 | 20.00975 | 160071.8 | 740.4204 | oligopepti | N |
| 436 | QrtoE | 395.2053 | 2 | 16.55079 | 120378.5 | 788.396 | oligopepti | N |
| 437 | QrtoE | 424.2204 | 2 | 21.15673 | 103873.6 | 846.4263 | oligopepti | N |
| 438 | QrtoE | 460.6674 | 2 | 14.23063 | 171688 | 919.3202 | oligopepti | N |

| ID | Protein | mz | charge | retentionti | raw.abund | mass | type | Match |
|---|---|---|---|---|---|---|---|---|
| 2 | Wheat | 231.1711 | 1 | 5.370767 | 666071.8 | 230.1638 | di/tripepti | Y |
| 3 | Wheat | 231.1711 | 1 | 9.6898 | 652493.9 | 230.1638 | di/tripepti | Y |
| 4 | Wheat | 231.1712 | 1 | 12.05777 | 317444.2 | 230.1639 | di/tripepti | Y |
| 5 | Wheat | 233.1501 | 1 | 2.189133 | 143073.8 | 232.1428 | di/tripepti | Y |
| 11 | Wheat | 245.1867 | 1 | 16.43059 | 1817012 | 244.1794 | di/tripepti | Y |
| 13 | Wheat | 245.1866 | 1 | 14.54314 | 1382481 | 244.1793 | di/tripepti | Y |
| 14 | Wheat | 245.1869 | 1 | 17.68958 | 145748.2 | 244.1796 | di/tripepti | Y |
| 15 | Wheat | 246.1454 | 1 | 1.952483 | 373811.4 | 245.1381 | di/tripepti | Y |
| 15 | Wheat | 246.1454 | 1 | 2.189133 | 285700.7 | 245.1382 | di/tripepti | Y |
| 16 | Wheat | 246.1456 | 1 | 2.661 | 269029 | 245.1383 | di/tripepti | Y |
| 20 | Wheat | 247.1294 | 1 | 2.189133 | 334347.9 | 246.1222 | di/tripepti | Y |
| 22 | Wheat | 247.1298 | 1 | 4.715917 | 173441 | 246.1225 | di/tripepti | Y |
| 23 | Wheat | 249.1274 | 1 | 2.803825 | 742508.1 | 248.1201 | di/tripepti | Y |
| 24 | Wheat | 250.1784 | 1 | 34.86167 | 860455.5 | 249.1711 | di/tripepti | Y |
| 27 | Wheat | 253.1189 | 1 | 2.306058 | 336752.7 | 252.1116 | di/tripepti | Y |
| 30 | Wheat | 260.1611 | 1 | 2.189133 | 879771.8 | 259.1539 | di/tripepti | Y |
| 31 | Wheat | 260.1613 | 1 | 2.661 | 381089.1 | 259.154 | di/tripepti | Y |
| 34 | Wheat | 261.1452 | 1 | 2.189133 | 146087.8 | 260.1379 | di/tripepti | Y |
| 36 | Wheat | 261.1455 | 1 | 5.448867 | 240124.4 | 260.1382 | di/tripepti | Y |
| 38 | Wheat | 263.1434 | 1 | 11.52553 | 284511.4 | 262.1361 | di/tripepti | Y |
| 42 | Wheat | 269.1614 | 1 | 2.08405 | 164839.5 | 268.1541 | di/tripepti | Y |
| 47 | Wheat | 288.1928 | 1 | 16.07955 | 115631.9 | 287.1855 | di/tripepti | Y |
| 48 | Wheat | 290.1718 | 1 | 2.389692 | 333863.8 | 289.1645 | di/tripepti | Y |
| 50 | Wheat | 294.1456 | 1 | 2.341733 | 130713 | 293.1383 | di/tripepti | Y |
| 51 | Wheat | 295.1301 | 1 | 3.341692 | 158206.3 | 294.1228 | di/tripepti | Y |
| 57 | Wheat | 302.2085 | 1 | 19.86337 | 163987.7 | 301.2012 | tri/oligope | Y |
| 58 | Wheat | 303.1671 | 1 | 2.189133 | 137609.3 | 302.1599 | tri/oligope | Y |
| 64 | Wheat | 318.167 | 1 | 2.200575 | 190049 | 317.1597 | tri/oligope | Y |
| 70 | Wheat | 325.1139 | 1 | 1.733992 | 173264.8 | 324.1066 | tri/oligope | Y |
| 72 | Wheat | 331.1621 | 1 | 1.898875 | 294826 | 330.1548 | tri/oligope | Y |
| 73 | Wheat | 331.1663 | 1 | 2.847033 | 107225.5 | 330.159 | tri/oligope | Y |
| 79 | Wheat | 333.1572 | 1 | 11.04772 | 120039.1 | 332.1499 | tri/oligope | Y |
| 82 | Wheat | 342.2402 | 1 | 21.80854 | 133644.8 | 341.2329 | tri/oligope | Y |
| 86 | Wheat | 344.2557 | 1 | 20.19083 | 278917.1 | 343.2484 | tri/oligope | Y |
| 89 | Wheat | 358.2713 | 1 | 24.00683 | 1704230 | 357.264 | tri/oligope | Y |
| 90 | Wheat | 359.2301 | 1 | 2.481733 | 241363.9 | 358.2228 | tri/oligope | Y |
| 91 | Wheat | 360.1966 | 1 | 18.7834 | 121297.9 | 359.1893 | tri/oligope | Y |
| 112 | Wheat | 415.2126 | 1 | 36.47357 | 207144.3 | 414.2053 | tri/oligope | Y |
| 144 | Wheat | 488.2516 | 1 | 19.12215 | 192371.4 | 487.2443 | tri/oligope | Y |
| 159 | Wheat | 520.2789 | 1 | 21.12522 | 181091.8 | 519.2716 | tri/oligope | Y |
| 174 | Wheat | 274.1682 | 2 | 35.89189 | 104890.5 | 546.3218 | tri/oligope | Y |
| 208 | Wheat | 643.3327 | 1 | 20.7454 | 108512.3 | 642.3255 | oligopepti | Y |
| 297 | Wheat | 990.7902 | 1 | 1.616408 | 325861.3 | 989.7829 | oligopepti | Y |
| 306 | Wheat | 534.2818 | 2 | 22.60479 | 106278.1 | 1066.549 | oligopepti | Y |
| 338 | Wheat | 223.1085 | 1 | 5.04975 | 143675.2 | 222.1012 | dipeptide | N |
| 339 | Wheat | 223.1085 | 1 | 10.4576 | 100831.6 | 222.1012 | dipeptide | N |
| 340 | Wheat | 229.1555 | 1 | 9.015492 | 525740.7 | 228.1482 | di/tripepti | N |
| 341 | Wheat | 229.1556 | 1 | 11.89632 | 265757.9 | 228.1483 | di/tripepti | N |
| 342 | Wheat | 231.1711 | 1 | 8.946125 | 158765.8 | 230.1638 | di/tripepti | N |

| | | | | | | | | |
|---|---|---|---|---|---|---|---|---|
| 343 | Wheat | 232.1297 | 1 | 2.166758 | 150864.8 | 231.1224 | di/tripepti | N |
| 344 | Wheat | 233.1137 | 1 | 2.189133 | 150159.4 | 232.1064 | di/tripepti | N |
| 345 | Wheat | 244.1297 | 1 | 1.804925 | 321045.2 | 243.1224 | di/tripepti | N |
| 346 | Wheat | 244.1298 | 1 | 2.08405 | 173101.4 | 243.1225 | di/tripepti | N |
| 347 | Wheat | 252.9632 | 1 | 7.0306 | 187169.8 | 251.9559 | di/tripepti | N |
| 348 | Wheat | 263.1398 | 1 | 14.45386 | 617579.9 | 262.1325 | di/tripepti | N |
| 349 | Wheat | 275.1358 | 1 | 1.698542 | 381413.7 | 274.1285 | di/tripepti | N |
| 350 | Wheat | 276.1562 | 1 | 2.189133 | 149393.1 | 275.1489 | di/tripepti | N |
| 351 | Wheat | 279.1712 | 1 | 19.91159 | 237269 | 278.164 | di/tripepti | N |
| 352 | Wheat | 279.2325 | 1 | 37.00883 | 261662.2 | 278.2252 | di/tripepti | N |
| 353 | Wheat | 290.1717 | 1 | 2.189133 | 103218.5 | 289.1644 | di/tripepti | N |
| 354 | Wheat | 292.1302 | 1 | 9.447125 | 115922.8 | 291.123 | di/tripepti | N |
| 355 | Wheat | 292.151 | 1 | 1.929767 | 153365.5 | 291.1437 | di/tripepti | N |
| 356 | Wheat | 295.2276 | 1 | 37.374 | 130564.4 | 294.2203 | di/tripepti | N |
| 357 | Wheat | 297.1181 | 1 | 1.698542 | 112820.6 | 296.1108 | di/tripepti | N |
| 358 | Wheat | 297.2433 | 1 | 36.13935 | 107629.6 | 296.236 | di/tripepti | N |
| 359 | Wheat | 300.2178 | 1 | 35.24678 | 106008.4 | 299.2105 | di/tripepti | N |
| 360 | Wheat | 302.1719 | 1 | 2.282167 | 158436.5 | 301.1647 | tri/oligope | N |
| 361 | Wheat | 302.172 | 1 | 2.493417 | 311180.8 | 301.1647 | tri/oligope | N |
| 362 | Wheat | 302.2086 | 1 | 7.277817 | 219556.9 | 301.2013 | tri/oligope | N |
| 363 | Wheat | 304.1878 | 1 | 6.612292 | 234280.8 | 303.1806 | tri/oligope | N |
| 364 | Wheat | 304.2491 | 1 | 37.56288 | 103072.9 | 303.2418 | tri/oligope | N |
| 365 | Wheat | 313.2381 | 1 | 35.80724 | 174658.4 | 312.2308 | tri/oligope | N |
| 366 | Wheat | 314.2087 | 1 | 13.00219 | 144195.6 | 313.2015 | tri/oligope | N |
| 367 | Wheat | 315.1671 | 1 | 2.072142 | 141060.5 | 314.1598 | tri/oligope | N |
| 368 | Wheat | 315.2538 | 1 | 36.13935 | 339272.3 | 314.2466 | tri/oligope | N |
| 369 | Wheat | 316.1881 | 1 | 13.05888 | 125323.2 | 315.1809 | tri/oligope | N |
| 370 | Wheat | 316.2241 | 1 | 19.08293 | 382923.7 | 315.2168 | tri/oligope | N |
| 371 | Wheat | 317.1829 | 1 | 2.189133 | 186687.9 | 316.1756 | tri/oligope | N |
| 372 | Wheat | 317.183 | 1 | 2.423558 | 267195.3 | 316.1757 | tri/oligope | N |
| 373 | Wheat | 318.1495 | 1 | 3.652992 | 324419.7 | 317.1423 | tri/oligope | N |
| 374 | Wheat | 318.2033 | 1 | 2.341733 | 170113.1 | 317.196 | tri/oligope | N |
| 375 | Wheat | 318.2034 | 1 | 14.59489 | 186303.6 | 317.1961 | tri/oligope | N |
| 376 | Wheat | 318.2649 | 1 | 37.83672 | 128218.3 | 317.2576 | tri/oligope | N |
| 377 | Wheat | 328.2244 | 1 | 17.63673 | 448632.9 | 327.2172 | tri/oligope | N |
| 378 | Wheat | 328.2245 | 1 | 16.8859 | 195103.8 | 327.2172 | tri/oligope | N |
| 379 | Wheat | 328.2343 | 1 | 15.48358 | 103910.1 | 327.227 | tri/oligope | N |
| 380 | Wheat | 329.1831 | 1 | 2.544158 | 222642.1 | 328.1758 | tri/oligope | N |
| 381 | Wheat | 330.2037 | 1 | 11.89632 | 466463.3 | 329.1964 | tri/oligope | N |
| 382 | Wheat | 331.1985 | 1 | 2.40165 | 462315.8 | 330.1913 | tri/oligope | N |
| 383 | Wheat | 331.1989 | 1 | 6.36225 | 141151.6 | 330.1916 | tri/oligope | N |
| 384 | Wheat | 332.1827 | 1 | 9.713975 | 786844.8 | 331.1754 | tri/oligope | N |
| 385 | Wheat | 332.2191 | 1 | 18.59551 | 466377.4 | 331.2118 | tri/oligope | N |
| 386 | Wheat | 333.1778 | 1 | 2.189133 | 250241.8 | 332.1705 | tri/oligope | N |
| 387 | Wheat | 334.1774 | 1 | 18.03428 | 433548.4 | 333.1701 | tri/oligope | N |
| 388 | Wheat | 336.1932 | 1 | 23.0508 | 105320.2 | 335.1859 | tri/oligope | N |
| 389 | Wheat | 337.2358 | 1 | 36.13935 | 142209.1 | 336.2285 | tri/oligope | N |
| 390 | Wheat | 342.2401 | 1 | 18.74299 | 360119 | 341.2328 | tri/oligope | N |
| 391 | Wheat | 342.2403 | 1 | 21.29544 | 159116.5 | 341.233 | tri/oligope | N |
| 392 | Wheat | 343.1987 | 1 | 2.189133 | 114446.5 | 342.1914 | tri/oligope | N |

| | | | | | | | | |
|---|---|---|---|---|---|---|---|---|
| 393 | Wheat | 350.1725 | 1 | 17.50723 | 200268.6 | 349.1652 | tri/oligope | N |
| 394 | Wheat | 351.0965 | 1 | 30.84455 | 125197.2 | 350.0893 | tri/oligope | N |
| 395 | Wheat | 357.2143 | 1 | 2.211933 | 255006.3 | 356.207 | tri/oligope | N |
| 396 | Wheat | 357.2145 | 1 | 2.544158 | 731017.1 | 356.2072 | tri/oligope | N |
| 397 | Wheat | 357.2148 | 1 | 12.48922 | 206727.4 | 356.2076 | tri/oligope | N |
| 398 | Wheat | 357.2148 | 1 | 13.26202 | 196346 | 356.2076 | tri/oligope | N |
| 399 | Wheat | 357.2149 | 1 | 14.98073 | 133218.8 | 356.2076 | tri/oligope | N |
| 400 | Wheat | 358.1732 | 1 | 1.929767 | 181459.8 | 357.1659 | tri/oligope | N |
| 401 | Wheat | 358.2961 | 1 | 36.2718 | 164990.5 | 357.2889 | tri/oligope | N |
| 402 | Wheat | 360.2144 | 1 | 17.6224 | 106532.5 | 359.2071 | tri/oligope | N |
| 403 | Wheat | 361.2092 | 1 | 2.200575 | 119846.8 | 360.202 | tri/oligope | N |
| 404 | Wheat | 362.2091 | 1 | 21.35706 | 139629.9 | 361.2019 | tri/oligope | N |
| 405 | Wheat | 362.2123 | 1 | 18.17774 | 132028.2 | 361.205 | tri/oligope | N |
| 406 | Wheat | 372.1888 | 1 | 1.952483 | 478441.3 | 371.1815 | tri/oligope | N |
| 407 | Wheat | 372.1888 | 1 | 2.189133 | 1054658 | 371.1816 | tri/oligope | N |
| 408 | Wheat | 373.2458 | 1 | 18.81772 | 208859.5 | 372.2386 | tri/oligope | N |
| 409 | Wheat | 373.2462 | 1 | 3.161967 | 152504.3 | 372.2389 | tri/oligope | N |
| 410 | Wheat | 373.2462 | 1 | 10.8438 | 129612.4 | 372.2389 | tri/oligope | N |
| 411 | Wheat | 373.3799 | 1 | 37.59178 | 128892.1 | 372.3726 | tri/oligope | N |
| 412 | Wheat | 374.2044 | 1 | 2.189133 | 277246.7 | 373.1971 | tri/oligope | N |
| 413 | Wheat | 376.2275 | 1 | 20.34591 | 1033480 | 375.2202 | tri/oligope | N |
| 414 | Wheat | 378.1709 | 1 | 10.53301 | 314982.8 | 377.1637 | tri/oligope | N |
| 415 | Wheat | 380.1832 | 1 | 11.38504 | 841076.7 | 379.1759 | tri/oligope | N |
| 416 | Wheat | 384.3117 | 1 | 37.32066 | 100712.2 | 383.3045 | tri/oligope | N |
| 417 | Wheat | 385.2461 | 1 | 19.16666 | 136298.1 | 384.2388 | tri/oligope | N |
| 418 | Wheat | 389.2041 | 1 | 2.189133 | 173705 | 388.1968 | tri/oligope | N |
| 419 | Wheat | 391.1991 | 1 | 17.2908 | 371465.6 | 390.1918 | tri/oligope | N |
| 420 | Wheat | 391.1992 | 1 | 14.68162 | 612563.4 | 390.1919 | tri/oligope | N |
| 421 | Wheat | 391.2203 | 1 | 3.010458 | 529232.6 | 390.213 | tri/oligope | N |
| 422 | Wheat | 391.2859 | 1 | 1.278858 | 140136 | 390.2787 | tri/oligope | N |
| 423 | Wheat | 403.2204 | 1 | 11.78113 | 109819.4 | 402.2131 | tri/oligope | N |
| 424 | Wheat | 404.1786 | 1 | 1.813992 | 103594.3 | 403.1714 | tri/oligope | N |
| 425 | Wheat | 413.2674 | 1 | 36.74438 | 667595.9 | 412.2602 | tri/oligope | N |
| 426 | Wheat | 415.2357 | 1 | 27.25548 | 291771.1 | 414.2284 | tri/oligope | N |
| 427 | Wheat | 415.3904 | 1 | 37.03262 | 106951.5 | 414.3831 | tri/oligope | N |
| 428 | Wheat | 428.252 | 1 | 15.45089 | 130038.8 | 427.2447 | tri/oligope | N |
| 429 | Wheat | 428.2521 | 1 | 16.89828 | 191382.9 | 427.2449 | tri/oligope | N |
| 430 | Wheat | 428.3379 | 1 | 37.27349 | 172705.5 | 427.3306 | tri/oligope | N |
| 431 | Wheat | 431.2512 | 1 | 14.22363 | 1320132 | 430.2439 | tri/oligope | N |
| 432 | Wheat | 220.1192 | 2 | 2.024483 | 124258.6 | 438.2238 | tri/oligope | N |
| 433 | Wheat | 441.3091 | 1 | 24.46216 | 108320.3 | 440.3018 | tri/oligope | N |
| 434 | Wheat | 442.2672 | 1 | 19.01901 | 821764.8 | 441.2599 | tri/oligope | N |
| 435 | Wheat | 443.226 | 1 | 2.189133 | 151337 | 442.2187 | tri/oligope | N |
| 436 | Wheat | 443.2878 | 1 | 17.37238 | 383505.6 | 442.2805 | tri/oligope | N |
| 437 | Wheat | 444.2463 | 1 | 2.189133 | 276065.7 | 443.239 | tri/oligope | N |
| 438 | Wheat | 445.3038 | 1 | 25.31478 | 196396 | 444.2965 | tri/oligope | N |
| 439 | Wheat | 446.2052 | 1 | 20.65447 | 117262.1 | 445.1979 | tri/oligope | N |
| 440 | Wheat | 446.208 | 1 | 2.317925 | 486242.9 | 445.2008 | tri/oligope | N |
| 441 | Wheat | 447.2255 | 1 | 17.07339 | 144960.7 | 446.2183 | tri/oligope | N |
| 442 | Wheat | 447.2621 | 1 | 23.57239 | 122806 | 446.2548 | tri/oligope | N |

| | | | | | | | | |
|---|---|---|---|---|---|---|---|---|
| 443 | Wheat | 449.2412 | 1 | 22.39721 | 252534.9 | 448.2339 | tri/oligope | N |
| 444 | Wheat | 454.2677 | 1 | 14.49811 | 210567.1 | 453.2604 | tri/oligope | N |
| 445 | Wheat | 454.2677 | 1 | 17.36105 | 108123.2 | 453.2605 | tri/oligope | N |
| 446 | Wheat | 455.3247 | 1 | 26.09888 | 272265.6 | 454.3174 | tri/oligope | N |
| 447 | Wheat | 455.3249 | 1 | 26.81138 | 123678.6 | 454.3176 | tri/oligope | N |
| 448 | Wheat | 456.2835 | 1 | 13.74982 | 441219.5 | 455.2762 | tri/oligope | N |
| 449 | Wheat | 456.2835 | 1 | 12.9017 | 218125.1 | 455.2763 | tri/oligope | N |
| 450 | Wheat | 459.2209 | 1 | 2.024483 | 474682.9 | 458.2136 | tri/oligope | N |
| 451 | Wheat | 461.2364 | 1 | 2.189133 | 134329 | 460.2291 | tri/oligope | N |
| 452 | Wheat | 461.2443 | 1 | 20.25217 | 229034.9 | 460.237 | tri/oligope | N |
| 453 | Wheat | 469.2416 | 1 | 2.200575 | 727148.3 | 468.2343 | tri/oligope | N |
| 454 | Wheat | 470.2993 | 1 | 15.03004 | 109214.5 | 469.292 | tri/oligope | N |
| 455 | Wheat | 471.2575 | 1 | 2.282167 | 699397.9 | 470.2502 | tri/oligope | N |
| 456 | Wheat | 472.242 | 1 | 12.89518 | 615723.1 | 471.2348 | tri/oligope | N |
| 457 | Wheat | 472.3642 | 1 | 37.23528 | 109615.2 | 471.357 | tri/oligope | N |
| 458 | Wheat | 475.2521 | 1 | 2.189133 | 270159.6 | 474.2448 | tri/oligope | N |
| 459 | Wheat | 478.2313 | 1 | 18.56444 | 274920 | 477.224 | tri/oligope | N |
| 460 | Wheat | 478.2314 | 1 | 10.83716 | 187399 | 477.2242 | tri/oligope | N |
| 461 | Wheat | 485.2728 | 1 | 2.189133 | 289038.7 | 484.2656 | tri/oligope | N |
| 462 | Wheat | 485.2735 | 1 | 15.45774 | 647214.1 | 484.2662 | tri/oligope | N |
| 463 | Wheat | 485.2737 | 1 | 13.70414 | 359215.9 | 484.2664 | tri/oligope | N |
| 464 | Wheat | 485.2737 | 1 | 12.6889 | 126047.5 | 484.2664 | tri/oligope | N |
| 465 | Wheat | 486.2578 | 1 | 13.40017 | 100818.6 | 485.2505 | tri/oligope | N |
| 466 | Wheat | 488.2523 | 1 | 20.36193 | 249082.7 | 487.245 | tri/oligope | N |
| 467 | Wheat | 489.2677 | 1 | 2.189133 | 184980 | 488.2605 | tri/oligope | N |
| 468 | Wheat | 489.2685 | 1 | 13.77724 | 124873.5 | 488.2612 | tri/oligope | N |
| 469 | Wheat | 489.2724 | 1 | 19.77329 | 387546.3 | 488.2651 | tri/oligope | N |
| 470 | Wheat | 489.2726 | 1 | 17.68958 | 516003.5 | 488.2654 | tri/oligope | N |
| 471 | Wheat | 489.3578 | 1 | 36.45119 | 107311.8 | 488.3505 | tri/oligope | N |
| 472 | Wheat | 494.2731 | 1 | 2.189133 | 175446.7 | 493.2658 | tri/oligope | N |
| 473 | Wheat | 247.6404 | 2 | 2.200575 | 570464.7 | 493.2662 | tri/oligope | N |
| 474 | Wheat | 500.2473 | 1 | 2.072142 | 696182.8 | 499.2401 | tri/oligope | N |
| 475 | Wheat | 502.2634 | 1 | 2.189133 | 177156.3 | 501.2562 | tri/oligope | N |
| 476 | Wheat | 504.284 | 1 | 22.75258 | 181550.2 | 503.2767 | tri/oligope | N |
| 477 | Wheat | 505.2675 | 1 | 21.91673 | 102107.4 | 504.2602 | tri/oligope | N |
| 478 | Wheat | 511.1763 | 1 | 1.698542 | 119708.8 | 510.1691 | tri/oligope | N |
| 479 | Wheat | 512.3465 | 1 | 26.21672 | 152896.6 | 511.3392 | tri/oligope | N |
| 480 | Wheat | 519.2578 | 1 | 16.43059 | 1428516 | 518.2505 | tri/oligope | N |
| 481 | Wheat | 519.2579 | 1 | 17.992 | 112976.9 | 518.2507 | tri/oligope | N |
| 482 | Wheat | 260.1327 | 2 | 16.39715 | 355146.7 | 518.2509 | tri/oligope | N |
| 483 | Wheat | 529.2629 | 1 | 2.189133 | 164063.9 | 528.2556 | tri/oligope | N |
| 484 | Wheat | 529.2995 | 1 | 15.53552 | 475527.6 | 528.2922 | tri/oligope | N |
| 485 | Wheat | 530.3206 | 1 | 21.39409 | 131369 | 529.3133 | tri/oligope | N |
| 486 | Wheat | 530.3334 | 1 | 36.67673 | 101064.8 | 529.3261 | tri/oligope | N |
| 487 | Wheat | 533.2399 | 1 | 2.211933 | 131580.6 | 532.2326 | tri/oligope | N |
| 488 | Wheat | 535.253 | 1 | 14.75852 | 295984.3 | 534.2457 | tri/oligope | N |
| 489 | Wheat | 535.253 | 1 | 13.18105 | 159693 | 534.2457 | tri/oligope | N |
| 490 | Wheat | 539.3202 | 1 | 15.54583 | 641726.8 | 538.313 | tri/oligope | N |
| 491 | Wheat | 270.1639 | 2 | 15.54583 | 537548.5 | 538.3132 | tri/oligope | N |
| 492 | Wheat | 541.3356 | 1 | 22.84101 | 1561098 | 540.3283 | tri/oligope | N |

| | | | | | | | | |
|---|---|---|---|---|---|---|---|---|
| 493 | Wheat | 543.2466 | 1 | 16.70721 | 804114.4 | 542.2393 | tri/oligope | N |
| 494 | Wheat | 543.2947 | 1 | 26.16046 | 268496.6 | 542.2874 | tri/oligope | N |
| 495 | Wheat | 545.2948 | 1 | 14.2172 | 246477.6 | 544.2875 | tri/oligope | N |
| 496 | Wheat | 547.2743 | 1 | 13.9547 | 107045.5 | 546.267 | tri/oligope | N |
| 497 | Wheat | 556.2741 | 1 | 2.189133 | 336390.4 | 555.2669 | tri/oligope | N |
| 498 | Wheat | 278.6593 | 2 | 13.36025 | 362918.8 | 555.304 | tri/oligope | N |
| 499 | Wheat | 556.3114 | 1 | 13.36661 | 245003.2 | 555.3041 | tri/oligope | N |
| 500 | Wheat | 559.3109 | 1 | 15.86658 | 306824.4 | 558.3036 | tri/oligope | N |
| 501 | Wheat | 566.2948 | 1 | 2.200575 | 195615.8 | 565.2876 | tri/oligope | N |
| 502 | Wheat | 567.8937 | 1 | 1.616408 | 141594.5 | 566.8864 | tri/oligope | N |
| 503 | Wheat | 285.1875 | 2 | 22.05974 | 203605.9 | 568.3605 | tri/oligope | N |
| 504 | Wheat | 569.368 | 1 | 22.05974 | 284404.9 | 568.3608 | tri/oligope | N |
| 505 | Wheat | 291.667 | 2 | 15.9513 | 143288.8 | 581.3195 | tri/oligope | N |
| 506 | Wheat | 582.3269 | 1 | 15.9513 | 155971.2 | 581.3196 | tri/oligope | N |
| 507 | Wheat | 584.3426 | 1 | 19.2453 | 180708.2 | 583.3353 | tri/oligope | N |
| 508 | Wheat | 585.3263 | 1 | 20.26817 | 466602.7 | 584.319 | tri/oligope | N |
| 509 | Wheat | 587.2799 | 1 | 2.166758 | 208261.3 | 586.2726 | tri/oligope | N |
| 510 | Wheat | 588.3419 | 1 | 20.59639 | 315050.1 | 587.3347 | tri/oligope | N |
| 511 | Wheat | 597.3006 | 1 | 2.189133 | 474023.8 | 596.2933 | tri/oligope | N |
| 512 | Wheat | 299.1541 | 2 | 2.200575 | 112194 | 596.2936 | tri/oligope | N |
| 513 | Wheat | 598.358 | 1 | 14.68162 | 171679.8 | 597.3507 | tri/oligope | N |
| 514 | Wheat | 299.6827 | 2 | 14.68162 | 255721.8 | 597.3508 | tri/oligope | N |
| 515 | Wheat | 598.8385 | 1 | 1.6396 | 103937.8 | 597.8313 | tri/oligope | N |
| 516 | Wheat | 300.1515 | 2 | 14.61748 | 148261 | 598.2885 | tri/oligope | N |
| 517 | Wheat | 599.3162 | 1 | 2.317925 | 165411.7 | 598.3089 | tri/oligope | N |
| 518 | Wheat | 300.1621 | 2 | 2.3299 | 102096.9 | 598.3096 | tri/oligope | N |
| 519 | Wheat | 600.2684 | 1 | 18.87723 | 385201.8 | 599.2611 | tri/oligope | N |
| 520 | Wheat | 601.3318 | 1 | 2.189133 | 106272.4 | 600.3245 | tri/oligope | N |
| 521 | Wheat | 601.3369 | 1 | 23.87813 | 148753.6 | 600.3296 | tri/oligope | N |
| 522 | Wheat | 301.1721 | 2 | 23.89423 | 128714.8 | 600.3297 | tri/oligope | N |
| 523 | Wheat | 605.3164 | 1 | 14.23678 | 209380.3 | 604.3091 | tri/oligope | N |
| 524 | Wheat | 303.6488 | 2 | 16.26488 | 596243.9 | 605.2831 | tri/oligope | N |
| 525 | Wheat | 606.2903 | 1 | 16.27448 | 499820.9 | 605.2831 | tri/oligope | N |
| 526 | Wheat | 303.6542 | 2 | 2.095875 | 136376.8 | 605.2939 | tri/oligope | N |
| 527 | Wheat | 612.2798 | 1 | 19.58876 | 101095 | 611.2726 | tri/oligope | N |
| 528 | Wheat | 613.3326 | 1 | 14.35231 | 357241.6 | 612.3253 | tri/oligope | N |
| 529 | Wheat | 307.17 | 2 | 14.35921 | 294243.2 | 612.3254 | tri/oligope | N |
| 530 | Wheat | 613.333 | 1 | 13.52729 | 109205.2 | 612.3258 | tri/oligope | N |
| 531 | Wheat | 307.1702 | 2 | 13.51691 | 196226.8 | 612.3258 | tri/oligope | N |
| 532 | Wheat | 307.1702 | 2 | 12.02376 | 114766.5 | 612.3259 | tri/oligope | N |
| 533 | Wheat | 615.3123 | 1 | 2.189133 | 199222.2 | 614.3051 | oligopepti | N |
| 534 | Wheat | 616.3112 | 1 | 20.36193 | 707698.7 | 615.3039 | oligopepti | N |
| 535 | Wheat | 616.3114 | 1 | 17.232 | 235901.3 | 615.3041 | oligopepti | N |
| 536 | Wheat | 616.3114 | 1 | 18.19935 | 352588.1 | 615.3041 | oligopepti | N |
| 537 | Wheat | 308.6593 | 2 | 17.19618 | 299018.8 | 615.3041 | oligopepti | N |
| 538 | Wheat | 308.6594 | 2 | 18.18406 | 351267.7 | 615.3042 | oligopepti | N |
| 539 | Wheat | 309.6671 | 2 | 15.63941 | 150764.4 | 617.3196 | oligopepti | N |
| 540 | Wheat | 618.3269 | 1 | 15.64733 | 118151.2 | 617.3196 | oligopepti | N |
| 541 | Wheat | 622.2852 | 1 | 14.66581 | 191051.5 | 621.2779 | oligopepti | N |
| 542 | Wheat | 313.176 | 2 | 2.189133 | 112583.4 | 624.3374 | oligopepti | N |

| | | | | | | | | |
|---|---|---|---|---|---|---|---|---|
| 543 | Wheat | 628.3065 | 1 | 2.189133 | 362314.6 | 627.2993 | oligopepti | N |
| 544 | Wheat | 315.1566 | 2 | 14.58643 | 1071877 | 628.2987 | oligopepti | N |
| 545 | Wheat | 315.628 | 2 | 16.24348 | 123795 | 629.2415 | oligopepti | N |
| 546 | Wheat | 632.3433 | 1 | 24.75281 | 148220.8 | 631.336 | oligopepti | N |
| 547 | Wheat | 633.3228 | 1 | 2.938692 | 160933.8 | 632.3155 | oligopepti | N |
| 548 | Wheat | 635.8814 | 1 | 1.616408 | 145587.3 | 634.8741 | oligopepti | N |
| 549 | Wheat | 319.1491 | 2 | 14.35231 | 131183.5 | 636.2837 | oligopepti | N |
| 550 | Wheat | 647.317 | 1 | 16.4071 | 476929.6 | 646.3097 | oligopepti | N |
| 551 | Wheat | 324.1622 | 2 | 16.4071 | 1237269 | 646.3099 | oligopepti | N |
| 552 | Wheat | 324.6545 | 2 | 17.05995 | 146983.3 | 647.2945 | oligopepti | N |
| 553 | Wheat | 652.3374 | 1 | 23.26678 | 208181.8 | 651.3301 | oligopepti | N |
| 554 | Wheat | 326.7246 | 2 | 15.54583 | 250310.4 | 651.4347 | oligopepti | N |
| 555 | Wheat | 655.3798 | 1 | 17.6224 | 114142.5 | 654.3725 | oligopepti | N |
| 556 | Wheat | 328.1936 | 2 | 17.6224 | 228989.7 | 654.3727 | oligopepti | N |
| 557 | Wheat | 328.693 | 2 | 26.44917 | 774176.2 | 655.3715 | oligopepti | N |
| 558 | Wheat | 656.379 | 1 | 26.44187 | 756193.9 | 655.3717 | oligopepti | N |
| 559 | Wheat | 329.1725 | 2 | 21.87318 | 692177.9 | 656.3305 | oligopepti | N |
| 560 | Wheat | 657.3379 | 1 | 21.85995 | 208129.5 | 656.3306 | oligopepti | N |
| 561 | Wheat | 668.6209 | 1 | 37.49818 | 190141.1 | 667.6136 | oligopepti | N |
| 562 | Wheat | 670.3537 | 1 | 2.189133 | 308069.2 | 669.3465 | oligopepti | N |
| 563 | Wheat | 335.6806 | 2 | 2.223333 | 116033.3 | 669.3465 | oligopepti | N |
| 564 | Wheat | 336.1412 | 2 | 16.43059 | 313631.2 | 670.2679 | oligopepti | N |
| 565 | Wheat | 672.338 | 1 | 26.08382 | 307226.8 | 671.3308 | oligopepti | N |
| 566 | Wheat | 336.6727 | 2 | 26.09888 | 142789.7 | 671.3308 | oligopepti | N |
| 567 | Wheat | 341.1517 | 2 | 21.85995 | 102231 | 680.2889 | oligopepti | N |
| 568 | Wheat | 341.2017 | 2 | 15.90067 | 149730.3 | 680.3888 | oligopepti | N |
| 569 | Wheat | 684.3329 | 1 | 2.189133 | 184202.7 | 683.3256 | oligopepti | N |
| 570 | Wheat | 343.1359 | 2 | 16.43059 | 150965.2 | 684.2572 | oligopepti | N |
| 571 | Wheat | 344.1783 | 2 | 18.24824 | 263569 | 686.342 | oligopepti | N |
| 572 | Wheat | 688.3695 | 1 | 23.96225 | 215349.9 | 687.3623 | oligopepti | N |
| 573 | Wheat | 692.3274 | 1 | 14.6904 | 254333.3 | 691.3201 | oligopepti | N |
| 574 | Wheat | 346.6676 | 2 | 14.71693 | 133914.9 | 691.3207 | oligopepti | N |
| 575 | Wheat | 346.6913 | 2 | 2.5241 | 278957.1 | 691.3681 | oligopepti | N |
| 576 | Wheat | 348.2096 | 2 | 22.89433 | 102082.7 | 694.4047 | oligopepti | N |
| 577 | Wheat | 348.2153 | 2 | 15.20952 | 100542.4 | 694.4161 | oligopepti | N |
| 578 | Wheat | 350.694 | 2 | 22.4133 | 230535.7 | 699.3735 | oligopepti | N |
| 579 | Wheat | 703.3434 | 1 | 18.79538 | 966370.7 | 702.3361 | oligopepti | N |
| 580 | Wheat | 352.1755 | 2 | 18.80696 | 186023.8 | 702.3365 | oligopepti | N |
| 581 | Wheat | 708.5136 | 1 | 37.36333 | 841671.1 | 707.5063 | oligopepti | N |
| 582 | Wheat | 355.6968 | 2 | 17.5135 | 198417 | 709.3791 | oligopepti | N |
| 583 | Wheat | 355.6969 | 2 | 16.82491 | 135712.5 | 709.3793 | oligopepti | N |
| 584 | Wheat | 356.1473 | 2 | 16.92817 | 157478 | 710.2801 | oligopepti | N |
| 585 | Wheat | 357.1863 | 2 | 21.08953 | 132958.5 | 712.358 | oligopepti | N |
| 586 | Wheat | 364.1544 | 2 | 18.79538 | 177207.6 | 726.2943 | oligopepti | N |
| 587 | Wheat | 364.1915 | 2 | 2.3299 | 108193.2 | 726.3685 | oligopepti | N |
| 588 | Wheat | 728.3275 | 1 | 19.09289 | 241918.1 | 727.3202 | oligopepti | N |
| 589 | Wheat | 364.6915 | 2 | 22.87449 | 109053.9 | 727.3685 | oligopepti | N |
| 590 | Wheat | 729.3955 | 1 | 20.33465 | 101898 | 728.3882 | oligopepti | N |
| 591 | Wheat | 365.2017 | 2 | 20.35348 | 292306.3 | 728.3889 | oligopepti | N |
| 592 | Wheat | 371.1995 | 2 | 14.647 | 242381.9 | 740.3845 | oligopepti | N |

| | | | | | | | | |
|---|---|---|---|---|---|---|---|---|
| 593 | Wheat | 371.1998 | 2 | 13.87792 | 298785.6 | 740.3851 | oligopepti | N |
| 594 | Wheat | 742.3492 | 1 | 2.189133 | 102689.7 | 741.3419 | oligopepti | N |
| 595 | Wheat | 372.6889 | 2 | 19.5179 | 307268.3 | 743.3633 | oligopepti | N |
| 596 | Wheat | 372.6889 | 2 | 20.3054 | 273835.8 | 743.3633 | oligopepti | N |
| 597 | Wheat | 372.689 | 2 | 18.16271 | 280954.5 | 743.3635 | oligopepti | N |
| 598 | Wheat | 375.6996 | 2 | 2.472525 | 119161 | 749.3847 | oligopepti | N |
| 599 | Wheat | 376.2069 | 2 | 25.81879 | 107051 | 750.3991 | oligopepti | N |
| 600 | Wheat | 376.7019 | 2 | 20.5231 | 628331.6 | 751.3893 | oligopepti | N |
| 601 | Wheat | 377.1897 | 2 | 15.05987 | 100714.6 | 752.3648 | oligopepti | N |
| 602 | Wheat | 377.2044 | 2 | 2.189133 | 114303.3 | 752.3942 | oligopepti | N |
| 603 | Wheat | 756.3916 | 1 | 14.70883 | 106843.9 | 755.3843 | oligopepti | N |
| 604 | Wheat | 378.6996 | 2 | 14.71693 | 185900.8 | 755.3846 | oligopepti | N |
| 605 | Wheat | 384.7177 | 2 | 16.04879 | 217117.6 | 767.4209 | oligopepti | N |
| 606 | Wheat | 386.7103 | 2 | 16.61057 | 175171.3 | 771.406 | oligopepti | N |
| 607 | Wheat | 386.7103 | 2 | 15.70572 | 247907.6 | 771.406 | oligopepti | N |
| 608 | Wheat | 388.1919 | 2 | 16.54799 | 166706.8 | 774.3692 | oligopepti | N |
| 609 | Wheat | 389.7281 | 2 | 18.76152 | 231472 | 777.4416 | oligopepti | N |
| 610 | Wheat | 392.2047 | 2 | 2.306058 | 229331.6 | 782.3948 | oligopepti | N |
| 611 | Wheat | 394.7079 | 2 | 12.6889 | 118919.8 | 787.4012 | oligopepti | N |
| 612 | Wheat | 396.736 | 2 | 19.81438 | 310104.4 | 791.4574 | oligopepti | N |
| 613 | Wheat | 397.7489 | 2 | 19.01901 | 1638227 | 793.4833 | oligopepti | N |
| 614 | Wheat | 399.2128 | 2 | 15.73055 | 694779.1 | 796.4111 | oligopepti | N |
| 615 | Wheat | 400.702 | 2 | 19.62874 | 548263 | 799.3894 | oligopepti | N |
| 616 | Wheat | 800.3968 | 1 | 19.62874 | 220442.9 | 799.3895 | oligopepti | N |
| 617 | Wheat | 400.7024 | 2 | 24.40623 | 135786.7 | 799.3902 | oligopepti | N |
| 618 | Wheat | 404.185 | 2 | 2.317925 | 118664.5 | 806.3555 | oligopepti | N |
| 619 | Wheat | 406.2387 | 2 | 20.33465 | 195664.7 | 810.4629 | oligopepti | N |
| 620 | Wheat | 408.2076 | 2 | 18.51542 | 193811.5 | 814.4007 | oligopepti | N |
| 621 | Wheat | 408.7179 | 2 | 20.63295 | 272643.8 | 815.4213 | oligopepti | N |
| 622 | Wheat | 410.721 | 2 | 14.20928 | 196196.6 | 819.4274 | oligopepti | N |
| 623 | Wheat | 411.1917 | 2 | 15.72163 | 135450.9 | 820.3689 | oligopepti | N |
| 624 | Wheat | 412.2388 | 2 | 22.53373 | 521375.6 | 822.463 | oligopepti | N |
| 625 | Wheat | 412.239 | 2 | 19.31338 | 164808.4 | 822.4634 | oligopepti | N |
| 626 | Wheat | 412.6809 | 2 | 19.63558 | 114284.3 | 823.3473 | oligopepti | N |
| 627 | Wheat | 416.2049 | 2 | 18.86862 | 395090 | 830.3952 | oligopepti | N |
| 628 | Wheat | 416.205 | 2 | 16.21573 | 119263.5 | 830.3954 | oligopepti | N |
| 629 | Wheat | 416.205 | 2 | 20.19791 | 216267.4 | 830.3954 | oligopepti | N |
| 630 | Wheat | 419.7262 | 2 | 14.20339 | 756009.2 | 837.4379 | oligopepti | N |
| 631 | Wheat | 420.1766 | 2 | 16.46486 | 195854.1 | 838.3386 | oligopepti | N |
| 632 | Wheat | 421.2154 | 2 | 19.495 | 266485.4 | 840.4162 | oligopepti | N |
| 633 | Wheat | 421.2157 | 2 | 21.30374 | 162879.2 | 840.4169 | oligopepti | N |
| 634 | Wheat | 421.7133 | 2 | 22.14129 | 100600.3 | 841.4121 | oligopepti | N |
| 635 | Wheat | 421.7257 | 2 | 20.59639 | 220860.4 | 841.4369 | oligopepti | N |
| 636 | Wheat | 423.2129 | 2 | 18.28563 | 375254.2 | 844.4113 | oligopepti | N |
| 637 | Wheat | 424.6872 | 2 | 16.53934 | 116119.8 | 847.3599 | oligopepti | N |
| 638 | Wheat | 426.2256 | 2 | 18.88708 | 149466.7 | 850.4367 | oligopepti | N |
| 639 | Wheat | 428.1996 | 2 | 17.97488 | 268560.5 | 854.3846 | oligopepti | N |
| 640 | Wheat | 429.2128 | 2 | 16.58339 | 114335.8 | 856.4111 | oligopepti | N |
| 641 | Wheat | 429.2313 | 2 | 22.58882 | 133337.9 | 856.448 | oligopepti | N |
| 642 | Wheat | 870.789 | 1 | 1.6396 | 113043.8 | 869.7818 | oligopepti | N |

| | | | | | | | | |
|---|---|---|---|---|---|---|---|---|
| 643 | Wheat | 436.7182 | 2 | 17.95167 | 833911 | 871.4218 | oligopepti | N |
| 644 | Wheat | 444.2183 | 2 | 23.84257 | 411345.3 | 886.422 | oligopepti | N |
| 645 | Wheat | 887.4294 | 1 | 23.84885 | 103265 | 886.4221 | oligopepti | N |
| 646 | Wheat | 446.2703 | 2 | 23.9556 | 165754.1 | 890.5261 | oligopepti | N |
| 647 | Wheat | 447.7574 | 2 | 17.47848 | 467439.3 | 893.5003 | oligopepti | N |
| 648 | Wheat | 449.1795 | 2 | 16.86275 | 123779 | 896.3445 | oligopepti | N |
| 649 | Wheat | 459.7208 | 2 | 18.7834 | 658196.6 | 917.4271 | oligopepti | N |
| 650 | Wheat | 460.6671 | 2 | 14.26665 | 169924.2 | 919.3196 | oligopepti | N |
| 651 | Wheat | 463.2627 | 2 | 26.12448 | 154960.3 | 924.5109 | oligopepti | N |
| 652 | Wheat | 468.2527 | 2 | 15.73985 | 151845.2 | 934.4908 | oligopepti | N |
| 653 | Wheat | 470.2522 | 2 | 22.15009 | 459573.7 | 938.4899 | oligopepti | N |
| 654 | Wheat | 471.6998 | 2 | 18.79538 | 138735 | 941.3851 | oligopepti | N |
| 655 | Wheat | 471.7397 | 2 | 21.02148 | 217689.3 | 941.4648 | oligopepti | N |
| 656 | Wheat | 473.7424 | 2 | 17.47848 | 109510.6 | 945.4703 | oligopepti | N |
| 657 | Wheat | 476.2682 | 2 | 19.47519 | 379594.4 | 950.5219 | oligopepti | N |
| 658 | Wheat | 477.7576 | 2 | 23.07053 | 596393.3 | 953.5007 | oligopepti | N |
| 659 | Wheat | 479.2554 | 2 | 16.0377 | 126481.6 | 956.4962 | oligopepti | N |
| 660 | Wheat | 480.2344 | 2 | 20.19791 | 192942 | 958.4543 | oligopepti | N |
| 661 | Wheat | 480.7434 | 2 | 19.01206 | 107312.4 | 959.4723 | oligopepti | N |
| 662 | Wheat | 485.2448 | 2 | 19.28934 | 624467.8 | 968.4751 | oligopepti | N |
| 663 | Wheat | 487.2422 | 2 | 19.84766 | 267392.7 | 972.4699 | oligopepti | N |
| 664 | Wheat | 491.7422 | 2 | 16.29992 | 911125.2 | 981.4698 | oligopepti | N |
| 665 | Wheat | 505.271 | 2 | 19.17697 | 115631.4 | 1008.527 | oligopepti | N |
| 666 | Wheat | 506.779 | 2 | 24.44628 | 415070.7 | 1011.543 | oligopepti | N |
| 667 | Wheat | 510.2999 | 2 | 22.51147 | 159614.5 | 1018.585 | oligopepti | N |
| 668 | Wheat | 513.7867 | 2 | 25.31478 | 441640.5 | 1025.559 | oligopepti | N |
| 669 | Wheat | 516.2637 | 2 | 23.40999 | 162552.1 | 1030.513 | oligopepti | N |
| 670 | Wheat | 523.7503 | 2 | 18.80696 | 330986.5 | 1045.486 | oligopepti | N |
| 671 | Wheat | 524.7947 | 2 | 18.62988 | 140753.2 | 1047.575 | oligopepti | N |
| 672 | Wheat | 355.5309 | 3 | 18.06803 | 187312.9 | 1063.571 | oligopepti | N |
| 673 | Wheat | 541.7874 | 2 | 21.67182 | 223329.1 | 1081.56 | oligopepti | N |
| 674 | Wheat | 541.7874 | 2 | 23.43084 | 159664.4 | 1081.56 | oligopepti | N |
| 675 | Wheat | 364.8663 | 3 | 18.59551 | 363299 | 1091.577 | oligopepti | N |
| 676 | Wheat | 548.8702 | 2 | 1.627925 | 117107.5 | 1095.726 | oligopepti | N |
| 677 | Wheat | 549.2742 | 2 | 19.61592 | 182450 | 1096.534 | oligopepti | N |
| 678 | Wheat | 552.7463 | 2 | 17.85991 | 128433.4 | 1103.478 | oligopepti | N |
| 679 | Wheat | 557.2907 | 2 | 21.79439 | 160892.6 | 1112.567 | oligopepti | N |
| 680 | Wheat | 559.8028 | 2 | 18.90566 | 202553.9 | 1117.591 | oligopepti | N |
| 681 | Wheat | 561.7335 | 2 | 15.59232 | 100407 | 1121.452 | oligopepti | N |
| 682 | Wheat | 563.3218 | 2 | 27.99865 | 115393.8 | 1124.629 | oligopepti | N |
| 683 | Wheat | 574.283 | 2 | 23.16401 | 245753.4 | 1146.551 | oligopepti | N |
| 684 | Wheat | 582.8063 | 2 | 16.07075 | 184543.3 | 1163.598 | oligopepti | N |
| 685 | Wheat | 588.314 | 2 | 16.43059 | 183824.9 | 1174.614 | oligopepti | N |
| 686 | Wheat | 596.3115 | 2 | 17.50723 | 220328.2 | 1190.608 | oligopepti | N |
| 687 | Wheat | 597.801 | 2 | 20.22518 | 401616.1 | 1193.587 | oligopepti | N |
| 688 | Wheat | 597.801 | 2 | 19.82518 | 194385.1 | 1193.588 | oligopepti | N |
| 689 | Wheat | 1194.752 | 1 | 1.616408 | 119313.9 | 1193.745 | oligopepti | N |
| 690 | Wheat | 598.3301 | 2 | 26.85128 | 120798.4 | 1194.646 | oligopepti | N |
| 691 | Wheat | 599.7985 | 2 | 19.46938 | 658090.3 | 1197.582 | oligopepti | N |
| 692 | Wheat | 408.1875 | 3 | 19.45954 | 141319.2 | 1221.541 | oligopepti | N |

| | | | | | | | | |
|---|---|---|---|---|---|---|---|---|
| 693 | Wheat | 613.3039 | 2 | 18.80696 | 390177.6 | 1224.593 | oligopepti( | N |
| 694 | Wheat | 619.8197 | 2 | 16.27448 | 166253.1 | 1237.625 | oligopepti( | N |
| 695 | Wheat | 624.8473 | 2 | 1.6396 | 119724.5 | 1247.68 | oligopepti( | N |
| 696 | Wheat | 627.3096 | 2 | 21.1766 | 100156.6 | 1252.605 | oligopepti( | N |
| 697 | Wheat | 638.3123 | 2 | 22.51147 | 111815.9 | 1274.61 | oligopepti( | N |
| 698 | Wheat | 646.8566 | 2 | 25.29243 | 209216.7 | 1291.699 | oligopepti( | N |
| 699 | Wheat | 661.8306 | 2 | 19.5179 | 137709.1 | 1321.647 | oligopepti( | N |
| 700 | Wheat | 696.85 | 2 | 14.9669 | 613452.2 | 1391.686 | oligopepti( | N |
| 701 | Wheat | 701.8601 | 2 | 17.44475 | 160552.1 | 1401.706 | oligopepti( | N |
| 702 | Wheat | 705.347 | 2 | 20.15254 | 387314.3 | 1408.679 | oligopepti( | N |
| 703 | Wheat | 472.8885 | 3 | 14.97448 | 155407.7 | 1415.644 | oligopepti( | N |
| 704 | Wheat | 718.3728 | 2 | 21.85995 | 331598.3 | 1434.731 | oligopepti( | N |
| 705 | Wheat | 726.8269 | 2 | 1.616408 | 109221.2 | 1451.639 | oligopepti( | N |
| 706 | Wheat | 732.3763 | 2 | 17.0188 | 250674.8 | 1462.738 | oligopepti( | N |
| 707 | Wheat | 758.3688 | 2 | 25.17736 | 118855.2 | 1514.723 | oligopepti( | N |
| 708 | Wheat | 776.3843 | 2 | 19.62874 | 119545.5 | 1550.754 | oligopepti( | N |
| 709 | Wheat | 794.8146 | 2 | 1.616408 | 106007 | 1587.615 | oligopepti( | N |
| 710 | Wheat | 804.4051 | 2 | 26.43405 | 153814.7 | 1606.796 | oligopepti( | N |
| 711 | Wheat | 896.7969 | 2 | 1.627925 | 111893 | 1791.579 | oligopepti( | N |
| 712 | Wheat | 922.8025 | 2 | 1.616408 | 395674.2 | 1843.59 | oligopepti( | N |
| 713 | Wheat | 641.9863 | 3 | 25.52629 | 180353.8 | 1922.937 | oligopepti( | N |
| 714 | Wheat | 678.3501 | 3 | 25.77462 | 168916.5 | 2032.028 | oligopepti( | N |
| 715 | Wheat | 683.3396 | 3 | 23.9556 | 158281 | 2046.997 | oligopepti( | N |
| 716 | Wheat | 1024.783 | 2 | 1.616408 | 162879.4 | 2047.552 | oligopepti( | N |
| 717 | Wheat | 684.6727 | 3 | 23.2907 | 102251.8 | 2050.996 | oligopepti( | N |
| 718 | Wheat | 692.3533 | 3 | 24.38337 | 108215.4 | 2074.038 | oligopepti( | N |
| 719 | Wheat | 702.9715 | 3 | 23.11389 | 137250.6 | 2105.893 | oligopepti( | N |
| 720 | Wheat | 717.0238 | 3 | 24.4759 | 716765.5 | 2148.05 | oligopepti( | N |
| 721 | Wheat | 724.6817 | 3 | 24.4759 | 150359.9 | 2171.023 | oligopepti( | N |
| 722 | Wheat | 1092.77 | 2 | 1.616408 | 137542.8 | 2183.526 | oligopepti( | N |
| 723 | Wheat | 745.6577 | 3 | 23.0574 | 113453.7 | 2233.951 | oligopepti( | N |
| 724 | Wheat | 753.387 | 3 | 25.68066 | 138182.5 | 2257.139 | oligopepti( | N |
| 725 | Wheat | 1160.757 | 2 | 1.605 | 103111.3 | 2319.5 | oligopepti( | N |
| 726 | Wheat | 778.7185 | 3 | 26.11214 | 117383.6 | 2333.134 | oligopepti( | N |
| 727 | Wheat | 821.4131 | 3 | 27.24807 | 155953.6 | 2461.217 | oligopepti( | N |
| 728 | Wheat | 833.4161 | 3 | 24.35449 | 103407.9 | 2497.227 | oligopepti( | N |
| 729 | Wheat | 911.1304 | 3 | 28.61151 | 171073.2 | 2730.369 | oligopepti( | N |

| ID | Protein | mz | charge | retentiont | raw.abund | mass | type | Match |
|---|---|---|---|---|---|---|---|---|
| 2 | Corn | 231.1712 | 1 | 5.396017 | 158307.3 | 230.1639 | di/tripepti | Y |
| 3 | Corn | 231.1712 | 1 | 9.687583 | 1375387 | 230.1639 | di/tripepti | Y |
| 4 | Corn | 231.1709 | 1 | 12.05802 | 504631 | 230.1636 | di/tripepti | Y |
| 5 | Corn | 233.1501 | 1 | 2.193917 | 628819.8 | 232.1428 | di/tripepti | Y |
| 6 | Corn | 233.1502 | 1 | 2.79675 | 348150.9 | 232.1429 | di/tripepti | Y |
| 11 | Corn | 245.1867 | 1 | 16.43508 | 690402.1 | 244.1794 | di/tripepti | Y |
| 12 | Corn | 245.1869 | 1 | 15.90157 | 119932.8 | 244.1796 | di/tripepti | Y |
| 13 | Corn | 245.1868 | 1 | 14.56174 | 236287.6 | 244.1795 | di/tripepti | Y |
| 14 | Corn | 245.187 | 1 | 17.69018 | 706226.6 | 244.1797 | di/tripepti | Y |
| 15 | Corn | 246.1454 | 1 | 1.949375 | 508336.1 | 245.1381 | di/tripepti | Y |
| 15 | Corn | 246.1454 | 1 | 2.182567 | 297023.3 | 245.1382 | di/tripepti | Y |
| 16 | Corn | 246.1455 | 1 | 2.664908 | 643265.2 | 245.1382 | di/tripepti | Y |
| 17 | Corn | 246.1457 | 1 | 3.409842 | 260183 | 245.1384 | di/tripepti | Y |
| 20 | Corn | 247.1295 | 1 | 2.182567 | 166699.4 | 246.1222 | di/tripepti | Y |
| 22 | Corn | 247.1298 | 1 | 4.743933 | 227432.5 | 246.1225 | di/tripepti | Y |
| 24 | Corn | 250.1784 | 1 | 34.86191 | 816965.1 | 249.1711 | di/tripepti | Y |
| 25 | Corn | 250.1784 | 1 | 37.25553 | 148185.1 | 249.1711 | di/tripepti | Y |
| 27 | Corn | 253.1189 | 1 | 2.489325 | 1404604 | 252.1116 | di/tripepti | Y |
| 28 | Corn | 253.1191 | 1 | 9.327025 | 214991.2 | 252.1118 | di/tripepti | Y |
| 30 | Corn | 260.161 | 1 | 2.193917 | 2704490 | 259.1537 | di/tripepti | Y |
| 32 | Corn | 260.1615 | 1 | 3.7618 | 1519225 | 259.1542 | di/tripepti | Y |
| 34 | Corn | 261.1452 | 1 | 2.182567 | 162573.2 | 260.1379 | di/tripepti | Y |
| 36 | Corn | 261.1455 | 1 | 5.516133 | 207755.8 | 260.1382 | di/tripepti | Y |
| 37 | Corn | 262.1405 | 1 | 1.961492 | 263812.8 | 261.1332 | di/tripepti | Y |
| 39 | Corn | 263.1971 | 1 | 2.193917 | 106503.8 | 262.1898 | di/tripepti | Y |
| 40 | Corn | 265.1557 | 1 | 16.28073 | 177791.6 | 264.1484 | di/tripepti | Y |
| 42 | Corn | 269.1614 | 1 | 2.182567 | 481306 | 268.1541 | di/tripepti | Y |
| 46 | Corn | 281.1507 | 1 | 7.137492 | 117675.9 | 280.1434 | di/tripepti | Y |
| 47 | Corn | 288.1929 | 1 | 15.94198 | 197663.6 | 287.1856 | di/tripepti | Y |
| 48 | Corn | 290.1718 | 1 | 2.718258 | 542986.1 | 289.1645 | di/tripepti | Y |
| 50 | Corn | 294.1457 | 1 | 2.337883 | 141425.2 | 293.1384 | di/tripepti | Y |
| 51 | Corn | 295.1301 | 1 | 3.356242 | 121027.8 | 294.1228 | di/tripepti | Y |
| 52 | Corn | 295.1664 | 1 | 16.42814 | 1690716 | 294.1592 | di/tripepti | Y |
| 57 | Corn | 302.2084 | 1 | 19.85123 | 255201.7 | 301.2011 | tri/oligope | Y |
| 58 | Corn | 303.1671 | 1 | 2.182567 | 266528.3 | 302.1598 | tri/oligope | Y |
| 64 | Corn | 318.1669 | 1 | 2.205492 | 130550.4 | 317.1596 | tri/oligope | Y |
| 65 | Corn | 318.1672 | 1 | 9.149433 | 324312.8 | 317.16 | tri/oligope | Y |
| 73 | Corn | 331.1662 | 1 | 2.848308 | 252746.3 | 330.1589 | tri/oligope | Y |
| 80 | Corn | 334.1619 | 1 | 2.193917 | 108305 | 333.1546 | tri/oligope | Y |
| 94 | Corn | 363.156 | 1 | 2.193917 | 133228.8 | 362.1487 | tri/oligope | Y |
| 112 | Corn | 415.2125 | 1 | 36.47488 | 160288.5 | 414.2052 | tri/oligope | Y |
| 141 | Corn | 479.3111 | 1 | 37.21091 | 118698.6 | 478.3038 | tri/oligope | Y |
| 144 | Corn | 488.252 | 1 | 19.12538 | 190063.4 | 487.2447 | tri/oligope | Y |
| 151 | Corn | 500.2889 | 1 | 25.18949 | 152810 | 499.2817 | tri/oligope | Y |
| 159 | Corn | 520.2785 | 1 | 21.18292 | 102050.8 | 519.2713 | tri/oligope | Y |
| 297 | Corn | 990.7901 | 1 | 1.610975 | 351365.3 | 989.7828 | oligopepti | Y |
| 338 | Corn | 229.1555 | 1 | 9.030592 | 231401.1 | 228.1482 | di/tripepti | N |
| 339 | Corn | 229.1556 | 1 | 11.96377 | 4366027 | 228.1483 | di/tripepti | N |
| 340 | Corn | 232.1298 | 1 | 1.912475 | 123972.1 | 231.1225 | di/tripepti | N |

| | | | | | | | | |
|---|---|---|---|---|---|---|---|---|
| 341 | Corn | 233.1504 | 1 | 4.096017 | 332920.7 | 232.1431 | di/tripepti | N |
| 342 | Corn | 237.1242 | 1 | 11.20312 | 789891.2 | 236.1169 | di/tripepti | N |
| 343 | Corn | 237.1242 | 1 | 4.635675 | 122518.3 | 236.1169 | di/tripepti | N |
| 344 | Corn | 261.1203 | 1 | 1.69265 | 173963.5 | 260.113 | di/tripepti | N |
| 345 | Corn | 263.1399 | 1 | 14.46187 | 691946.9 | 262.1327 | di/tripepti | N |
| 346 | Corn | 263.1433 | 1 | 13.13114 | 232306.1 | 262.136 | di/tripepti | N |
| 347 | Corn | 269.1138 | 1 | 2.182567 | 156450.1 | 268.1065 | di/tripepti | N |
| 348 | Corn | 269.114 | 1 | 2.337883 | 119950.9 | 268.1067 | di/tripepti | N |
| 349 | Corn | 274.1404 | 1 | 2.079408 | 380530 | 273.1331 | di/tripepti | N |
| 350 | Corn | 274.1767 | 1 | 2.489325 | 722792.5 | 273.1694 | di/tripepti | N |
| 351 | Corn | 274.1768 | 1 | 2.275783 | 281117.8 | 273.1695 | di/tripepti | N |
| 352 | Corn | 274.1769 | 1 | 2.894767 | 534145.2 | 273.1696 | di/tripepti | N |
| 353 | Corn | 274.177 | 1 | 9.687583 | 241279.1 | 273.1697 | di/tripepti | N |
| 354 | Corn | 274.1771 | 1 | 6.849683 | 335612.8 | 273.1698 | di/tripepti | N |
| 355 | Corn | 275.1357 | 1 | 1.704392 | 472490.6 | 274.1284 | di/tripepti | N |
| 356 | Corn | 276.1561 | 1 | 2.182567 | 254731.3 | 275.1488 | di/tripepti | N |
| 357 | Corn | 278.1176 | 1 | 2.182567 | 100959.8 | 277.1103 | di/tripepti | N |
| 358 | Corn | 279.135 | 1 | 3.783342 | 175937.2 | 278.1277 | di/tripepti | N |
| 359 | Corn | 279.138 | 1 | 2.182567 | 108668 | 278.1307 | di/tripepti | N |
| 360 | Corn | 279.1707 | 1 | 19.90277 | 335838.2 | 278.1634 | di/tripepti | N |
| 361 | Corn | 279.1715 | 1 | 21.02548 | 473756.7 | 278.1642 | di/tripepti | N |
| 362 | Corn | 281.1506 | 1 | 6.155742 | 150799.8 | 280.1434 | di/tripepti | N |
| 363 | Corn | 288.1561 | 1 | 2.182567 | 230096.8 | 287.1488 | di/tripepti | N |
| 364 | Corn | 288.1925 | 1 | 2.240483 | 109420.2 | 287.1853 | di/tripepti | N |
| 365 | Corn | 290.1717 | 1 | 2.193917 | 303802.6 | 289.1645 | di/tripepti | N |
| 366 | Corn | 297.118 | 1 | 1.704392 | 131204.9 | 296.1107 | di/tripepti | N |
| 367 | Corn | 300.1929 | 1 | 12.07529 | 139994.4 | 299.1856 | di/tripepti | N |
| 368 | Corn | 300.193 | 1 | 4.027008 | 133843.1 | 299.1857 | di/tripepti | N |
| 369 | Corn | 300.193 | 1 | 9.846067 | 255261.1 | 299.1857 | di/tripepti | N |
| 370 | Corn | 301.1515 | 1 | 2.121342 | 190535.9 | 300.1442 | di/tripepti | N |
| 371 | Corn | 302.2085 | 1 | 10.28011 | 403052.7 | 301.2013 | tri/oligope | N |
| 372 | Corn | 302.2086 | 1 | 11.14295 | 229869.5 | 301.2014 | tri/oligope | N |
| 373 | Corn | 304.1875 | 1 | 2.263875 | 584293.4 | 303.1802 | tri/oligope | N |
| 374 | Corn | 304.1876 | 1 | 2.584183 | 436108.2 | 303.1803 | tri/oligope | N |
| 375 | Corn | 310.1404 | 1 | 2.182567 | 545970.4 | 309.1332 | tri/oligope | N |
| 376 | Corn | 313.1765 | 1 | 2.182567 | 107425.8 | 312.1692 | tri/oligope | N |
| 377 | Corn | 316.1881 | 1 | 13.02993 | 5200033 | 315.1808 | tri/oligope | N |
| 378 | Corn | 316.2244 | 1 | 16.36875 | 129879.2 | 315.2171 | tri/oligope | N |
| 379 | Corn | 317.1828 | 1 | 2.193917 | 666098.9 | 316.1755 | tri/oligope | N |
| 380 | Corn | 317.183 | 1 | 2.760175 | 388664.2 | 316.1757 | tri/oligope | N |
| 381 | Corn | 318.2037 | 1 | 15.90739 | 245150.2 | 317.1965 | tri/oligope | N |
| 382 | Corn | 328.2245 | 1 | 18.06751 | 208407.2 | 327.2172 | tri/oligope | N |
| 383 | Corn | 328.2245 | 1 | 17.80022 | 289791.7 | 327.2173 | tri/oligope | N |
| 384 | Corn | 328.2247 | 1 | 16.85918 | 148257.3 | 327.2174 | tri/oligope | N |
| 385 | Corn | 328.2343 | 1 | 15.48559 | 114213.2 | 327.227 | tri/oligope | N |
| 386 | Corn | 329.1828 | 1 | 2.216942 | 205386.3 | 328.1756 | tri/oligope | N |
| 387 | Corn | 331.1985 | 1 | 2.193917 | 305414.6 | 330.1913 | tri/oligope | N |
| 388 | Corn | 331.1986 | 1 | 2.729283 | 442041.4 | 330.1913 | tri/oligope | N |
| 389 | Corn | 332.2193 | 1 | 19.43598 | 186956.4 | 331.212 | tri/oligope | N |
| 390 | Corn | 333.1778 | 1 | 2.022767 | 249091.8 | 332.1706 | tri/oligope | N |

| | | | | | | | | |
|---|---|---|---|---|---|---|---|---|
| 391 | Corn | 342.2402 | 1 | 19.85123 | 12301501 | 341.233 | tri/oligope | N |
| 392 | Corn | 343.1986 | 1 | 2.182567 | 258281.5 | 342.1913 | tri/oligope | N |
| 393 | Corn | 343.1988 | 1 | 13.38618 | 1065880 | 342.1916 | tri/oligope | N |
| 394 | Corn | 344.2465 | 1 | 19.85123 | 245701.3 | 343.2392 | tri/oligope | N |
| 395 | Corn | 345.2145 | 1 | 14.61305 | 110405.2 | 344.2073 | tri/oligope | N |
| 396 | Corn | 346.1732 | 1 | 1.912475 | 118890.3 | 345.1659 | tri/oligope | N |
| 397 | Corn | 346.2352 | 1 | 13.55542 | 165911.2 | 345.2279 | tri/oligope | N |
| 398 | Corn | 348.1777 | 1 | 2.193917 | 126454.7 | 347.1704 | tri/oligope | N |
| 399 | Corn | 350.1726 | 1 | 12.39429 | 162205.6 | 349.1653 | tri/oligope | N |
| 400 | Corn | 350.1728 | 1 | 10.41498 | 132505.9 | 349.1655 | tri/oligope | N |
| 401 | Corn | 350.1756 | 1 | 2.534883 | 132069.9 | 349.1683 | tri/oligope | N |
| 402 | Corn | 351.0965 | 1 | 30.86048 | 107204.5 | 350.0893 | tri/oligope | N |
| 403 | Corn | 357.2146 | 1 | 15.02083 | 1760807 | 356.2073 | tri/oligope | N |
| 404 | Corn | 359.2305 | 1 | 15.67595 | 182320.5 | 358.2232 | tri/oligope | N |
| 405 | Corn | 359.2306 | 1 | 18.51229 | 129561.8 | 358.2233 | tri/oligope | N |
| 406 | Corn | 361.2091 | 1 | 2.205492 | 182828.8 | 360.2018 | tri/oligope | N |
| 407 | Corn | 361.2093 | 1 | 2.374408 | 144989.5 | 360.2021 | tri/oligope | N |
| 408 | Corn | 362.1681 | 1 | 1.880458 | 146314.3 | 361.1608 | tri/oligope | N |
| 409 | Corn | 366.1675 | 1 | 10.4215 | 150271.2 | 365.1602 | tri/oligope | N |
| 410 | Corn | 372.1888 | 1 | 2.182567 | 129363.5 | 371.1815 | tri/oligope | N |
| 411 | Corn | 372.1889 | 1 | 2.055175 | 142540.3 | 371.1817 | tri/oligope | N |
| 412 | Corn | 373.2463 | 1 | 18.82 | 113259.8 | 372.239 | tri/oligope | N |
| 413 | Corn | 374.2043 | 1 | 2.205492 | 490812.8 | 373.197 | tri/oligope | N |
| 414 | Corn | 376.188 | 1 | 14.71723 | 105097 | 375.1807 | tri/oligope | N |
| 415 | Corn | 376.2245 | 1 | 22.17804 | 4468277 | 375.2172 | tri/oligope | N |
| 416 | Corn | 376.2246 | 1 | 25.40067 | 4434877 | 375.2173 | tri/oligope | N |
| 417 | Corn | 378.231 | 1 | 25.40067 | 118667.2 | 377.2237 | tri/oligope | N |
| 418 | Corn | 378.2311 | 1 | 22.17804 | 111013.7 | 377.2238 | tri/oligope | N |
| 419 | Corn | 387.225 | 1 | 2.751233 | 551356 | 386.2177 | tri/oligope | N |
| 420 | Corn | 387.2254 | 1 | 6.291717 | 113909 | 386.2181 | tri/oligope | N |
| 421 | Corn | 387.2978 | 1 | 19.85924 | 114396.3 | 386.2906 | tri/oligope | N |
| 422 | Corn | 388.1838 | 1 | 2.159617 | 255587.7 | 387.1765 | tri/oligope | N |
| 423 | Corn | 388.1838 | 1 | 1.979608 | 118602.8 | 387.1765 | tri/oligope | N |
| 424 | Corn | 388.2199 | 1 | 2.193917 | 716871.1 | 387.2126 | tri/oligope | N |
| 425 | Corn | 388.2205 | 1 | 3.606933 | 135160.6 | 387.2132 | tri/oligope | N |
| 426 | Corn | 388.2205 | 1 | 7.719858 | 108796 | 387.2133 | tri/oligope | N |
| 427 | Corn | 389.2043 | 1 | 2.47845 | 104492.1 | 388.197 | tri/oligope | N |
| 428 | Corn | 389.2047 | 1 | 8.24745 | 147489.7 | 388.1975 | tri/oligope | N |
| 429 | Corn | 391.22 | 1 | 2.350358 | 134382.7 | 390.2127 | tri/oligope | N |
| 430 | Corn | 391.286 | 1 | 0.020133 | 609909.3 | 390.2787 | tri/oligope | N |
| 431 | Corn | 392.2193 | 1 | 19.74492 | 114768.4 | 391.212 | tri/oligope | N |
| 432 | Corn | 393.1818 | 1 | 11.25363 | 166960.9 | 392.1745 | tri/oligope | N |
| 433 | Corn | 399.2616 | 1 | 16.40899 | 118159.5 | 398.2543 | tri/oligope | N |
| 434 | Corn | 399.2618 | 1 | 23.52373 | 469979.7 | 398.2545 | tri/oligope | N |
| 435 | Corn | 400.22 | 1 | 2.193917 | 119275 | 399.2127 | tri/oligope | N |
| 436 | Corn | 402.1993 | 1 | 2.182567 | 299310.5 | 401.192 | tri/oligope | N |
| 437 | Corn | 403.2206 | 1 | 13.83088 | 111375.7 | 402.2133 | tri/oligope | N |
| 438 | Corn | 407.1938 | 1 | 2.444425 | 191975.2 | 406.1865 | tri/oligope | N |
| 439 | Corn | 407.1941 | 1 | 11.14295 | 279338.1 | 406.1868 | tri/oligope | N |
| 440 | Corn | 407.1941 | 1 | 18.31378 | 368751.4 | 406.1868 | tri/oligope | N |

| | | | | | | | | |
|---|---|---|---|---|---|---|---|---|
| 441 | Corn | 413.2773 | 1 | 23.34236 | 194554.4 | 412.27 | tri/oligope | N |
| 442 | Corn | 413.2778 | 1 | 23.60378 | 348084.7 | 412.2705 | tri/oligope | N |
| 443 | Corn | 414.2359 | 1 | 2.522992 | 176117.8 | 413.2286 | tri/oligope | N |
| 444 | Corn | 414.236 | 1 | 14.65268 | 409086.4 | 413.2287 | tri/oligope | N |
| 445 | Corn | 415.2568 | 1 | 14.16772 | 261213.8 | 414.2495 | tri/oligope | N |
| 446 | Corn | 415.2569 | 1 | 16.67753 | 114975.6 | 414.2496 | tri/oligope | N |
| 447 | Corn | 418.2308 | 1 | 2.829542 | 633497.9 | 417.2236 | tri/oligope | N |
| 448 | Corn | 425.2774 | 1 | 19.41129 | 168611 | 424.2701 | tri/oligope | N |
| 449 | Corn | 427.2928 | 1 | 21.15314 | 1076153 | 426.2855 | tri/oligope | N |
| 450 | Corn | 428.2519 | 1 | 14.30353 | 750787.4 | 427.2446 | tri/oligope | N |
| 451 | Corn | 429.2727 | 1 | 23.96705 | 114953.6 | 428.2654 | tri/oligope | N |
| 452 | Corn | 430.2306 | 1 | 2.228658 | 167267.3 | 429.2233 | tri/oligope | N |
| 453 | Corn | 430.2314 | 1 | 13.94725 | 310405.9 | 429.2242 | tri/oligope | N |
| 454 | Corn | 430.267 | 1 | 14.68648 | 1103708 | 429.2598 | tri/oligope | N |
| 455 | Corn | 433.2464 | 1 | 23.9258 | 213494.6 | 432.2391 | tri/oligope | N |
| 456 | Corn | 433.2673 | 1 | 16.38399 | 114721.6 | 432.26 | tri/oligope | N |
| 457 | Corn | 436.344 | 1 | 0.020133 | 257768.3 | 435.3367 | tri/oligope | N |
| 458 | Corn | 439.2931 | 1 | 19.90277 | 800375.1 | 438.2858 | tri/oligope | N |
| 459 | Corn | 441.3089 | 1 | 24.74833 | 328827.2 | 440.3016 | tri/oligope | N |
| 460 | Corn | 442.2676 | 1 | 13.3633 | 212568.6 | 441.2603 | tri/oligope | N |
| 461 | Corn | 442.2679 | 1 | 15.72072 | 103760.3 | 441.2606 | tri/oligope | N |
| 462 | Corn | 443.2259 | 1 | 2.193917 | 234191.6 | 442.2186 | tri/oligope | N |
| 463 | Corn | 444.2469 | 1 | 11.69911 | 385719.8 | 443.2396 | tri/oligope | N |
| 464 | Corn | 445.2415 | 1 | 2.182567 | 151465.3 | 444.2343 | tri/oligope | N |
| 465 | Corn | 445.2419 | 1 | 2.885633 | 633838.8 | 444.2346 | tri/oligope | N |
| 466 | Corn | 447.262 | 1 | 20.74559 | 444748.9 | 446.2547 | tri/oligope | N |
| 467 | Corn | 447.262 | 1 | 25.55214 | 230948.3 | 446.2547 | tri/oligope | N |
| 468 | Corn | 451.2204 | 1 | 14.03378 | 426214.2 | 450.2132 | tri/oligope | N |
| 469 | Corn | 453.2728 | 1 | 27.5805 | 231617.1 | 452.2656 | tri/oligope | N |
| 470 | Corn | 455.3245 | 1 | 26.47475 | 100017.7 | 454.3172 | tri/oligope | N |
| 471 | Corn | 455.3246 | 1 | 26.80086 | 455916.6 | 454.3173 | tri/oligope | N |
| 472 | Corn | 455.3248 | 1 | 27.76863 | 175582 | 454.3176 | tri/oligope | N |
| 473 | Corn | 456.2832 | 1 | 22.19738 | 283522.5 | 455.2759 | tri/oligope | N |
| 474 | Corn | 456.2833 | 1 | 20.49612 | 474204.5 | 455.276 | tri/oligope | N |
| 475 | Corn | 459.2571 | 1 | 2.182567 | 615002.9 | 458.2498 | tri/oligope | N |
| 476 | Corn | 462.2571 | 1 | 2.38445 | 317838.9 | 461.2498 | tri/oligope | N |
| 477 | Corn | 463.2564 | 1 | 22.69029 | 1019681 | 462.2491 | tri/oligope | N |
| 478 | Corn | 463.2568 | 1 | 19.06212 | 206891.1 | 462.2495 | tri/oligope | N |
| 479 | Corn | 463.2572 | 1 | 25.77708 | 108852.7 | 462.2499 | tri/oligope | N |
| 480 | Corn | 470.298 | 1 | 23.12191 | 7142067 | 469.2907 | tri/oligope | N |
| 481 | Corn | 470.2987 | 1 | 15.02083 | 1287939 | 469.2914 | tri/oligope | N |
| 482 | Corn | 235.6531 | 2 | 15.01209 | 269174.8 | 469.2916 | tri/oligope | N |
| 483 | Corn | 471.2573 | 1 | 2.182567 | 276945.8 | 470.25 | tri/oligope | N |
| 484 | Corn | 473.2367 | 1 | 2.182567 | 111728.3 | 472.2294 | tri/oligope | N |
| 485 | Corn | 473.2774 | 1 | 22.05129 | 140183.9 | 472.2701 | tri/oligope | N |
| 486 | Corn | 475.252 | 1 | 2.182567 | 258107.8 | 474.2447 | tri/oligope | N |
| 487 | Corn | 475.2523 | 1 | 2.522992 | 134919.3 | 474.2451 | tri/oligope | N |
| 488 | Corn | 242.6666 | 2 | 12.99005 | 138787.9 | 483.3187 | tri/oligope | N |
| 489 | Corn | 485.2734 | 1 | 12.70335 | 1144959 | 484.2662 | tri/oligope | N |
| 490 | Corn | 485.2736 | 1 | 11.25363 | 159230 | 484.2663 | tri/oligope | N |

| | | | | | | | | |
|---|---|---|---|---|---|---|---|---|
| 491 | Corn | 485.2736 | 1 | 16.17312 | 577227.9 | 484.2664 | tri/oligope | N |
| 492 | Corn | 487.2573 | 1 | 29.3331 | 132183.8 | 486.25 | tri/oligope | N |
| 493 | Corn | 487.2892 | 1 | 15.63426 | 291668.7 | 486.2819 | tri/oligope | N |
| 494 | Corn | 489.2682 | 1 | 2.5683 | 111633.7 | 488.2609 | tri/oligope | N |
| 495 | Corn | 489.2729 | 1 | 20.61214 | 568516.7 | 488.2656 | tri/oligope | N |
| 496 | Corn | 490.2521 | 1 | 14.57228 | 819257 | 489.2448 | tri/oligope | N |
| 497 | Corn | 490.2679 | 1 | 21.78767 | 150210.8 | 489.2606 | tri/oligope | N |
| 498 | Corn | 499.2889 | 1 | 14.64181 | 159545.6 | 498.2816 | tri/oligope | N |
| 499 | Corn | 499.2892 | 1 | 13.47342 | 467868.3 | 498.2819 | tri/oligope | N |
| 500 | Corn | 500.3092 | 1 | 19.92463 | 187494.7 | 499.3019 | tri/oligope | N |
| 501 | Corn | 500.3099 | 1 | 23.74262 | 113619.5 | 499.3026 | tri/oligope | N |
| 502 | Corn | 501.268 | 1 | 2.534883 | 1140694 | 500.2607 | tri/oligope | N |
| 503 | Corn | 501.2681 | 1 | 15.45965 | 929826.6 | 500.2609 | tri/oligope | N |
| 504 | Corn | 501.2682 | 1 | 14.72839 | 776547.8 | 500.2609 | tri/oligope | N |
| 505 | Corn | 251.138 | 2 | 15.46473 | 109901.3 | 500.2615 | tri/oligope | N |
| 506 | Corn | 504.2828 | 1 | 24.82333 | 3444882 | 503.2755 | tri/oligope | N |
| 507 | Corn | 504.2833 | 1 | 22.97654 | 274978.7 | 503.276 | tri/oligope | N |
| 508 | Corn | 504.2835 | 1 | 19.33528 | 318501.1 | 503.2762 | tri/oligope | N |
| 509 | Corn | 506.2628 | 1 | 15.90739 | 110675.8 | 505.2555 | tri/oligope | N |
| 510 | Corn | 254.6264 | 2 | 23.11553 | 147769.4 | 507.2383 | tri/oligope | N |
| 511 | Corn | 512.3451 | 1 | 28.89864 | 2895232 | 511.3378 | tri/oligope | N |
| 512 | Corn | 513.3051 | 1 | 18.8046 | 295723.9 | 512.2978 | tri/oligope | N |
| 513 | Corn | 515.2837 | 1 | 2.407683 | 118520.7 | 514.2765 | tri/oligope | N |
| 514 | Corn | 515.2838 | 1 | 14.71723 | 343090 | 514.2765 | tri/oligope | N |
| 515 | Corn | 515.284 | 1 | 13.26439 | 116655 | 514.2767 | tri/oligope | N |
| 516 | Corn | 515.2841 | 1 | 12.03925 | 350117.8 | 514.2768 | tri/oligope | N |
| 517 | Corn | 520.2785 | 1 | 18.49042 | 439136.2 | 519.2712 | tri/oligope | N |
| 518 | Corn | 522.3304 | 1 | 19.98531 | 105382.5 | 521.3231 | tri/oligope | N |
| 519 | Corn | 526.3253 | 1 | 23.09667 | 1570616 | 525.318 | tri/oligope | N |
| 520 | Corn | 526.3612 | 1 | 28.90485 | 269459 | 525.3539 | tri/oligope | N |
| 521 | Corn | 527.1853 | 1 | 2.444425 | 128831.3 | 526.178 | tri/oligope | N |
| 522 | Corn | 527.3208 | 1 | 16.95263 | 157565.6 | 526.3135 | tri/oligope | N |
| 523 | Corn | 530.2579 | 1 | 2.182567 | 198903.2 | 529.2506 | tri/oligope | N |
| 524 | Corn | 531.2613 | 1 | 19.38906 | 137485.3 | 530.254 | tri/oligope | N |
| 525 | Corn | 531.2615 | 1 | 14.23405 | 270422.2 | 530.2543 | tri/oligope | N |
| 526 | Corn | 541.3359 | 1 | 19.11893 | 890820.9 | 540.3286 | tri/oligope | N |
| 527 | Corn | 271.1717 | 2 | 19.12538 | 553041.6 | 540.3289 | tri/oligope | N |
| 528 | Corn | 542.2945 | 1 | 2.240483 | 309734.3 | 541.2872 | tri/oligope | N |
| 529 | Corn | 543.3154 | 1 | 21.58534 | 501710 | 542.3081 | tri/oligope | N |
| 530 | Corn | 543.3154 | 1 | 20.04357 | 105232 | 542.3082 | tri/oligope | N |
| 531 | Corn | 544.2736 | 1 | 2.193917 | 138879.4 | 543.2664 | tri/oligope | N |
| 532 | Corn | 549.2685 | 1 | 12.86502 | 153646.1 | 548.2612 | tri/oligope | N |
| 533 | Corn | 276.66 | 2 | 33.84269 | 102886.4 | 551.3055 | tri/oligope | N |
| 534 | Corn | 556.3107 | 1 | 14.84938 | 736337.9 | 555.3034 | tri/oligope | N |
| 535 | Corn | 556.3113 | 1 | 12.35729 | 137749.1 | 555.304 | tri/oligope | N |
| 536 | Corn | 557.3311 | 1 | 19.83533 | 167236.3 | 556.3239 | tri/oligope | N |
| 537 | Corn | 559.274 | 1 | 2.407683 | 209712 | 558.2667 | tri/oligope | N |
| 538 | Corn | 560.3105 | 1 | 24.54669 | 130390.2 | 559.3032 | tri/oligope | N |
| 539 | Corn | 560.3106 | 1 | 20.71625 | 198781.8 | 559.3033 | tri/oligope | N |
| 540 | Corn | 560.3216 | 1 | 14.42962 | 284901.4 | 559.3143 | tri/oligope | N |

| | | | | | | | | |
|---|---|---|---|---|---|---|---|---|
| 541 | Corn | 570.3272 | 1 | 15.83318 | 178211.6 | 569.3199 | tri/oligope | N |
| 542 | Corn | 572.3053 | 1 | 2.182567 | 308430.4 | 571.2981 | tri/oligope | N |
| 543 | Corn | 572.3465 | 1 | 26.21093 | 1583364 | 571.3393 | tri/oligope | N |
| 544 | Corn | 575.3209 | 1 | 23.35982 | 549359.4 | 574.3136 | tri/oligope | N |
| 545 | Corn | 581.3317 | 1 | 26.3655 | 282903.8 | 580.3244 | tri/oligope | N |
| 546 | Corn | 583.3827 | 1 | 28.98068 | 2769437 | 582.3754 | tri/oligope | N |
| 547 | Corn | 583.383 | 1 | 23.12191 | 101107.4 | 582.3757 | tri/oligope | N |
| 548 | Corn | 292.1953 | 2 | 28.96868 | 140773.7 | 582.3761 | tri/oligope | N |
| 549 | Corn | 584.3421 | 1 | 19.70915 | 858209.8 | 583.3348 | tri/oligope | N |
| 550 | Corn | 584.3428 | 1 | 17.24573 | 107379.4 | 583.3355 | tri/oligope | N |
| 551 | Corn | 585.3263 | 1 | 19.14884 | 748268 | 584.319 | tri/oligope | N |
| 552 | Corn | 586.3209 | 1 | 2.193917 | 324173.3 | 585.3136 | tri/oligope | N |
| 553 | Corn | 586.3214 | 1 | 15.02083 | 1428076 | 585.3141 | tri/oligope | N |
| 554 | Corn | 293.6645 | 2 | 15.01209 | 127967 | 585.3144 | tri/oligope | N |
| 555 | Corn | 587.306 | 1 | 16.1594 | 350305.2 | 586.2988 | tri/oligope | N |
| 556 | Corn | 590.2959 | 1 | 16.04208 | 122779.8 | 589.2886 | tri/oligope | N |
| 557 | Corn | 591.3155 | 1 | 22.07953 | 125966.2 | 590.3082 | tri/oligope | N |
| 558 | Corn | 598.3572 | 1 | 23.20908 | 4265186 | 597.35 | tri/oligope | N |
| 559 | Corn | 299.6824 | 2 | 23.20203 | 1058825 | 597.3503 | tri/oligope | N |
| 560 | Corn | 598.3583 | 1 | 22.91849 | 921363.8 | 597.351 | tri/oligope | N |
| 561 | Corn | 299.6828 | 2 | 15.6513 | 223429.7 | 597.3511 | tri/oligope | N |
| 562 | Corn | 598.3584 | 1 | 15.6447 | 168191.2 | 597.3512 | tri/oligope | N |
| 563 | Corn | 299.6829 | 2 | 20.23801 | 546063.5 | 597.3512 | tri/oligope | N |
| 564 | Corn | 598.3586 | 1 | 20.23801 | 444305.5 | 597.3513 | tri/oligope | N |
| 565 | Corn | 598.3586 | 1 | 18.61213 | 508694.8 | 597.3514 | tri/oligope | N |
| 566 | Corn | 299.683 | 2 | 18.60463 | 189103.7 | 597.3515 | tri/oligope | N |
| 567 | Corn | 300.6437 | 2 | 13.00488 | 786453.5 | 599.2728 | tri/oligope | N |
| 568 | Corn | 602.2992 | 1 | 15.99201 | 263615.1 | 601.2919 | tri/oligope | N |
| 569 | Corn | 602.3163 | 1 | 2.522992 | 368182.3 | 601.3091 | tri/oligope | N |
| 570 | Corn | 602.3167 | 1 | 14.44234 | 717505.6 | 601.3094 | tri/oligope | N |
| 571 | Corn | 301.6621 | 2 | 14.44234 | 150353.9 | 601.3096 | tri/oligope | N |
| 572 | Corn | 602.3577 | 1 | 25.08481 | 154119 | 601.3504 | tri/oligope | N |
| 573 | Corn | 603.3371 | 1 | 19.37633 | 672852.4 | 602.3298 | tri/oligope | N |
| 574 | Corn | 604.2827 | 1 | 18.28582 | 307615.7 | 603.2754 | tri/oligope | N |
| 575 | Corn | 605.3308 | 1 | 22.01991 | 3401053 | 604.3235 | tri/oligope | N |
| 576 | Corn | 605.331 | 1 | 23.19293 | 119274.9 | 604.3237 | tri/oligope | N |
| 577 | Corn | 303.1693 | 2 | 22.01991 | 3259027 | 604.324 | tri/oligope | N |
| 578 | Corn | 607.3112 | 1 | 20.40947 | 222176.9 | 606.304 | tri/oligope | N |
| 579 | Corn | 304.1828 | 2 | 19.72981 | 335145.9 | 606.351 | tri/oligope | N |
| 580 | Corn | 304.1832 | 2 | 17.19853 | 258879.9 | 606.3518 | tri/oligope | N |
| 581 | Corn | 305.6435 | 2 | 15.01209 | 173335.7 | 609.2724 | tri/oligope | N |
| 582 | Corn | 305.6829 | 2 | 21.77626 | 115990.3 | 609.3513 | tri/oligope | N |
| 583 | Corn | 612.3741 | 1 | 21.14709 | 129768.7 | 611.3668 | tri/oligope | N |
| 584 | Corn | 612.3743 | 1 | 20.75984 | 141997 | 611.367 | tri/oligope | N |
| 585 | Corn | 307.1702 | 2 | 16.62097 | 483966.4 | 612.3259 | tri/oligope | N |
| 586 | Corn | 613.3332 | 1 | 16.60424 | 228256.3 | 612.3259 | tri/oligope | N |
| 587 | Corn | 613.3939 | 1 | 28.95146 | 336203.4 | 612.3866 | tri/oligope | N |
| 588 | Corn | 614.3527 | 1 | 19.85123 | 441196.6 | 613.3454 | oligopepti | N |
| 589 | Corn | 614.353 | 1 | 21.61801 | 880579.7 | 613.3457 | oligopepti | N |
| 590 | Corn | 307.6802 | 2 | 21.62668 | 303047 | 613.3457 | oligopepti | N |

| | | | | | | | | |
|---|---|---|---|---|---|---|---|---|
| 591 | Corn | 616.3321 | 1 | 14.70728 | 428634.5 | 615.3248 | oligopepti( | N |
| 592 | Corn | 309.1877 | 2 | 25.5411 | 569429.4 | 616.3608 | oligopepti( | N |
| 593 | Corn | 617.3681 | 1 | 25.5411 | 666602.9 | 616.3609 | oligopepti( | N |
| 594 | Corn | 618.2933 | 1 | 2.288042 | 279242.1 | 617.286 | oligopepti( | N |
| 595 | Corn | 620.2775 | 1 | 15.27055 | 177726.8 | 619.2703 | oligopepti( | N |
| 596 | Corn | 620.3066 | 1 | 13.5986 | 127026 | 619.2993 | oligopepti( | N |
| 597 | Corn | 310.657 | 2 | 13.59221 | 103268.6 | 619.2994 | oligopepti( | N |
| 598 | Corn | 310.6672 | 2 | 23.20908 | 126410.5 | 619.3199 | oligopepti( | N |
| 599 | Corn | 311.1689 | 2 | 28.96868 | 110217.7 | 620.3232 | oligopepti( | N |
| 600 | Corn | 311.6614 | 2 | 23.20908 | 476032.3 | 621.3083 | oligopepti( | N |
| 601 | Corn | 312.638 | 2 | 15.01209 | 104258 | 623.2614 | oligopepti( | N |
| 602 | Corn | 313.641 | 2 | 14.44234 | 155813.6 | 625.2675 | oligopepti( | N |
| 603 | Corn | 314.1538 | 2 | 22.01058 | 101571.1 | 626.2931 | oligopepti( | N |
| 604 | Corn | 627.3487 | 1 | 16.17959 | 196004 | 626.3414 | oligopepti( | N |
| 605 | Corn | 315.1481 | 2 | 22.01058 | 436918.1 | 628.2816 | oligopepti( | N |
| 606 | Corn | 629.3281 | 1 | 2.193917 | 123269.7 | 628.3208 | oligopepti( | N |
| 607 | Corn | 631.3477 | 1 | 21.56201 | 139130.5 | 630.3404 | oligopepti( | N |
| 608 | Corn | 632.3104 | 1 | 19.52863 | 139073.9 | 631.3031 | oligopepti( | N |
| 609 | Corn | 632.3432 | 1 | 24.77268 | 367105 | 631.3359 | oligopepti( | N |
| 610 | Corn | 634.3226 | 1 | 21.41064 | 102500.6 | 633.3153 | oligopepti( | N |
| 611 | Corn | 318.6561 | 2 | 23.20908 | 274492.3 | 635.2976 | oligopepti( | N |
| 612 | Corn | 319.1494 | 2 | 16.62097 | 110160.2 | 636.2843 | oligopepti( | N |
| 613 | Corn | 319.1575 | 2 | 2.030925 | 112645.5 | 636.3004 | oligopepti( | N |
| 614 | Corn | 319.6594 | 2 | 21.62668 | 144736.8 | 637.3042 | oligopepti( | N |
| 615 | Corn | 321.1753 | 2 | 22.19738 | 101926.8 | 640.336 | oligopepti( | N |
| 616 | Corn | 642.3851 | 1 | 24.0136 | 135788.2 | 641.3778 | oligopepti( | N |
| 617 | Corn | 322.1427 | 2 | 22.01991 | 272867.9 | 642.2709 | oligopepti( | N |
| 618 | Corn | 643.343 | 1 | 14.73782 | 211957.8 | 642.3357 | oligopepti( | N |
| 619 | Corn | 322.1753 | 2 | 14.73782 | 207328.1 | 642.3361 | oligopepti( | N |
| 620 | Corn | 322.1755 | 2 | 12.48661 | 100324.9 | 642.3365 | oligopepti( | N |
| 621 | Corn | 323.1911 | 2 | 15.05633 | 110406.7 | 644.3676 | oligopepti( | N |
| 622 | Corn | 646.3216 | 1 | 14.88147 | 345036.8 | 645.3143 | oligopepti( | N |
| 623 | Corn | 323.6645 | 2 | 14.88952 | 302013.1 | 645.3145 | oligopepti( | N |
| 624 | Corn | 648.304 | 1 | 14.88952 | 141268.8 | 647.2967 | oligopepti( | N |
| 625 | Corn | 648.3379 | 1 | 18.21343 | 208803 | 647.3307 | oligopepti( | N |
| 626 | Corn | 324.6727 | 2 | 18.21343 | 201084.5 | 647.3308 | oligopepti( | N |
| 627 | Corn | 326.6422 | 2 | 23.19293 | 106396.8 | 651.2698 | oligopepti( | N |
| 628 | Corn | 655.38 | 1 | 18.05075 | 492222.4 | 654.3727 | oligopepti( | N |
| 629 | Corn | 328.1936 | 2 | 18.05075 | 252031.3 | 654.3727 | oligopepti( | N |
| 630 | Corn | 329.1543 | 2 | 14.87133 | 378404.6 | 656.2941 | oligopepti( | N |
| 631 | Corn | 330.199 | 2 | 18.34035 | 794034.6 | 658.3834 | oligopepti( | N |
| 632 | Corn | 334.1612 | 2 | 20.42355 | 101293.1 | 666.3078 | oligopepti( | N |
| 633 | Corn | 673.3543 | 1 | 15.93387 | 164648.5 | 672.347 | oligopepti( | N |
| 634 | Corn | 673.3953 | 1 | 24.16437 | 151029.2 | 672.388 | oligopepti( | N |
| 635 | Corn | 677.3279 | 1 | 13.3789 | 192834.6 | 676.3206 | oligopepti( | N |
| 636 | Corn | 339.1678 | 2 | 13.3633 | 467941.8 | 676.3211 | oligopepti( | N |
| 637 | Corn | 688.3697 | 1 | 23.98862 | 224659.9 | 687.3625 | oligopepti( | N |
| 638 | Corn | 698.2517 | 1 | 17.48576 | 259592.2 | 697.2444 | oligopepti( | N |
| 639 | Corn | 351.1467 | 2 | 13.3789 | 175686.5 | 700.2789 | oligopepti( | N |
| 640 | Corn | 703.3798 | 1 | 23.32184 | 238260.1 | 702.3725 | oligopepti( | N |

| | | | | | | | | |
|---|---|---|---|---|---|---|---|---|
| 641 | Corn | 352.1937 | 2 | 23.32184 | 194881 | 702.3729 | oligopepti( | N |
| 642 | Corn | 354.1991 | 2 | 25.81663 | 679004.5 | 706.3836 | oligopepti( | N |
| 643 | Corn | 356.2248 | 2 | 26.13642 | 1202231 | 710.435 | oligopepti( | N |
| 644 | Corn | 711.4425 | 1 | 26.13642 | 370493.8 | 710.4352 | oligopepti( | N |
| 645 | Corn | 718.4169 | 1 | 27.19068 | 101267.1 | 717.4097 | oligopepti( | N |
| 646 | Corn | 359.7122 | 2 | 27.19068 | 154301.4 | 717.4098 | oligopepti( | N |
| 647 | Corn | 360.1914 | 2 | 18.08528 | 105958.8 | 718.3682 | oligopepti( | N |
| 648 | Corn | 360.725 | 2 | 23.20908 | 384169.3 | 719.4354 | oligopepti( | N |
| 649 | Corn | 361.6085 | 2 | 17.48576 | 147232.4 | 721.2025 | oligopepti( | N |
| 650 | Corn | 363.1424 | 2 | 22.01991 | 104669 | 724.2703 | oligopepti( | N |
| 651 | Corn | 363.7123 | 2 | 19.58842 | 370876.5 | 725.41 | oligopepti( | N |
| 652 | Corn | 363.7123 | 2 | 22.97654 | 408083.8 | 725.41 | oligopepti( | N |
| 653 | Corn | 368.2039 | 2 | 26.14323 | 192481.4 | 734.3933 | oligopepti( | N |
| 654 | Corn | 369.1532 | 2 | 14.19923 | 138647.4 | 736.2919 | oligopepti( | N |
| 655 | Corn | 371.2 | 2 | 16.68396 | 300055.6 | 740.3854 | oligopepti( | N |
| 656 | Corn | 760.4266 | 1 | 23.11553 | 312462.7 | 759.4193 | oligopepti( | N |
| 657 | Corn | 380.717 | 2 | 23.09667 | 166966.2 | 759.4195 | oligopepti( | N |
| 658 | Corn | 767.3855 | 1 | 20.97956 | 104991.1 | 766.3782 | oligopepti( | N |
| 659 | Corn | 384.1966 | 2 | 21.01863 | 251893.9 | 766.3787 | oligopepti( | N |
| 660 | Corn | 384.7361 | 2 | 21.4909 | 245696.7 | 767.4576 | oligopepti( | N |
| 661 | Corn | 385.2155 | 2 | 17.21054 | 260170.3 | 768.4164 | oligopepti( | N |
| 662 | Corn | 769.4237 | 1 | 17.21939 | 175653.7 | 768.4164 | oligopepti( | N |
| 663 | Corn | 390.2146 | 2 | 24.82333 | 128089.2 | 778.4146 | oligopepti( | N |
| 664 | Corn | 392.223 | 2 | 19.36508 | 385366.7 | 782.4315 | oligopepti( | N |
| 665 | Corn | 783.4389 | 1 | 19.36508 | 241395.9 | 782.4316 | oligopepti( | N |
| 666 | Corn | 392.7334 | 2 | 22.69993 | 306940.6 | 783.4523 | oligopepti( | N |
| 667 | Corn | 786.4431 | 1 | 25.90492 | 358052.3 | 785.4358 | oligopepti( | N |
| 668 | Corn | 393.7252 | 2 | 25.91128 | 371461 | 785.4359 | oligopepti( | N |
| 669 | Corn | 400.689 | 2 | 2.205492 | 132110.8 | 799.3634 | oligopepti( | N |
| 670 | Corn | 402.6726 | 2 | 16.19513 | 103214.9 | 803.3307 | oligopepti( | N |
| 671 | Corn | 404.2021 | 2 | 19.36508 | 110355 | 806.3896 | oligopepti( | N |
| 672 | Corn | 409.7258 | 2 | 21.68743 | 139314.3 | 817.437 | oligopepti( | N |
| 673 | Corn | 412.2261 | 2 | 15.5482 | 693335.3 | 822.4376 | oligopepti( | N |
| 674 | Corn | 413.2469 | 2 | 26.60053 | 110524.4 | 824.4793 | oligopepti( | N |
| 675 | Corn | 841.3103 | 1 | 20.50386 | 142090 | 840.303 | oligopepti( | N |
| 676 | Corn | 857.4802 | 1 | 26.46451 | 245611.4 | 856.4729 | oligopepti( | N |
| 677 | Corn | 429.2438 | 2 | 26.43755 | 527062.4 | 856.473 | oligopepti( | N |
| 678 | Corn | 870.7886 | 1 | 1.634117 | 121397.1 | 869.7813 | oligopepti( | N |
| 679 | Corn | 450.2673 | 2 | 28.37774 | 943931.1 | 898.52 | oligopepti( | N |
| 680 | Corn | 899.5277 | 1 | 28.36293 | 429008.9 | 898.5204 | oligopepti( | N |
| 681 | Corn | 452.2619 | 2 | 27.20595 | 135405.3 | 902.5092 | oligopepti( | N |
| 682 | Corn | 454.2352 | 2 | 22.87473 | 201039.7 | 906.4558 | oligopepti( | N |
| 683 | Corn | 460.2589 | 2 | 23.18525 | 225862.5 | 918.5033 | oligopepti( | N |
| 684 | Corn | 460.6673 | 2 | 14.26923 | 130321.1 | 919.3201 | oligopepti( | N |
| 685 | Corn | 473.7544 | 2 | 21.95956 | 1363195 | 945.4942 | oligopepti( | N |
| 686 | Corn | 946.502 | 1 | 21.95956 | 113751.4 | 945.4947 | oligopepti( | N |
| 687 | Corn | 478.2555 | 2 | 23.74262 | 132849.3 | 954.4964 | oligopepti( | N |
| 688 | Corn | 479.7371 | 2 | 17.14424 | 226288.4 | 957.4596 | oligopepti( | N |
| 689 | Corn | 485.7333 | 2 | 21.95183 | 141419.4 | 969.452 | oligopepti( | N |
| 690 | Corn | 324.1582 | 3 | 21.95956 | 255105.1 | 969.4529 | oligopepti( | N |

| | | | | | | | | |
|---|---|---|---|---|---|---|---|---|
| 691 | Corn | 485.786 | 2 | 28.70022 | 648638.8 | 969.5574 | oligopepti( | N |
| 692 | Corn | 970.565 | 1 | 28.70022 | 154372.8 | 969.5577 | oligopepti( | N |
| 693 | Corn | 485.7864 | 2 | 29.42978 | 120972.9 | 969.5583 | oligopepti( | N |
| 694 | Corn | 328.8213 | 3 | 21.95956 | 163587.8 | 983.4421 | oligopepti( | N |
| 695 | Corn | 493.7838 | 2 | 29.72158 | 402074.4 | 985.5531 | oligopepti( | N |
| 696 | Corn | 986.5606 | 1 | 29.71512 | 139164.9 | 985.5533 | oligopepti( | N |
| 697 | Corn | 502.7786 | 2 | 29.42218 | 107942.8 | 1003.543 | oligopepti( | N |
| 698 | Corn | 529.3024 | 2 | 30.03608 | 330984.5 | 1056.59 | oligopepti( | N |
| 699 | Corn | 1057.598 | 1 | 30.02927 | 103196.7 | 1056.59 | oligopepti( | N |
| 700 | Corn | 529.3025 | 2 | 29.73788 | 107592.6 | 1056.591 | oligopepti( | N |
| 701 | Corn | 548.8702 | 2 | 1.634117 | 137416.4 | 1095.726 | oligopepti( | N |
| 702 | Corn | 565.7491 | 2 | 21.85148 | 144722.6 | 1129.484 | oligopepti( | N |
| 703 | Corn | 1194.753 | 1 | 1.610975 | 118491.4 | 1193.746 | oligopepti( | N |
| 704 | Corn | 598.3114 | 2 | 19.12538 | 138468.4 | 1194.608 | oligopepti( | N |
| 705 | Corn | 609.8198 | 2 | 24.71795 | 211653.5 | 1217.625 | oligopepti( | N |
| 706 | Corn | 624.8459 | 2 | 1.634117 | 109724.7 | 1247.677 | oligopepti( | N |
| 707 | Corn | 433.926 | 3 | 24.79603 | 157337.1 | 1298.756 | oligopepti( | N |
| 708 | Corn | 654.8291 | 2 | 26.86219 | 1063134 | 1307.644 | oligopepti( | N |
| 709 | Corn | 658.8389 | 2 | 1.634117 | 107272.2 | 1315.663 | oligopepti( | N |
| 710 | Corn | 691.3515 | 2 | 26.13001 | 1004758 | 1380.688 | oligopepti( | N |
| 711 | Corn | 692.8334 | 2 | 1.634117 | 128214.7 | 1383.652 | oligopepti( | N |
| 712 | Corn | 469.2227 | 3 | 26.13001 | 213933.5 | 1404.646 | oligopepti( | N |
| 713 | Corn | 473.8856 | 3 | 26.13001 | 159725.5 | 1418.635 | oligopepti( | N |
| 714 | Corn | 726.8266 | 2 | 1.622633 | 128678.8 | 1451.639 | oligopepti( | N |
| 715 | Corn | 760.8213 | 2 | 1.622633 | 133128.7 | 1519.628 | oligopepti( | N |
| 716 | Corn | 794.8147 | 2 | 1.610975 | 116345.5 | 1587.615 | oligopepti( | N |
| 717 | Corn | 828.8083 | 2 | 1.634117 | 125133.6 | 1655.602 | oligopepti( | N |
| 718 | Corn | 854.8145 | 2 | 1.622633 | 547858.8 | 1707.615 | oligopepti( | N |
| 719 | Corn | 896.796 | 2 | 1.634117 | 105134.2 | 1791.577 | oligopepti( | N |
| 720 | Corn | 930.7894 | 2 | 1.634117 | 109844.8 | 1859.564 | oligopepti( | N |
| 721 | Corn | 964.7838 | 2 | 1.622633 | 105788.4 | 1927.553 | oligopepti( | N |
| 722 | Corn | 1024.783 | 2 | 1.622633 | 176518.1 | 2047.552 | oligopepti( | N |
| 723 | Corn | 1092.771 | 2 | 1.622633 | 144535.3 | 2183.527 | oligopepti( | N |

**Supplementary Figure 3.** Bayesian Information Criterion (BIC) analysis of the proteomic data set.

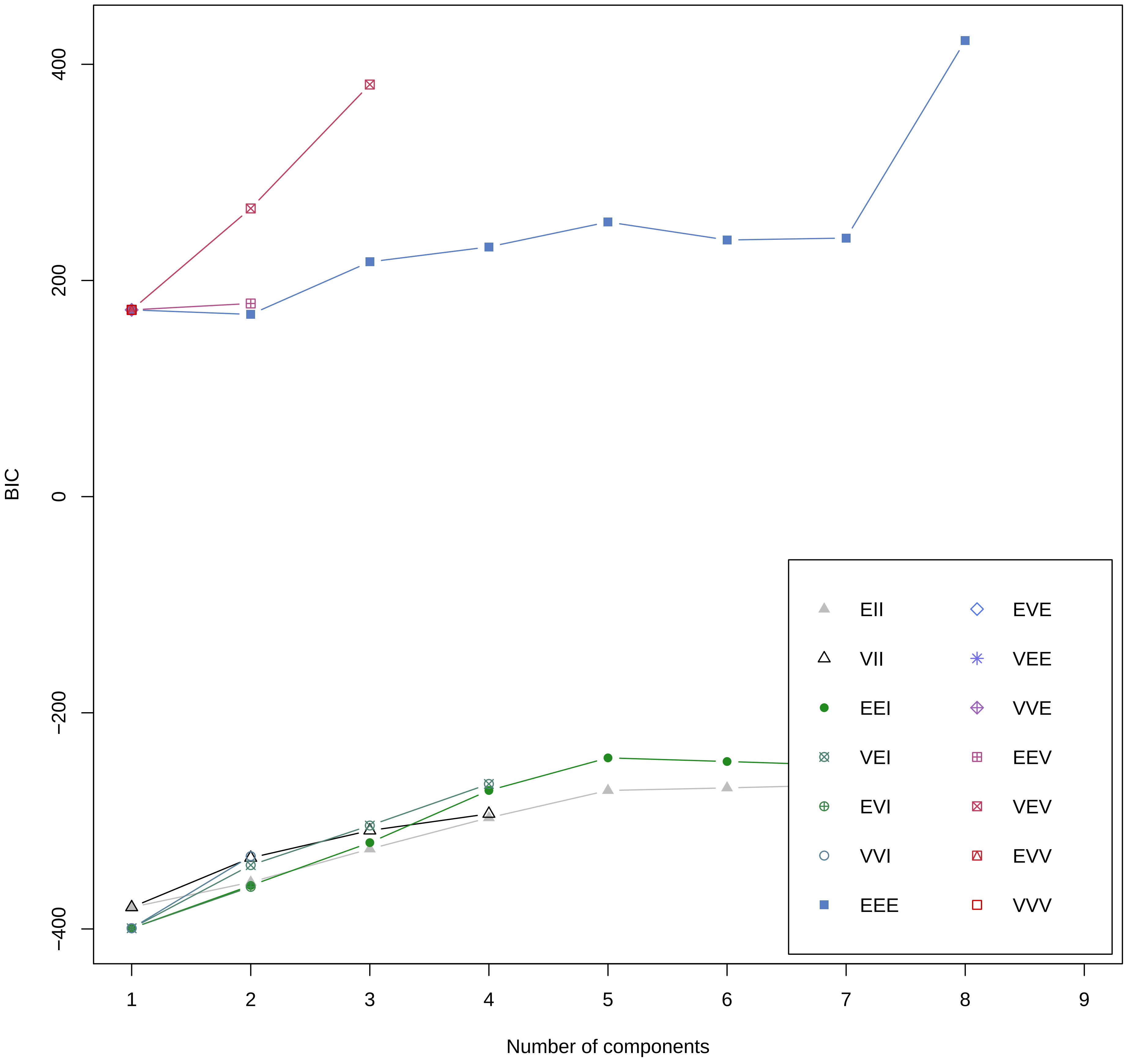

**Supplementary Figure 4.** All varieties of two component plots showing the protein clustering based on proteomic data.

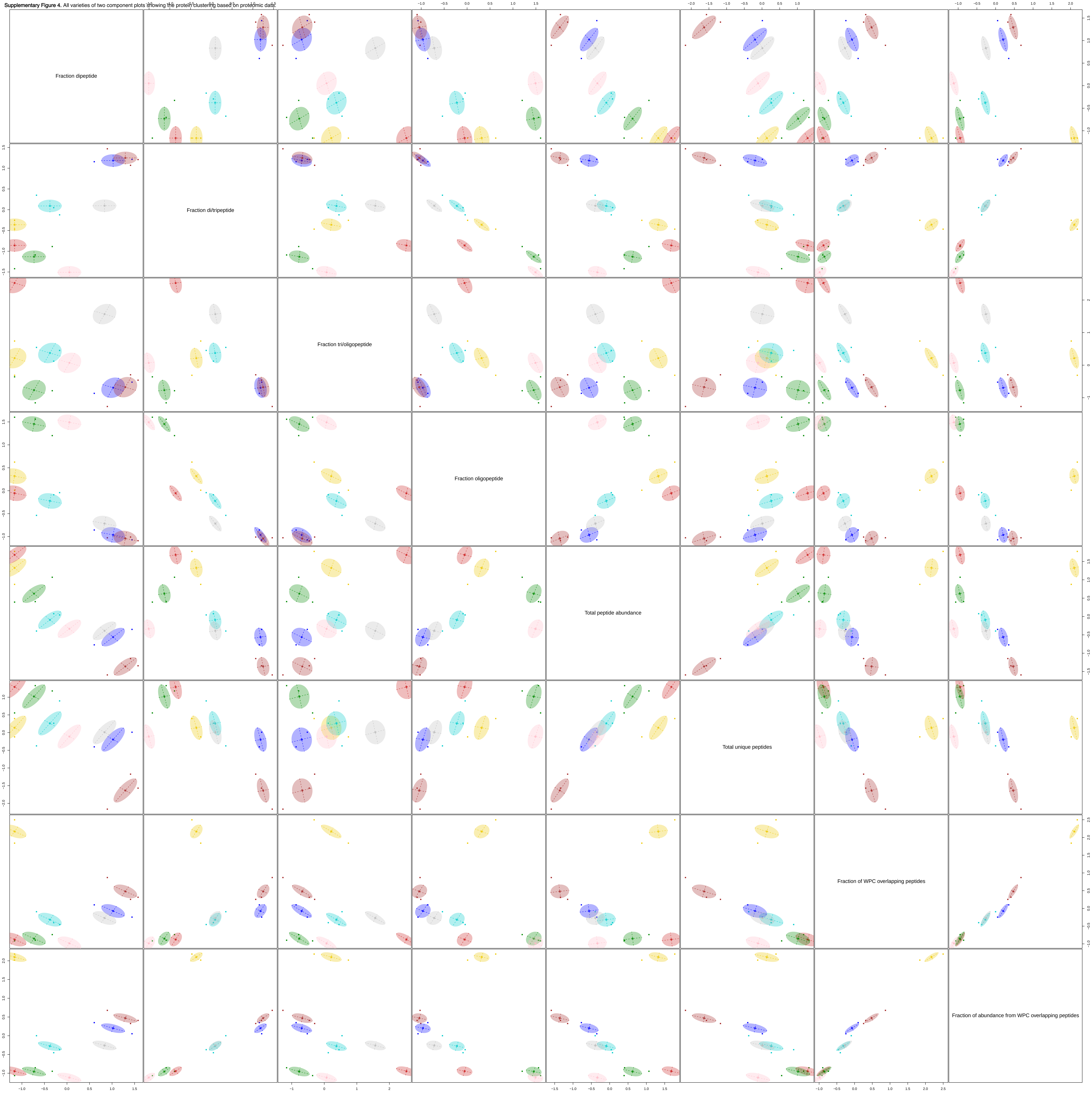

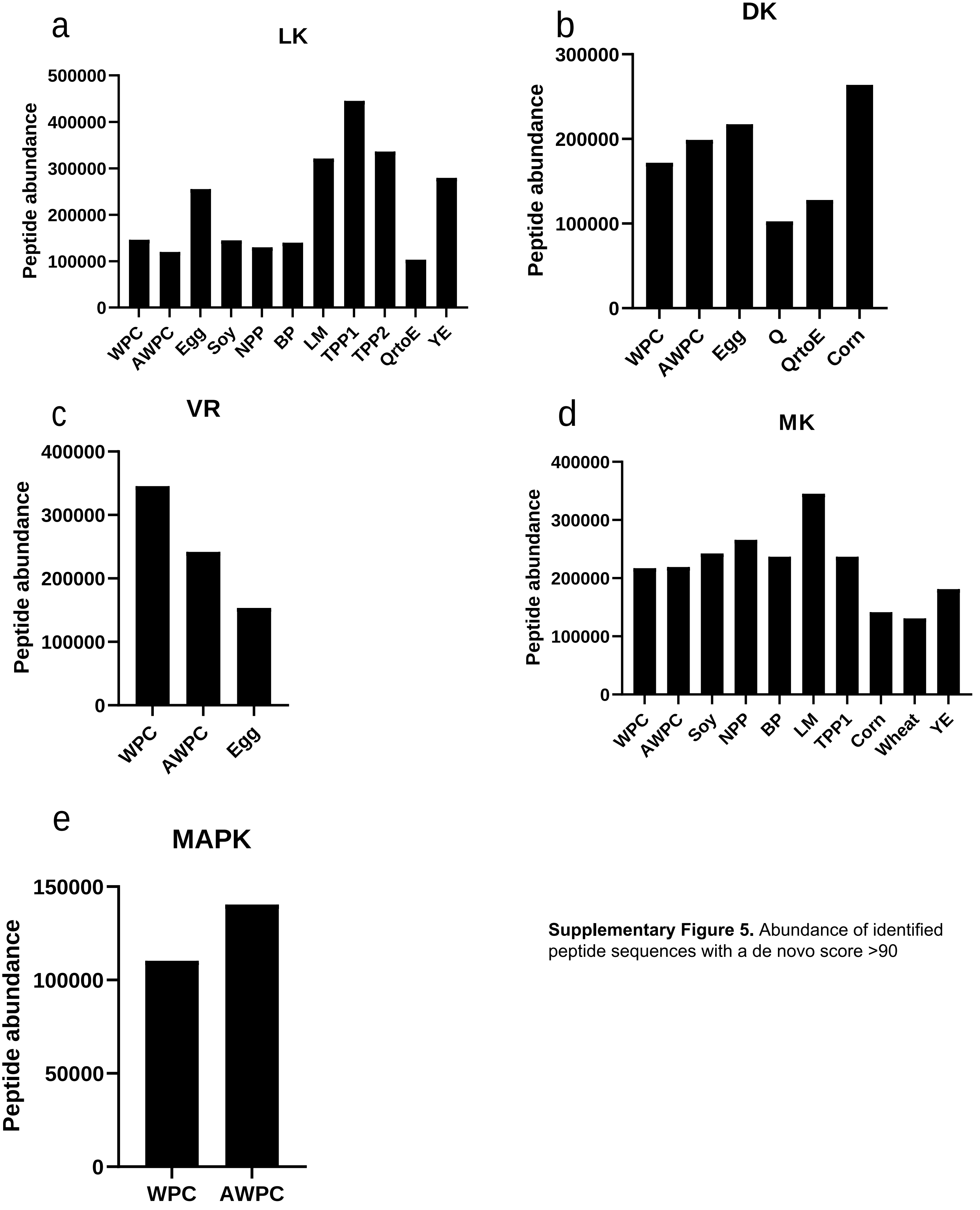

**Supplementary Figure 5.** Abundance of identified peptide sequences with a de novo score >90

**Supplementary Table 2.** Overview of the 18 dietary proteins evaluated including protein concentration based on DUMAS method and endotoxin content.

| Full name | Protein Source | Abbreviation | Protein concentration (mg/mL) | Endotoxin (EU/mL*) |
|---|---|---|---|---|
| Digestive enzyme mixtures | - | Blank | 7.7 ± 0.4 | 43374 |
| Whey protein concentrate | Whey | WPC | 12.8 ± 0.0 | 56019 |
| Acidic whey protein concentrate | Whey | AWPC | 11.6 ± 0.4 | 57911 |
| Egg | Egg white | Egg | 11.3 ± 0.1 | >65000 |
| Soy | Soya bean | Soy | 10.0 ± 0.3 | >65000 |
| NPP | Pea | NPP | 9.9 ± 0.6 | >65000 |
| NWP | Wheat | NWP | 11.4 ± 0.2 | >65000 |
| Bovine plasma | Blood protein | BP | 9.8 ± 1.8 | 13477 |
| Peptan | Blood protein | Peptan | 14.8 ± 0.2 | 63077 |
| Lesser mealworm | Insect | LM | 8.7 ± 0.5 | 55907 |
| Lesser mealworm concentrate 2 | Insect | LMC2 | 11.2 ± 0.1 | 61471 |
| TPP1 | Potato | TPP1 | 4.4 ± 0.6 | >65000 |
| TPP2 | Potato | TPP2 | 2.9 ± 0.2 | 10391 |
| Mycoprotein | Fungi | Q | 11.7 ± 3.5 | >65000 |
| Quorn ready to eat | Fungi/egg | QrtoE | 6.1 ± 0.5 | >65000 |
| Wheat | Wheat | Wheat | 4.9 ± 0.8 | 61577 |
| Corn | Corn | Corn | 9.1 ± 0.6 | >65000 |
| Yeast | Yeast | YE | 7.9 ± 0.5 | >65000 |

**a**

**b**

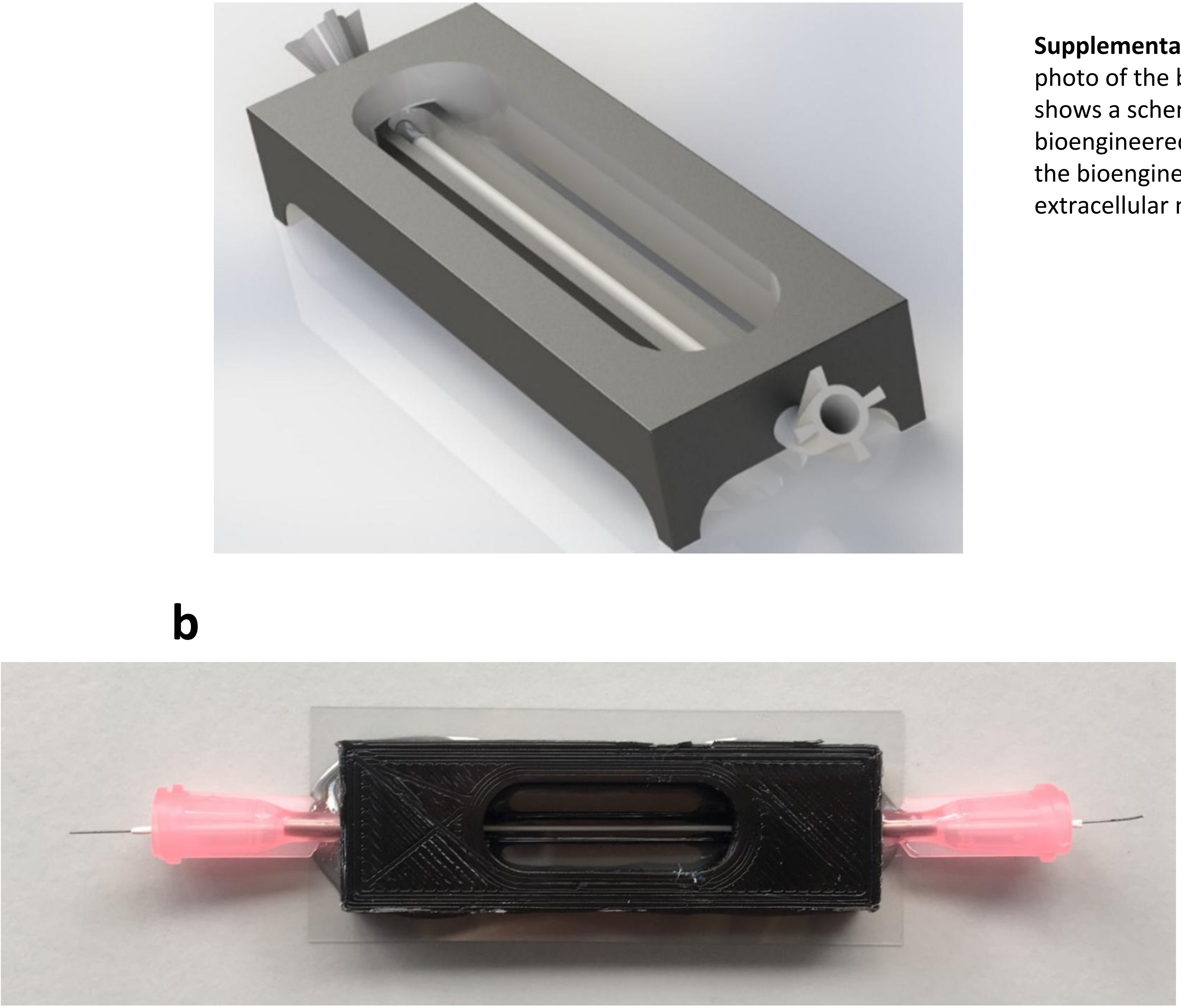

**Supplementary Figure 6.** A schematic image and photo of the bioengineered intestinal tubule. A) shows a schematic representation of the bioengineered intestinal tubules. B) shows a photo of the bioengineered intestinal tubules before extracellular matrix-coating and Caco-2 cell seeding.

**Supplementary Figure 7.** Biological effects in bioengineered intestinal tubules exposed to potential dietary proteins

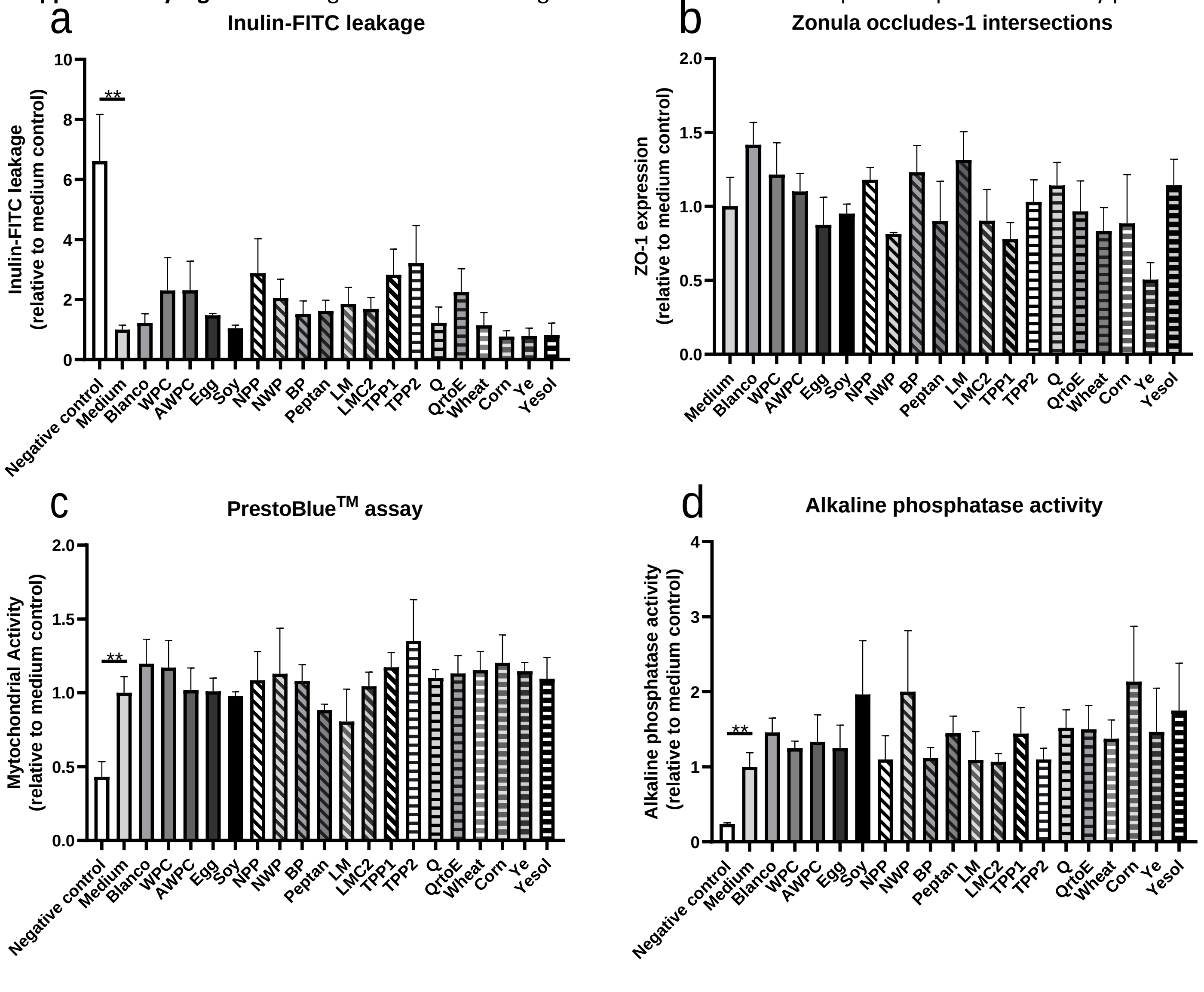

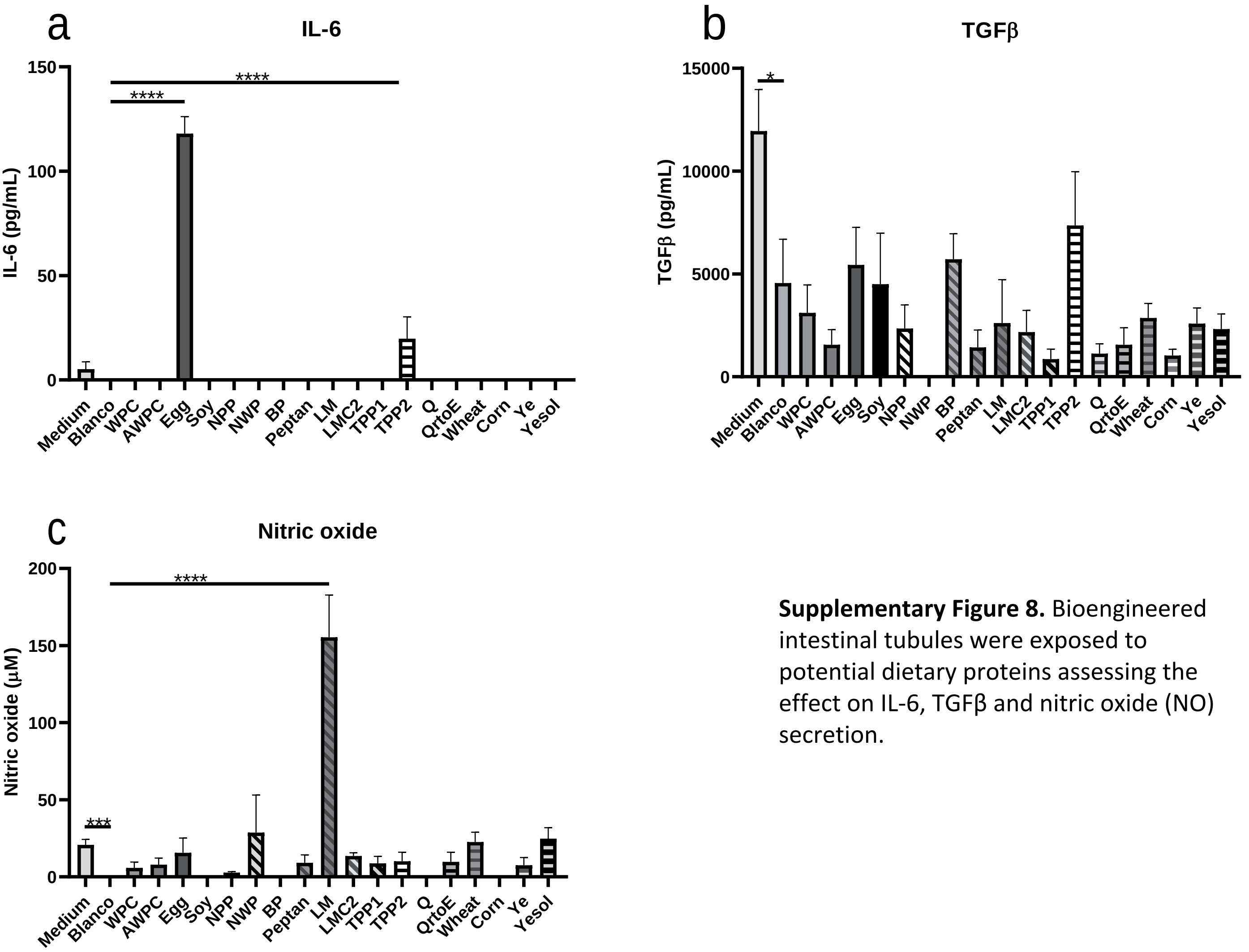

**Supplementary Figure 8.** Bioengineered intestinal tubules were exposed to potential dietary proteins assessing the effect on IL-6, TGFβ and nitric oxide (NO) secretion.

**Supplementary Figure 9.** Shows Bayesian Information Criterion (BIC) analysis of the in vitro biological efficacy data.

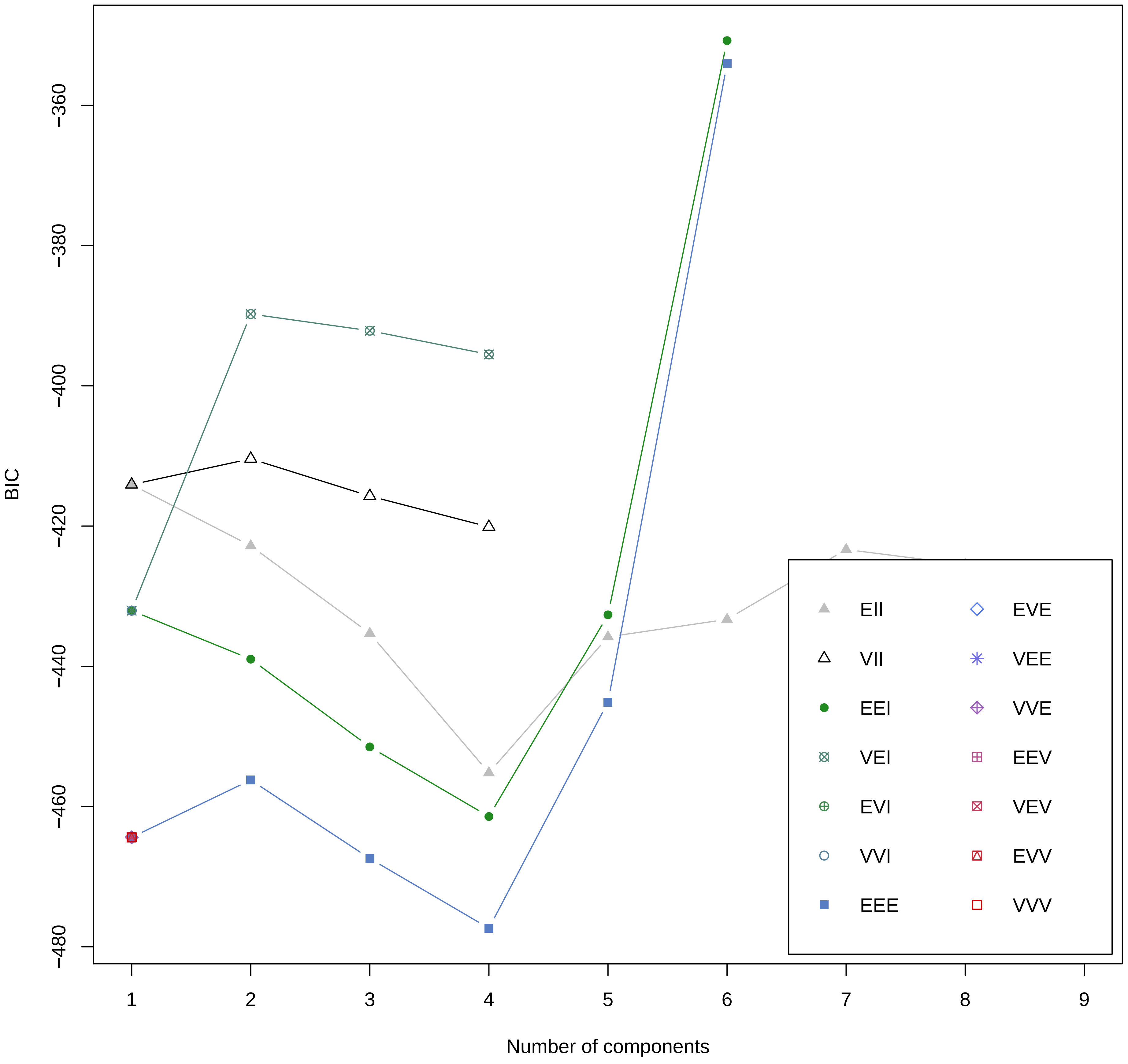

**Supplementary Figure 10.** All varieties of two component plots

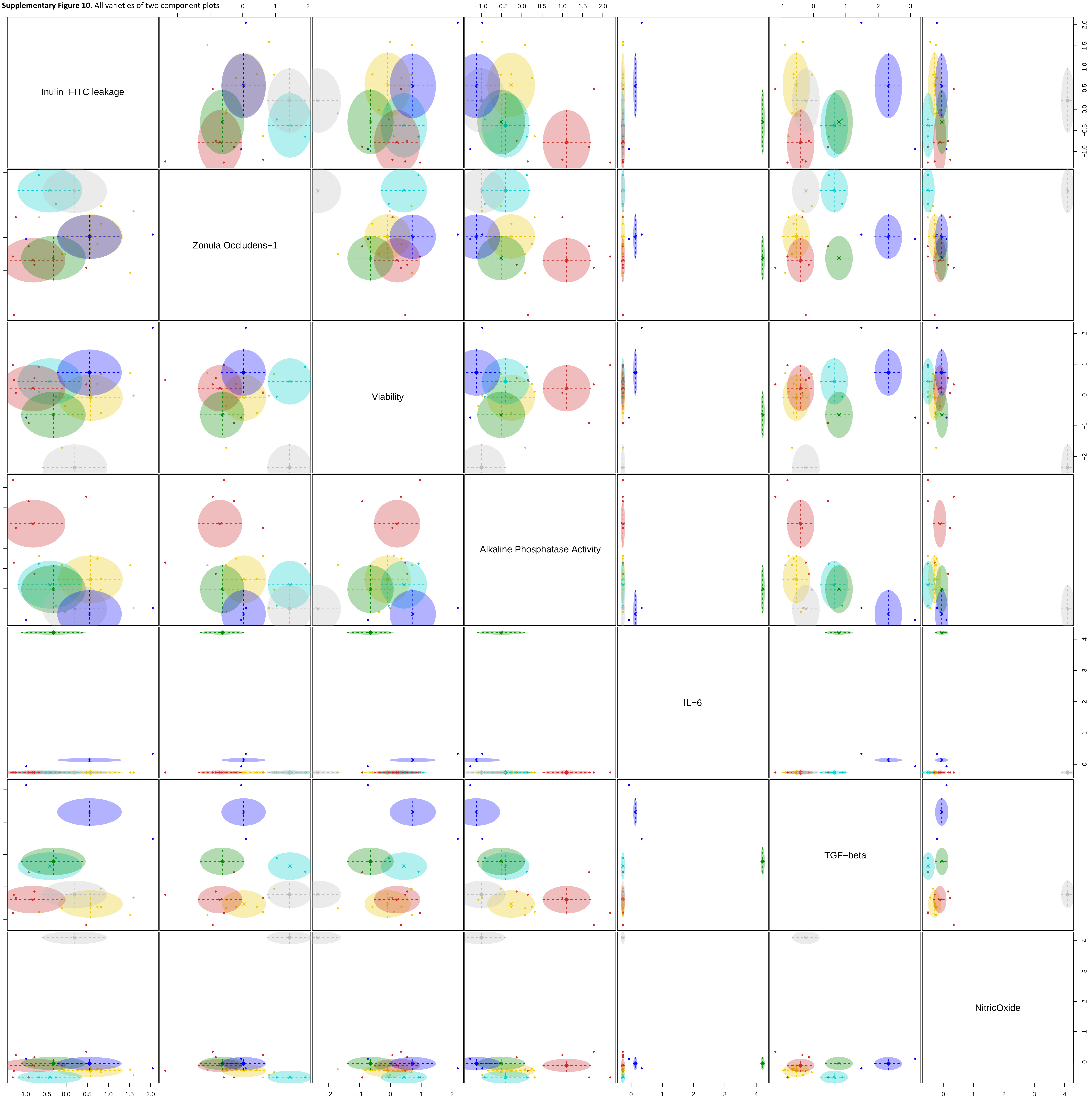

**Supplementary Figure 11.** Shows Bayesian Information Criterion (BIC) analysis of both proteomic and in vitro biological efficacy data

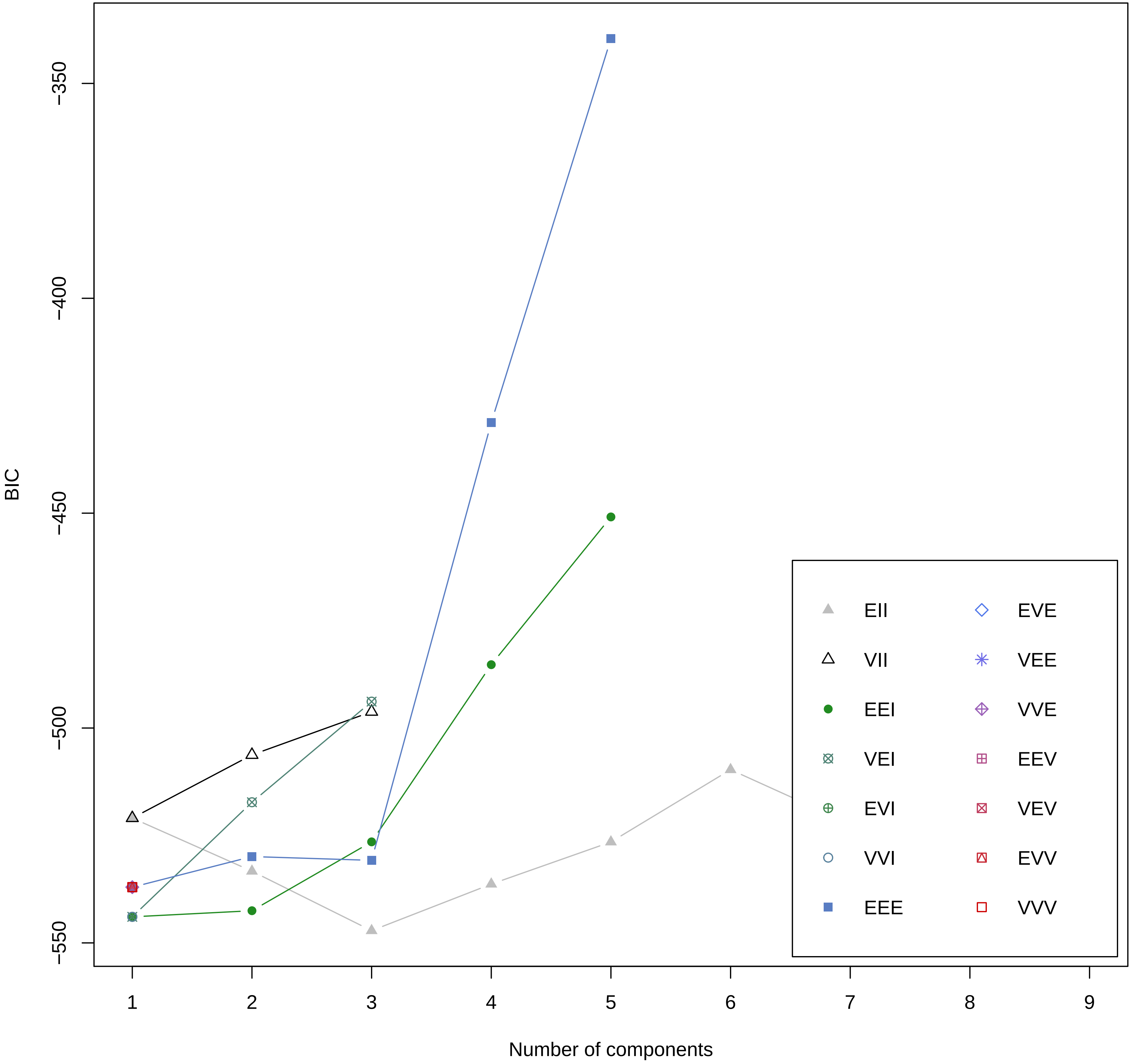

**Supplementary Figure 12.** All varieties of two component plots based on proteomic and in vitro biological efficacy data.

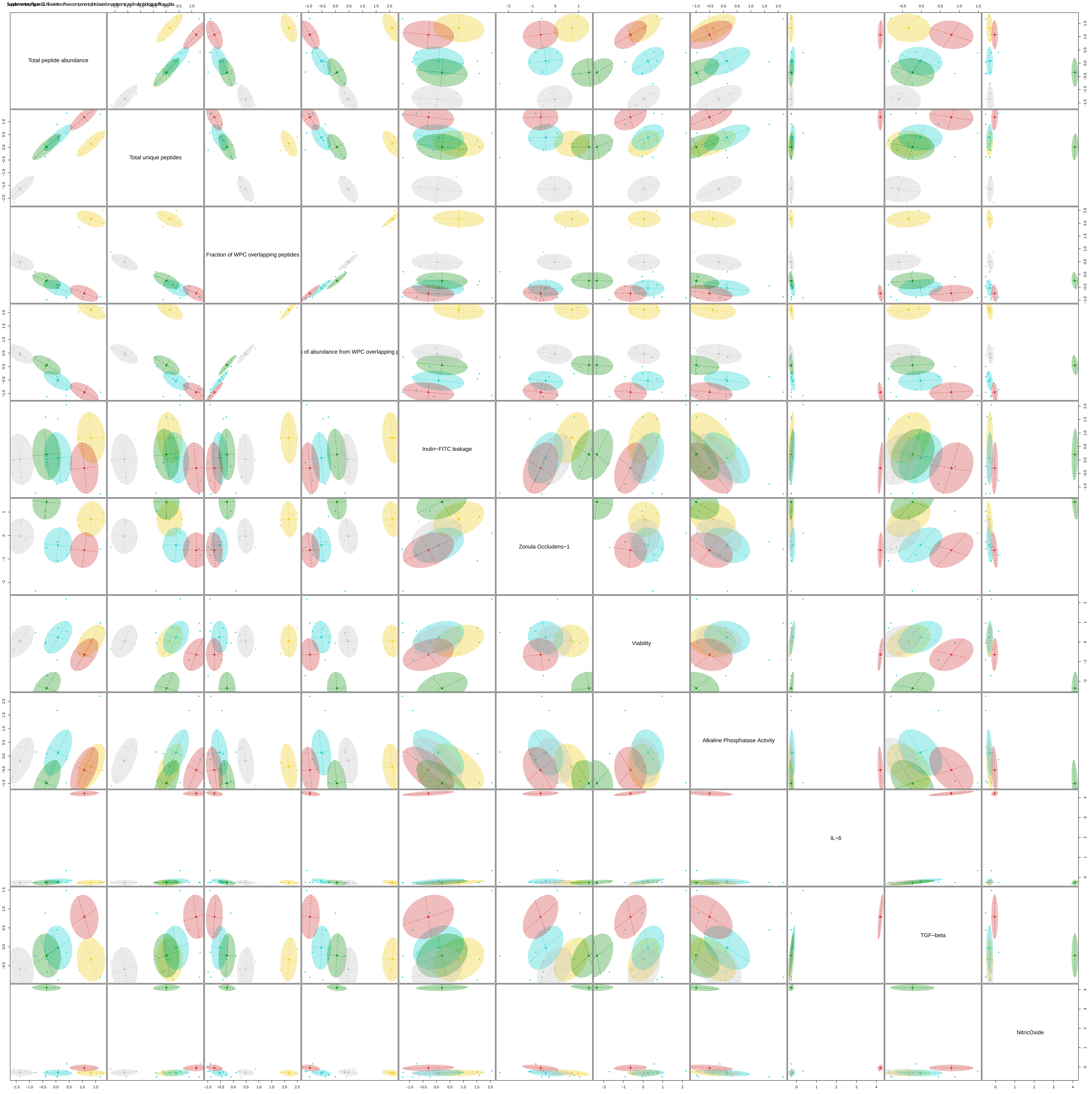

**Supplementary Figure 13.** Semi-quantification of ZO-1 as marker of barrier integrity.

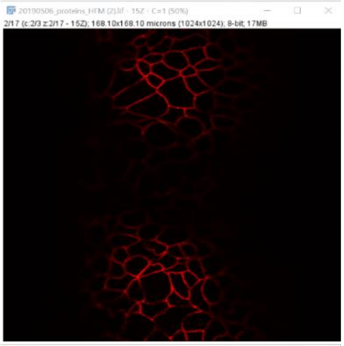

A single image of a bioengineered intestinal tubule Z-stack of the Zonula Occludens-1 network

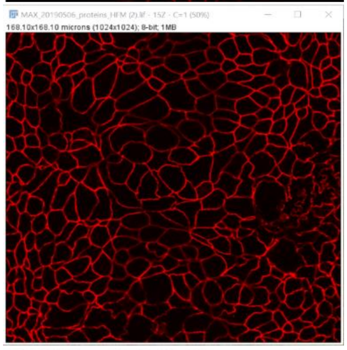

Maximum intensity projection (MIP) created from the Z-stack. Noise level was determined with the plot profile function

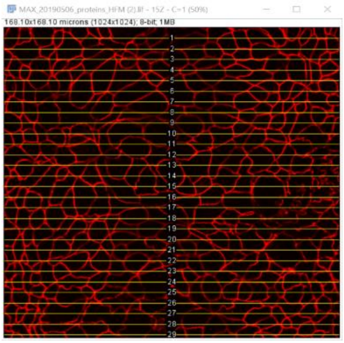

A custom made ROI SET is applied over the image. The ROI SET consists of 29 horizontal lines evenly distributed over the full image

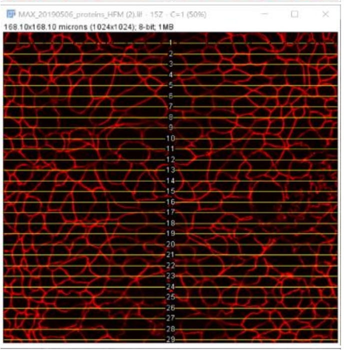

Peakfinder plugin is activated and tolerance is set to 2x noise level. First line is selected with the ROI manager and peakfinder displays a handle for each intersect on the line

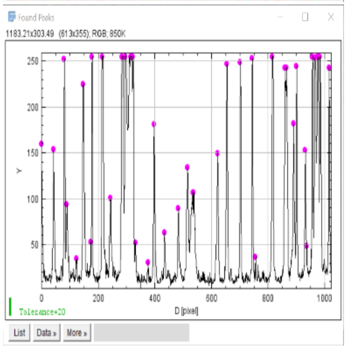

Peakfinder creates a graph displaying peaks for each intersect

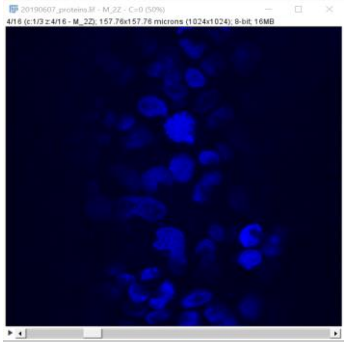

A single image of a bioengineered intestinal tubule Z-stack of the nuclei

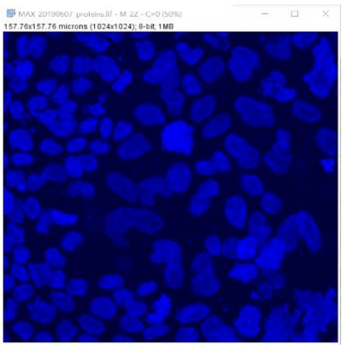

Maximum intensity projection (MIP) created from the Z-stack. This image is then used for counting nuclei

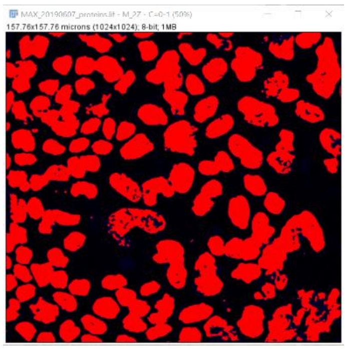

The threshold is determined and applied using Otsu method

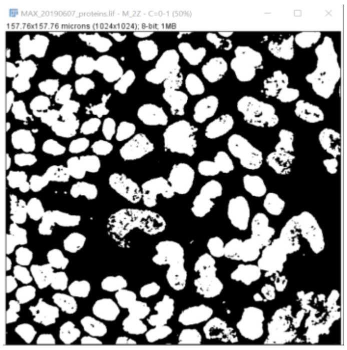

Imaged including threshold limit is converted to a binary image and the watershed function is applied

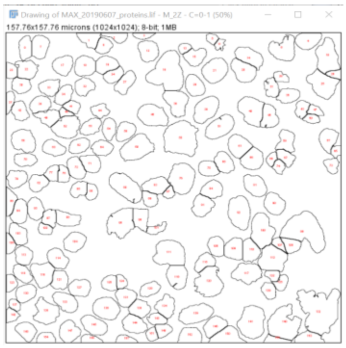

The nuclei are counted with the analyse particle function

**Supplementary Figure 14.** Schematic image of the 3-dimensional printed chamber including dimensions in mm.

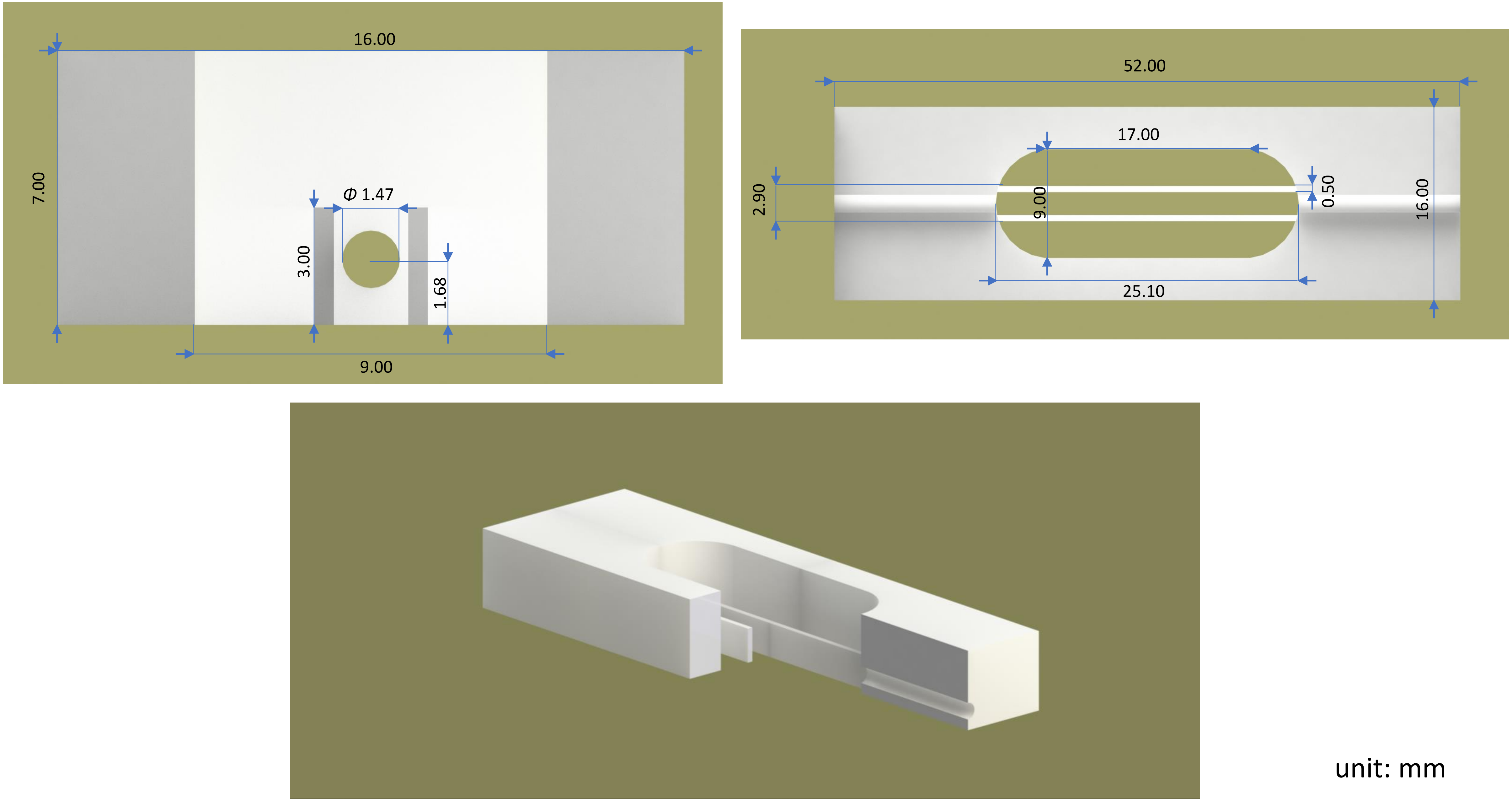
